# Identification of Late Pleistocene and Holocene fossil lizards from Hall’s Cave and a primer on morphological variation in North American lizard skulls

**DOI:** 10.1101/2023.07.25.549934

**Authors:** David T. Ledesma, Simon G. Scarpetta, John J. Jacisin, Antonio Meza, Melissa E. Kemp

## Abstract

Fossil identification practices have a profound effect on our interpretation of the past because these identifications form the basis for downstream analyses. Therefore, well-supported fossil identifications are paramount for examining the impact of past environmental changes on populations and communities. Here we apply an apomorphic identification framework in a case study identifying fossil lizard remains from Hall’s Cave, a late Quaternary fossil site located in Central Texas, USA. We present images and descriptions of a broad comparative sample of North American lizard cranial elements and compile new and previously reported apomorphic characters for identifying fossil lizards. Our fossil identifications from Hall’s Cave resulted in a minimum of 11 lizard taxa, including five lizard taxa previously unknown from the site. Most of the identified fossil lizard taxa inhabit the area around Hall’s Cave today, but we reinforce the presence of an extirpated species complex of horned lizard. A main goal of this work is to establish a procedure for making well-supported fossil lizard identifications across North America. The data from this study will assist researchers endeavoring to identify fossil lizards, increasing the potential for novel discoveries related to North American lizards and facilitating more holistic views of ancient faunal assemblages.

## Introduction

Fossils are valuable for providing a glimpse into past life on earth, and fossil data have played critical roles in understanding range shifts (Vasilyan and Bukhsianidze 2020), extinctions (Bochaton et al. 2019), and diversification during critical junctures in earth history (Smith et al. 2022). Paleontological data also lend a long-term perspective on the responses of taxa and ancient communities to environmental changes and have become increasingly important in conservation (Dietl et al. 2015). One of the first and most critical steps for incorporating fossils into ecological and evolutionary analyses is establishing fossil identifications. It is important to recognize that fossil identification practices have a profound effect on our interpretation of the past since these identifications form the basis for downstream analyses. Well-supported fossil identifications are paramount for examining past populations and communities and have been used to infer the impact of past environmental changes on North American Quaternary herpetofauna (non-avian reptiles and amphibians).

North American fossil herpetofaunas from the Quaternary period (2.56 Ma -present) were the subjects of numerous paleontological investigations during the 20^th^ century (see Holman 1995 and references therein). The cumulative findings of these investigators led to the formation of a hypothesis predicting relative taxonomic and biogeographic stability in North American herpetofauna throughout the Quaternary (Auffenberg and Milstead 1965), specifically during the last 1.8 million years (Holman 1995). This hypothesis was born out of the fact that paleoherpetologists who published on Quaternary fossils reported few extinct taxa, noted few to no speciation events, and found that identified species from fossil sites were generally congruent with species found in the area today (Brewer 1985). It was later argued, however, that this stability hypothesis was predicated on biased fossil identification practices that resulted in a circular argument for stability (Bell et al. 2010). Those biases include the historical practice of identifying Quaternary fossil herpetofauna using comparative extant specimens sourced from the immediate vicinity of the fossil deposit or within nearby circumscribed geopolitical or geographical regions. Using these geographically limited comparative datasets, fossils were commonly identified to the species level, sometimes based on ostensible vagueries (Bell et al. 2010). These practices effectively predetermined the taxonomic and geographic stability of Quaternary herpetofauna fossil assemblages (Bell et al. 2010). To address the circularity of the stability hypothesis, Bell et al. (2010) advocated for the use of apomorphies–evolutionary derived features–for making fossil identifications. Importantly, apomorphic identifications, when paired with an expansive comparative dataset, are less sensitive to geographic biases and provide a more objective and replicable system for identifying fossils.

Here we employ an apomorphy based identification framework to identify fossil lizard remains from a late Quaternary fossil site in Central Texas. Our study site, Hall’s Cave, is an exceptional study system because there is an abundance of well-preserved fossil material, including a sizable accumulation of fossil herpetofauna that would benefit from rigorous examination (Toomey 1993). Hall’s Cave contains the most complete late Quaternary stratigraphic sequence in Texas, spanning the last 20,000 years (Toomey 1993; Cook et al. 2003), and is located near the southern edge of the Edwards Plateau, an uplifted region occupying much of central Texas. Hall’s Cave was first excavated beginning in the late 1960s (then under the name Klein Cave; Roth 1972) and excavations continued in the late 80’s-early 90’s (Toomey 1993) as well as in recent years (Seersholm et al. 2020; Waters et al. 2021). Identification of fossil remains from these excavations largely focused on mammalian taxa (Toomey 1993; Smith et al. 2016) and plant microfossils (Cordova and Johnson 2019), but some herpetofauna were identified (Parmley 1988; Toomey 1993). Previous research on Hall’s Cave has also included bulk bone ancient DNA metabarcoding analyses, yet lizard ancient DNA was not recovered and amplified (Seersholm et al. 2020). Previous investigations of the fossil lizards from Hall’s Cave reported at least six different lizard taxa (Parmley 1988; Toomey 1993). Most of these identifications lacked a corresponding discussion of apomorphic morphological features supporting their taxonomic assignment. Here we reassess previously examined as well as new fossil lizard material from Hall’s Cave using an apomorphic identification framework.

In addition to identifying fossil lizards from Hall’s Cave, we also sought to establish a procedure for making well-supported fossil lizard identifications, particularly for North American lizards. We therefore compiled previously reported apomorphic characters as well as new potential apomorphies that can be used to identify fossil lizard remains from the Quaternary of North America. Furthermore, the Neogene and Quaternary fossil record for lizards is largely composed of tooth bearing elements from the upper and lower jaws (Holman 1995; Bell et al. 2010), yet it has been shown that there remain previously undiscovered apomorphies on other skeletal elements that are useful for fossil identification (Smith 2011). The authors of Bell et al. (2010) posited that the abundance of fossil lizard tooth bearing elements from the upper and lower jaws is due to a historical emphasis placed on mammal fossil collections and a general unfamiliarity of the larger lizard skeletal system. Therefore, for fossil skeletal elements examined here (Fig. 1), we also include images from a diverse set of North American lizard taxa to showcase morphological diversity and facilitate the recognition and identification of fossil lizard remains by paleontologists who do not specialize in lizard osteology.

**Figure 1.**
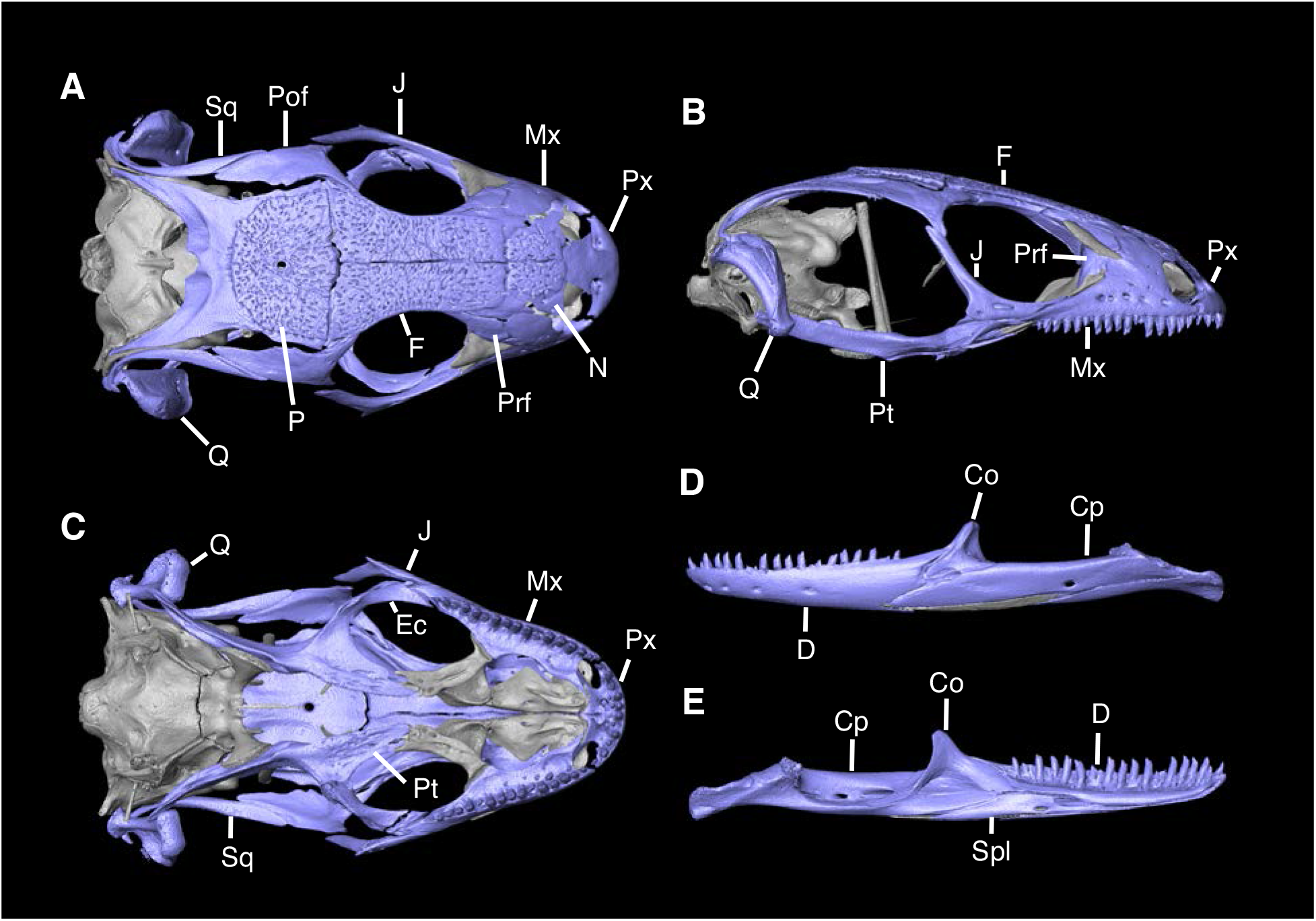
Skull of *Diploglossus bilobatus* TNHC 31933. A. Dorsal view; B. Lateral view; C. Ventral view; D. Mandible lateral view; E. Mandible medial view. Abbreviations: Co, Coronoid; Cp, Compound bone; D, Dentary; Ec, Ectopterygoid; F, Frontal; J, Jugal; Mx, Maxilla; N, Nasal; P, Parietal; Pof, Postorbitofrontal; Prf, Prefrontal; Pt, Pterygoid; Px, Premaxilla; Spl, Splenial; Sq, Squamosal; Q, Quadrate.

## Methods

Fossils in this study were previously excavated from Hall’s Cave and are accessioned at the University of Texas Vertebrate Paleontology Collections (TxVP), locality 41229. We largely restricted our osteological comparative material to North American (NA) lizard taxa (see table 1 in Scarpetta et al. 2020) including those that today live on or north of the Isthmus of Panama, as well as those that inhabit Caribbean islands. Our comparative dataset was chiefly based on dry skeletal specimens; however, we also examined skeletons from specimens scanned using high-resolution computed tomography (CT). We aimed to get at least one specimen for every North American lizard family for our comparative dataset and for some families (e.g., Phrynosomatidae) we were able to sample all or nearly all genera. However, it was difficult to obtain comparative material for a few NA lizard families (e.g., Gymnophthalmidae) and so we instead relied on morphological evidence from the published literature. The full list of comparative specimens can be found in Table S1. Our phylogenetic framework follows that of Burbrink et al. (2020) for higher squamate relationships.

**Table 1.** Identified fossil lizard taxa represented in Hall’s Cave from this study compared to previous studies.

| Family | Previously identified lizard taxa by Parmley (1988) | Previously identified lizard taxa by Toomey (1993) | Updated faunal list from this study |
| --- | --- | --- | --- |
| Crotaphytidae |  | *Crotaphytus* sp. | Crotaphytidae |
| Phrynosomatidae |  | unid. Sceloporinae | Sceloporinae |
|  |  |  | *Urosaurus* |
|  |  |  | Callisaurini |
|  | *Phrynosoma* cf. *P. cornutum* | *Phrynosoma cornutum* | *Phrynosoma* cf. *P. cornutum* |
|  |  | *Phrynosoma douglasii* | *Phrynosoma douglasii* species complex |
|  |  | *Phrynosoma modestum?* | *Phrynosoma* sp. and *Phrynosoma douglasii* species complex |
| Anguidae | *Gerrhonotus liocephalus* |  | *Gerrhonotus* |
|  |  | *Ophisaurus attenuatus* | *Ophisaurus* |
| Scincidae |  |  | *Scincella* |
|  |  |  | Scincinae |
| Teiidae |  |  | Teiidae |

We employed an apomorphy-based fossil identification framework using previously published apomorphies, as well as apomorphies we report here. Because our comparative dataset is largely restricted to NA lizard taxa, our fossil identifications are also geographically restricted on the continental scale. We were not able to examine every NA lizard species or clade and therefore take a conservative approach for fossil identifications. Apomorphies taken from the literature are listed in Table S2. Newly reported apomorphies can be found in the text and are based on our NA restricted comparative dataset. We note that newly reported apomorphies must be examined in non-NA taxa before being used in a global scale apomorphic diagnosis. Anatomical terminology follows that of Evans (2008) unless otherwise noted and many of the anatomical structures referenced in the text are labeled in Figure 2. We were able to base our fossil descriptions on one or a few exemplar fossil specimens because we observed no substantive differences between exemplar and other referred specimens, unless otherwise noted.

**Figure 2.**
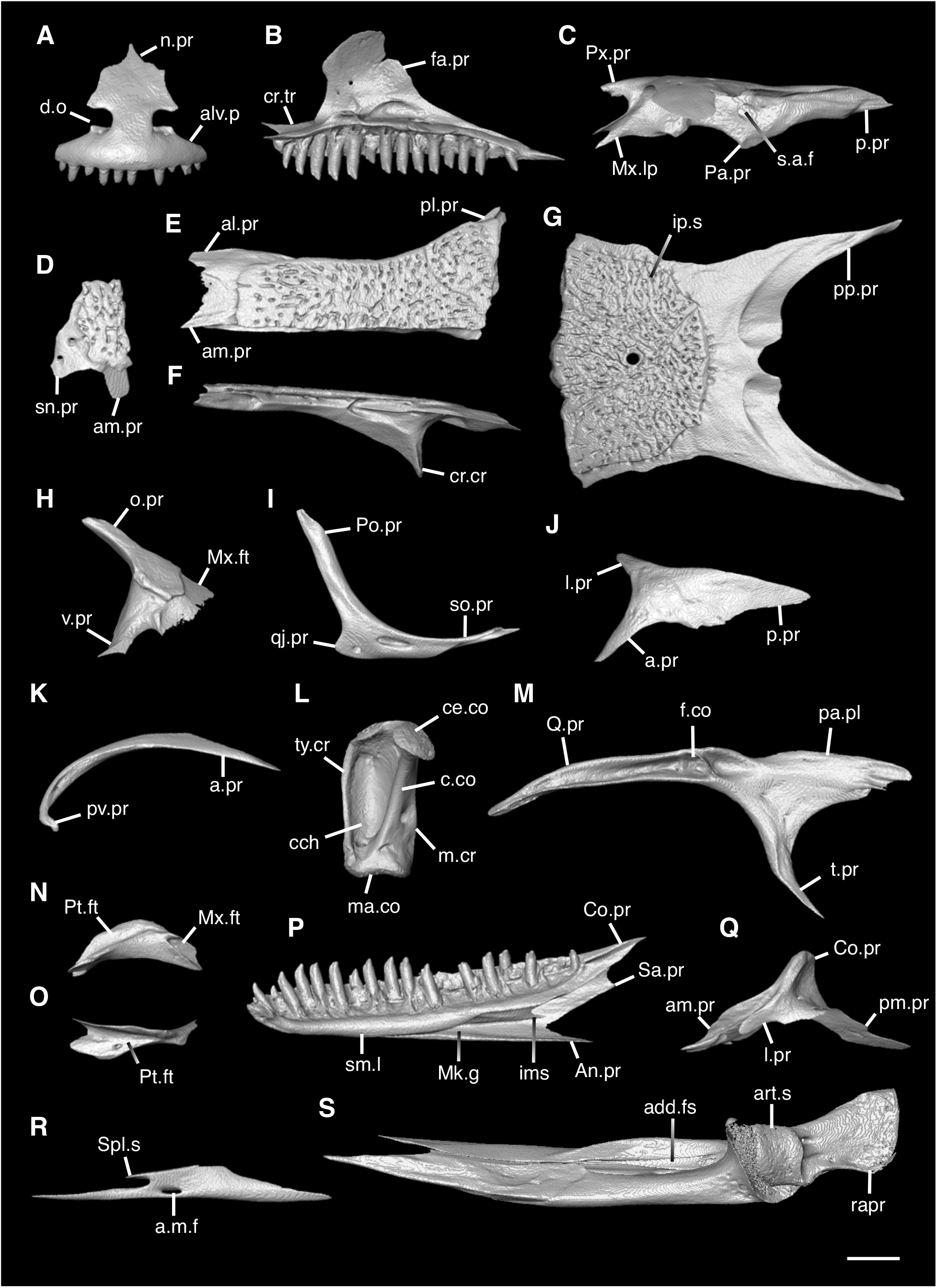
Cranial elements of *Diploglossus bilobatus* TNHC 31933 with relevant anatomical structures labeled”. **A**. Premaxilla in anterior view; **B-C**. Right maxilla in medial and dorsal views; **D**. Left nasal dorsal view; **E-F**. Right frontal dorsal and lateral views; **G**. Parietal in dorsal view; **H**. Right prefrontal lateral view; **I**. Right jugal lateral view; **J**. Right postorbitofrontal dorsal view; **K**. Right squamosal lateral view; **L**. Left quadrate posterior view; **M**. Right pterygoid dorsal view; **N-O**. Right ectopterygoid ventral and lateral views; **P**. Right dentary medial view; **Q**. Left coronoid lateral view; **R**. Right splenial medial view; **S**. Left compound bone dorsal view. Scale bar = 1 mm. **Abbreviations**: add.fo, adductor fossa; al.pr, anterolateral process; alv.p, alveolar plate; a.m.f, anterior mylohyoid foramen; am.pr, anteromedial process; An.pr, Angular process; a.pr, anterior process; art.s, articular surface; cch, conch; c.co, central column; ce.co, cephalic condyle; Co.pr, Coronoid process; cr.cr, cristae cranii; cr.tr, crista transversalis; d.o, dorsal ossification; fa.pr, facial process; f.co, fossa columella; ims, intramandibular septum; ip.s, interparietal shield; l.pr, lateral process; ma.co, mandibular condyle; m.cr, medial crest; Mk.g, Meckelian groove; Mx.ft, maxilla facet; Mx.lp, maxillary lappet; n.pr, nasal process; o.pr, orbital process; pa.pl, palatal plate; Pa.pr, palatine process; pl.pr, posterolateral process; pm.pr, posteromedial process; pp.pr, postparietal process; p.pr, posterior process; Pt.ft, Pterygoid facet; pv.pr, posteroventral process; Px.pr, premaxillary process; qj.pr, quadratojugal process; Q.pr, Quadrate process; rapr, retroarticular process; s.a.f, superior alveolar foramen; Sa.pr, surangular process; sm.l, suprameckelian lip; sn.pr, supranarial process; so.pr, suborbital process; Spl.s, Splenial spine; t.pr, transverse process; ty.cr, tympanic crest; v.pr, ventral process.

## Systematic Paleontology

**Iguania Cuvier, 1817**

**Pleurodonta Cope, 1864**

**Referred specimens**: See Table S3.

### Nasal

#### Description

TxVP 41229-25562 is a right nasal (Fig. 3). There is a long anteromedial process with a lateral articulation facet for the premaxilla. There is a shorter anterolateral process. The posterior end is narrow compared to the anterior end. The ventral surface is concave, and there are two foramina that pierce the bone vertically.

**Figure 3.**
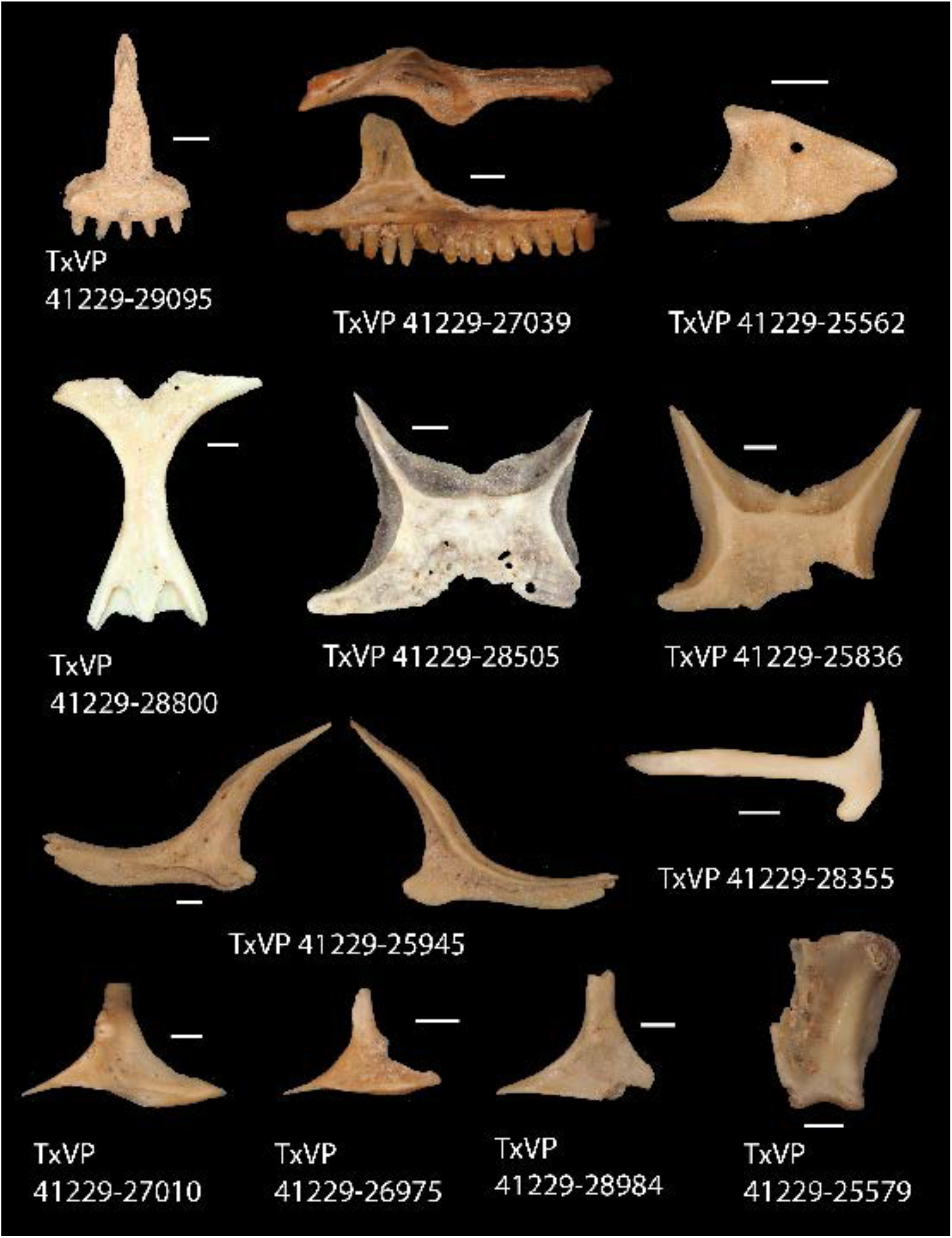
Fossil pleurodontans. TxVP 41229-29095 Anterior view of premaxilla; TxVP 41229-27039 Dorsal and medial view of right maxilla; TxVP 41229-25562 Dorsal view of nasal; TxVP 41229-28800 Dorsal view of frontal; TxVP 41229-28505 Dorsal view of parietal; TxVP 41229-25836 Dorsal view of parietal; TxVP 41229-25945 Lateral and medial view of left jugal; TxVP 41229-28355 Lateral view of left squamosal; TxVP 41229-27010 Dorsolateral view of left postorbital; TxVP 41229-26975 Dorsolateral view of left postorbital; TxVP 41229-28984 Dorsolateral view of left postorbital; TxVP 41229-25579 Posterior view of left quadrate. Scale bars = 1 mm.

#### Identification

The fossil shares with pleurodontans a supranarial process (Gauthier et al. 2012), although the process is much longer in some pleurodontan taxa (e.g., *Dipsosaurus dorsalis*), and a nasal that gradually narrows posteriorly. A supranarial process is also present in some non-iguanians but it is usually small except for *Diploglossus bilobatus* (Fig. 4). The fossil nasal distinctly narrows posteriorly similar to that observed in many examined North American lizards pleurodontans (Fig. 5). The fossil nasal differs from non-pleurodontan North American lizards, except for *Anelytropsis papillosus*, in rapidly tapering posteriorly, resulting in a posterior end that is much narrower compared to the anterior end. *Anelytropsis papillosus* differs from the fossil in lacking a supranarial process. Furthermore, many non-pleurodontan North American lizards, and anguimorphs in particular, differ from the fossil in often having a distinct rugose dorsal surface of the nasal.

**Figure 4.**
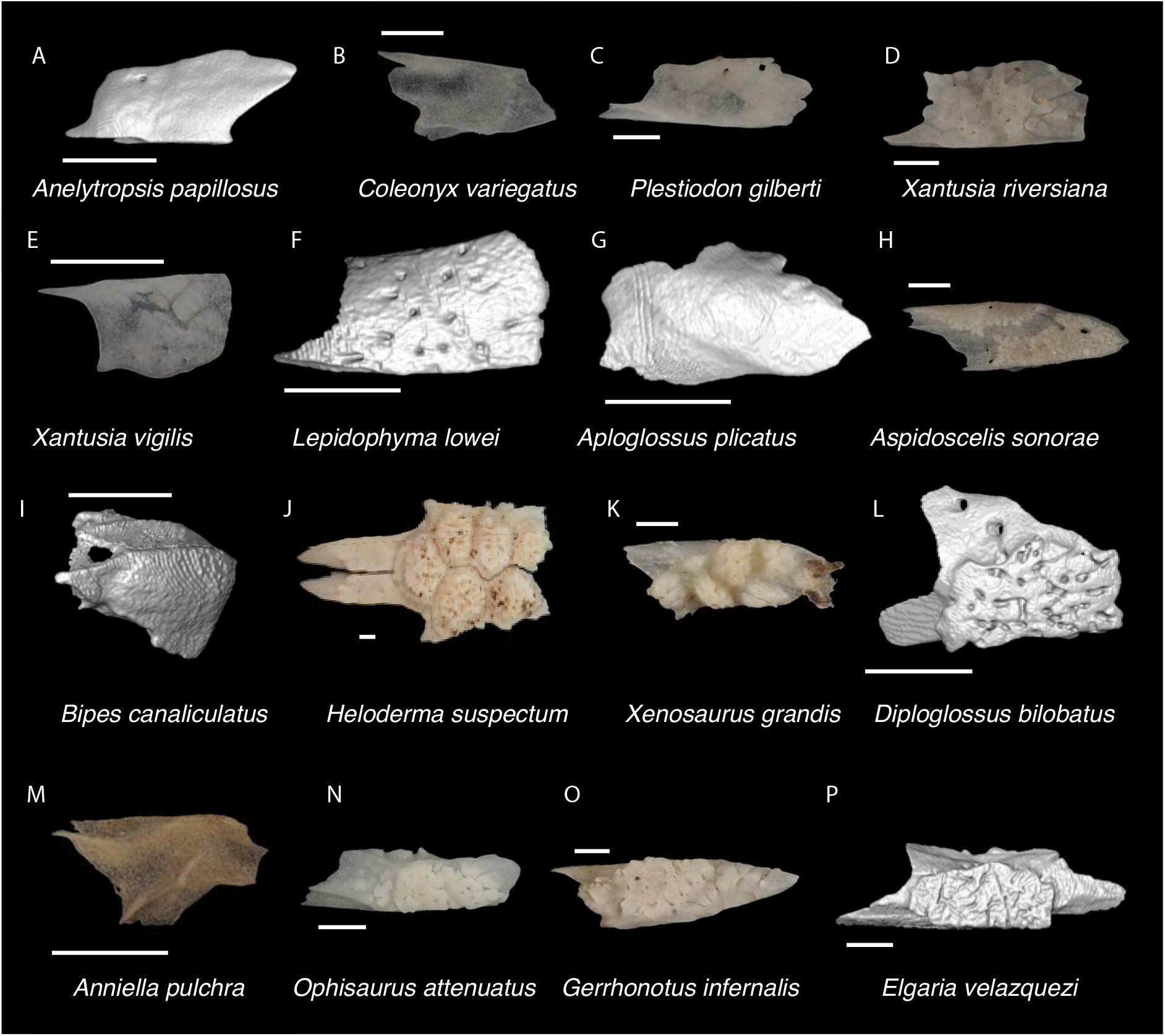
Non-Pleurodontan Nasals. Nasals in dorsal view– **A**. *Anelytropsis papillosus* UF 86708; **B**. *Coleonyx variegatus* TxVP M-12109; **C**. *Plestiodon gilberti* TxVP M-8587; **D**. *Xantusia riversiana* TxVP M-8505; **E**. *Xantusia virigatus* TxVP M-12130; **F**. *Lepidophyma lowei* LACM 143367; **G**. *Aploglossus plicatus* TNHC 34481; **H**. *Aspidoscelis sonorae* TxVP M-15670; **I**. *Bipes canaliculatus* CAS 134753; **J**. *Heloderma suspectum* TxVP M-9001; **K**. *Xenosaurus grandis* TxVP M-8960; **L**. *Diploglossus bilobatus* TNHC 31933; **M**. *Anniella pulchra* TxVP M-8678; **N**. *Ophisaurus attenuatus* TxVP M-8979; **O**. *Gerrhonotus infernalis* TxVP M-13441; **P**. *Elgaria velazquezi* SDNHM 68677. Scale bars = 1 mm.

**Figure 5.**
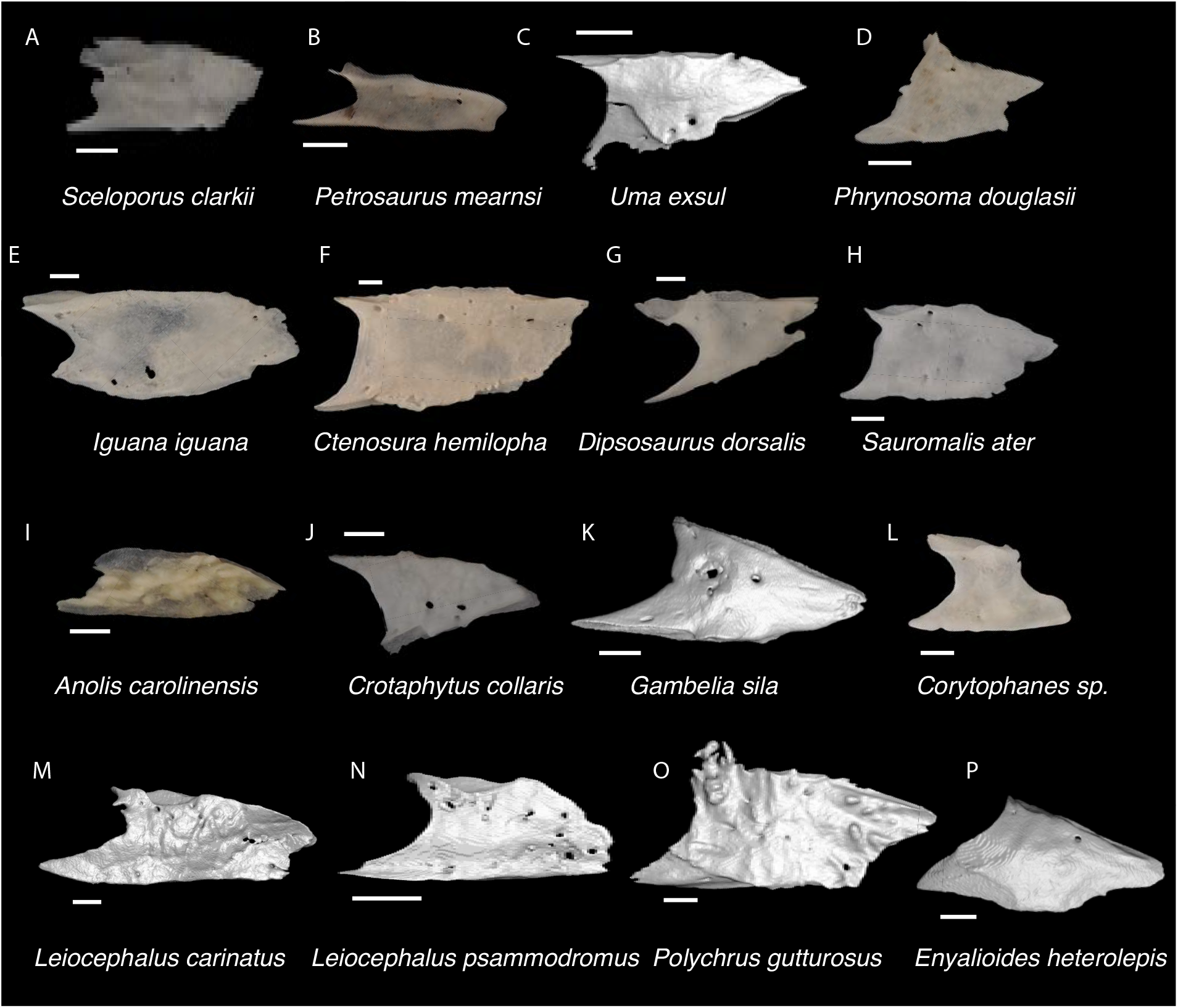
Pleurodontan Nasals. Nasals in dorsal view– **A**. *Sceloporus clarkii* TxVP M-12157; **B**. *Petrosaurus mearnsi* TxVP M-14910; **C**. *Uma exsul* TNHC 30247; **D**. *Phrynosoma douglasii* TxVP M-8526; **E**. *Iguana iguana* TxVP M-8454; **F**. *Ctenosaura hemilopha* TxVP M-8616; **G**. *Dipsosaurus dorsalis* TxVP M-13086; **H**. *Sauromalis ater* TxVP M-11599; **I**. *Anolis carolinensis* TxVP M-9042; **J**. *Crotaphytus collaris* TxVP M-12468; **K**. *Gambelia sila* TNHC 95261; **L**. *Corytophanes* sp. TxVP M-16765; **M**. *Leiocephalus carinatus* TNHC 89274; **N**. *Leiocephalus psammodromus* TNHC 103220; **O**. *Polychrus gutturosus* TNHC 24152; **P**. *Enyalioides heterolepis* UF 68015. Scale bars = 1 mm.

**Crotaphytidae Smith & Brodie, 1982**

**Referred specimens**: See Table S3.

### Premaxilla

#### Description

TxVP 41229-29095 is a premaxilla with five tooth positions filled with widely spaced unicuspid teeth (Fig. 3). It has a relatively flat anterior rostral surface and a long nasal process with distinct nasal articulation facets visible on the anterior and lateral surfaces. There are small foramina just lateral to the base of the nasal process and there are no anterior foramina. There are shallow maxillary facets laterally on the alveolar plate. Posteriorly, the palatal plate is steeply slanted and narrow. The short, rounded incisive process is slightly bilobed.

#### Identification

A fossil premaxilla is assigned to Pleurodonta based on being fused (Estes et al. 1988) with fewer than seven tooth positions (Smith 2009a). Some pleurodontans such as *Anolis*, *Polychrus*, corytophanids, and *Enyaliodes* differ from the fossil in having seven or more tooth positions (Smith 2011; Daza et al. 2012; Fig. 6) and examined specimens of *Anolis* have an incised posterior edge of the palatal process not found in the fossil. Members of Iguanidae differ in often having multicuspid teeth on the premaxilla (de Queiroz 1987). Unicuspid teeth are sometimes found in *Ctenosaura*, *Cyclura* and *Sauromalus* (Scarpetta 2019) but these taxa differ in that the anterior rostral face of the premaxilla in *Ctenosaura* is distinctly rounded (Scarpetta 2019) and the nasal process of *Ctenosaura*, *Cyclura*, and *Sauromalus* curves far posteriorly (Scarpetta 2019; Smith 2011). The nasals were reported to overlap the nasal process of the premaxilla in *Crotaphytus collaris* (Daza et al. 2012), and the fossil shares with many examined *Crotaphytus* distinct facets visible on the nasal process in anterior view. The nasals of phrynosomatids do not overlap the nasal process of the premaxilla (Daza et al. 2012) and no examined phrynosomatids have distinct nasal facets on the anterior face of the nasal process. The fossil premaxilla differs from corytophanids and phrynosomatines in lacking anterior premaxillary foramina, which are also absent in most sceloporines and crotaphytids (Smith 2009a; Scarpetta 2019). We found that *Crotaphytus collaris* TxVP M-9255 has one small anterior premaxilla foramen. *Leiocephalus* also lacks anterior premaxillary foramina (Smith 2009a). Examined *Leiocephalus* differ from the fossil in having at least seven tooth positions (see also Bochaton et al. 2021), but *Leiocephalus personatus* was reported to have six (Smith 2009a). Additionally, the fossil and many examined *Crotaphytus collaris* have a palatal shelf that does not extend as far posteriorly compared to examined *Sceloporus* and *Leiocephalus*. On this basis, the fossil was assigned to Crotaphytidae.

**Figure 6.**
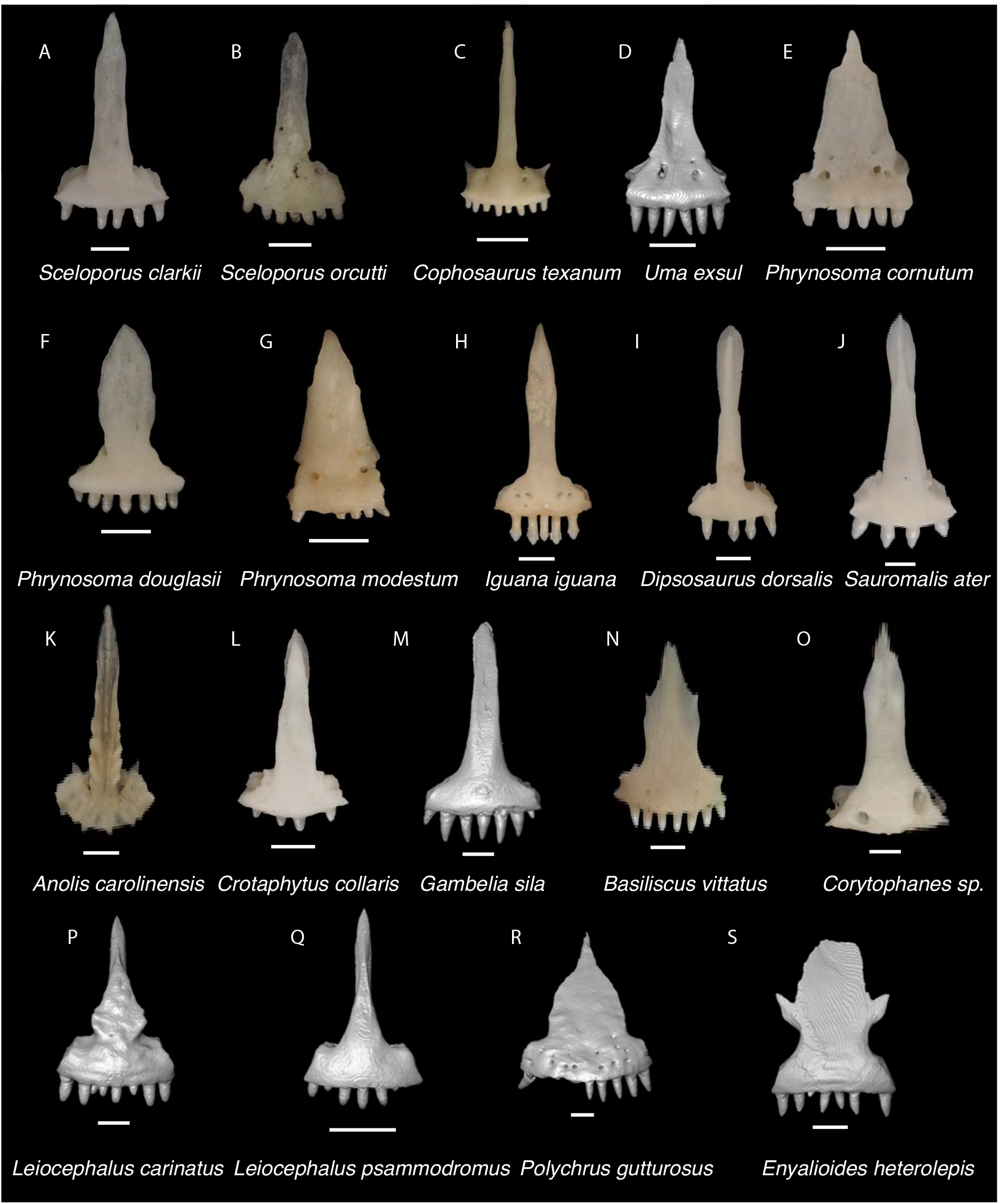
Pleurodontan Premaxillae. Premaxillae in anterior view– **A**. *Sceloporus clarkii* TxVP M-12157; **B**. *Sceloporus orcutti* TxVP M-12155; **C**. *Cophosaurus texanum* TxVP M-8527; **D**. *Uma exsul* TNHC 30247; **E**. *Phrynosoma cornutum* TxVP M-9621; **F**. *Phrynosoma douglasii* TxVP M-8526; **G**. *Phrynosoma modestum* TNHC 95921; **H**. *Iguana iguana* TxVP M-13054; **I**. *Dipsosaurus dorsalis* TxVP M-13086; **J**. *Sauromalis ater* TxVP M-11599; **K**. *Anolis carolinensis* TxVP M-9042; **L**. *Crotaphytus collaris* TxVP M-12468; **M**. *Gambelia sila* TNHC 95261; **N**. *Basiliscus vittatus* TxVP M-8556; **O**. *Corytophanes* sp. TxVP M-16765; **P**. *Leiocephalus carinatus* TNHC 89274; **Q**. *Leiocephalus psammodromus* TNHC 103220; **R**. *Polychrus gutturosus* TNHC 24152; **S**. *Enyalioides heterolepis* UF 68015. Scale bars = 1 mm.

### Maxilla

#### Description

TxVP 41229-27039 serves as the basis for our description (Fig. 3). TxVP 41229-27039 is a right maxilla with 15 tooth positions. The facial process is tall and narrow, and gently curves anteromedially where it diminishes and merges with the crista transversalis. There is an anterior inferior alveolar foramen anterior to the facial process, and an elongate opening for the subnarial artery. There is a sub-triangular, symmetrical palatine process and an anteroposteriorly elongate depression (gutter) on the palatal shelf housing the superior alveolar nerve and maxillary artery. The lateral wall of the posterior orbital process is short. The posterior orbital process is narrow, and elongated with a deep, narrow jugal groove. Teeth are tricuspid and the distal teeth are widened such that the base of the tooth is substantially wider than the crown. There are five nutrient foramina along the lateral surface of the bone, and a few foramina scattered on the lateral face of the facial process.

#### Identification

Maxillae were referred to Pleurodonta based on the presence of an elongate depression on the palatal shelf encompassing the superior alveolar nerve and maxillary artery (Smith 2009a; Fig. 7). This depression is absent in non-iguanian lizards (Smith 2006). Furthermore, maxillae can be assigned to Pleurodonta based on the presence of pleurodont teeth with tricuspid crowns, and two foramina on the dorsal surface of the premaxillary process (Smith 2009a). Maxillae were identified to Crotaphytidae based on having teeth with bases much wider than the crowns, and a deep jugal groove beginning lateral to the palatine process and running posteriorly along the posterior orbital process (Smith 2009a; Scarpetta 2021). A deep jugal groove is absent in phrynosomatids but is present in other North American pleurodontans, including Corytophanidae, Iguanidae, and Leiocephalidae (Smith 2009a; Scarpetta 2021). Many iguanids differ from crotaphytids in having multicuspid teeth that are flared at the crown (de Queiroz 1987). Moreover, among those taxa, only Crotaphytidae, some members of Corytophanidae, and *Leiocephalus* have a large palatine process (Scarpetta 2021). *Leiocephalus* is precluded because species within that genus have tightly packed teeth with flared crowns without a widened base (Smith 2009a). Corytophanids differ in having a wide posterior orbital process of maxilla, an anteroposteriorly long facial process, a tall lateral edge of the maxilla posterior to the facial process, and lacking teeth with bases much wider than the crowns (Scarpetta 2021).

**Figure 7.**
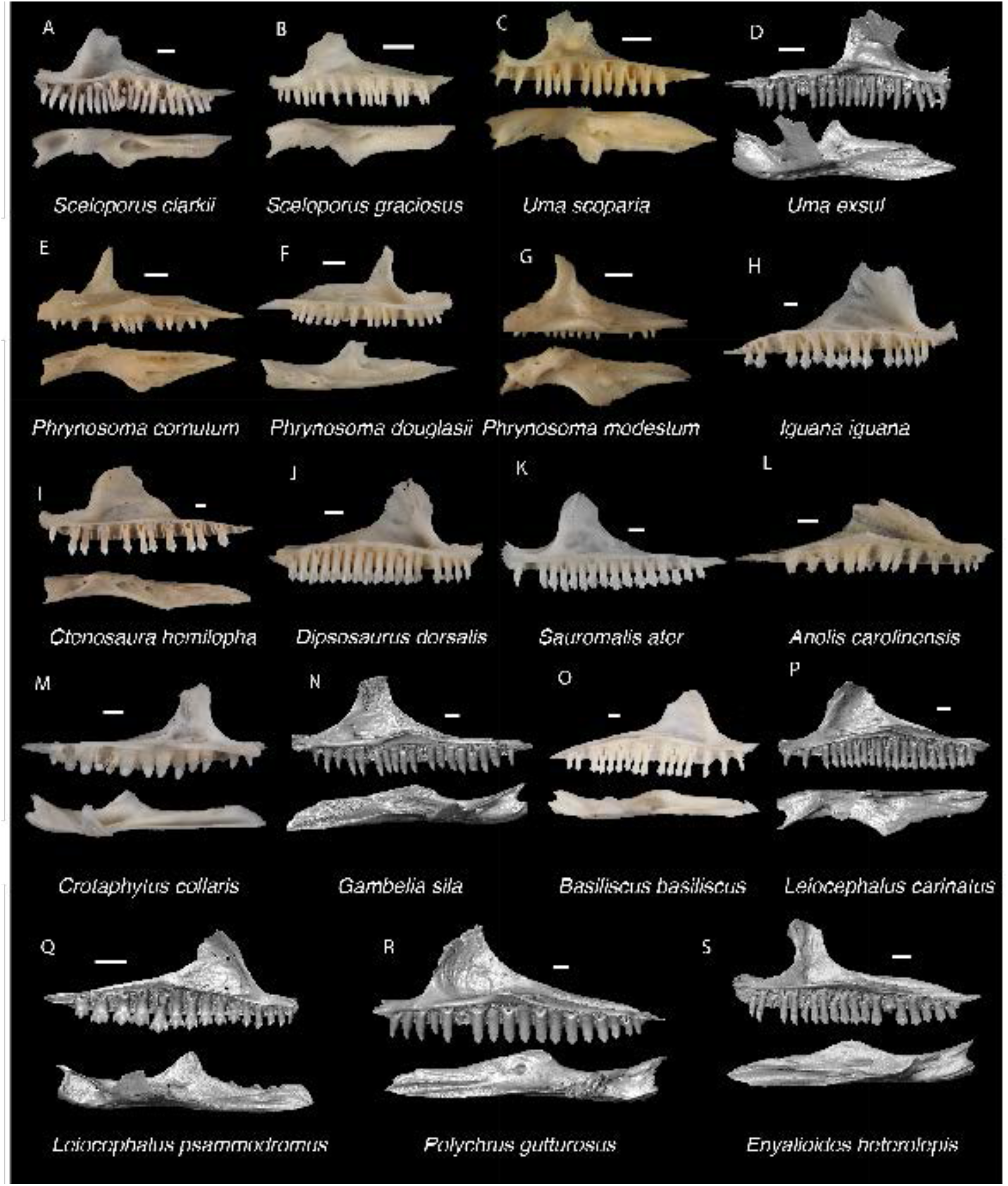
Pleurodontan Maxillae. Maxillae in medial and dorsal views– **A**. *Sceloporus clarkii* TxVP M-12157; **B**. *Sceloporus graciosus* TxVP M-14879; **C**. *Uma scoparia* TxVP M-8529; **D**. *Uma exsul* TNHC 30247; **E**. *Phrynosoma cornutum* TxVP M-6405; **F**. *Phrynosoma douglasii* TxVP M-8526; **G**. *Phrynosoma modestum* TNHC 95921; **H**. *Iguana iguana* TxVP M-8454; **I**. *Ctenosaura hemilopha* TxVP M-8616; **J**. *Dipsosaurus dorsalis* TxVP M-13086; **K**. *Sauromalis ater* TxVP M-11599; **L**. *Anolis carolinensis* TxVP M-9042; **M**. *Crotaphytus collaris* TxVP M-12468; **N**. *Gambelia sila* TNHC 95261; **O**. *Basiliscus basiliscus* TxVP M-11907; **P**. *Leiocephalus carinatus* TNHC 89274; **Q**. *Leiocephalus psammodromus* TNHC 103220; **R**. *Polychrus gutturosus* TNHC 24152; **S**. *Enyalioides heterolepis* UF 68015. Scale bars = 1 mm.

### Frontal

#### Description

TxVP 41229-28800 lacks only the end of the right posterolateral corner (Fig. 3). The bone has a narrow, waisted interorbital region with posterolateral margins that flare laterally. There is a large, deep midline notch on the posterior edge. The dorsal surface has a slightly rugose texture. The anterior end is triradiate with large, well defined nasal facets separated by an anteromedial process. There are distinct prefrontal facets on the anterolateral ends of the bone. TxVP 41229-28800 is slightly concave ventrally and has short cristae cranii that approach one another in the interorbital region and bound an indistinct groove for attachment of the solium supraseptale. The cristae cranii diverge posteriorly and extend along the lateral margins of the ventral surface.

#### Identification

TxVP 41229-28800 is assigned to Pleurodonta based on the presence of a fused frontal with reduced cristae cranii (Estes et al. 1988). Teiids, gymnophthalmoids, geckos, some anguids, and some skinks also have a fused frontal (Greer 1970; Estes et al 1988). Teiids were reported to differ from iguanians in lacking strongly constricted interorbital margins of the frontal (Estes et al. 1988); however, we observed some specimens of *Aspidoscelis* that have similarly constricted interorbital margins. Examined *Aspidoscelis*, *Mabuya*, and *Scincella* differ from the fossil and from other pleurodontans in having an interorbital edge that weakly curves posterolaterally. Gymnophthalmoids possess frontal tabs not found in iguanians (Estes et al. 1988). Geckos differ from iguanians in having the crista cranii meet to form an enclosed olfactory canal (Kluge 1967; Estes et al. 1988). Anguimorphs usually have co-ossified osteoderms and well-developed cristae cranii (Evans 2008). The fossil and Crotaphytidae share a deep, narrow notch in the posterior edge of the frontal for the parietal foramen, constricted interorbital margins of the frontal, and a distinctly anteromedial process on the anterior end of the frontal (Fig. 8). The frontal of *Anolis* generally does not contribute to the parietal/parietal foramen (Etheridge 1959; Smith 2011) and in *Anolis* and examined *Polychrus* the interorbital margins are considerably widened (Pregill 1988; Smith 2011). The interorbital margins of the frontal are also considerably widened in *Ctenosaurus similis* (Smith 2011), *Iguana iguana*, *Iguana delicatissima*, and some *Liocephalus* (Pregill 1992); however, this morphology varies ontogenetically (de Queiroz 1987; Bochaton et al. 2016a). *Anolis*, *Polychrus*, *Enyalioides heterolepis*, and some *Liocephalus* differ from crotaphytids and the fossil in having a distinct rugose texture on the dorsal surface of the frontal (Etheridge and de Queiroz 1988; Pregill 1992). The parietal foramen is located entirely within the frontal in *Basiliscus*, *Corytophanes*, some *Laemanctus*, some *Sauromalus*, and in many *Dipsosaurus dorsalis* (de Queiroz 1987). Among examined specimens of *Crotaphytus collaris*, the position of the parietal foramen is variably at the frontoparietal suture, largely within the frontal as demarcated by a deep, narrow notch in the posterior edge, or completely within the frontal (*C*. *collaris* TxVP M-8615). One examined specimen of *Gambelia* (*G. wislizenii* TxVP M-9974) also has the parietal foramen largely within the frontal. Among phrynosomatids, the parietal foramen is either located at the frontoparietal suture (e.g., some *Phrynosoma*, Presch 1969) or largely within the parietal (Etheridge 1964). However, most phrynosomatids lack the deep, narrow notch in the posterior edge of the frontal that is present in crotaphytid specimens. Furthermore, many examined phrynosomatids do not have a distinct anteromedial process on the frontal, and larger species of *Sceloporus* that are similar in size to *Crotaphytus* have relatively wider interorbital margins of the frontal. On this basis, the fossil was assigned to Crotaphytidae.

**Figure 8.**
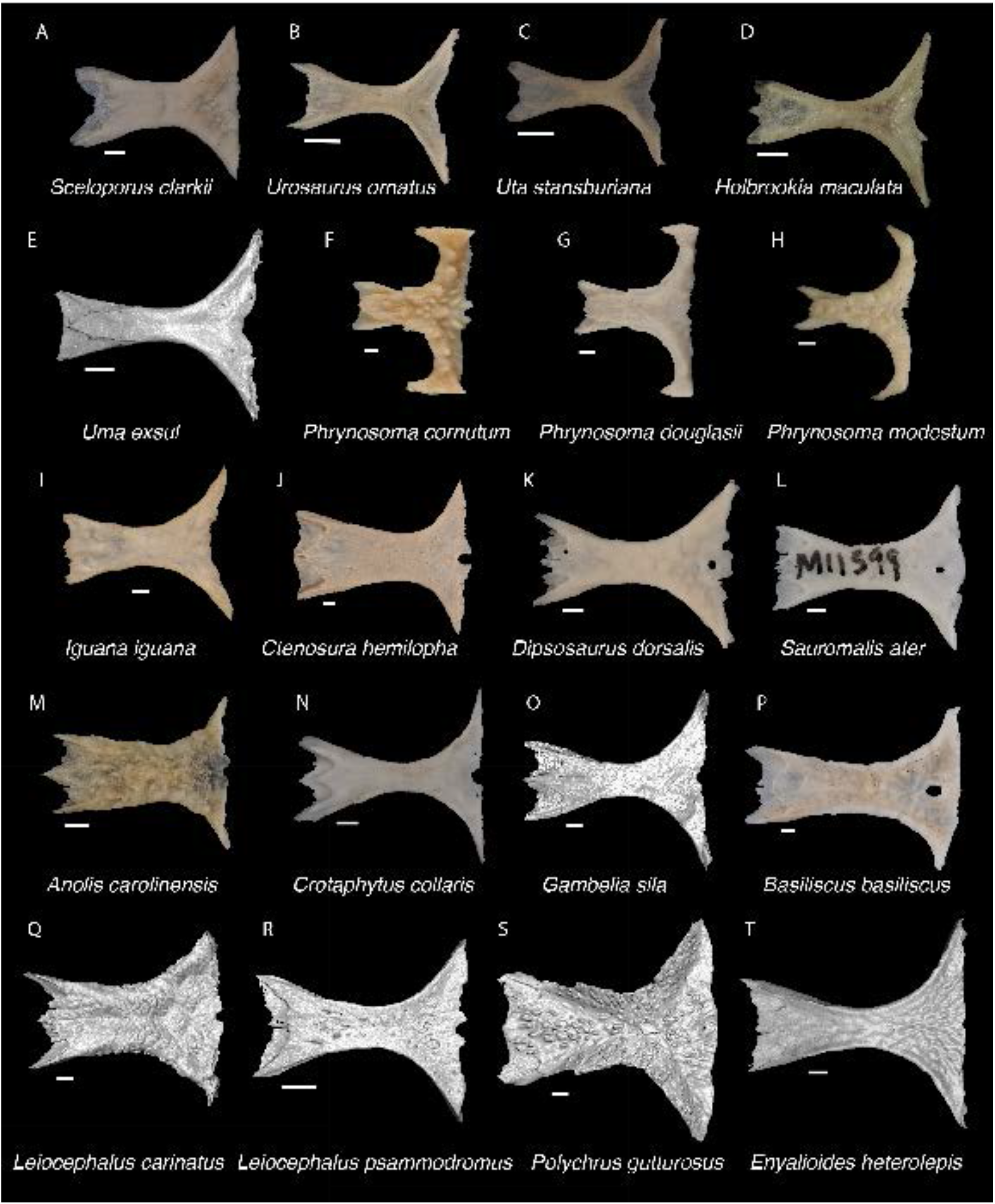
Pleurodontan Frontals. Frontals in dorsal view– **A**. *Sceloporus clarkii* TxVP M-12157; **B**. *Urosaurus ornatus* TxVP M-8638; **C**. *Uta stansburiana* TxVP M-14935; **D**. *Holbrookia maculata* TxVP M-14322; **E**. *Uma exsul* TNHC 30247; **F**. *Phrynosoma cornutum* TxVP M-6405; **G**. *Phrynosoma douglasii* TxVP M-8526; **H**. *Phrynosoma modestum* TNHC 95921; **I**. *Iguana iguana* TxVP M-13054; **J**. *Ctenosura hemilopha* TxVP M-8616; **K**. *Dipsosaurus dorsalis* TxVP M-13086; **L**. *Sauromalis ater* TxVP M-11599; **M**. *Anolis carolinensis* TxVP M-9042; **N**. *Crotaphytus collaris* TxVP M-12468; **O**. *Gambelia sila* TNHC 95261; **P**. *Basiliscus basiliscus* TxVP M-11907; **Q**. *Leiocephalus carinatus* TNHC 89274; **R**. *Leiocephalus psammodromus* TNHC 103220; **S**. *Polychrus gutturosus* TNHC 24152; **T**. *Enyalioides heterolepis* UF 68015. Scale bars = 1 mm.

### Parietal

#### Description

TxVP 41229-25836 is a parietal that is missing the anterolateral corner and the ventral tips of the postparietal processes (Fig. 3). There is a distinct anterolateral process that strongly curves laterally from the parietal table. The adductor crests do not approach one another posteriorly, giving the parietal table a trapezoidal shape. The ventrolateral crests are low without distinct epipterygoid processes and are easily visible in dorsal view. The posterior edge between the postparietal processes is characterized by two distinct depressions (nuchal fossae) separated by a small ridge above a small notch. The postparietal processes have dorsal crests that slant medially. On the ventral surface there are shallow depressions (cerebral vault) divided by a low ridge. There is a deep pit for the processus ascendens just anterior to the posterior edge. TxVP 41229-28505 is similar to TxVP 41229-25836 but has a larger notch at the anterior edge. This notch may be at least partially caused by erosion of the bone indicated by holes in the parietal table.

#### Identification

Parietals are assigned to Pleurodonta based on the presence of a fused parietal (Estes et al. 1988), the absence of co-ossified osteoderms (Estes et al. 1988; Conrad 2008; Evans 2008), the absence of parietal foramen fully enclosed by the parietal (Estes et al. 1988), and the absence of distinct ventrolateral crests (=parietal downgrowths of Estes et al. 1988). The parietal of xantusiids is often unfused (Estes et al. 1988). The parietal in *Xantusia virgilis* is often fused but differs from pleurodontans in being relatively rectangular with distinct ventral flanges extending below the parietal table (Young 1942). Cnemidophorines, gymnophthalmoids, and skinks generally have well-developed, tall ventrolateral crests or projections on the ventral surface of the parietal (Nash 1970; Estes et al. 1988) and anguimorphs often have co-ossified osteoderms on the dorsal surface of the parietal table (Bhullar 2011; Scarpetta et al. 2021). Parietals are assigned to Crotaphytidae based on the presence of a trapezoidal shaped parietal table, a distinct pit for the processus ascendens, medially slanted crests on the postparietal processes, and a supratemporal fossa that is broadly visible in dorsal view. Among pleurodontans, the adductor crests form a “Y” shape of the parietal table in corytophanids, many anoles, and some iguanids (Frost and Etheridge 1989; Smith 2011). The adductor crests form a “V” shape in *Iguana*, *Ctenosaura*, *Cyclura*, some hoplocercids, some *Leiocephalus*, and some anoles (Etheridge 1959; Etheridge and de Queiroz 1988; Pregill 1992) although there is substantial ontogenetic variation in this morphology (Etheridge 1959; de Queiroz 1987; Pregill 1992; Bochaton et al. 2016a). *Polychrus,* some *Anolis*, and some *Liocephalus* differ from crotaphytids in having distinct rugosities on the dorsal surface of the parietal (Etheridge and de Queiroz 1988; Pregill 1992; Bochaton et al. 2021). The pit for the processus ascendens in *Sauromalus* is reduced compared to other iguanids (Smith 2011).

In the remaining NA pleurodontans, there is substantial morphological variation of the parietal (Fig. 9), and it is possible additional variation not captured in our sample may make fossil identification to the family level more difficult. Here we list some tentative differences observed among examined specimens. In *Dipsosaurus dorsalis*, the postparietal processes gradually taper to a tip while in *Crotaphytus*, the postparietal processes are often widened until the posterior end. The dorsal crests on the postparietal processes in *Dipsosaurus dorsalis* are slanted laterally or lie directly midline along the process so that the medial surface of the postparietal process is visible in dorsal view. In crotaphytids, the crests on the postparietal processes are slanted medially, even just slightly, so that the medial surface of the postparietal process is obscured in dorsal view. The ventrolateral crests in crotaphytids are oriented more laterally compared to *Dipsosaurus dorsalis* and phrynosomatids, such that more of the supratemporal fossa is visible in dorsal view. The hoplocercid *Enyalioides heterolepis* has a large portion of the supratemporal fossa visible in dorsal view but has adductor crests that form a “V” shape. The posterior portion of the parietal table is reportedly more constricted in *Crotaphytus* compared to *Gambelia*, but it was noted that there is substantial ontogenetic and perhaps sexual variation (McGuire 1996), so we refrain from making generic identifications.

**Figure 9.**
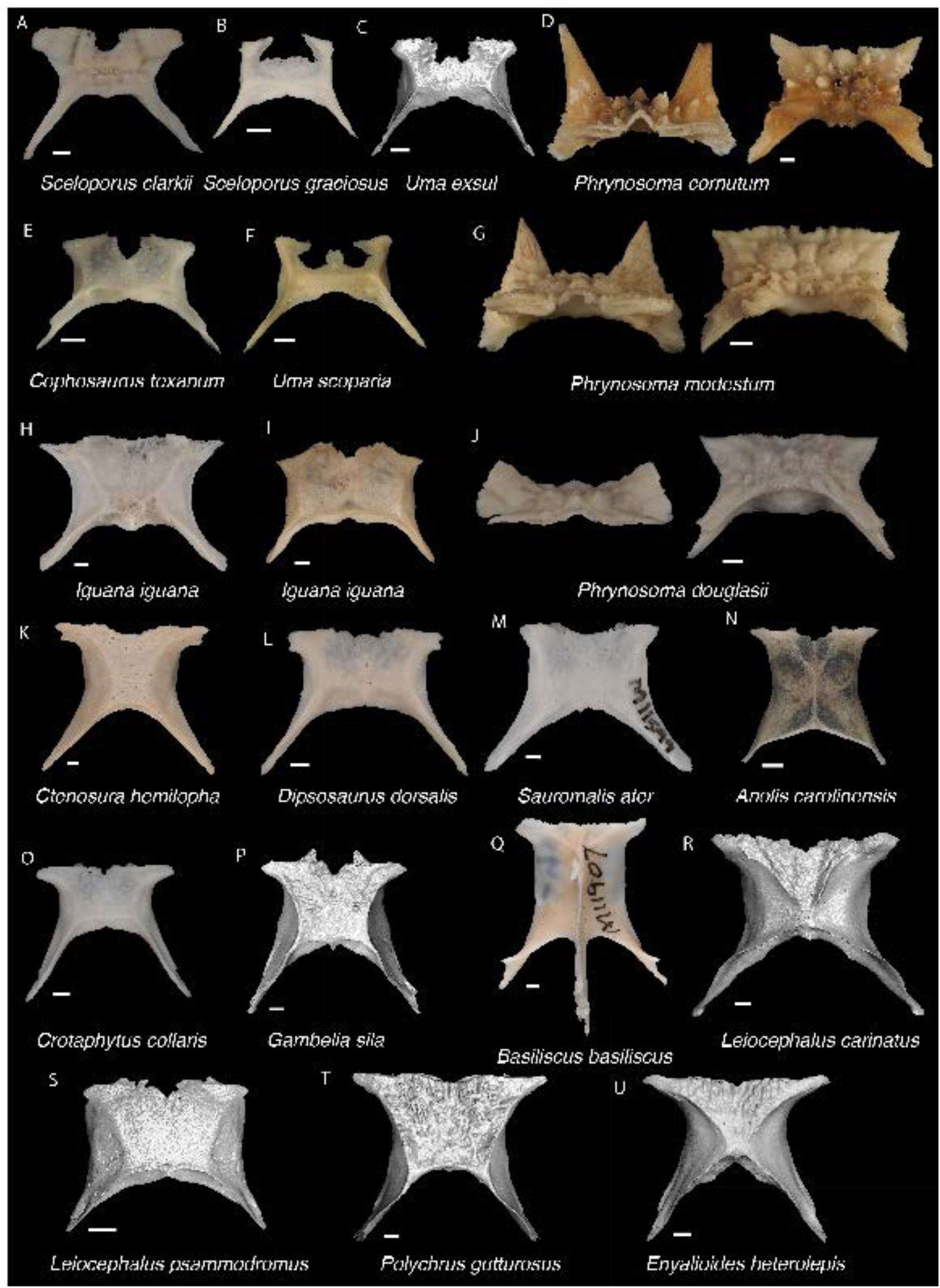
Pleurodontan Parietals. Parietals in dorsal view– **A**. *Sceloporus clarkii* TxVP M-12157; **B**. *Sceloporus graciosus* TxVP M-14879; **C**. *Uma exsul* TNHC 30247; **D**. *Phrynosoma cornutum* TxVP M-6405; **E**. *Cophosaurus texanum* TxVP M-8527; **F**. *Uma scoparia* TxVP M-8529; **G**. *Phrynosoma modestum* TNHC 95921; **H**. *Iguana iguana* TxVP M-8454; **I**. *Iguana iguana* TxVP M-13054; **J**. *Phrynosoma douglasii* TxVP M-8526; **K**. *Ctenosura hemilopha* TxVP M-8616; **L**. *Dipsosaurus dorsalis* TxVP M-13086; **M**. *Sauromalis ater* TxVP M-11599; **N**. *Anolis carolinensis* TxVP M-9042; **O**. *Crotaphytus collaris* TxVP M-12468; **P**. *Gambelia sila* TNHC 95261; **Q**. *Basiliscus basiliscus* TxVP M-11907; **R**. *Leiocephalus carinatus* TNHC 89274; **S**. *Leiocephalus psammodromus* TNHC 103220; **T**. *Polychrus gutturosus* TNHC 24152; **U**. *Enyalioides heterolepis* UF 68015. Scale bars = 1 mm.

### Jugal

#### Description

TxVP 41229-25945 serves as the basis for our description (Fig. 3). TxVP 41229-25945 is a left jugal. Laterally, there is a distinct maxillary facet on the ventral half of the suborbital process and a depression near the inflection point. There is a distinct, wide quadratojugal process. The postorbital process is long, curves posteriorly, and has a distinct postorbital facet on the anterodorsal surface. On the medial surface, there is a distinct ridge that is located at the dorsal margin of the suborbital process and that runs posteriorly on the anterior edge of the postorbital process. There are several foramina below the medial ridge and on the lateral surface including within the depression, on the postorbital process, and on the quadratojugal process.

#### Identification

Jugals were assigned to Pleurodonta based on the presence of a posteriorly deflected postorbital process (present in many but not all pleurodontan taxa, see Table S2) and a medial ridge that is located anteriorly on the postorbital process and medially on the suborbital process (Čerňanský et al. 2014). Geckos have a small, reduced jugal, and dibamids lack a jugal (Greer 1985; Evans 2008). In examined anguids and scincids, the postorbital process is oriented dorsally and the jugal is more angulated (Conrad 2008). Xantusiids differ in having a short suborbital process and *Xantusia riversiana* and *Lepidophyma* have an anteroposteriorly widened postorbital process (Savage 1963). Teiioids differ in having a distinct medial ectopterygoid process, which is also present, albeit smaller, in some gymnophthalmoids (Evans 2008). The position of the medial ridge varies among North American lizards, but the presence of a medial ridge that is located anteriorly on the postorbital process and medially on the suborbital process is consistent with several pleurodontan clades (Čerňanský et al. 2014). Examined phrynosomatids, *Leiocephalus*, *Enyalioides heterolepis*, *Polychrus* (except *P. femoralis*; Smith 2011), and iguanids differ from crotaphytids and the fossils in lacking a posteroventral process (quadratojugal process) (Smith 2011; Fig. 10). Additionally, the anterior suborbital process in *Iguana iguana* and *Dipsosaurus dorsalis* is shortened relative to crotaphytids and the fossils. Examined corytophanids also lacks a posteroventral process and the postorbital process is directed only dorsally, whereas in *Crotaphytus* the process is posteriorly deflected (Smith 2009a). Examined *Corytophanes* differ in having a posteriorly widened postorbital process. Examined *Anolis* also have a posteroventral process (Smith 2011), albeit somewhat less distinct than in the fossil and in crotaphytids. Examined anoles differ from crotaphytids and the fossils in having a thinner anterior suborbital process and a dorsal margin of the suborbital process that is everted laterally to a lesser degree. Additionally, examined *Anolis* differ in having a posteriorly widened postorbital process that tapers dorsally. On this basis, the fossil was assigned to Crotaphytidae.

**Figure 10.**
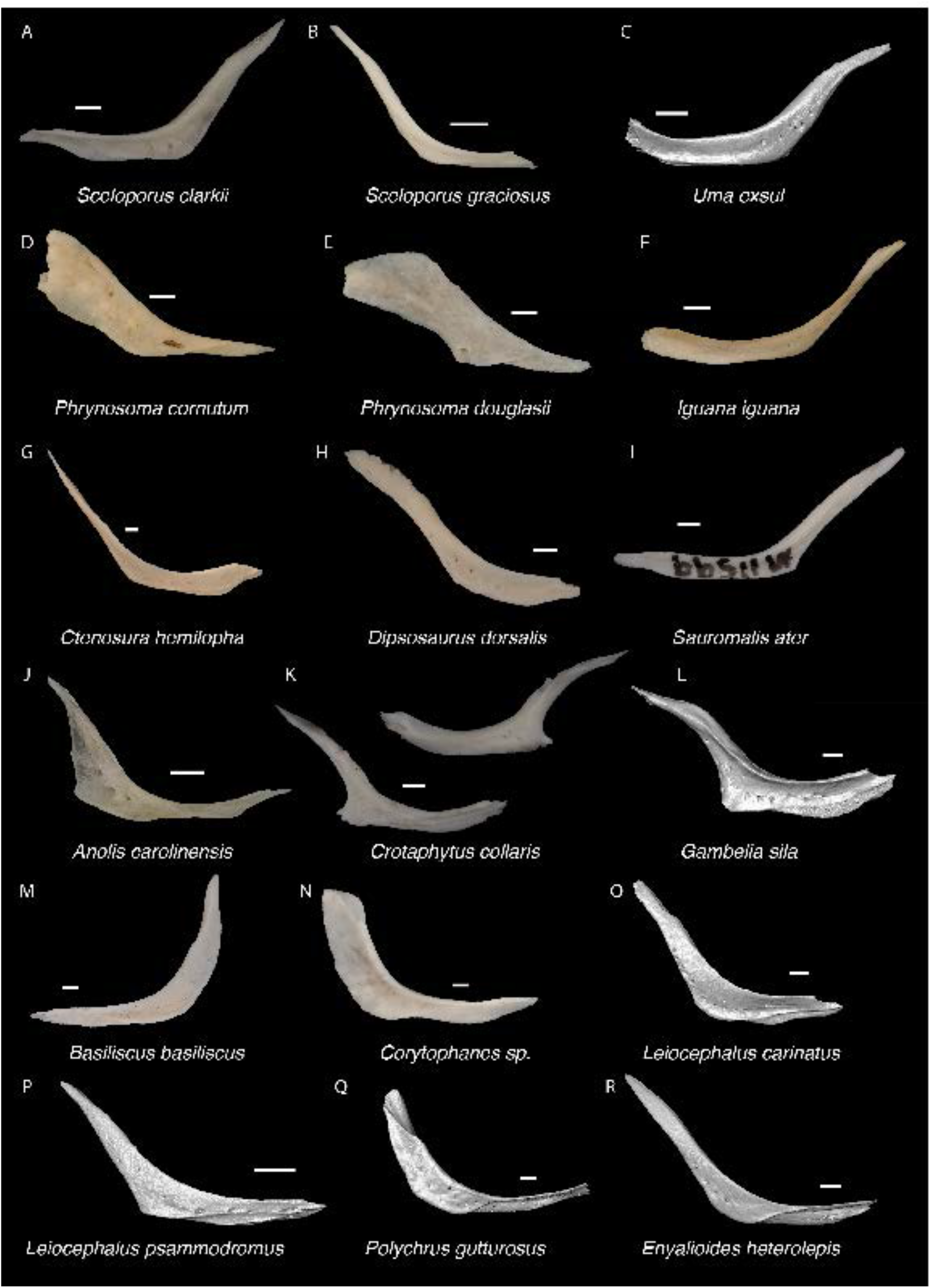
Pleurodontan Jugals. Jugals in lateral view– A. *Sceloporus clarkii* TxVP M-12157; B. *Sceloporus graciosus* TxVP M-14879; C. *Uma exsul* TNHC 30247; D. *Phrynosoma cornutum* TxVP M-6405; E. *Phrynosoma douglasii* TxVP M-8526; F. *Iguana iguana* TxVP M-13054; G. *Ctenosaura hemilopha* TxVP M-8616; H. *Dipsosaurus dorsalis* TxVP M-13086; I. *Sauromalis ater* TxVP M-11599; J. *Anolis carolinensis* TxVP M-9042; K. *Crotaphytus collaris* TxVP M-12468; L. *Gambelia sila* TNHC 95261; M. *Basiliscus basiliscus* TxVP M-11907; N. *Corytophanes* sp. TxVP M-16765; O. *Leiocephalus carinatus* TNHC 89274; P. *Leiocephalus psammodromus* TNHC 103220; Q. *Polychrus gutturosus* TNHC 24152; R. *Enyalioides heterolepis* UF 68015. Scale bars = 1 mm.

### Postorbital

#### Description

TxVP 41229-27010 is a well preserved left postorbital (Fig. 3). It is triradiate with a thin anterior process and thicker dorsal and posterior processes. There are distinct jugal and squamosal facets along the ventrolateral edge. The posterior edge is concave and there is a distinct tubercle near the middle of the anterior surface of the dorsal process. The tip of the dorsal process is squared off. In TxVP 41229-26975 the squamosal facet is less visible in dorsolateral view compared to TxVP 41229-27010, and the tubercle is on the posterior edge of the dorsal process. The posterior process and the tip of the dorsal process are broken in TxVP 41229-28984. TxVP 41229-28984 differs from the other specimens in having a smaller tubercle on the dorsal process.

#### Identification

Postorbitals were identified to Pleurodonta based on a sub-triangular shape (Fig. 11) with a distinct ventral process (Estes et al. 1988). Many anguimorphs and skinks differ in having a long posterior process of the postorbital, and teiids have a quadradiate postorbitofrontal. Among pleurodontans, crotaphytids and the fossils differ from phrynosomatids, *Enyalioides*, *Anolis*, and some *Leiocephalus* in having a large, round knob laterally on the postorbital (Smith 2009a, b). *Leiocephalus carinatus* has a knob but differs from the fossils in having a much taller posterior process and a posteriorly deflected dorsal process with an anterior articulation facet for the frontoparietal corner. *Polychrus* have a knob (Smith 2009a, b), but differ from the fossils in having a distinct longitudinal canthal crest (Smith 2011). Examined juveniles of *Iguana iguana* lack a large round knob, but large skeletally mature *Iguana iguana* have a large knob. In large *Iguana* the bone is wider closer to the articulation with the frontal. In iguanids and corytophanids, the knob is located closer to the articulation with the frontal compared to the fossils and crotaphytids (Smith 2009a). In addition, corytophanids differ in having a convex dorsal margin of the posterior process where it borders the supratemporal fenestra (Smith 2009a). On this basis, the fossil was assigned to Crotaphytidae.*rolepis*

**Figure 11.**
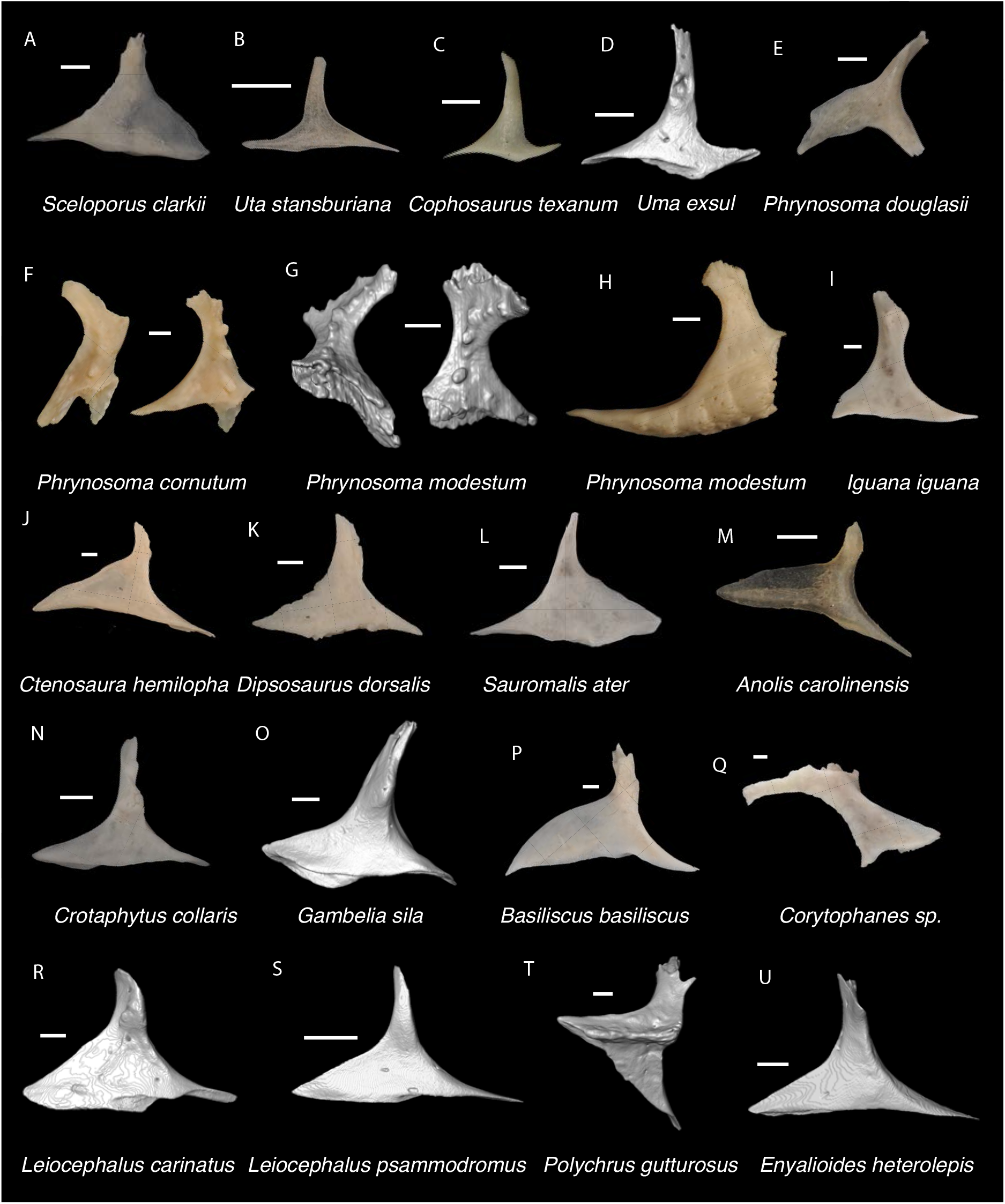
Pleurodontan Postorbitals. Postorbitals in dorsolateral and dorsal views– **A**. *Sceloporus clarkii* TxVP M-12157; **B**. *Uta stansburiana* TxVP M-14935; **C**. *Cophosaurus texanum* TxVP M-8527; **D**. *Uma exsul* TNHC 30247; **E**. *Phrynosoma douglasii* TxVP M-8526; **F**. *Phrynosoma cornutum* TxVP M-6405; **G**. *Phrynosoma modestum* TNHC 95921; **H**. *Phrynosoma modestum* TNHC 95921; **I**. *Iguana iguana* TxVP M-8454; **J**. *Ctenosura hemilopha* TxVP M-8616; **K**. *Dipsosaurus dorsalis* TxVP M-13086; **L**. *Sauromalis ater* TxVP M-11599; **M**. *Anolis carolinensis* TxVP M-9042; **N**. *Crotaphytus collaris* TxVP M-12468; **O**. *Gambelia sila* TNHC 95261; **P**. *Basiliscus basiliscus* TxVP M-11907; **Q**. *Corytophanes* sp. TxVP M-16765; **R**. *Leiocephalus carinatus* TNHC 89274; **S**. *Leiocephalus psammodromus* TNHC 103220; **T**. *Polychrus gutturosus* TNHC 24152; **U**. *Enyalioides heterolepis* UF 68015. Scale bars = 1 mm.

### Squamosal

#### Description

TxVP 41229-28355 is a left squamosal (Fig. 3). The main shaft of the bone is concave ventrally and there are distinct posterodorsal and posteroventral processes. The posterodorsal process is elongated and pointed and the posteroventral process is rounded and points anteroventrally. There is an elongate facet of the posterodorsal process for articulation with the postparietal process.

#### Identification

TxVP 41229-28355 shares with Pleurodonta, some teiids, and xenosaurids a posterodorsal process (Estes et al. 1988; Fig. 12). Some examined *Aspidoscelis* (e.g., *A. tigris* TxVP M-13877) and other teiids (e.g., *Ameiva sp*. TxVP M-8459) also have a distinct dorsal and ventral projection on the posterior end of the squamosal (Tedesco et al., 1999; Evans 2008). However, the posteroventral process projects ventrally or posteriorly in examined *Ameiva* (Evans 2008) and *Aspidoscelis*, and the posteroventral process is often shorter and less distinct in examined *Aspidoscelis* (Fig. 13). The dorsal process of *Xenosaurus* differs in that it is developed into a broad sheet and more anteriorly located along the squamosal (Bhullar 2011). On this basis, the fossil was assigned to Pleurodonta. The fossil and crotaphytids differ from iguanids, *Enyalioides heterolepis*, and phrynosomatids in having a longer and more distinct posterodorsal process of the squamosal (de Queiroz 1987). In some crotaphytids (e.g., *C. bicinctores* TxVP M-8612, M-8947) the posterodorsal process is less distinct than in other *Crotaphytus* because it is connected with the main rod by a broad sheet of bone, but no fossils were found with this morphology. Most corytophanids, except *Laemanctus longipes*, have a relatively shorter posterodorsal process (Smith 2011), and *Laemanctus longipes* differs in having a distinctly downturned main rod of the squamosal. *Anolis* and *Leiocephalus* also have a shorter posterodorsal process compared to the fossil and crotaphytids (Pregill 1992; Smith 2011). The posterodorsal process is long among *Polychrus* (Smith 2011); however, *Polychrus* differs in having a relatively shorter main rod of the squamosal and may have lateral protuberances (Smith 2011). On this basis, the fossil was assigned to Crotaphytidae.

**Figure 12.**
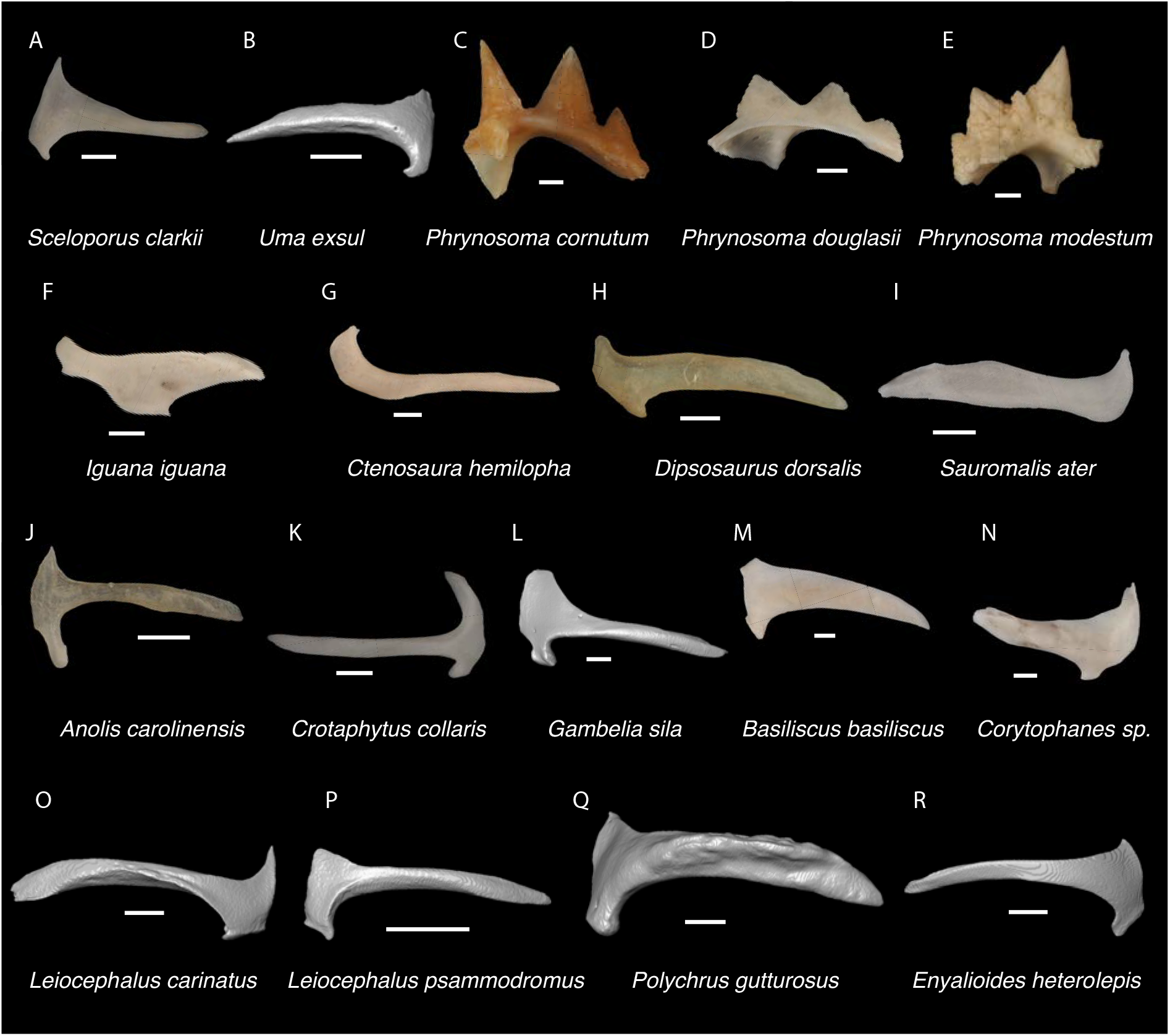
Pleurodontan Squamosals. Squamosals in lateral and posterior views– **A**. *Sceloporus clarkii* TxVP M-12157; **B**. *Uma exsul* TNHC 30247; **C**. *Phrynosoma cornutum* TxVP M-6405; **D**. *Phrynosoma douglasii* TxVP M-8526; **E**. *Phrynosoma modestum* TNHC 95921; **F**. *Iguana iguana* TxVP M-8454; **G**. *Ctenosaura hemilopha* TxVP M-8616; **H**. *Dipsosaurus dorsalis* TxVP M-9285; **I**. *Sauromalis ater* TxVP M-11599; **J**. *Anolis carolinensis* TxVP M-9042; **K**. *Crotaphytus collaris* TxVP M-12468; **L**. *Gambelia sila* TNHC 95261; **M**. *Basiliscus basiliscus* TxVP M-11907; **N**. *Corytophanes* sp. TxVP M-16765; **O**. *Leiocephalus carinatus* TNHC 89274; **P**. *Leiocephalus psammodromus* TNHC 103220; **Q**. *Polychrus gutturosus* TNHC 24152; **R**. *Enyalioides heterolepis* UF 68015. Scale bars = 1 mm.

**Figure 13.**
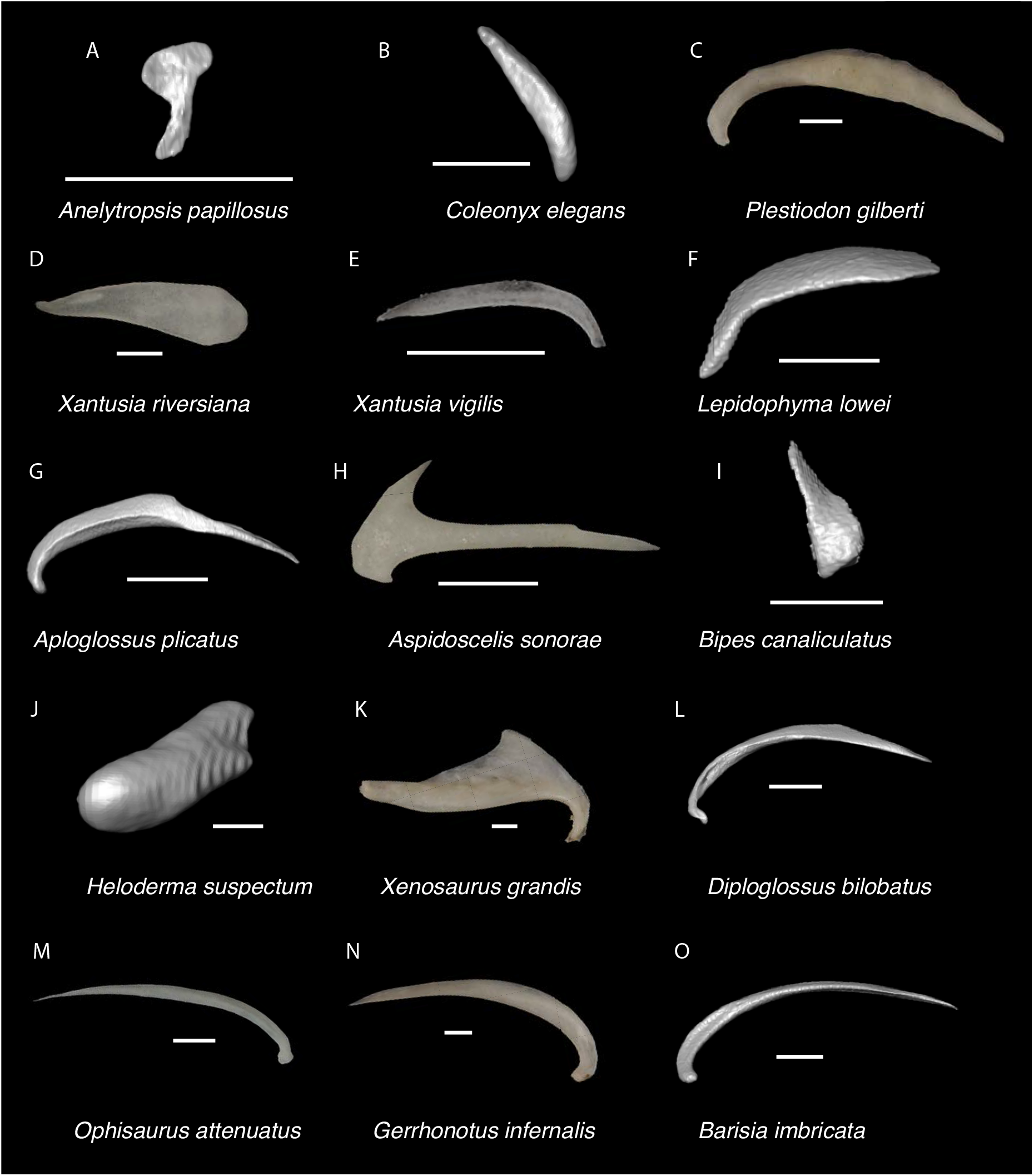
Non-Pleurodontan Squamosals. Squamosals in lateral view– **A**. *Anelytropsis papillosus* UF 86708; **B**. *Coleonyx elegans* UF 11258; **C**. *Plestiodon gilberti* TxVP M-8587; **D**. *Xantusia riversiana* TxVP M-8505; **E**. *Xantusia virigatus* TxVP M-12130; **F**. *Lepidophyma lowei* LACM 143367; **G**. *Aploglossus plicatus* TNHC 34481; **H**. *Aspidoscelis sonorae* TxVP M-15670; **I**. *Bipes canaliculatus* CAS 134753; **J**. *Heloderma suspectum* TNHC 62766; **K**. *Xenosaurus grandis* TxVP M-8960; **L**. *Diploglossus bilobatus* TNHC 31933; **M**. *Ophisaurus attenuatus* TxVP M-8979; **N**. *Gerrhonotus infernalis* TxVP M-13441; **O**. *Barisia imbricata* TNHC 76984. Scale bars = 1 mm.

### Quadrate

#### Description

TxVP 41229-25579 is a left quadrate that is well preserved but is missing the dorsolateral portion of the bone (Fig. 3). The central column is wide, and the bone narrows ventrally. There is no pterygoid lappet, but there is a well-developed and anteromedially directed medial crest. The conch is deep, gradually slants laterally from the central column, and narrows ventrally. The cephalic condyle projects posteriorly and there is no extensive ossification dorsally. The dorsal margin is straight past the lateral extent of the mandibular condyle. There is a foramen medial to the central column.

#### Identification

TxVP 41229-25579 shares with geckos, *Scincella*, *Xantusia*, some pleurodontans, and anguimorphs (besides *Heloderma*) the absence of a distinct pterygoid lappet (Estes et al. 1988; Evans 2008). The fossil quadrate differs from that of geckos and *Xantusia* in being relatively wide overall and with a wide central column (Kluge 1962; Evans 2008). The fossil and pleurodontans differ from anguimorphs in having a quadrate that widens dorsally. On this basis, the fossil was identified to Pleurodonta. Compared to the fossil and examined crotaphytids, the quadrate of *Dipsosaurus dorsalis* is much wider dorsally relative to the articular surface, the conch is deeper, and the pterygoid lamina does not extend as far anteriorly (Fig. 14). The quadrate of a small *Iguana iguana* (TxVP M-13054; SVL = 90 mm) is much slenderer and has a reduced pterygoid lamina compared to the fossil and crotaphytids. Larger *Iguana iguana* differ from the fossil and crotaphytids in that the pterygoid lamina is extended medially such that the central column is vertically oriented and near the midline of the bone (*Iguana iguana* OUVC 10612). The quadrate of most examined *Sauromalus* (except *Sauromalis ater* TNHC 18483) differ in having a low dorsolateral portion, such that the dorsal margin of the bone is sloped ventrolaterally. Additionally, the quadrate of *Sauromalus* has a dorsomedial expansion that contacts the paroccipital process (Evans 2008) and which is especially prominent in large specimens. *Ctenosaura* differs in having a more distinct and curved tympanic crest and a deeper conch. In *Anolis* there is a distinct boss at the ventromedial margin of the bone below the articulation with the pterygoid. In *Anolis* and *Polychrus*, the lateral and medial margins are more parallel compared to the fossil and crotaphytids. *Leiocephalus barahonensis* (USNM 260564) and *L. carinatus* differ from crotaphytids in having a medially expanded pterygoid lamina that extends medially beyond the dorsomedial corner of the quadrate (see also fig. 6 in Bochaton et al. 2021). *Leiocephalus psammodromus* differs from crotaphytids in having a quadrate that is not as distinctly widened dorsally and having a distinct medial notch where the pterygoid articulates. The quadrate conch of *Enyalioides heterolepis* is more shallow compared to the fossil and crotaphytids. The quadrate of *Basiliscus basiliscus* is proportionally taller and the central column more slender. The quadrate is exceptionally slender in examined *Corytophanes* and *Laemanctus* (Lang 1989). The quadrate of phrynosomatids, except for *Phrynosoma*, differs from crotaphytids in having a more curved lateral margin (tympanic crest) (Mead 1988). Additionally, the central column is more slender near the base and the conch is deeper in examined large-bodied *Sceloporus* (e.g., *S. clarkii*, Fig. 14). The quadrate of *Phrynosoma* differs from the fossil and crotaphytids in having a reduced pterygoid lamina, a dorsal portion of the bone that is much wider than the ventral portion, and a conch that remains deep lateral to the central column. Based on these differences, quadrates were assigned to Crotaphytidae.

**Figure 14.**
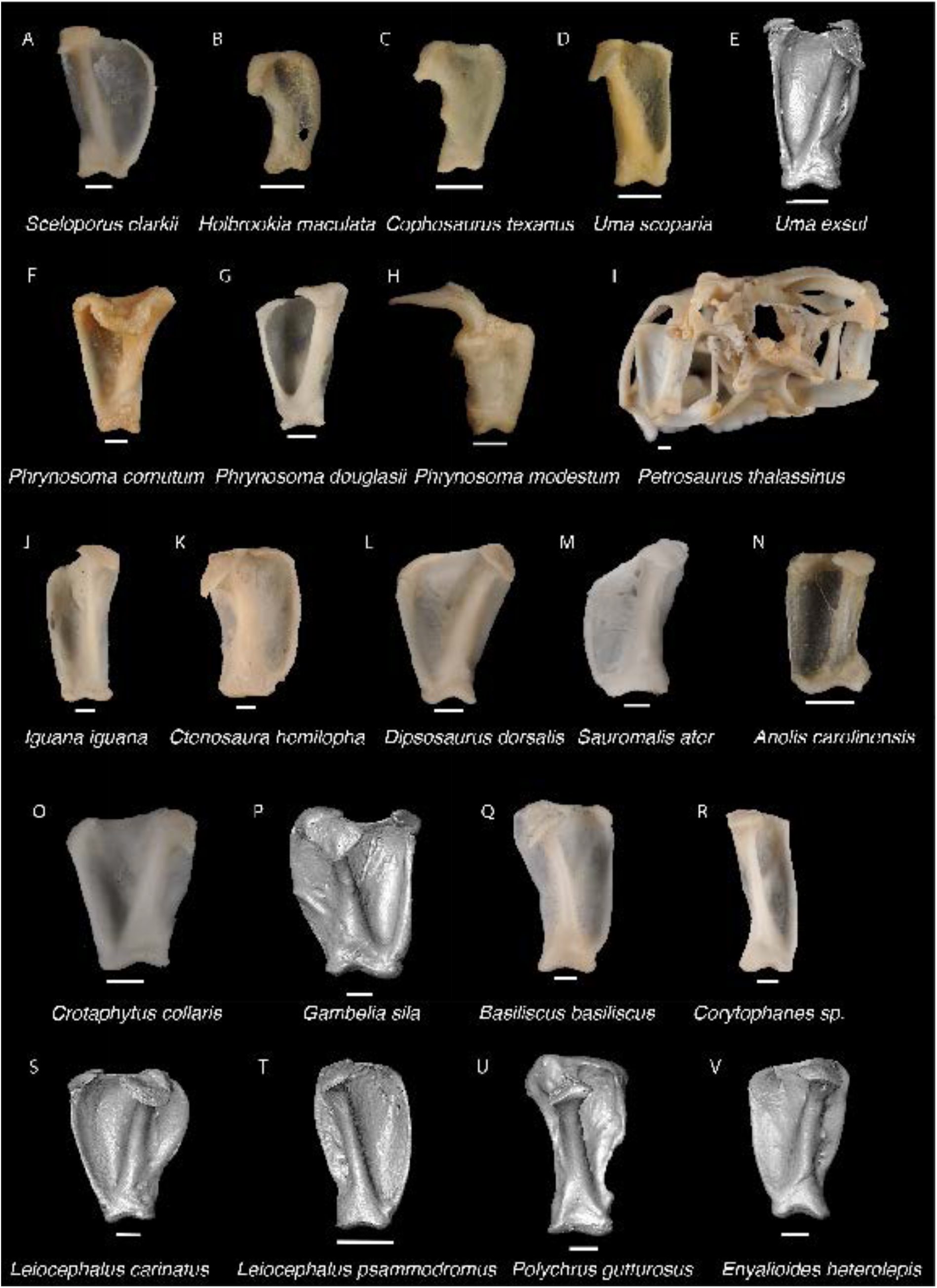
Pleurodontan Quadrates. Quadrates in posterior view– **A**. *Sceloporus clarkii* TxVP M-12157; **B**. *Holbrookia maculata* TxVP M-14322; **C**. *Cophosaurus texanum* TxVP M-8527; **D**. *Uma scoparia* TxVP M-8529; **E**. *Uma exsul* TNHC 30247; **F**. *Phrynosoma cornutum* TxVP M-6405; **G**. *Phrynosoma douglasii* TxVP M-8526; **H**. *Phrynosoma modestum* TNHC 95921; **I**. *Petrosaurus thalassinus* TxVP M-9612; **J**. *Iguana iguana* TxVP M-8454; **K**. *Ctenosura hemilopha* TxVP M-8616; **L**. *Dipsosaurus dorsalis* TxVP M-13086; **M**. *Sauromalis ater* TxVP M-11599; **N**. *Anolis carolinensis* TxVP M-9042; **O**. *Crotaphytus collaris* TxVP M-12468; **P**. *Gambelia sila* TNHC 95261; **Q**. *Basiliscus basiliscus* TxVP M-11907; **R**. *Corytophanes* sp. TxVP M-16765 **S**. *Leiocephalus carinatus* TNHC 89274; **T**. *Leiocephalus psammodromus* TNHC 103220; **U**. *Polychrus gutturosus* TNHC 24152; **V**. *Enyalioides heterolepis* UF 68015. Scale bars = 1 mm.

### Pterygoid

#### Description

TxVP 41229-28513 is a pterygoid that is missing only the distal end of the palatine process (Fig. 15). The palatine process is thin near its base and the transverse process extends laterally at nearly a 90-degree angle with the quadrate process. The transverse process is dorsoventrally tall, and the distal end is oriented near vertical. There is a distinct ridge on the dorsal surface for insertion of the superficial pseudotemporal muscle and a distinct ectopterygoid facet just anterior to that. There is a large ridge on the ventral surface for insertion of the pterygomandibular muscle and a small, deep fossa columella that continues posteriorly as a short, narrow groove. The quadrate process is elongated, and the medial surface has an elongate groove that serves for insertion of the pterygoideus muscle. There is a ventromedial projection at the floor of the basipterygoid fossa. There is a long patch of 17 tooth positions filled with 16 pterygoid teeth. There are four small foramina on the dorsal surface in the area from the palatine process to just anterior to the fossa columella. There are two foramina just anterior to the ridge for insertion of the pterygomandibular muscle. TxVP 41229-27144 and TxVP 41229-28866 are missing portions of the palatine and quadrate processes, and do not differ substantively from TxVP 41229-28513.

**Figure 15.**
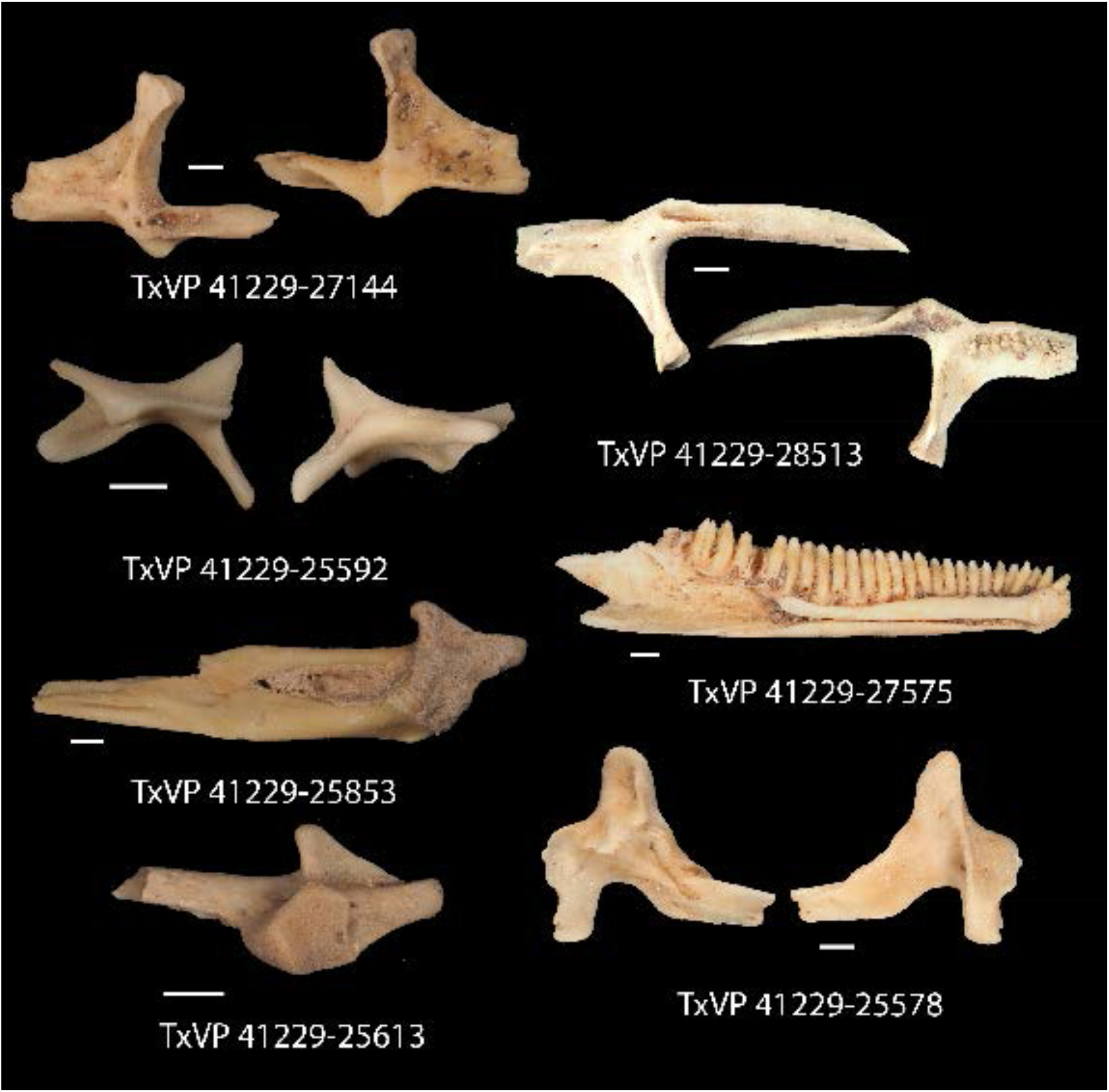
Fossil crotaphytids. TxVP 41229-27144 Dorsal and ventral view of right pterygoid; TxVP 41229-28513 Dorsal and ventral view of left pterygoid; TxVP 41229-25592 Posterior and ventral view of right ectopterygoid; TxVP 41229-27575 Medial view of left dentary; TxVP 41229-25853 Dorsal view of left compound bone; TxVP 41229-25613 Dorsal view of left articular; TxVP 41229-25578 Lateral and medial view of right coronoid. Scale bars = 1 mm.

#### Identification

Pterygoids are assigned to Pleurodonta based on the presence of a well-developed ventromedial projection at the floor of the basipterygoid fossa (Evans 2008; Smith 2009b; Fig. 16). A ventromedial projection at the floor of the basipterygoid fossa is also in some non-pleurodontans (e.g., *Ophisaurus*), but it is less distinct than in the fossil and in many pleurodontans (e.g., *Crotaphytus*). Among pleurodontans, pterygoid teeth are absent in phrynosomatids and are variably present in *Anolis*, *Polychrus*, *Dipsosaurus*, and *Leiocephalus* (Etheridge and de Queiroz 1988). The palatal plate in *Anolis* and phrynosomatids is broad relative to the fossils. Compared with the fossils, *Sauromalus* has a shorter transverse process and a shorter quadrate process. The quadrate process in *Iguana iguana* is more distinctly short and the fossa columella elongated. Pterygoid teeth of examined *Ctenosaura* are oriented more oblique (Mahler and Kearney 2006), and the palatal plate is stepped along the anterior edge (Oelrich 1956). An examined *Ctenosaura clarki* (TxVP M-14824) has a unique ridge on the ventral surface of the pterygoid along which the teeth are positioned. *Dipsosaurus dorsalis* has a narrow linear notch on the transverse process not seen on the fossils. The transverse process is shorter and less robust in *Enyalioides heterolepis*. *Basiliscus* and *Corytophanes* have a taller quadrate process and tend to have longer pterygoid teeth which may be slightly recurved in *Basiliscus* (Lang 1989). *Basiliscus* and *Laemanctus* generally have fewer pterygoid teeth ranging from three to five in number (Lang 1989). The shelf under the basipterygoid fossa is less extensive in *Basiliscus basiliscus*. The quadrate process is anteroposterioly shorter, and the transverse process is exceptionally tall in *Corytophanes*. Among North American pleurodontans, the largest number of pterygoid teeth is found in crotaphytids and *Ctenosaura* which can have a similar number of tooth positions to the fossils (Taylor 1940). However, there is a large amount of variation in the number of pterygoid teeth among crotaphytids (Taylor 1940). The transverse process of the fossils and of *Crotaphytus* is more dorsoventrally elongated and less pointed in dorsal view compared to examined *Leiocephalus*. Pterygoids are assigned to Crotaphytidae based on the differences from other NA pleurodontans described above.

**Figure 16.**
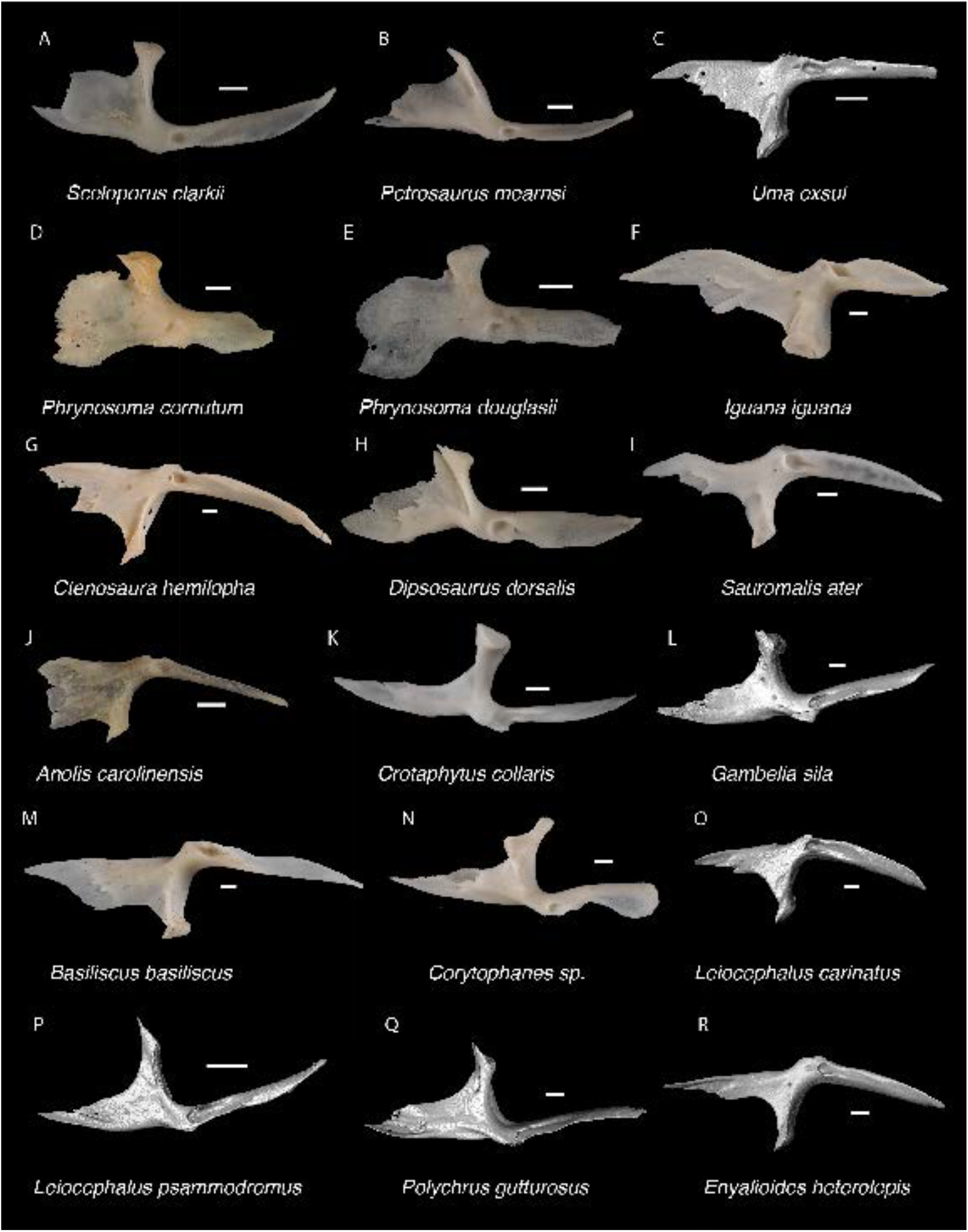
Pleurodontan Pterygoids. Pterygoids in dorsal view– **A**. *Sceloporus clarkii* TxVP M-12157; **B**. *Petrosaurus mearnsi* TxVP M-14910; **C**. *Uma exsul* TNHC 30247; **D**. *Phrynosoma cornutum* TxVP M-6405; **E**. *Phrynosoma douglasii* TxVP M-8526; **F**. *Iguana iguana* TxVP M-8454; **G**. *Ctenosaura hemilopha* TxVP M-8616; **H**. *Dipsosaurus dorsalis* TxVP M-13086; **I**. *Sauromalis ater* TxVP M-11599; **J**. *Anolis carolinensis* TxVP M-9042; **K**. *Crotaphytus collaris* TxVP M-12468; **L**. *Gambelia sila* TNHC 95261; **M**. *Basiliscus basiliscus* TxVP M-11907; **N**. *Corytophanes* sp. TxVP M-16765; **O**. *Leiocephalus carinatus* TNHC 89274; **P**. *Leiocephalus psammodromus* TNHC 103220; **Q**. *Polychrus gutturosus* TNHC 24152; **R**. *Enyalioides heterolepis* UF 68015. Scale bars = 1 mm.

### Ectopterygoid

#### Description

TxVP 41229-25592 is a right ectopterygoid (Fig. 15). It has distinct anterolateral and posterolateral processes. The ventral corner of the posterolateral process is narrow and extends far ventrally. There is a groove for articulation with the maxilla on the anterolateral edge of the bone below the dorsal corner. The dorsal surface is slightly concave at the lateral end and the pterygoid process extends far ventrally. The pterygoid facet is deep with a distinct overhanging corner of bone dorsally. The main shaft of the ectopterygoid is orthogonal to the lateral edge of the bone and there are two small foramina within the jugal facet.

#### Identification

TxVP 41229-25592 share with iguanians, xantusiids, and xenosaurids the presence of an elongate posterolateral process on the ectopterygoid (Smith 2009b). The ectopterygoid (=os transversum of Young 1942) of *Xantusia* can be distinguished from that of pleurodontans in being a broad triangular plate (Young 1942; Fig. 17). The ectopterygoid of xenosaurids differs from pleurodontans in having the main shaft of the bone oriented more anteroposteriorly in the skull and in having a much shorter posterolateral process (Bhullar 2011). Among NA pleurodontans, *Anolis* differs from the fossil and Crotaphytidae in lacking a flange on the lateral head that articulates ventrally or ventromedially with the maxilla, and in having a main shaft that is oblique relative to the lateral face of the bone (Smith 2009b; Fig. 18). The main shaft of the ectopterygoid is slightly oblique relative to the lateral face of the bone in some corytophanids, including *Corytophanes hernandesii* and *Laemanctus longipes* (Smith 2011). In phrynosomatids, *Leiocephalus*, and *Enyalioides heterolepis*, the main shaft of the ectopterygoid is also slightly oblique relative to the lateral face of the bone, while in the fossil and *Crotaphytus*, the main shaft is orthogonal to the lateral face of the bone. Additionally, the shaft of the bone is much more slender in non-*Phrynosoma* phrynosomatids compared to the fossil and *Crotaphytus*. The ventral corner of the posterolateral process of the ectopterygoid in the fossil and *Crotaphytus* is developed into a distinct wedge that projects ventrally, similar to that seen in corytophanids (Smith 2011). The ventral corner of the posterolateral process is well-developed in crotaphytids, corytophanids, and *Dipsosaurus dorsalis*, semi-developed in *Ctenosaura hemilopha*, and undeveloped in *Sauromalus ater* and *Ctenosaura similis* (Smith 2011). Although the ventral corner of the posterolateral process is well-developed in *Dipsosaurus dorsalis*, it does not project as far ventrally as in corytophanids and *Crotaphytus collaris*. In *Crotaphytus collaris* there is a corner of bone dorsal to the posterior pterygoid facet that is downturned. This downturned corner is also present in *Laemanctus longipes*, some *Anolis*, and *Enyalioides laticeps*, although it is absent in other hoplocercids (Smith 2011). A downturned corner is absent in *Polychrus* and *Gambelia wislizenii* (Smith 2011). The fossil ectopterygoid can be referred to Crotaphytidae based on possessing a main shaft that is nearly orthogonal to the lateral face of the bone, a wide shaft of the ectopterygoid, a ventral corner of the posterolateral process of the ectopterygoid developed into a distinct wedge that projects ventrally, and a downturned corner of bone dorsal to the posterior pterygoid facet. A downturned corner of bone dorsal to the posterior pterygoid facet is absent in *Gambelia wislizenii* (Smith 2011). A slightly downturned corner is present in *Gambelia sila* (TxVP M-95261), although not to the extent seen in *Crotaphytus*. More information on this feature among species of *Gambelia* is needed before a confident generic assignment can be made.

**Figure 17.**
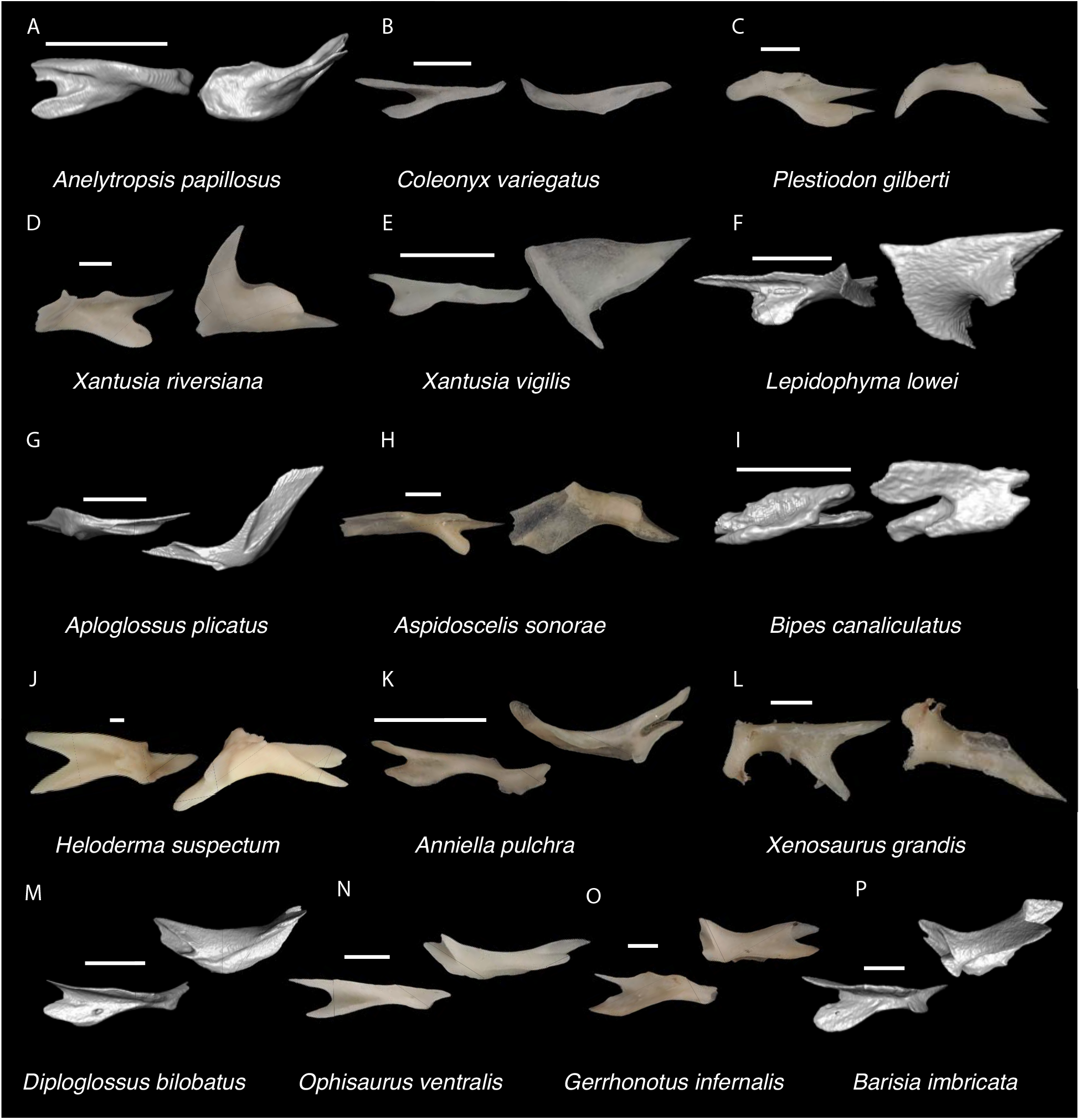
Non-Pleurodontan Ectopterygoids. Ectopterygoids in lateral and ventral views– **A**. *Anelytropsis papillosus* UF 86708; **B**. *Coleonyx variegatus* TxVP M-12109; **C**. *Plestiodon gilberti* TxVP M-8587; **D**. *Xantusia riversiana* TxVP M-8505; **E**. *Xantusia virigatus* TxVP M-12130; **F**. *Lepidophyma lowei* LACM 143367; **G**. *Aploglossus plicatus* TNHC 34481; **H**. *Aspidoscelis sonorae* TxVP M-15670; **I**. *Bipes canaliculatus* CAS 134753; **J**. *Heloderma suspectum* TxVP M-9001; **K**. *Anniella pulchra* TxVP M-8678; **L**. *Xenosaurus grandis* TxVP M-8960; **M**. *Diploglossus bilobatus* TNHC 31933; **N**. *Ophisaurus ventralis* TxVP M-8585; **O**. *Gerrhonotus infernalis* TxVP M-13441; **P**. *Barisia imbricata* TNHC 76984. Scale bars = 1 mm.

**Figure 18.**
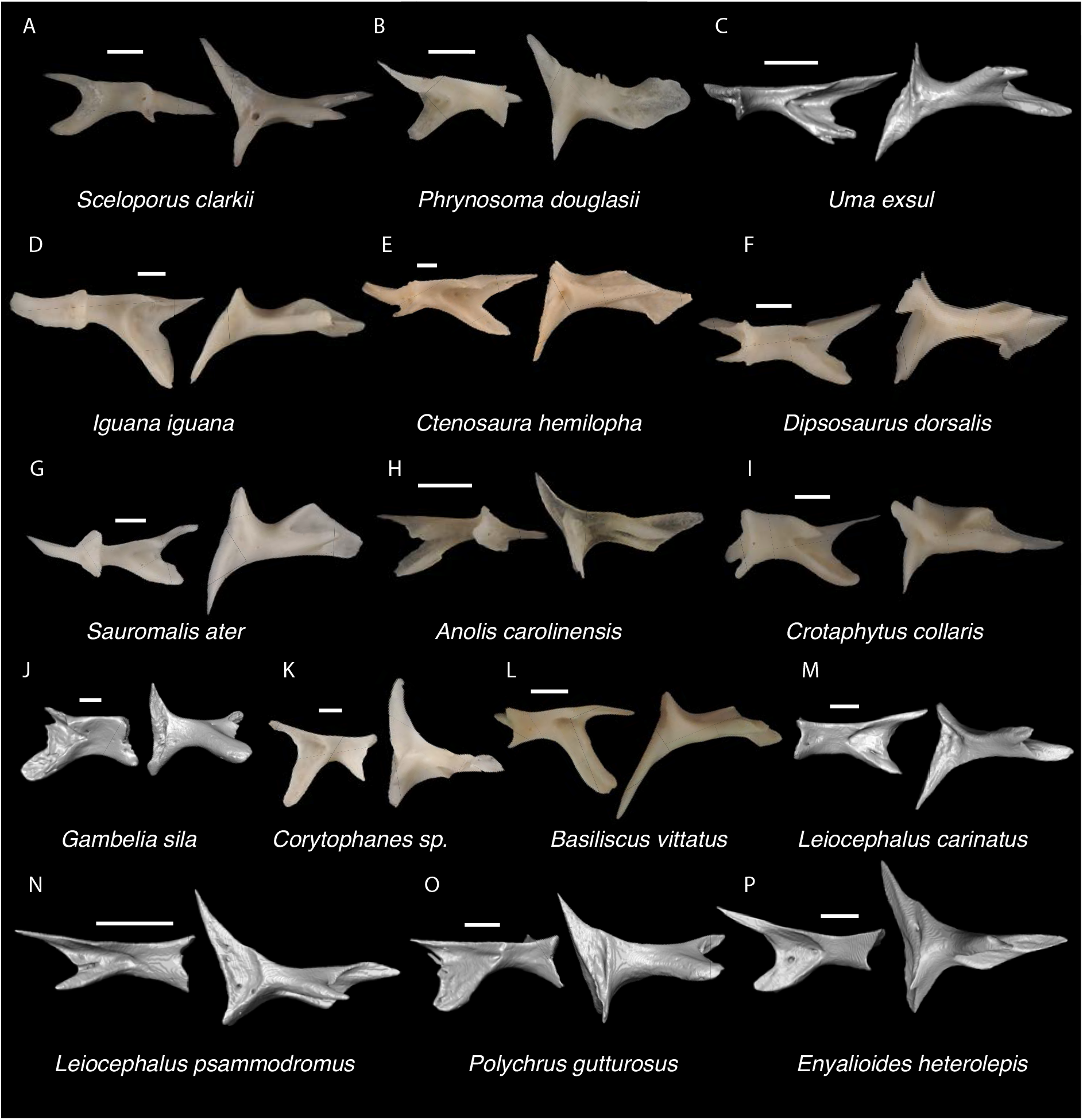
Pleurodontan Ectopterygoids. Ectopterygoids in posterior and ventral views– **A**. *Sceloporus clarkii* TxVP M-12157; **B**. *Phrynosoma douglasii* TxVP M-8526; **C**. *Uma exsul* TNHC 30247; **D**. *Iguana iguana* TxVP M-8454; **E**. *Ctenosura hemilopha* TxVP M-8616; **F**. *Dipsosaurus dorsalis* TxVP M-13086; **G**. *Sauromalis ater* TxVP M-11599; **H**. *Anolis carolinensis* TxVP M-9042; **I**. *Crotaphytus collaris* TxVP M-12468; **J**. *Gambelia sila* TNHC 95261; **K**. *Corytophanes* sp. TxVP M-16765; **L**. *Basiliscus vittatus* TxVP M-8556; **M**. *Leiocephalus carinatus* TNHC 89274; **N**. *Leiocephalus psammodromus* TNHC 103220; **O**. *Polychrus gutturosus* TNHC 24152; **P**. *Enyalioides heterolepis* UF 68015. Scale bars = 1 mm.

### Dentary

#### Description

TxVP 41229-27575 serves as the basis for our description (Fig. 15). TxVP 41229-27575 is a right dentary with 26 tooth positions. The posterior end is bifurcated and the Meckelian canal is open, with tall suprameckelian and inframeckelian lips. There is a distinct intramandibular lamella and the intramandibular septum is restricted to the anterior portion of the dentary. There is a narrow dental shelf that becomes wider closer to the symphysis. The teeth are slightly eroded at the crown, but some teeth display tricuspid morphology. The tooth bases are much wider than the crown for the more distal teeth. There are six nutrient foramina arranged in a row on the lateral surface near the anterior end of the dentary.

#### Identification

Dentaries were placed in Pleurodonta based on the presence of pleurodont tricuspid teeth and a closed or partially closed Meckelian groove bounded by the suprameckelian and inframeckelian lips (Scarpetta 2021; Fig. 19). Other non-pleurodontan lizards with tricuspid teeth include *Xantusia riversiana* (Savage 1963), some teiids (Tedesco et al. 1999), and some gymnophthalmids (Bell et al. 2003). *Xantusia* differs from pleurodontans in having a fused spleniodentary (Mead and Bell 2001). Teiids differ in having a tall and open Meckelian groove and substantial cementum deposits at the tooth bases (Estes et al. 1988; Scarpetta 2020). In gymnophthalmoids, the Meckelian groove is not bounded by suprameckelian and inframeckelian lips (Morales et al. 2019) or is completely fused to almost the level of the posterior-most tooth position (Bell et al. 2003). Dentaries were identified to Crotaphytidae based on mesiodistally expanded tooth bases, a narrow subdental shelf, and an anteriorly tall suprameckelian lip (Scarpetta 2021). Among pleurodontans, the Meckelian groove is fused in iguanids, dactyloids, extant *Leiocephalus*, tropidurids, polychrotids, leiosaurids, and some liolaemids and oplurids (Smith 2011; Scarpetta 2021). Corytophanids differ from crotaphytids in lacking mesiodistally expanded tooth bases, and in *Basiliscus*, *Corytophanes*, some *Laemanctus*, and *Enyalioides* the tooth crowns are flared (Smith 2009a; Scarpetta 2021). Examined hoplocercids besides *Enyalioides laticeps* (FMNH 206132) have an open Meckelian groove that is not bounded by suprameckelian and inframeckelian lips (Frost and Etheridge 1989; Scarpetta 2021). Phrynosomatids have relatively more slender teeth that do not widen at the base (Hollenshead and Mead 2006), often have a larger subdental shelf, and have an anteriorly short suprameckelian lip (Scarpetta 2021).

**Figure 19.**
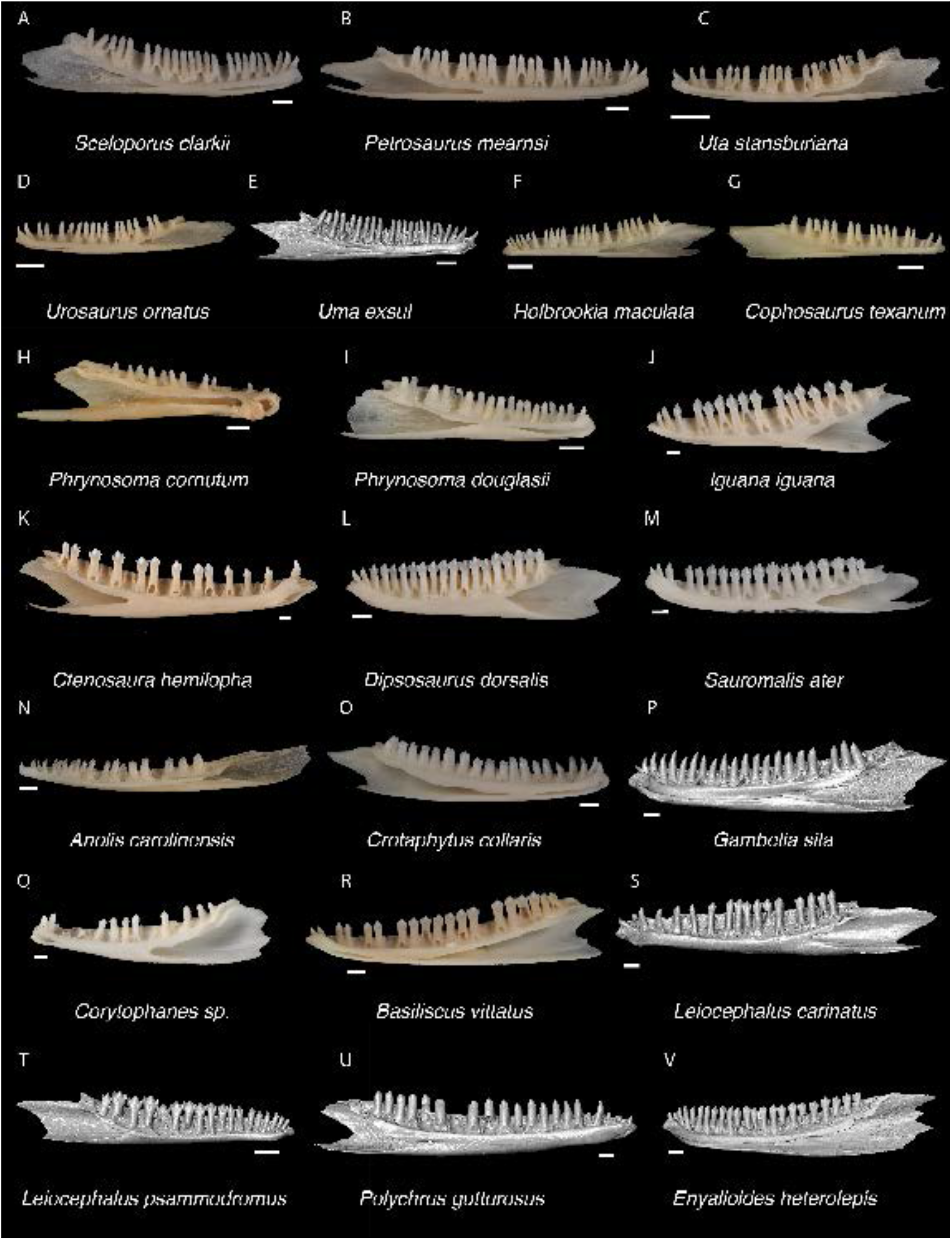
Pleurodontan Dentaries. Dentaries in medial view– **A**. *Sceloporus clarkii* TxVP M-12157; **B**. *Petrosaurus mearnsi* TxVP M-14910; **C**. *Uta stansburiana* TxVP M-14935; **D**. *Urosaurus ornatus* TxVP M-14330; **E**. *Uma exsul* TNHC 30247; **F**. *Holbrookia maculata* TxVP M-14322; **G**. *Cophosaurus texanum* TxVP M-8527; **H**. *Phrynosoma cornutum* TxVP M-6405; **I**. *Phrynosoma douglasii* TxVP M-8526; **J**. *Iguana iguana* TxVP M-8454; **K**. *Ctenosaura hemilopha* TxVP M-8616; **L**. *Dipsosaurus dorsalis* TxVP M-13086; **M**. *Sauromalis ater* TxVP M-11599; **N**. *Anolis carolinensis* TxVP M-9042; **O**. *Crotaphytus collaris* TxVP M-8395; **P**. *Gambelia sila* TNHC 95261; **Q**. *Corytophanes* sp. TxVP M-16765; **R**. *Basiliscus vittatus* TxVP M-8556; **S**. *Leiocephalus carinatus* TNHC 89274; **T**. *Leiocephalus psammodromus* TNHC 103220; **U**. *Polychrus gutturosus* TNHC 24152; **V**. *Enyalioides heterolepis* UF 68015. Scale bars = 1 mm.

### Coronoid

#### Description

TxVP 41229-25578 is a right coronoid (Fig. 15). The coronoid process is tall and rounded, the anteromedial process is elongated, and the tip is missing. The posteromedial process is oriented ventrally with an expanded lamina of bone posteriorly to articulate the surangular medially and dorsally. There is a distinct medial crest that extends from the coronoid process onto the posteromedial process. There is a small, rounded lateral process. There is a distinct vertically oriented lateral crest that ends at the anterior margin of the lateral process. There is a relatively broad facet for dorsal articulation with the surangular that has a narrow groove medial to the lateral process.

#### Identification

The fossil coronoid shares with several pleurodontans and xantusiids the lack of an anteriorly projecting lateral process that overlaps the dentary (Estes et al. 1988; Fig. 20). Although xantusiids also lack an anteriorly projecting lateral process (Estes et al. 1988), they differ from pleurodontans in having an anterior groove extending onto the coronoid process for articulation with the coronoid process of the spleniodentary (Savage 1963). Additionally, xantusiids differ from most pleurodontans and the fossil in having a relatively short and blunt anteromedial process and, except for *Xantusia riversiana*, in having a wide and low dorsal coronoid process (Savage 1963). Crotaphytids lack an anterolateral process of the coronoid, while iguanids, *Enyalioides heterolepis*, *Anolis*, and *Leiocephalus* have an anteriorly projecting anterolateral process that strongly articulates with the lateral portion of the dentary (Smith 2011; Bochaton et al. 2021). A smaller specimen of *Iguana iguana* (TxVP M-13054), similar in size to adult *Crotaphytus*, has a tall, pointed coronoid process, although this morphology varies ontogenetically in *Iguana* (Bochaton et al. 2016a). The fossil and crotaphytids are further distinguished from *Dipsosaurus* in that the posteromedial process in *Dipsosaurus dorsalis* does not extend far ventrally below the notch formed between the posteromedial and anteromedial processes. Furthermore, the anteromedial process is short in *Dipsosaurus dorsalis* relative to examined *Crotaphytus*. It was reported that *Dipsosaurus dorsalis* lacks a concavity on the posterior margin of the posterolateral process (Smith 2011); however, this feature is variable since a concavity is present on at least one specimen of *Dipsosaurus dorsalis* (Digimorph specimen YPM 14376). In corytophanids, the lateral crest on the coronoid process continues ventrally onto the medial or posterior margin of the lateral process (see fig. 69 of Smith 2011), whereas crotaphytids and the fossil have a lateral crest that merges with the anterior margin of the lateral process. Examined *Polychrus*, *Basiliscus basiliscus*, and *Corytophanes* differ from the fossil in lacking an expanded lamina of bone posterior to the coronoid process to articulate with the surangular dorsally. Phrynosomatids differ in having a relatively narrow facet on the ventral surface for articulation with the dorsal surface of the surangular. Additionally, in large species of *Sceloporus* (e.g., *S. clarkii*), the coronoid process is tall relative to the body of the bone. The posteromedial process in *Crotaphytus* is directed almost ventrally, whereas the process is directed posteromedially (∼45 degrees from vertical) in *Gambelia wislizenii* and *Gambelia copei* (Norell 1989; McGuire 1996). The more vertically oriented posteromedial process on the fossil coronoid makes it likely that the fossils represent *Crotaphytus*; however, this is tentative given a similar morphology also occurs in *Gambelia sila* (McGuire 1996).

**Figure 20.**
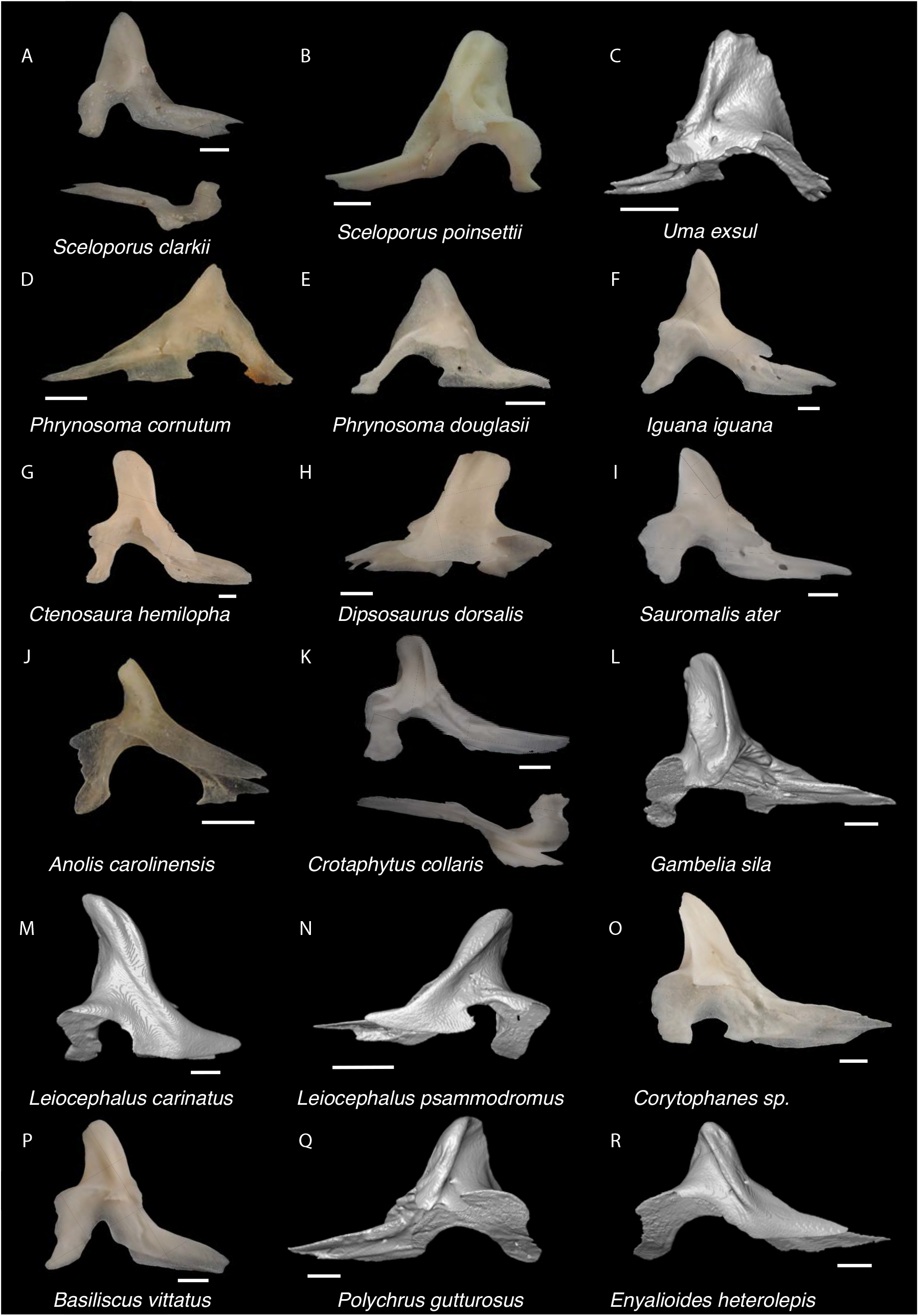
Pleurodontan Coronoids. Coronoids in lateral and ventral views– **A**. *Sceloporus clarkii* TxVP M-12157; **B**. *Sceloporus poinsettii* TxVP M-8373; **C**. *Uma exsul* TNHC 30247; **D**. *Phrynosoma cornutum* TxVP M-6405; **E**. *Phrynosoma douglasii* TxVP M-8526; **F**. *Iguana iguana* TxVP M-8454; **G**. *Ctenosaura hemilopha* TxVP M-8616; **H**. *Dipsosaurus dorsalis* TxVP M-13086; **I**. *Sauromalis ater* TxVP M-11599; **J**. *Anolis carolinensis* TxVP M-9042; **K**. *Crotaphytus collaris* TxVP M-12468; **L**. *Gambelia sila* TNHC 95261; **M**. *Leiocephalus carinatus* TNHC 89274; **N**. *Leiocephalus psammodromus* TNHC 103220; **O**. *Corytophanes* sp. TxVP M-16765; **P**. *Basiliscus vittatus* TxVP M-8556; **Q**. *Polychrus gutturosus* TNHC 24152; **R**. *Enyalioides heterolepis* UF 68015. Scale bars = 1 mm.

### Compound bone

#### Description

TxVP 41229-25853 is a left compound bone (fused prearticular, articular, and surangular), with only the anterior portion of the prearticular missing (Fig. 15). The adductor fossa is elongate. The dorsal face of the articular surface and the retroarticular process is coated in precipitant. The retroarticular process is narrow and elongate. There is an anteromedially-oriented angular process with a small lamina of bone connecting the retroarticular and angular processes. There is a medial crest (tympanic crest of McGuire 1996) that extends along the retroarticular process, and a distinct ventral ridge on the ventral surface of the process. The margin just lateral to the articular surface is devolved into a small boss. There is no lateral process nor a shelf connecting the body of the surangular to the tubercle (medial process of McGuire 1996) anterior to the articular surface. There is a distinct dentary articulation facet on the lateral surface. There is a foramen in the surangular just anterior to the adductor fossa and the surangular foramen in near the dorsal edge of the bone on the lateral surface. TxVP 41229-25613 is a dermarticular (fused articular and prearticular) that does not differ substantially from TxVP 41229-25853 in the features preserved.

#### Identification

Compound bones are referred to Pleurodonta based on the presence of an angular process and a posteriorly directed retroarticular process that is not broadened posteriorly (Estes et al. 1988; Fig. 21). Teiids also have a posteriorly directed retroarticular process that is not broadened posteriorly; however, the adductor fossa in teiids is greatly expanded for insertion of the m. adductor mandibulae posterior into Meckel’s canal (Estes et al. 1988) and the angular process is deflected further ventrally compared to pleurodontans (Tedesco et al. 1999; Morales et al. 2019). Among NA pleurodontans, the retroarticular process is reduced in *Polychrus* and the angular process is more horizontally oriented in *Dipsosaurus dorsalis* (Smith 2011) compared to crotaphytids. The medial crest is more distinct in *Dipsosaurus dorsalis*, *Iguana iguana*, *Ctenosaura similis*, and *Enyalioides* compared to crotaphytids (Smith 2011). *Sauromalus* differs from the fossil and crotaphytids in having a short triangular angular process (Smith 2011). In *Laemanctus* and *Corytophanes*, the angular process is short, and the angular process in *Basiliscus basiliscus* does not project as far medially compared to *Crotaphytus collaris* (see fig. 71 in Smith 2011). *Leiocephalus* differs from the fossil and crotaphytids in having a wider and rounded retroarticular process (Bochaton et al. 2021). In phrynosomatids and in *Anolis*, the angular process is connected to the retroarticular process by a sheet of bone that is more extensive than seen in examined crotaphytid specimens; however, one large specimen of *Sceloporus clarkii* (TxVP M-12202) has a reduced connection between the angular and retroarticular process. That specimen lacks a medial crest and a relatively wide retroarticular process, distinguishing it from examined crotaphytids. Fossils are assigned to Crotaphytidae based on an elongate, thin retroarticular process, an elongate and ventrally slanted angular process, a short medial crest on the retroarticular process, and a reduced lamina of bone connecting the angular process and retroarticular process. *Crotaphytus* differs from *Gambelia* in having a distinct knob-like process just anterolateral to the articular surface (lateral process of McGuire 1996) and having a more reduced shelf of bone connecting the body of the surangular to the dorsal tubercle anterior to the articular surface (McGuire 1996). A distinct knob-like process just anterolateral to the articular surface was not observed on some examined specimens of *Crotaphytus collaris*, so this feature likely varies intra- or interspecifically within *Crotaphytus*. A shelf of bone anterior to the dorsal tubercle is absent in most specimens of *Crotaphytus*, and a small shelf is present in specimens of *Gambelia wislizenii*. A small shelf is also reported in other species of *Crotaphytus* (McGuire 1996) so the presence of a shelf alone cannot be used to diagnose a fossil to *Gambelia*; however, only *Crotaphytus* lack the shelf (McGuire 1996). A distinct ridge on the dorsolateral surface of the surangular reportedly occurs in *Crotaphytus* and is reduced or absent in *Gambelia* (McGuire 1996). We could not identify a distinct ridge on the dorsolateral surface of the surangular in examined specimens of *Crotaphytus*, so this feature likely varies intra- or interspecifically within *Crotaphytus* and may be related to the development of the m. adductor mandibularis externus (McGuire 1996).

**Figure 21.**
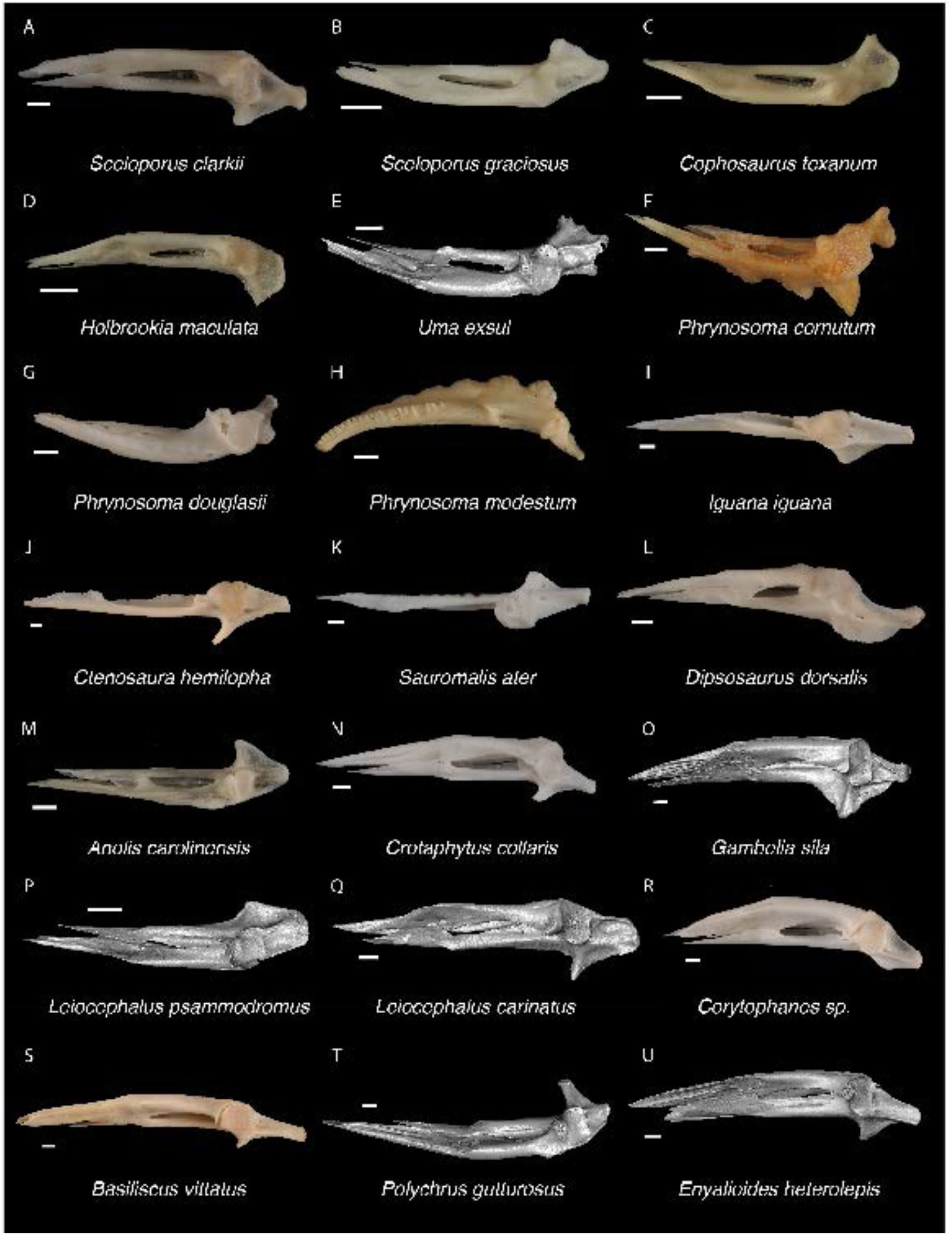
Pleurodontan Compound bones. Compound bones in dorsal view– **A**. *Sceloporus clarkii* TxVP M-12157; **B**. *Sceloporus graciosus* TxVP M-14879; **C**. *Cophosaurus texanum* TxVP M-8527; **D**. *Holbrookia maculata* TxVP M-14322; **E**. *Uma exsul* TNHC 30247; **F**. *Phrynosoma cornutum* TxVP M-6405; **G**. *Phrynosoma douglasii* TxVP M-8526; **H**. *Phrynosoma modestum* TNHC 95921; **I**. *Iguana iguana* TxVP M-8454; **J**. *Ctenosaura hemilopha* TxVP M-8616; **K**. *Sauromalis ater* TxVP M-11599; **L**. *Dipsosaurus dorsalis* TxVP M-13086; **M**. *Anolis carolinensis* TxVP M-9042; **N**. *Crotaphytus collaris* TxVP M-12468; **O**. *Gambelia sila* TNHC 95261; **P**. *Leiocephalus psammodromus* TNHC 103220; **Q**. *Leiocephalus carinatus* TNHC 89274; **R**. *Corytophanes* sp. TxVP M-16765; **S**. *Basiliscus vittatus* TxVP M-8554; **T**. *Polychrus gutturosus* TNHC 24152; **U**. *Enyalioides heterolepis* UF 68015. Scale bars = 1 mm.

**Phrynosomatidae Fitzinger, 1843**

**Referred specimens**: See Table S3.

### Maxilla

#### Description

TxVP 41229-27461 and TxVP 41229-27044 serve as the basis for our description (Fig. 22). TxVP 41229-27461 is a right maxilla with 22 tooth positions. Teeth are tricuspid except for the mesial teeth, and teeth are slender throughout the tooth row with bases near equal in width compared to the crowns. The facial process is broken dorsally and is curved medially with a distinct canthal crest. The facial process diminishes and merges with the crista transversalis, which extends anteromedially. There is a depression dorsally on the premaxillary process. The palatine process is present but broken. Dorsally the postorbital process has a groove that widens posteriorly for articulation with the jugal. Laterally there is a longitudinal ridge on the postorbital process. There is a superior alveolar foramen medial to the palatine process. There are three foramina on the premaxillary process, one anterior to the facial process (anterior inferior alveolar foramen), one on the crista transversalis (anterior inferior alveolar foramen), and one anteromedially on the premaxillary process (subnarial arterial foramen). There are eight lateral nutrient foramina. TxVP 41229-27044 differs in having 17 tooth positions, a less defined dorsal depression on the premaxillary process, no longitudinal ridge on the postorbital process, a flat dorsal surface of the postorbital process, a distinct asymmetric palatine process, two superior alveolar foramina, and four lateral nutrient foramina.

**Figure 22.**
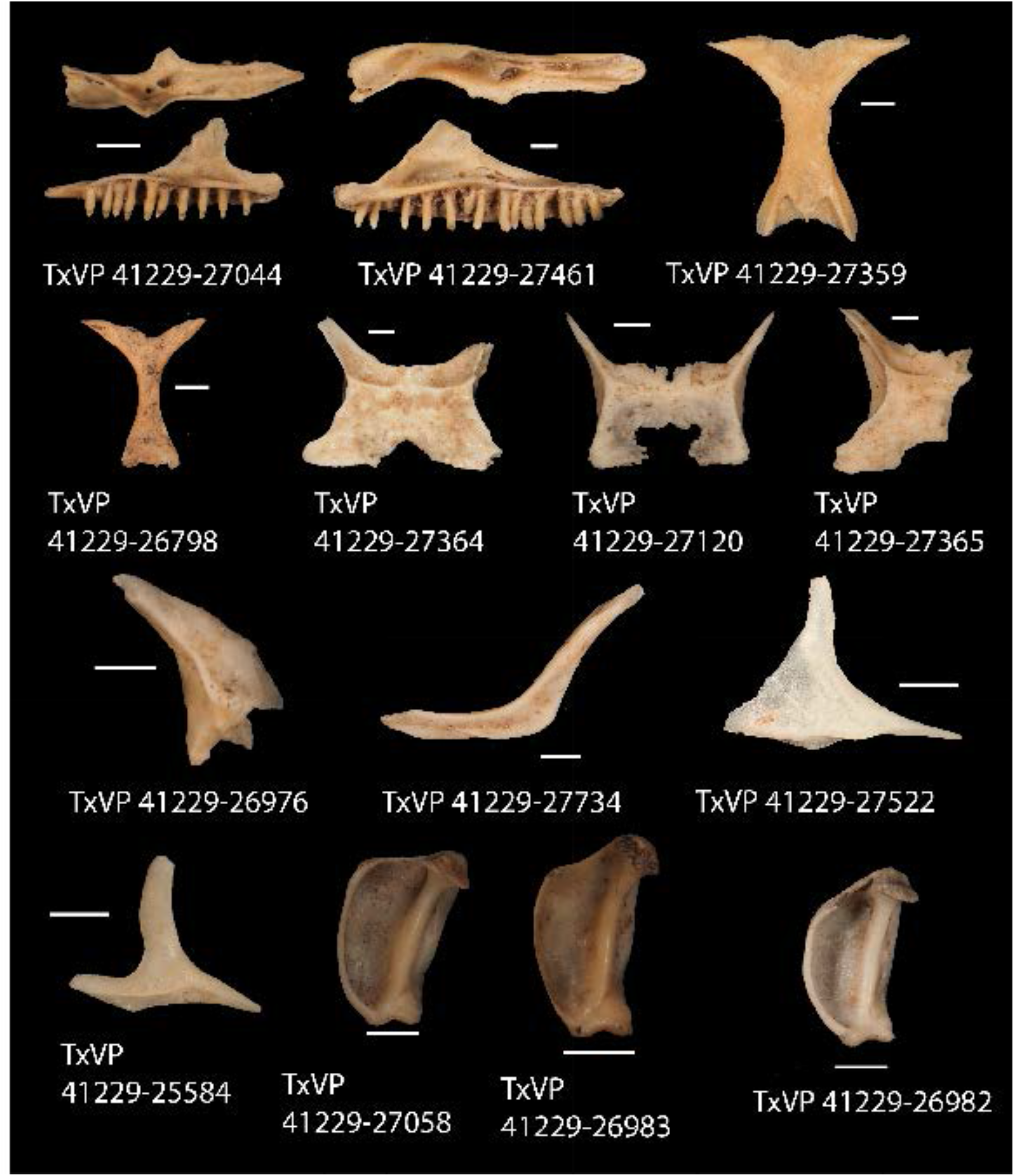
Fossil phrynosomatids. TxVP 41229-27044 Dorsal and medial view of left maxilla; TxVP 41229-27461 Dorsal and medial view of right maxilla; TxVP 41229-27359 Dorsal view of frontal; TxVP 41229-26798 Dorsal view of frontal; TxVP 41229-27364 Dorsal view of parietal; TxVP 41229-27120 Dorsal view of parietal; TxVP 41229-27365 Dorsal view of parietal; TxVP 41229-26976 Lateral view of right prefrontal; TxVP 41229-27734 Lateral view of left jugal; TxVP 41229-27522 Dorsolateral view of right postorbital; TxVP 41229-25584 Dorsolateral view of right postorbital; TxVP 41229-27058 Posterior view of left quadrate; TxVP 41229-26983 Posterior view of left quadrate; TxVP 41229-26982 Posterior view of left quadrate. Scale bars = 1 mm.

#### Identification

Maxillae were assigned to Pleurodonta based on the presence of pleurodont tricuspid teeth, having two foramina on the premaxillary process, and having a medially folded facial process with an anteroventally trending cathal crest (Smith 2009a). Phrynosomatids and the fossil maxillae differ from crotaphytids in having slender teeth throughout the tooth row with bases near equal in width compared to the crowns (Hollenshead and Mead 2006). Furthermore, the fossils differ from crotaphytids in having a facial process that curves anteromedially and reaches the medial edge of the premaxillary process, as opposed to a facial process that is more weakly folded and diminishes into a low ridge far from the medial edge of the premaxillary process (Smith 2009a). Fossil maxillae share with phrynosomatids, anoles, and *Polychrus* a relatively flat dorsal surface of the palatal plate (Smith 2009a). *Anolis* differs from phrynosomatids and the fossils in having an anteroposteriorly extended facial process with a canthal crest closer to horizontal, and *Polychrus* differs in having a facial process medially folded only at the dorsal tip (Smith 2009a).

### Frontal

#### Description

TxVP 41229-27359 and TxVP 41229-26798 serve as the basis for our description (Fig. 22). There is a small amount of sculpting on the posterodorsal surface, and the bone is slightly concave ventrally. Anteriorly there are lateral prefrontal facets and two distinct dorsal nasal facets defined by distinct anterolateral processes and separated by a smaller anteromedial process. The interorbital margins are waisted and the posterolateral processes flare laterally. The posterior edge is wavy and has a wide midline notch. There are small postfrontal facets laterally on the posterolateral processes and distinct parietal facets posteriorly. The cristae cranii are short, but approach one another in the interorbital region, and bound an indistinct groove for attachment of the solium supraseptale. The cristae cranii diverge posteriorly and extend along the lateral margins of the ventral surface. TxVP 41229-26798 differs in having less distinct nasal facets, minute cristae cranii, a larger notch in the posterior edge, and an especially thin interorbital region.

#### Identification

Fossil frontals share with Pleurodonta and Teiioidea a fused frontal with reduced cristae cranii (Estes et al. 1988). Fossil frontals differ from teiids in having strongly constricted interorbital margins of the frontal that strongly curves posterolaterally (Estes et al. 1988). Fossils differ from gymnophthalmoids in lacking frontal tabs (Estes et al. 1988). *Mabuya* and *Scincella* also have a fused frontal with reduced cristae cranii (Greer 1970; Jerez et al. 2015) but differ from the fossils in having a well-developed anteromedial process on the frontal and a relatively wider anterior end. The fossils and phrynosomatids differ from many iguanids (see above) in lacking a parietal foramen enclosed within the frontal, and differ from *Anolis, Polychrus*, and some *Leiocephalus* (e.g., *L. carinatus*) in having more slender interorbital margins (Pregill 1988; Pregill 1992; Smith 2011). Examined *Liocephalus* have a small, narrow notch at the posterior margin of the frontal (see also Bochaton et al. 2021) that is not found in the fossils or phrynosomatids. *Enyalioides heterolepis* differs from the fossil in having a bumpy, knob-like texture on the dorsal surface of the frontal. *Ctenosaura* and adult *Iguana* have wider interorbital margins compared to phrynosomatids, but the interorbital margins of juvenile *Iguana iguana* are comparable to that of larger *Sceloporus* and only differ in having more slender posterolateral processes (Bochaton et al. 2016a). Fossil frontals differ from crotaphytids in lacking a distinctly triradiate anterior end of the frontal. Additionally, smaller fossil frontals differ from crotaphytids in having a wider concave margin in the posterior edge, and larger fossils more closely resemble *Sceloporus* in having wider interorbital margins of the frontal compared to similar sized crotaphytids.

### Parietal

#### Description

TxVP 41229-27364 and TxVP 41229-27120 serve as the basis for our description (Fig. 22). TxVP 41229-27364 is a parietal with some sculpturing on the dorsal surface. The left anterolateral process and distal portions of the postparietal processes are broken. There is a large midline notch on the anterior edge. The adductor crests do not meet posteriorly, and the parietal table has a trapezoidal appearance. The ventrolateral crests are short and obscured in dorsal view by the adductor crests. The anterolateral process flares laterally. The posterior edge between the postparietal processes is characterized by two distinct depressions (nuchal fossae). The postparietal process has a dorsal crest that slants laterally. The ventral surface has shallow depressions (cerebral vault) divided by a low ridge. There is a deep pit for the processus ascendens just anterior to the posterior edge. TxVP 41229-27120 differs in lacking laterally flared anterolateral processes, having a large fontanelle on the anteromedial portion of the parietal, having adductor crests that do not cover the ventrolateral crests in dorsal view, lacking dorsal crests on the postparietal processes, and having a shallow pit for the processus ascendens. TxVP 41229-27120 has a notch on the posterior edge, but this is likely due to erosion of the bone because there are many pitted areas.

#### Identification

Parietals are assigned to Pleurodonta based on the presence of a fused parietal (Estes et al. 1988), the absence of co-ossified osteoderms (Estes et al. 1988; Evans 2008), the absence of parietal foramen fully enclosed by the parietal (Estes et al. 1988), and the absence of distinct ventrolateral crests (=parietal downgrowths of Estes et al. 1988). Parietals are assigned to Phrynosomatidae based on the presence of a large unossified anteromedial portion of the parietal around the location of the parietal foramen (Evans 2008), which is largely ossified in other NA pleurodontans. A few *Crotaphytus* (e.g., *C*. *collaris* TxVP M-8354) do have a large unossified anteromedial portion of the parietal, but in *Crotaphytus*, the ventrolateral crests are oriented more laterally. A large unossified anteromedial portion of the parietal was noted to occur in smaller-bodied phrynosomatid genera, such as many phrynosomatines (Evans 2008), but we observed this feature in smaller species of *Sceloporus* (e.g., *S. occidentalis*) and *Uta* (see also Maisano 2002a) as well.

### Prefrontal

#### Description

TxVP 41229-26976 is a right prefrontal (Fig. 22). It is triradiate with a long and pointed orbital process, a short ventral process, and an anterior main body developed into a sheet of bone. The anterior sheet is slightly broken and has a broad articulation facet for the facial process of the maxilla. There is a distinct ridge on the lateral surface near the base of the orbital process that is continuous with a large lateral boss. The ventral process is narrow and squared-off. There is a distinct notch for the lacrimal foramen, and the ventral process forms the posterior border of the foramen. Medially, the boundary of the olfactory chamber is a smooth, rounded, and concave surface. Dorsal to the olfactory chamber is a shallow groove for articulation with the frontal. The orbitonasal flange is narrow.

#### Identification

The fossil shares with pleurodontans and teiids the presence of a distinct prefrontal boss (Gauthier et al. 2012; Fig. 23). The fossil and prefrontals of many pleurodontans differ from teiids in having a boss that widens ventrally and is semicircular or tear-dropped shaped (Smith 2009a). Additionally, teiids differ in having a thin, laterally projecting lamina with a distinct articulation facet for the facial process of the maxilla (Tedesco et al. 1999). *Basiliscus*, *Polychrus*, and *Anolis* differ from the fossil in often having a more prominent boss or a strong lateral canthal ridge (Smith 2011). The prefrontal boss is larger in adult *Cyclura* and *Iguana* compared to the fossil (Smith 2011), and juvenile *Iguana* differ in having an exceptionally thin ventral process. *Corytophanes* and *Laemanctus* differ from the fossil in having rugosities on the dorsal surface (Smith 2011). Furthermore, *Corytophanes* differs from the fossil in having a long supraorbital process, and *Laemanctus* differs in having a posterior process that is directed more posteriorly (Smith 2011). *Crotaphytus* differs from the fossil in having the prefrontal boss slightly obscure the lacrimal notch in lateral view. Examined *Gambelia* differ in having a more distinct boss that projects farther laterally. The ridge dorsal to the prefrontal boss is wider in examined *Leiocephalus* compared to the fossil. *Enyalioides heterolepis* differs from the fossil in having a more bulbous prefrontal boss. The fossil differs from other NA pleurodontans and shares several features with phrynosomatids excluding *Phrynosoma*, which has a long supraorbital process, and phrynosomatines that have a prefrontal boss developed into a laterally projecting thin lamina (e.g., *Cophosaurus* and *Holbrookia*).*heterolepis*

**Figure 23.**
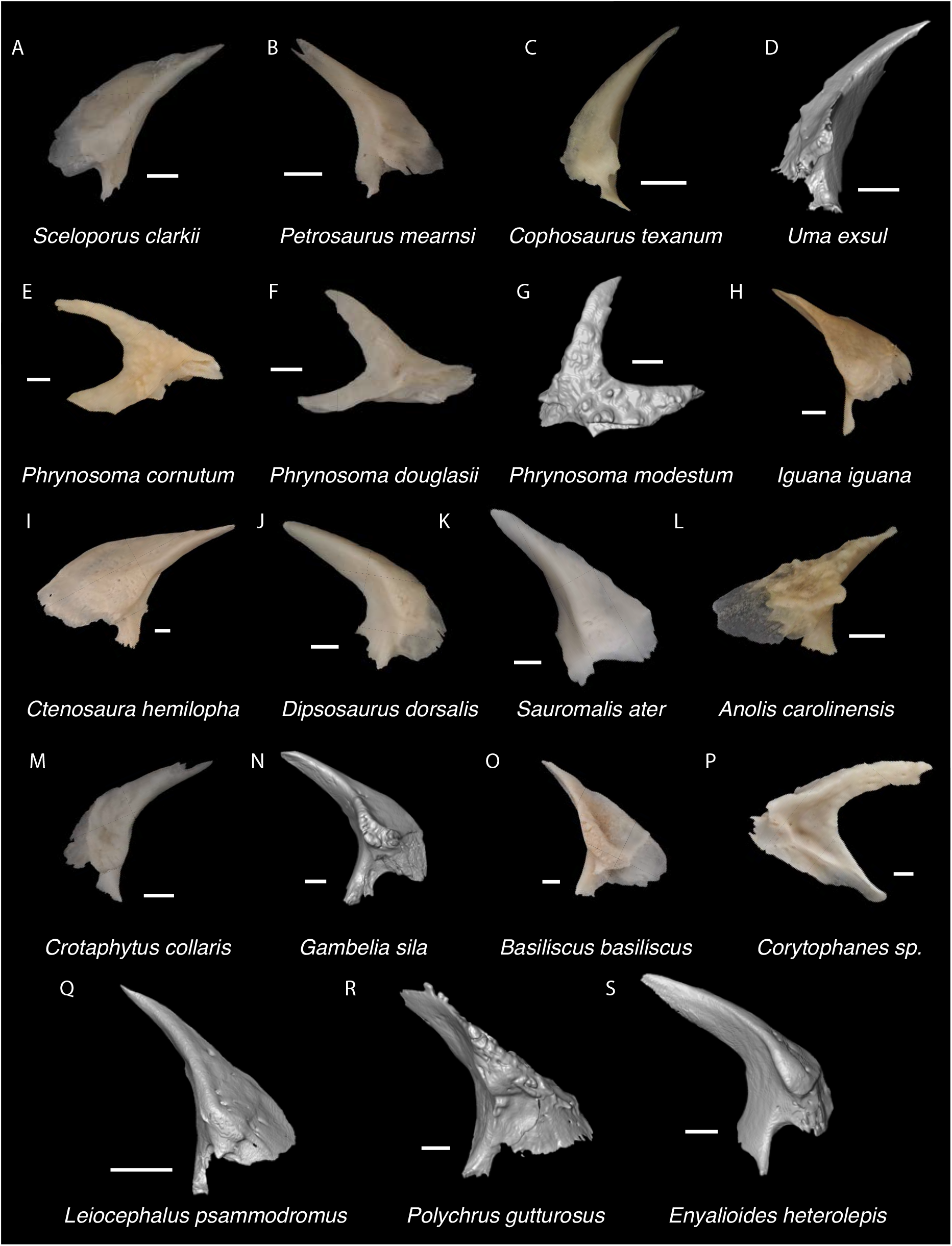
Pleurodontan Prefrontals. Prefrontals in lateral view– **A**. *Sceloporus clarkii* TxVP M-12157; **B**. *Petrosaurus mearnsi* TxVP M-14910; **C**. *Cophosaurus texanum* TxVP M-8527; **D**. *Uma exsul* TNHC 30247; **E**. *Phrynosoma cornutum* TxVP M-6405; **F**. *Phrynosoma douglasii* TxVP M-8526; **G**. *Phrynosoma modestum* TNHC 95921; **H**. *Iguana iguana* TxVP M-13054; **I**. *Ctenosura hemilopha* TxVP M-8616; **J**. *Dipsosaurus dorsalis* TxVP M-13086; **K**. *Sauromalis ater* TxVP M-11599; **L**. *Anolis carolinensis* TxVP M-9042; **M**. *Crotaphytus collaris* TxVP M-12468; **N**. *Gambelia sila* TNHC 95261; **O**. *Basiliscus basiliscus* TxVP M-11907; **P**. *Corytophanes* sp. TxVP M-16765; **Q**. *Leiocephalus psammodromus* TNHC 103220; **R**. *Polychrus gutturosus* TNHC 24152; **S**. *Enyalioides heterolepis* UF 68015. Scale bars = 1 mm.

### Jugal

#### Description

TxVP 41229-27734 is a left jugal (Fig. 22). There is a maxillary facet laterally and ventrally on the suborbital process. The dorsal margin of the suborbital process is everted laterally and there is a groove on the medial surface. There is no quadratojugal process. There is a ridge on the anteromedial edge of the postorbital process and a postorbital facet anteriorly. The postorbital process is posteriorly directed and concave posteriorly. There is a foramen medially near the inflection point, two foramina anteriorly on the postorbital process, and many small foramina along the lateral surface. TxVP 41229-27968 differs in having an elongate depression dorsally on the suborbital process, a tall lateral edge of the suborbital process with an upturned corner at the anterior end, and a postorbital process that curves posteromedially.

#### Identification

Jugals were assigned to Pleurodonta based on the absence of a quadratojugal process (Gauthier et al. 1988; Smith 2009a). *Xantusia* also lack a quadratojugal process but differ from pleurodontans in having an exceptionally short and thin suborbital process (Young 1942). Among NA pleurodontans, the fossil shares with Phrynosomatidae and Crotaphytidae a posteriorly deflected distal end of the postorbital process (Smith 2009a); however, crotaphytids differ in having a quadratojugal process.

### Postorbital

#### Description

Morphotype A: TxVP 41229-27522 is a right postorbital (Fig. 22). The fossil is triradiate with thin anterior process and dorsal processes and a wider posterior process. The anterior and dorsal processes are relatively straight. There is a small articulation facet anteromedially on the orbital process and there are distinct jugal and squamosal facets along the ventrolateral edge.

Morphotype B: TxVP 41229-25584 is a right postorbital (Fig. 22). The bone is triradiate with thin anterior, dorsal and posterior processes. The ventrolateral margin is concave below the anterior portion and the posterior process is slightly curved dorsally. The dorsal process is concave anteriorly without an anterior articulation facet. There are distinct jugal and squamosal facets that are visible in dorsolateral view.

#### Identification

Postorbitals were identified to Pleurodonta based on a sub-triangular morphology with a distinct ventral process (Estes et al. 1988). The fossils share with phrynosomatids, *Anolis*, some iguanids, and some *Leiocephalus* a smooth lateral face of the dorsal process (Smith 2009a, b; Smith 2011). The posterior process in *Ctenosaura* (see also fig. 40G of Smith 2011) is proportionally more elongated compared to the fossils. *Dipsosaurus dorsalis* differs in often having a flangelike expansion on the orbital process (Smith 2011). *Iguana* generally has a wider orbital process compared to the fossil and phrynosomatids excluding *Phrynosoma*. The postorbital of *Sauromalus* and *Dipsosaurus dorsalis* is more straight in anterior view compared to the fossils and phrynosomatids (Smith 2011). *Anolis* differs from the fossils and phrynosomatids in having an articulation facet visible on the lateral or anterolateral surface of the orbital process as it underlaps the frontoparietal corner (Smith 2009a). The postorbital of *Leiocephalus* differs from the fossils in lacking a distinct squamosal facet along the ventrolateral edge (see also fig. 3F of Bochaton et al. 2021). The postorbitals of examined *Uta*, *Urosaurus*, *Cophosaurus*, and *Callisaurus* have a slender build relative to *Sceloporus* and are more similar to Morphotype B. Sand lizards differ from sceloporines in lacking a postfrontal (Cox and Tanner 1977) and therefore also lack a postfrontal articulation surface on the postorbital. Fossils identified to morphotype A likely represent *Sceloporus* or *Petrosaurus* and fossils identified to morphotype B likely represent sand lizards.

### Pterygoid

#### Description

TxVP 41229-27488 and TxVP 41229-27011 serve as the basis for our description (Fig. 24). TxVP 41229-27488 is a left pterygoid that is missing the distal end of the transverse process. The palatine process is broad at the base and has a palatine facet anteriorly. The transverse process extends laterally at nearly a 90-degree angle with the quadrate process. The transverse process is dorsoventrally tall and bears a distinct ectopterygoid facet. There is a distinct ridge on the dorsal surface for insertion of the superficial pseudotemporal muscle. There is a large ridge on the ventral surface for insertion of the pterygomandibular muscle and a deep fossa columella without a pterygoid groove. The quadrate process is elongated, and the medial surface has a groove that serves for insertion of the pterygoideus muscle. There is a medial groove dorsal to a shelf-like projection at the floor of the basipterygoid fossa. There are no pterygoid teeth, but there is a large hole on the ventral surface of the palatal plate along with several smaller foramina on both the dorsal and ventral surface. TxVP 41229-27011 is smaller than TxVP 41229-27488 and the distal end of the transverse process is pointed and tall vertically.

**Figure 24.**
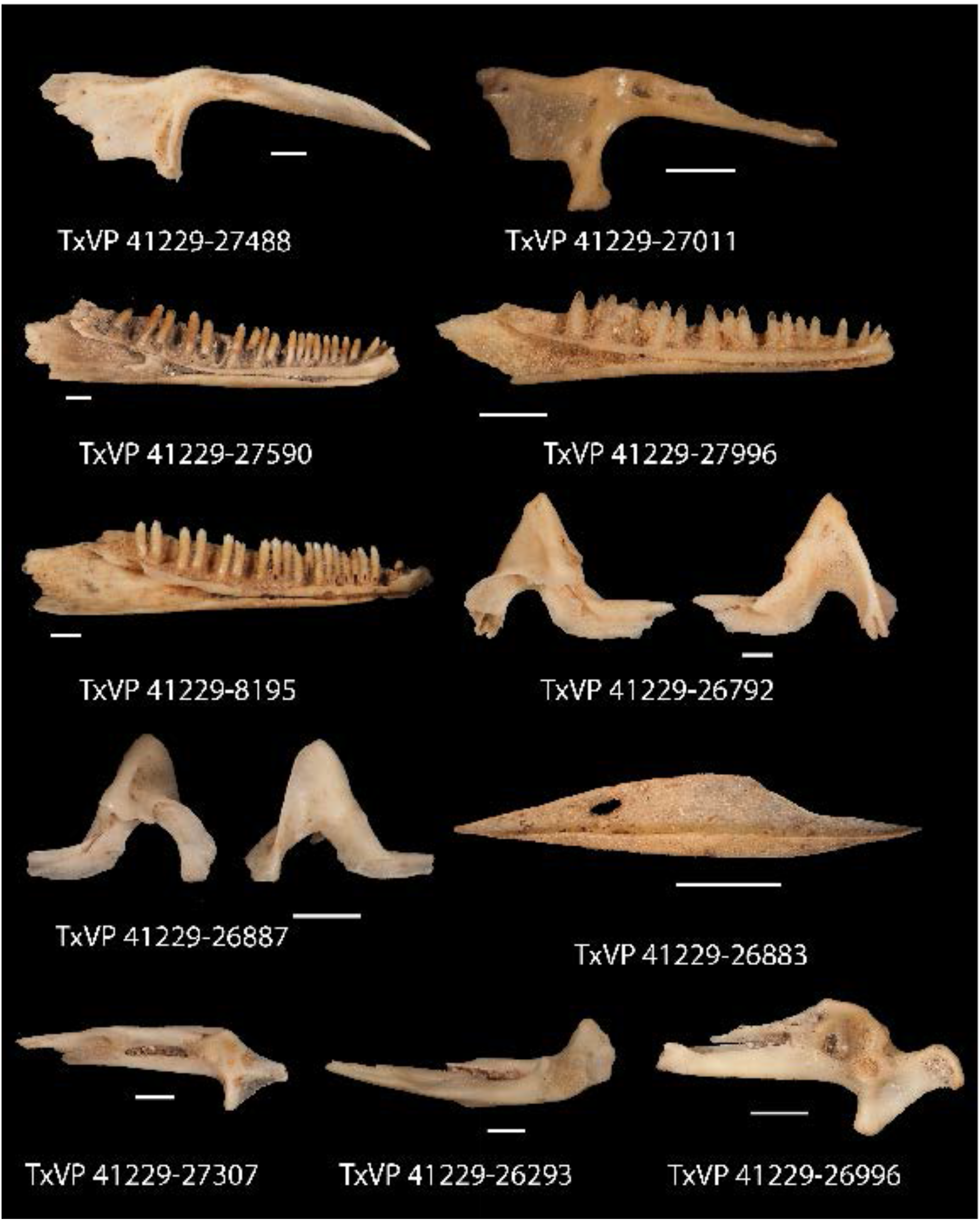
Fossil phrynosomatids. TxVP 41229-27488 Dorsal view of left pterygoid; TxVP 41229-27011 Dorsal view of left pterygoid; TxVP 41229-27590 Lateral view of left dentary; TxVP 41229-27996 Lateral view of left dentary; TxVP 41229-8195 Lateral view of left dentary; TxVP 41229-26792 Lateral and medial view of right coronoid; TxVP 41229-26887 Lateral and medial view of left coronoid; TxVP 41229-26883 Lateral view of left splenial; TxVP 41229-27307 Dorsal view of right compound bone; TxVP 41229-26293 Dorsal view of left compound bone; TxVP 41229-26996 Dorsal view of right compound bone. Scale bars = 1 mm.

#### Identification

Among examined NA lizards, only in pleurodontans is the transverse process oriented medially, creating a near right angle with the quadrate process. In other NA lizards, the transverse process is oriented anteromedially. This distinction in transverse process orientation likely only applies to NA lizards since at least some non-pleurodontans (e.g., scincids; Paluh and Bauer 2017) from other geographic regions share a medially oriented transverse process. Fossil pterygoids have medially oriented transverse process and were identified to Pleurodonta. Among NA pleurodontans, pterygoid teeth are absent in phrynosomatids and are variably present in *Anolis*, *Polychrus*, *Dipsosaurus*, and *Leiocephalus* (Etheridge and de Queiroz 1988; Smith 2009a). Pterygoid teeth were reported as present in *Sauromalus*, but we observed one specimen (*Sauromalis ater* TxVP M-9782) without pterygoid teeth. *Dipsosaurus dorsalis* differs from phrynosomatids in having a narrower palatal plate and a narrow linear notch on the transverse process. Compared with the fossils, *Sauromalus* has a narrower palatal plate and a taller quadrate process. *Leiocephalus* differs from the fossils in having a narrower palatal plate and a more distinct ventromedial projection at the floor of the basipterygoid fossa. Examined *Anolis* and *Polychrus* differ from the fossils and examined phrynosomatids in having a taller ridge for the insertion of the pterygomandibular muscle (Villa and Delfino 2019) on the ventral surface of the pterygoid extending from the transverse process to the basipterygoid fossa. In addition, examined *A. carolinensis* have a ventral projection near the basipterygoid fossa not seen in the fossils or phrynosomatids. Based on the differences from other NA pleurodontans, fossils were assigned to Phrynosomatidae.

### Dentary

#### Description

TxVP 41229-27590 and TxVP 41229-8195 serve as the basis for our description (Fig. 24). TxVP 41229-27590 is a left dentary with 26 tooth positions. Distal teeth are tricuspid and relatively slender. The suprameckelian and inframeckelian lips approach one another midway along the tooth row, but the Meckelian groove is open for its entire length. The suprameckelian lip is relatively short anteriorly. The dental shelf is narrow but widens slightly anteriorly. There is a distinct intramandibular lamella, and the intramandibular septum reaches to the level of the fourth most distal tooth position. There is a small posterior projection from the intramandibular septum. The posterior end is broken, but there is a coronoid facet with a dorsally projecting corner of bone. There are four nutrient foramina on the anterolateral surface of the bone. TxVP 41229-8195 differs from TxVP 41229-27590 in lacking the small posterior projection from the intramandibular septum, having a bifurcated posterior end, and having five nutrient foramina on the anterolateral surface.

#### Identification

Dentaries were placed in Pleurodonta based on pleurodont tricuspid teeth and a closed or partially closed Meckelian groove bounded by suprameckelian and inframeckelian lips (Scarpetta 2021). Among NA pleurodontans, the fossils differ from iguanids, dactyloids, extant *Leiocephalus*, and polychrotids, which all have a fused Meckelian groove (Smith 2011; Scarpetta 2021). *Basiliscus*, *Corytophanes*, and some *Laemanctus* differ from the fossils in having flared the tooth crowns (Smith 2009a; Scarpetta 2021). Some *Laemanctus* have a fused Meckelian groove (Smith 2011; Scarpetta 2021), but it is unfused in *Laemanctus serratus* (Lang1989; Smith 2009a). Hoplocercids, except besides *Enyalioides laticeps* (FMNH 206132), differ in having an open Meckelian groove that is not bounded by suprameckelian and inframeckelian lips (Frost and Etheridge 1989; Scarpetta 2021). Crotaphytids differ from the fossils in having a relatively tall suprameckelian lip on the anterior half of the dentary (Scarpetta 2021). Furthermore, *Crotaphytus* differs in having teeth that widen towards the base (Hollenshead and Mead 2006) and *Gambelia* differs in having sharper and more recurved mesial teeth (Scarpetta 2021). Based on these differences with other NA pleurodontans, fossils were assigned to Phrynosomatidae.

### Coronoid

#### Description

TxVP 41229-26792 and TxVP 41229-26887 serve as the basis for our description (Fig. 24). TxVP 41229-26792 is a right coronoid. The coronoid process is tall and pointed and the anteromedial process is elongated. The posteromedial process is missing the distal end, but the remaining portion is ventrally oriented. There is a distinct medial crest that extends from the coronoid process onto the posteromedial process. There is a small, rounded lateral process. There is a vertically oriented lateral crest that ends at the anterior margin of the lateral process. The ventral surface is characterized by a narrow concave facet for articulation with the surangular dorsally. The ventral surface also bears a small lateral groove. TxVP 41229-26887 differs from TxVP 41229-26792 in being smaller, having a more rounded coronoid process, and having a more distinct lateral crest.

#### Identification

Fossil coronoids share with some members of Pleurodonta and xantusiids the absence of a distinct anteriorolateral process (Estes et al. 1988). Xantusiids differ from the fossils and pleurodontans in having an anterior groove extending onto the coronoid process for articulation with the coronoid process of the spleniodentary. Fossils differ from iguanids, *Enyalioides heterolepis*, *Anolis*, and *Leiocephalus*, which all have a anterolateral process (Smith 2011; Bochaton et al. 2021). The fossils differ from corytophanids (see also fig. 69 of Smith 2011) but share with phrynosomatids and crotaphytids a lateral crest that merges with the anterior margin of the lateral process. Additionally, corytophanids differ in having the apex of the coronoid process posterodorsally deflected to a greater degree compared to the fossils (see also fig. 69 of Smith 2011). Fossils differ from crotaphytids in having a narrow facet on the ventral surface of the coronoid for articulation with the dorsal surface of the surangular. Moreover, *Crotaphytus* differs from the fossils in having more extensive lamina of bone posterior to the coronoid process to articulate with the surangular dorsally and *Gambelia*, except for *G. sila*, differs in having a posteromedial process directed more posteriorly (Norell 1989; McGuire 1996). Based on these differences with respect to other NA pleurodontans, fossils were assigned to Phrynosomatidae.

### Splenial

#### Description

TxVP 41229-26883 is a left splenial (Fig. 24). It is slender with elongate anterior and posterior processes. There is a large anterior inferior mylohyoid foramen that is positioned anterodorsal to the smaller anterior mylohyoid foramen. There is a medial shelf that curls dorsally and obscures the anterior mylohyoid foramen in medial view.

#### Identification

The fossil shares an enclosed anterior inferior alveolar foramen with pleurodontans, teiids, some gymnophthalmoids, and some anguids (Evans 2008; Gauthier et al. 2012; Morales et al. 2019; Fig. 25). Teiinae differ from the fossil and other North American lizards in having an inferior alveolar foramen that is larger and posterodorsally located relative to the anterior mylohyoid foramen (Tedesco et al. 1999; Fisher and Tanner 1970; Gauthier et al. 2012; Fig. 26). Anguids usually have an anterior inferior alveolar foramen that is not entirely enclosed by the splenial, but some individuals have been observed with a completely enclosed foramen (Bochaton et al. 2016b; Ledesma et al. 2021). Many anguids differ from the fossil and pleurodontans in having the posterior end of the splenial bifurcated into a distinct dorsal and ventral process (Ledesma and Scarpetta 2018). There is substantial variation in splenial morphology among gymnophthalmoids. Some gymnopthhalmoids differ from the fossil in having an anteroposteriorly short splenial (Hoyos 1998; Evans 2008; Yanez-Munoz et al. 2021). Other gymnophthalmoids (e.g., *Gymnophthalmus speciosus*) differ from the fossil in lacking an anterior inferior alveolar foramen fully enclosed within the splenial (Evans 2008). Some gymnophthalmoids (e.g., see fig. 7 of *Ptychoglossus vallensis* in Morales et al. 2019) resemble the fossil, but are excluded based on geography. The fossil splenial is identified to Pleurodonta.

**Figure 25.**
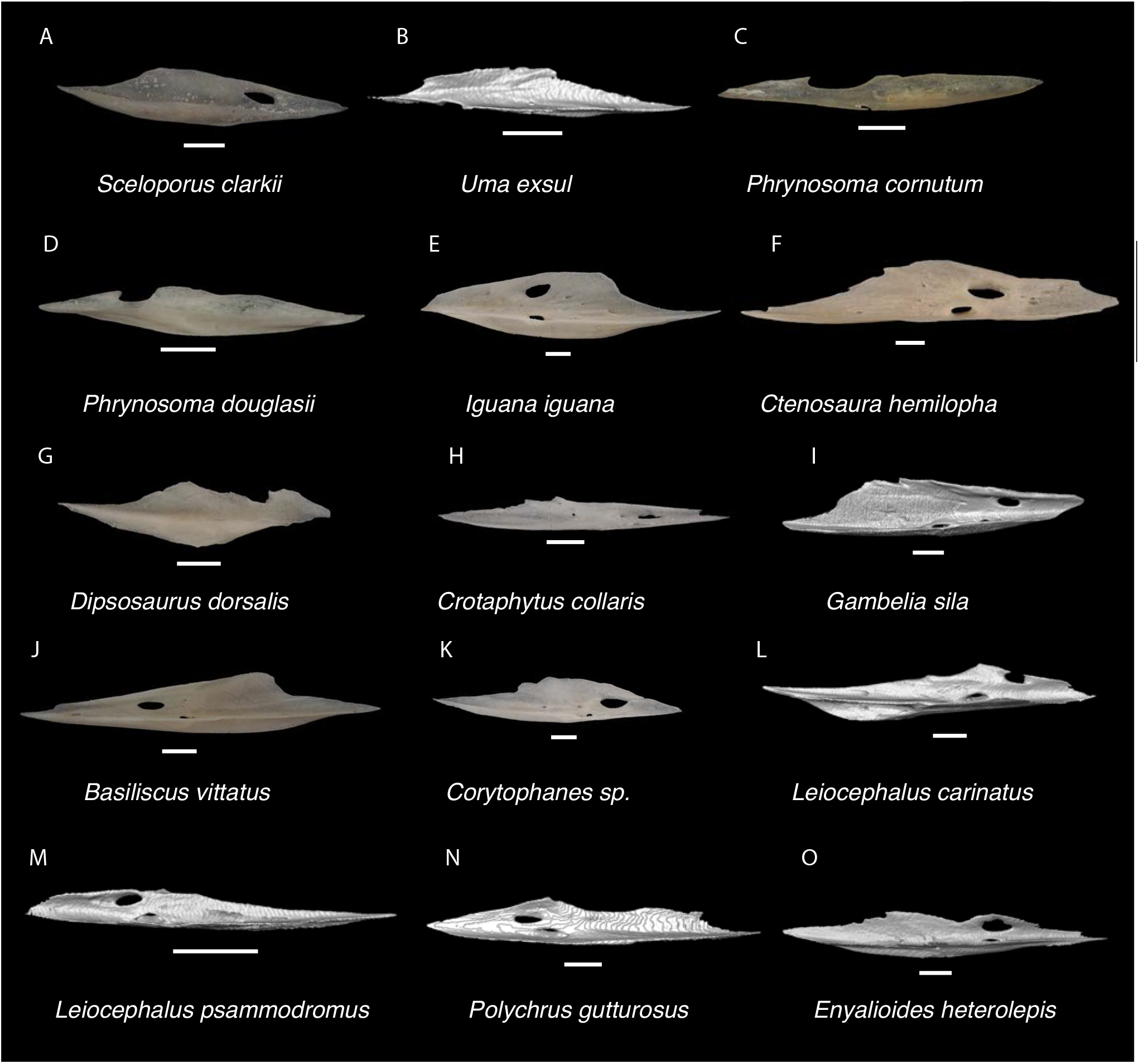
Pleurodontan Splenials. Splenials in lateral view– **A**. *Sceloporus clarkii* TxVP M-12157; **B**. *Uma exsul* TNHC 30247; **C**. *Phrynosoma cornutum* TxVP M-6405; **D**. *Phrynosoma douglasii* TxVP M-8526; **E**. *Iguana iguana* TxVP M-8454; **F**. *Ctenosura hemilopha* TxVP M-8616; **G**. *Dipsosaurus dorsalis* TxVP M-13086; **H**. *Crotaphytus collaris* TxVP M-12468; **I**. *Gambelia sila* TNHC 95261; **J**. *Basiliscus vittatus* TxVP M-8556; **K**. *Corytophanes* sp. TxVP M-16765 **L**. *Leiocephalus carinatus* TNHC 89274; **M**. *Leiocephalus psammodromus* TNHC 103220; **N**. *Polychrus gutturosus* TNHC 24152; **O**. *Enyalioides heterolepis* UF 68015. Scale bars = 1 mm.

**Figure 26.**
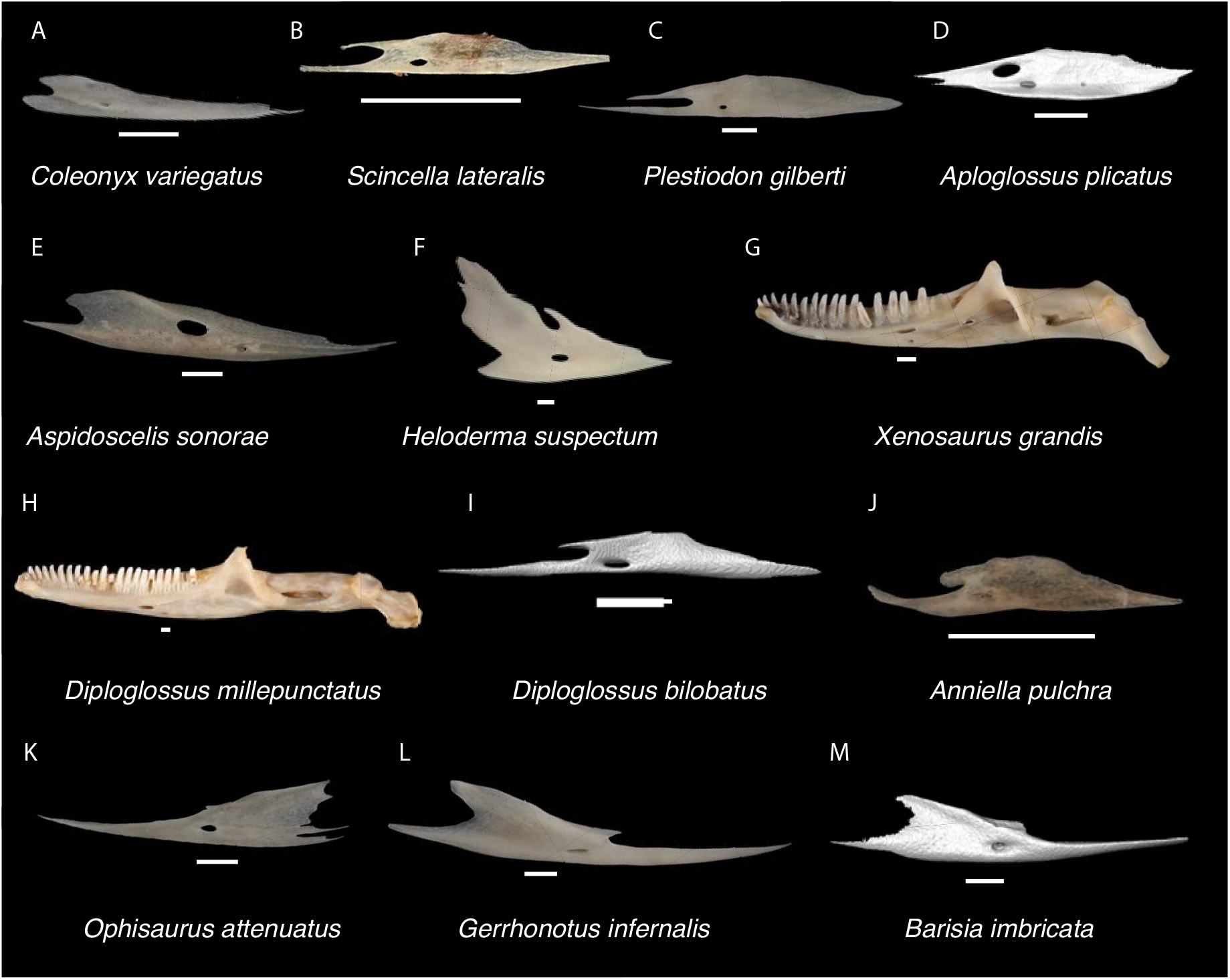
Non-Pleurodontan Splenials. Splenials in medial view– **A**. *Coleonyx variegatus* TxVP M-12109; **B**. *Scincella lateralis* TxVP M-5531; **C**. *Plestiodon gilberti* TxVP M-8587; **D**. *Aploglossus plicatus* 34481; **E**. *Aspidoscelis sonorae* TxVP M-15670; **F**. *Heloderma suspectum* TxVP M-9001; **G**. *Xenosaurus grandis* TxVP M-8960; **H**. *Diploglossus millepunctatus* TxVP M-9010; **I**. *Diploglossus bilobatus* TNHC 31933; **J**. *Anniella pulchra* TxVP M-8678; **K**. *Ophisaurus attenuatus* TxVP M-8979; **L**. *Gerrhonotus infernalis* TxVP M-13441; **M**. *Barisia imbricata* TNHC 76984. Scale bars = 1 mm.

The splenial of crotaphytids is much more slender and elongate compared to that of the fossil. The prearticular crest in *Iguana iguana*, *Ctenosaura*, *Dipsosaurus dorsalis*, *Sauromalus*, *Corytophanes*, and *Basiliscus vittatus* is flat, whereas it curves dorsally in the fossil and in most examined *Sceloporus* (except for some *Sceloporus jarrovii*) such that the anterior mylohyoid foramen is obscured in lateral view. *Iguana* (Conrad and Norell 2010) and *Polychrus marmoratus* UF 65135 differ from the fossil in having the anterior inferior alveolar foramen positioned directly dorsal to the anterior mylohyoid foramen. *Polychrus gutturosus* differs in having a relatively shorter splenial along the posterior half of the bone. The splenials of *Leiocephalus* and *Enyalioides heterolepis* differ from the fossil in being relatively longer and more slender. *Anolis* differs in either lacking a separate and distinct splenial bone or having a short splenial (Etheridge and de Queiroz 1988). Based on these differences with other NA pleurodontans, fossils were assigned to Phrynosomatidae.

### Compound bone

#### Description

Morphotype A: TxVP 41229-27307 serves as the basis for our description and is a right compound bone that is missing the distal end of the prearticular (Fig. 24). The adductor fossa is elongate and open. The retroarticular process narrows posteriorly and is squared off at the posterior end. There is a medially oriented angular process with a lamina of bone connecting the retroarticular and angular processes. There is a medial crest (tympanic crest of McGuire 1996) that extends longitudinally along the dorsal surface of the retroarticular process, and a longitudinal ridge on the ventral surface of the process. The medial process is tall. There is a small crest dorsally on the surangular and a distinct dentary articulation facet on the lateral surface. There are two posterior surangular foramina, one lateral to the medial process and the other slightly ventral, and one anterior surangular foramen lateral to the crest on the surangular. TxVP 41229-26996 is an unusual dermarticular with a lateral notch on the retroarticular process. The margin just lateral to the articular surface is developed into a small boss.

Morphotype B: TxVP 41229-26293 serves as the basis for our description (Fig. 24). TxVP 41229-26293 is a left compound bone that is missing the distal end of the prearticular. The adductor fossa is elongate and open. The retroarticular process is broad and flat. There is a posteromedially-oriented angular process with a low dorsal ridge. There is no distinct medial crest and a ventral ridge on the retroarticular process. The medial process is tall. There is a small crest dorsally on the surangular and distinct dentary articulation facet on the lateral surface.

#### Identification

Fossils are referred to Pleurodonta based on having a posteriorly directed retroarticular process that is not broadened posteriorly and that is lacking a widely opened adductor fossa (Estes et al. 1988). *Polychrus* differs from the fossils in having the retroarticular process reduced (Smith 2011). The angular process does not project as far medially in *Dipsosaurus dorsalis*, *Sauromalus*, and *Iguana iguana* compared to the fossils (see also fig. 48 in Smith 2011). The medial crest tends to be more distinct in *Dipsosaurus dorsalis*, *Iguana iguana*, *Ctenosaura similis*, and *Enyalioides* compared to crotaphytids (Smith 2011). In *Laemanctus* and *Corytophanes*, the angular process is shorter compared to the fossils and in *Basiliscus basiliscus* the medial crest is more distinct and the retroarticular process is thinner (see fig. 71 in Smith 2011). Examined *Anolis* resemble fossils from Morphotype A in having a distinct angular process connected to the retroarticular process by a sheet of bone but differ in having a horizontally oriented posterior terminus of the retroarticular process. The posterior terminus of the retroarticular process is slightly angled in the fossils, similar to many phrynosomatids (Presch 1969). Additionally, examined *Anolis carolinensis* (also in *A. garmani*, see fig 25 in Smith 2011) have a broad depression on the dorsal surface of the surangular just anterior to the articular surface not seen in the fossils. Based on these differences with other NA pleurodontans, fossils were assigned to Phrynosomatidae.

**Sceloporinae Savage, 1958**

**Referred specimens**: See Table S3.

### Premaxilla

#### Description

TxVP 41229-29096 and TxVP 41229-27523 serve as the basis for our description (Fig. 27). TxVP 41229-29096 is a premaxilla with six unicuspid, widely spaced teeth. It has a slightly rounded rostral surface and a long nasal process with distinct lateral nasal articulation facets. There is an ossified bridge extending laterally from the nasal process and enclosing the medial ethmoidal foramen on the right side and an incomplete bridge of bone on the left side. There are small foramina just lateral to the base of the nasal process that pierce ventrally. There are shallow maxillary facets laterally on the alveolar plate. Posteriorly, the palatal plate is steeply slanted with distinct vomer articulation facets. The incisive process is short, squared-off, and slightly bilobed. TxVP 41229-27523 differs from TxVP 41229-29096 in having seven tooth positions, a narrow nasal process with a small anterior nasal facet on the left side, and in lacking an ossified bridge.

**Figure 27.**
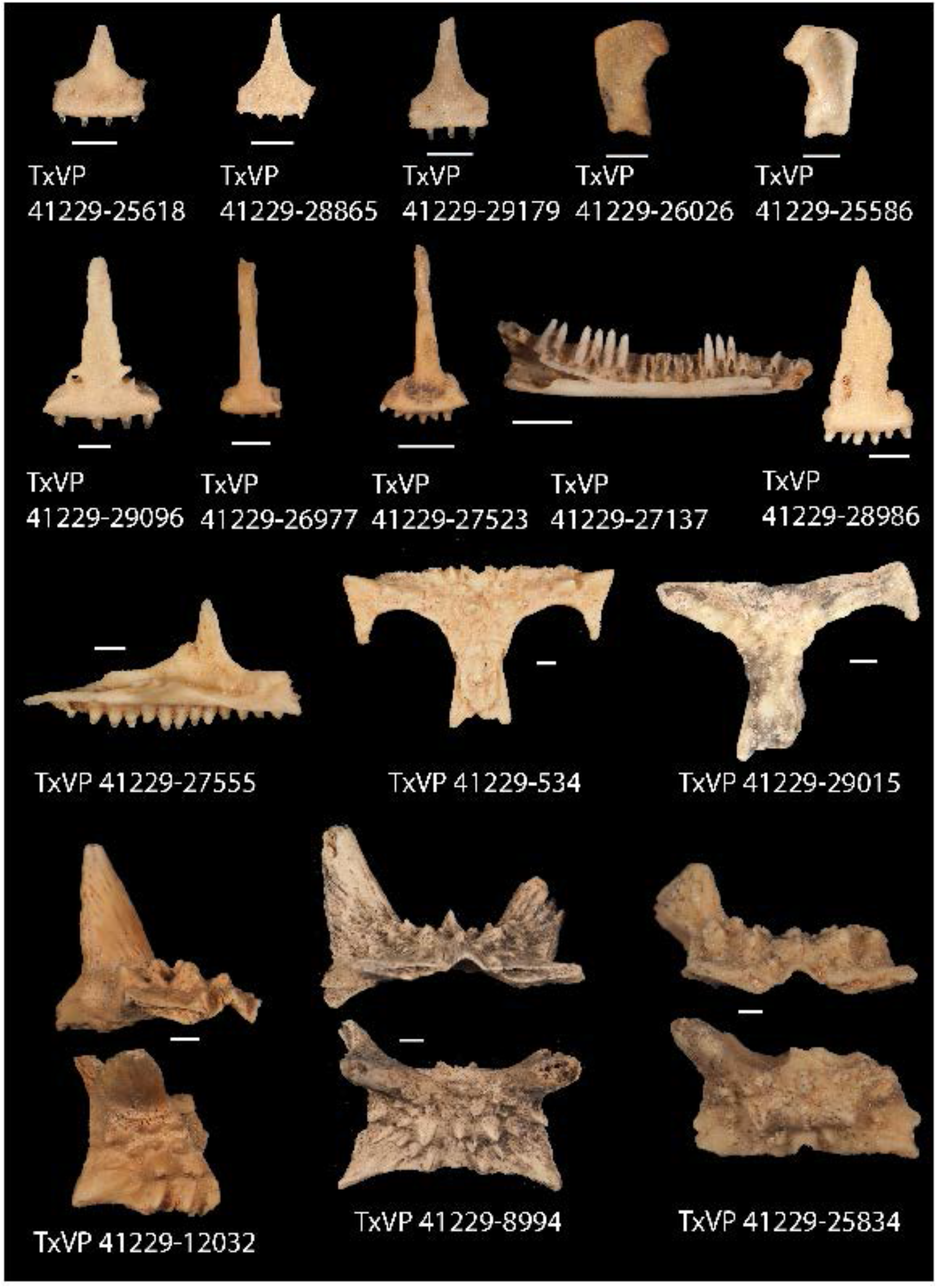
Fossil phrynosomatids. TxVP 41229-25618 Anterior view of premaxilla; TxVP 41229-28865 Anterior view of premaxilla; TxVP 41229-29179 Anterior view of premaxilla; TxVP 41229-26026 Posterior view of left quadrate; TxVP 41229-25586 Posterior view of right quadrate; TxVP 41229-29096 Anterior view of premaxilla; TxVP 41229-26977 Anterior view of premaxilla; TxVP 41229-27523 Anterior view of premaxilla; TxVP 41229-27137 Lateral view of left dentary; TxVP 41229-28986 Anterior view of premaxilla; TxVP 41229-27555 Medial view of left maxilla; TxVP 41229-534 Dorsal view of frontal; TxVP 41229-29015 Dorsal view of frontal; TxVP 41229-12032 Anterior and dorsal view of parietal; TxVP 41229-8994 Anterior and dorsal view of parietal; TxVP 41229-25834 Anterior and dorsal view of parietal. Scale bars = 1 mm.

#### Identification

Premaxillae are assigned to Pleurodonta based on having a fused premaxilla (Estes et al. 1988) with less than seven tooth positions (Smith 2009a). The seven tooth positions of TxVP 41229-27523 overlaps with other NA lizards including scincids, teiids, xantusiids, and *Anniella* (Evans 2008; Smith 2009a). Scincids differ in having an unfused premaxilla (Gauthier et al. 1988), teiids differ in lacking an incisive process (Gao and Norell 1998; Conrad 2008; Morales et al. 2019), xantusiids differ in having a much thinner nasal process and a broad, flat palatal plate (Savage 1963), and *Anniella* have a more posteriorly directed nasal process (Toerien 1950; Evans 2008). Among NA pleurodontans, *Anolis* and *Polychrus* differ from the fossils in having greater than seven tooth positions (Smith 2011; Daza et al. 2012). Many members of Iguanidae differ in having multicuspid teeth on the premaxilla (de Queiroz 1987). Unicuspid teeth are sometimes found in *Ctenosaura*, *Cyclura* and *Sauromalus* (Scarpetta 2019), but these taxa differ in that the anterior rostral face of the premaxilla in *Ctenosaura* is distinctly rounded (Scarpetta 2019) and the nasal processes of *Ctenosaura*, *Cyclura*, and *Sauromalus* curve far posteriorly (Scarpetta 2019; Smith 2011). *Basiliscus*, *Corytophanes*, and *Enyaliodes* differ from the fossils in having a broader nasal process (Smith 2009a). *Laemanctus* differs from the fossils in having a strongly tapering nasal process (Smith 2011). TxVP 41229-29096 differs from examined *Crotaphytus* in lacking facets on the nasal process in anterior view; however, facets may be absent in some species of *Gambelia* (Daza et al. 2012). TxVP 41229-27523 has a small anterior nasal facet on the left side but differs from many *Crotaphytus* in having a more slender nasal process (McGuire 1996). *Gambelia* differ from the fossils in having longer and sharper teeth and a vertical ridge on the posterior edge of the palatal plate (McGuire 1996). Examined *Leiocephalus* differ from the fossil in having at least seven tooth positions (see also Bochaton et al. 2021), but *Leiocephalus personatus* was reported to have six tooth positions (Smith 2009a). Furthermore, examined *Leiocephalus* differ from the fossils in having a more triangular shaped nasal process that tapers posterodorsally (see also Bochaton et al. 2021). Based on these differences with other NA pleurodontans, fossils were assigned to Phrynosomatidae.

Among phrynosomatids, anterior premaxillary foramina are lacking in *Sceloporus*, *Urosaurus*, and *Petrosaurus*, are present in *Uma* and *Phrynosoma*, and are variably present in *Callisaurus*, *Cophosaurus*, *Holbrookia*, and *Uta stansburiana* (Scarpetta 2019). Some *Sceloporus* (e.g., *S. orcutii*) have an ossified bridge extending laterally from the nasal process and enclosing the medial ethmoidal foramen, but these foramina, when present, are positioned farther dorsolaterally compared to the anterior premaxillary foramina in other phrynosomatids. Sand-lizards (*Callisaurus*, *Cophosaurus*, *Holbrookia*, and *Uma*) typically have a flat rostral face of the premaxilla, while sceloporines (*Sceloporus*, *Urosaurus*, *Petrosaurus*, and *Uta*) have a more rounded rostral face (Scarpetta 2019). Fossil premaxillae are assigned to Sceloporinae on that basis.

### Quadrate

#### Description

TxVP 41229-27058 and TxVP 41229-26983 serve as the basis for our description (Fig. 22). TxVP 41229-27058 is a left quadrate. There is a thin, medially slanted central column, and the bone narrows ventrally. There is a moderately developed medial crest but no pterygoid lappet. The conch is deep and gradually slants laterally from the central column. The cephalic condyle projects posteriorly without extensive dorsal ossification and the condyle has a small dorsal tubercle. The dorsolateral margin of the tympanic crest is rounded. There is a foramen on the anteroventral surface (quadrate foramen of Villa and Delfino 2019), and a small hole on the dorsal margin of the conch that may be a taphonomic alteration. TxVP 41229-26983 differs from TxVP 41229-27058 in having a dorsal margin that slopes ventrolaterally in a step-like fashion.

#### Identification

Fossil quadrates are differentiated from NA lizards except for pleurodontans based on the absence of a pterygoid lappet (Estes et al. 1988; Evans 2008), having a wide central column (Kluge 1962; Evans 2008), and having the dorsal portion much wider compared to the articular surface (see crotaphytid section above for additional discussion on identifying quadrates to Pleurodonta). Among NA pleurodontans, in *Anolis* and *Polychrus* the lateral and medial margins are more parallel compared to the fossils. Additionally, in *Anolis* there is a distinct boss at the ventromedial margin of the quadrate not seen in the fossils. The quadrate of *Basiliscus basiliscus*, *Corytophanes*, and *Laemanctus* is proportionally taller relative to its width compared to the fossils. The quadrate of *Enyalioides heterolepis* has a more shallow conch and straighter lateral margin (tympanic crest) compared to the fossils. *Ctenosaura*, large *Iguana,* and *Leiocephalus* differ from the fossils in having a pterygoid lamina that extends farther medially. The quadrate of small *Iguana* is much more slender than the fossils. The quadrates of most *Sauromalus* have a ventrolaterally slanted dorsolateral margin, resembling TxVP 41229-26983, but differ in having a distinct dorsomedial expansion at the dorsal margin of the pterygoid lamina (Evans 2008). Some examined *Sceloporus* (*S. olivaceous* TxVP M-8375 and *S. orcutti* TxVP M-12155) also have a ventrolaterally slanted dorsolateral margin. Crotaphytids differ from the fossils in having a straight lateral margin (tympanic crest) (Mead 1988). Based on these differences with other NA pleurodontans, quadrates were assigned to Phrynosomatidae. The quadrate of phrynosomatines (except for *Uma* and *Phrynosoma*) differs from the fossils in having a very shallow conch (Evans 2008) and a strongly curved medial edge, including a medially curved central column. *Uma* and *Phrynososma* differ from the fossils in having a straight lateral margin (tympanic crest) (Mead 1988). Based on these differences fossils were assigned to Sceloporinae.

***Urosaurus* Hallowell, 1854**

**Referred specimens**: See Table S3.

### Dentary

#### Description

TxVP 41229-27603 is a left dentary with 25 tooth positions (Fig. 27). Distal teeth are weakly tricuspid and slender. The Meckelian groove is fused for about 12 tooth positions and opens at the anterior end of the dentary. The dental shelf is narrow, but widens slightly anteriorly. There is a small intramandibular lamella. The posterior end bears a distinct surangular and angular process and there is a coronoid facet within a dorsally projecting corner of bone (coronoid process). There are four nutrient foramina on the anterolateral surface. TxVP 41229-27137 is missing the anterior and posterior ends. In TxVP 41229-27137, the Meckelian groove is fused for nine tooth positions.

#### Identification

Fossil dentaries share with Pleurodonta, Xantusiidae, and some gymnophthalmoids pleurodont tricuspid teeth and a fused Meckelian groove (Evans 2008; Scarpetta 2021). Xantusiids differ in having a fused spleniodentary (Savage 1963; Mead and Bell 2001; Evans 2008). In gymnophthalmoids with tricuspid teeth and a fused Meckelian groove, the groove is completely fused to almost the level of the posterior-most tooth position (Bell et al. 2003). Among NA pleurodontans, the fossils share with some phrynosomatids, iguanids, dactyloids, extant *Leiocephalus*, polychrotids, and some corytophanids a fused Meckelian groove (Smith 2011; Scarpetta 2021). Fossils differ from these taxa, except for some phrynosomatids, in having weakly tricuspid teeth. Additionally, iguanids, *Leiocephalus*, *Basiliscus*, *Corytophanes*, and some *Laemanctus* differ from fossils in having flared tooth crowns (Smith 2009a; Scarpetta 2021) and *Polychrus* differ in having labial and lingual striations on the teeth (Smith 2009a). On this basis, fossils were identified to Phrynosomatidae. Among phrynosomatids, having a fused Meckelian groove and weakly tricuspid teeth is a combination of features only found among *Urosaurus* (Mead et al. 1985; Scarpetta 2021). A fused Meckelian groove is occasionally present in other phrynosomatids (e.g., *Petrosaurus mearnsi* TxVP M-14910); but the teeth have more prominent accessory cusps.

**Phrynosomatinae Wiens et al. 2010**

**Callisaurini “sand lizards” (*Callisaurus*, *Cophosaurus*, *Holbrookia*, and *Uma*) Wiens et al. 2010**

**Referred specimens**: See Table S3.

### Premaxilla

#### Description

TxVP 41229-29179 serves as the basis for our description (Fig. 27). TxVP 41229-29179 is a premaxilla with seven tooth positions and unicuspid teeth. It has a flat rostral surface with two anterior premaxillary foramina. The long, thin nasal process is missing the distal tip. There is a ventral keel separating the nasal facets. There are small foramina posteriorly on the palatal shelf. There are indistinct maxillary facets laterally on the alveolar plate. Posteriorly, the palatal plate is short and steeply slanted. The incisive process is minute and narrow. TxVP 41229-28865 has eight tooth positions.

#### Identification

Fossil premaxillae share with some pleurodontans, some amphisbaenians, some anguimorphs (*Ophisaurus*, gerrhonotines, and xenosaurids) anterior premaxillary foramina (Smith 2009b; Bhullar 2011; Scarpetta 2019). Examined amphisbaenians differ in having an enlarged median tooth (Smith 2009b). Anguimorphs differ from the fossils in having a more distinctly bilobed and elongate incisive process (Evans 2008). Furthermore, *Xenosaurus* differ in having a rugose rostral surface of the premaxilla (Bhullar 2011). Fossils are assigned to Pleurodonta.

Among NA pleurodontans, crotaphytids and *Leiocephalus* differ in lacking anterior premaxillary foramina (Smith 2009a). The South American hoplocercid *Enyalioides oshaughnessyi* was reported to have anterior premaxillary foramina (Smith 2009a), but they are absent in examined *E. heterolepis*. Many iguanids differ from the fossils in having multicuspid teeth on the premaxilla (de Queiroz 1987), and the remaining iguanids differ in having a rounded anterior rostral face of the premaxilla (Scarpetta 2019). *Anolis*, *Polychrus*, and corytophanids also differ from the fossils in having a more rounded anterior rostral face of the premaxilla (see also Smith 2011). Furthermore, *Basiliscus*, *Corytophanes*, and *Enyalioides* differ from the fossils in having a broader and parallel-sided nasal process (Smith 2009a, b). Based on differences with other NA pleurodontans, fossils were identified to Phrynosomatidae.

Among phrynosomatids anterior premaxillary foramina are known in *Uma*, *Phrynosoma*, *Callisaurus*, *Cophosaurus*, *Holbrookia*, and *Uta stansburiana* (Scarpetta 2019). *Uta* differs from phrynosomatines in having a rounded rostral face of the premaxilla (Scarpetta 2019). Fossils premaxillae are assigned to Phrynosomatinae based on the presence of anterior premaxillary foramina and a flat anterior rostral face of the premaxilla. Among phrynosomatines, *Phrynosoma* differs from sand-lizards in having a nasal process that is directed dorsally and a nasal process that at its base is as wide as the body of the premaxilla (Montanucci 1987; Scarpetta 2019). Premaxillae are assigned to the sand lizard clade based on having a nasal process that is directed posterodorsally and a nasal process that is narrower than the body of the premaxilla. The premaxilla of *Uma* differs from other sand-lizards in having a nasal process that resembles an isosceles triangle. Fossil premaxillae lack that morphology and so fossils likely represent *Callisaurus*, *Cophosaurus*, or *Holbrookia*.

### Quadrate

#### Description

TxVP 41229-26026 is a left quadrate (Fig. 27). The central column is indistinct, curves medially, and has a distinct medial groove. The bone narrows ventrally and there is no pterygoid lappet nor medial crest. The conch is shallow. The cephalic condyle projects posteriorly without extensive ossification dorsally. The dorsolateral margin of the tympanic crest is rounded. There is an anteriorly slanted dorsal tubercle. TxVP 41229-25586 differs in having an ossified squamosal foramen and having a dorsal surface with two depressions separated by a slanted medial ridge.

#### Identification

Fossils share the absence of a distinct pterygoid lappet with geckos, *Scincella*, *Xantusia*, some pleurodontans, and anguimorphs beside *Heloderma* (Estes et al. 1988; Evans 2008). Geckos, *Xantusia*, *Scincella*, and anguimorphs differ from the fossils in lacking a quadrate that widens dorsally. On this basis the fossil was identified to Pleurodonta. Among NA pleurodontans, the fossils share the absence of a well-developed pterygoid lamina (medial conch of Smith 2009a) with *Anolis*, *Polychrus*, *Corytophanes*, *Laemanctus*, and some phrynosomatines (Lang 1989; Smith 2009a). Corytophanids and *Polychrus* differ in having a low ridge within the lateral concha (Smith 2009a). *Anolis* differs in having a distinct boss at the ventromedial margin of the bone below the articulation with the pterygoid. The fossils share with phrynosomatines (except for *Uma* and *Phrynosoma*) a very shallow conch (Evans 2008) and a strongly curved medial edge, including a medially curved central column. This combination of features is unique to sand lizards among NA pleurodontans. Fossils are assigned to the sand lizard clade on this basis. The quadrate of *Xenosaurus grandis* also has a shallow conch strongly curved medial edge but differs from phrynosomatines in having an expanded medial pterygoid lamina and a proportionally wider mandibular condyle.

**Phrynosomatidae Fitzinger, 1843**

**Phrynosomatinae Wiens et al. 2010**

***Phrynosoma* Wiegmann, 1828**

**Referred specimens**: See Table S3.

### Premaxilla

#### Description

TxVP 41229-28986 is a premaxilla with six tooth positions and unicuspid teeth (Fig. 27). It has a flat rostral surface with two anterior premaxillary foramina on the right side. The base of the nasal process is notched on the left side without enclosed anterior foramina. There is a long, wide nasal process with small, irregularly spaced anterior foramina. The nasal process has a ventral keel separating the nasal facets. There are shallow maxillary facets laterally on the alveolar plate. Posteriorly, the palatal plate is short and steeply slanted. The incisive process is missing. TxVP 41229-25725 differs in having five tooth positions and only one right anterior foramen below a notched nasal process on both sides.

#### Identification

Premaxillae are assigned to Pleurodonta based on having a fused premaxilla (Estes et al. 1988) with less than seven tooth positions (Smith 2009a). Premaxillae are assigned to Phrynosomatinae based on the presence of less than seven tooth positions, unicuspid teeth, anterior premaxillary foramina, and a flat anterior rostral face of the premaxilla (see discussion above; Scarpetta 2019). Premaxillae can be referred to *Phrynosoma* based on a nasal process that is directly dorsally and a nasal process that at its base is as wide as the body of the premaxilla (Montanucci 1987; Scarpetta 2019).

### Maxilla

#### Description

TxVP 41229-27555 serves as the basis for our description (Fig. 27). TxVP 41229-27555 is a left maxilla with 13 tooth positions. Teeth are unicuspid and slender throughout the tooth row. The facial process is a narrow projection that diminishes anteriorly to merge with the tall crista transversalis. The palatal shelf is flat, and the palatine process is asymmetrical. The superior alveolar foramen pierces a raised area medial to the palatine process. There is a deep depression on the premaxillary process, housing two foramina, one anterior (anterior inferior alveolar foramen) and one slightly posterior (subnarial arterial foramen). There are four lateral nutrient foramina.

#### Identification

Fossils were assigned to Pleurodonta based on the presence of two foramina on the premaxillary process (Smith 2009a). The fossils share with *Phrynosoma* a facial process that is exceptionally narrow anteroposteriorly and triangular to sub-triangular in shape (Bell et al. 2004; Scarpetta 2021). This morphology of the facial process is unlike that of any other NA lizard and fossils were assigned to *Phrynosoma* on that basis. Additionally, maxillae of most *Phrynosoma* differ from other pleurodontans in lacking multicuspid teeth and differ from many pleurodontans in having an asymmetrical palatine process in dorsal and ventral view (Smith 2009a). The crista transversalis in many species of *Phrynosoma* is tall compared to other phrynosomatids (Scarpetta 2021).

### Frontal

#### Description

Morphotype A: TxVP 41229-534 serves as the basis for our description (Fig. 27). TxVP 41229-534 is a frontal with two posterolateral processes on either side, each with a small anteriorly projecting anterior superciliary process (*sensu* Montanucci 1987). The frontal is dorsoventrally tall near the base of the posterolateral processes. There are many tall, peaked tubercles on the dorsal surface beginning midway at the interorbital region and posterior. There are long superciliary horns on the posterolateral process. Anteriorly, there are bilateral facets for articulation with the nasal and prefrontal. The cristae cranii do not project far ventrally. There is a small midline notch on the posterior edge of the bone, and posterolaterally there are deep parietal facets.

Morphotype B: TxVP 41229-29015 serves as the basis for our description (Fig. 27). The fossil is mostly complete but is missing the distal end of the right posterolateral process and the left prefrontal and nasal facets. The bone is tall near the base of the posterolateral processes. The left posterolateral process has a small anteriorly projecting anterior superciliary process. There are low rounded tubercles posteriorly on the dorsal surface, and the superciliary horns on the posterolateral process are short. Anteriorly, there is a facet for articulation with the nasal and prefrontal. The cristae cranii are short and anteriorly border a deep depression on the ventral surface.

#### Identification

Fossils share with Pleurodonta and Teiioidea a fused frontal with reduced cristae cranii (Estes et al. 1988). Fossils differ from all other NA lizards, except for *Phrynosoma*, in having a supraorbital process extending anteriorly from each posterolateral portion of the frontal (Presch 1969). Fossils are assigned to *Phrynosoma* on that basis. A major difference between frontals of morphotype A and B is the morphology of the ornamentation. Fossils of both morphotypes differ from *Phrynosoma asio*, which has a smooth dorsal surface, and from *P. mcallii* which has dorsal tubercles over nearly the entire length of the frontal (Montanucci 1987).

Morphotype A: Fossils of morphotype A most closely resemble species including *P. cornutum*, *P. modestum*, *P. platyrhinos*, *P. goodei*, *P. coronatum* (reportedly variable; Montanucci 1987), *P. blainvillii*, *P. cerroense*, *P. ditmarsi*, *P. taurus, P. braconnieri,* and *P. obiculare* in having peaked tubercles on the posterolateral process (Montanucci 1987). *Phrynosoma modestum*, *P. platyrhinos*, and *P. goodei* differ from the fossils in having relatively shorter superciliary horns (see fig. 1 in Powell et al. 2017). Furthermore, *P. modestum* differs in having an anterolaterally-oriented supraorbital process. *Phrynosoma coronatum*, *P. blainvillii*, *P. cerroense*, *P. ditmarsi*, *P. modestum*, *P. platyrhinos*, and *P. obiculare* differ from the fossils in having relatively shorter tubercles and peaked tubercles largely restricted to the posterolateral process (Montanucci 1987; Powell et al. 2017). The peaked tubercles appear to be shorter in *P. braconnieri* compared to fossils, while *P. taurus* has longer superciliary horns (Powell et al. 2017). The fossils most closely resemble *P. cornutum*; however, we did not examine several species of *Phrynosoma* (e.g., *P. sherbrookei*), so we do not make a species or infrageneric identification.

Morphotype B: Fossils of morphotype B differ from morphotype A in having low rounded tubercles posteriorly on the dorsal surface, as in species in the *P. douglasii* species complex (Montanucci 1987). It was previously suggested that several fossil frontals may represent *P. modestum* (Toomey 1993). However, *P. modestum* has shorter superciliary horns and more distinctively rugose tubercles compared to the fossils and examined species in the *P. douglasii* species complex.

### Parietal

#### Description

Morphotype A: TxVP 41229-8994 serves as the basis for our description (Fig. 27). The fossil is nearly complete, only missing the distal end of the left lateral horn and the left postparietal process. The parietal table is rectangular with distinct anterolateral processes. The anterior edge has deep, bilateral facets for the frontal. The dorsal surface is covered in long tubercles. The right posterolateral parietal horn is long and there is a small horn between the main posterolateral horns. The posterior edge between the postparietal processes is characterized by two depressions (nuchal fossae) separated by a small midline posterior projection. The right postparietal process is short and projects posteroventrally and bears a lateral facet for the squamosal and a medial facet for the paroccipital process of the exoccipital. The ventrolateral crests are low without distinct epipterygoid processes. On the ventral surface, there are shallow depressions (cerebral vault) divided by a low ridge. There is no pit for the processus ascendens.

Morphotype B: TxVP 41229-25834 serves as the basis for our description (Fig. 27). TxVP 41229-25834 is a parietal missing the left posterolateral portion and the right postparietal process. The parietal table is rectangular. The anterior edge has a small midline notch between deep, bilateral facets for the frontal. The dorsal surface is covered in rounded tubercles. The right posterolateral parietal horn is short. The lateral surface has a deep depression for articulation with the squamosal. The posterior edge between the postparietal processes is characterized by two depressions (nuchal fossae) separated by a small midline posterior projection. The ventrolateral crests are low without distinct epipterygoid processes. On the ventral surface there are shallow depressions (cerebral vault) divided by a low ridge. There is no pit for the processus ascendens.

#### Identification

The broad, rectangular shape of the parietal and the presence of posterolateral dorsal horns differentiate *Phrynosoma* from those of all other North American lizards (Presch 1969). Fossils were identified to *Phrynosoma* on that basis.

Morphotype A: The length of the parietal horns varies among extant *Phrynosoma*. Fossil parietals of morphotype A have long lateral parietal horns, a small medial horn, and long dorsal bony tubercles. The length of the lateral horns of morphotype A is similar to *P. cornutum, P. solare*, *P. mcallii, P. platyrhinos, P. goodei*, *P. blainvillii*, *P. cerroense*, and *P. coronatum* (Norell 1989; Parmley and Bahn 2012; see also fig. 1 in Powell et al. 2017). It was previously suggested that a few fossil parietals (e.g., TxVP 41229-12032) may represent *P. modestum* (Toomey 1993); however, *P. modestum*, *P. obiculare*, *P. braconnieri*, and *P. asio* have slightly shorter lateral horns, and *P*. *modestum* and *P. asio* lack the long dorsal tubercles present in morphotype A (Mead et al. 1985; Montanucci 1987; Parmley and Bahn 2012). Additionally, *P. modestum* lacks a small medial horn, similar to *P. asio* and some *P. platyrhinos* (Presch 1969; Norell 1989). *Phrynosoma mcallii* differ from the fossils in having short dorsal tubercles, and *P. solare* differs in having four well developed parietal horns (Montanucci 1987). The long dorsal tubercles present in fossil parietals are similar to *P. cornutum, P. coronatum, P. solare*, *P. blainvillii*, *P. cerroense*, *P. braconnieri*, and *P. orbiculare*, but the latter two species have shorter parietal horns compared to the fossils. Based on those features, fossils are most similar to *P. cornutum, P. coronatum, P. blainvillii*, and *P. cerroense*.

Morphotype B: Fossil parietals of morphotype B have short lateral horns with no medial horn and rounded tubercles on the dorsal surface. The short lateral horns of morphotype B are similar to *P. taurus* and species in the *P. douglasii* species complex (Montanucci 1987; Norell 1989; Parmley and Bahn 2012). The tubercles in *P. taurus* are more numerous and pointed compared to the fossils (see fig. 1 in Powell et al. 2017) and the closely related *P. sherbrookei* has longer parietal horns (Nieto-Montes de Oca et al. 2014). Fossils are most similar to species in the *P. douglasii* species complex.

### Prefrontal

#### Description

Morphotype A: TxVP 41229-27205 is a left prefrontal that serves as the basis for our description (Fig. 28). It is triradiate with a small anterior orbital process, a long posteromedial process, and a long supraorbital process. The supraorbital process thins distally and curves medially. The medial surface is characterized by deep articulation facets for the frontal posteriorly and the maxilla anteriorly. Dorsal to the frontal facet, there is a small groove for the nasal articulation. The dorsal surface has several distinct tubercles. The ventral surface is smooth, concave, and has a small anterior foramen.

**Figure 28.**
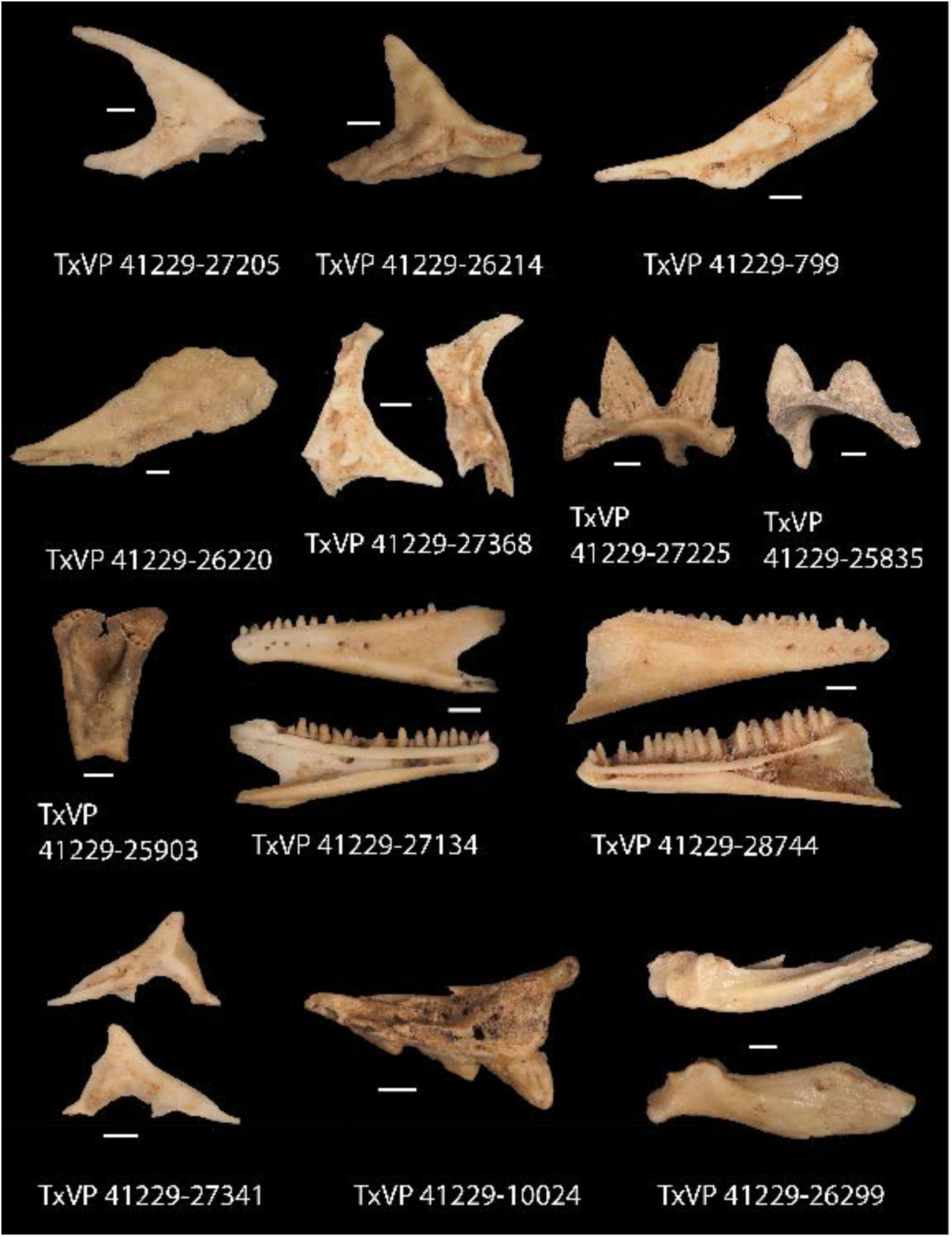
Fossil phrynosomatids. TxVP 41229-27205 Dorsal view of left prefrontal; TxVP 41229-26214 Dorsal view of left prefrontal; TxVP 41229-799 Lateral view of left jugal; TxVP 41229-26220 Lateral view of left jugal; TxVP 41229-27368 Lateral and posterior view of postorbital; TxVP 41229-27225 Anterior view of squamosal; TxVP 41229-25835 Anterior view of squamosal; TxVP 41229-25903 Posterior view quadrate; TxVP 41229-27134 Lateral and medial view of dentary; TxVP 41229-28744 Lateral and medial view of dentary; TxVP 41229-27341 Lateral and medial view of coronoid; TxVP 41229-10024 Dorsal view of compound bone; TxVP 41229-26299 Dorsal and lateral of compound bone. Scale bars = 1 mm.

Morphotype B: TxVP 41229-26214 is also a left prefrontal that serves as the basis for our description (Fig. 28). It is triradiate and the processes are nearly the same length. The anterior orbital process is notched at the anterior end for passage of the lacrimal foramen. The supraorbital process is directed posterolaterally and is relatively straight. The dorsal surface has a few low, round tubercles. The ventral surface is smooth, concave, and has a small foramen medially.

#### Identification

The prefrontal of *Phrynosoma* is distinct from nearly all NA lizards (except for *Corytophanes*) in having a long supraorbital process that extends posteriorly to partially or fully enclose the supraorbital fenestra (Presch 1969; Lang 1989). The prefrontal of *Corytophanes* differs from that of *Phrynosoma* in having a deep depression on the lateral surface of the bone. Fossils were identified to *Phrynosoma* on that basis.

Morphotype A: Fossils of morphotype A have a long supraorbital process that curves medially. A long supraorbital process also occurs in *P. cornutum*, *P. mcallii*, *P. solare*, *P. taurus*, *P. asio*, and somewhat in *P. braconnieri* (Montanucci 1987; Powell et al. 2017). The supraorbital process of *P. solare* appears to be relatively longer compared to the fossils and other species of *Phrynosoma* (Montanucci 1987; Powell et al. 2017).

Morphotype B: Fossils of morphotype B have a shorter supraorbital process that is straighter. A short supraorbital process also occurs in *P. coronatum*, *P. cerroense*, *P. blainvilli*, *P. ditmarsi*, *P. modestum*, *P. obiculare*, *P. platyrhinos*, and species in the *P. douglasii* species complex (Montanucci 1987; Powell et al. 2017). *P. modestum* differs from the fossil in having a more distinctly rugose surface and having a large anterior notch.

### Jugal

#### Description

Morphotype A: TxVP 41229-799 is a left jugal (Fig. 28). The anterior orbital process is thin with a ventral maxillary articulation facet. The postorbital process is broad, is directed posterodorsally, and has a distinct anterior postorbital articulation facet. The lateral surface of the postorbital process has a row of three long tubercles. The medial surface is smooth, and there is a depression on the postorbital process. There is a foramen on the lateral surface near the inflection point.

Morphotype B: TxVP 41229-26220 serves as the basis for our description and is a well preserved left jugal (Fig. 28). TxVP 41229-26220 differs from morphotype A in having the anterior orbital process widened relative to the postorbital process, a broad, flat posterodorsal surface in lateral view, and the lateral surface of the postorbital process has a row of several low, rounded tubercles.

#### Identification

Fossil jugals share with *Phrynosoma*, *Anolis*, *Corytophanes*, *Lepidophyma*, and *Xantusia riversiana* an expanded postorbital process (Savage 1963; Montanucci 1987; Lang 1989; Smith 2011). *Anolis* differs in having a relatively less expanded postorbital process (Smith 2011). Fossil jugals differ from *Anolis*, *Corytophanes*, *Xantusia riversiana*, and *Lepidophyma* in having a more posteriorly oriented postorbital process (Savage 1963; Smith 2011; Lang 1989). Fossils were identified to *Phrynosoma*.

Morphotype A: Fossils of morphotype A have a row of long tubercles on the lateral surface. This is similar to the condition described for *P. ditmarsi*, *P. coronatum*, *P. solare*, *P. cornutum*, *P. modestum*, *P. mcallii*, and *P. platyrhinos* (Montanucci 1987). *Phrynosoma braconnieri*, *P. taurus*, *P. obiculare*, *P. douglassi*, and *P. hernandesi* have lower and more rounded tubercles compared to the fossils (Montanucci 1987). We were unable to examine this feature in all species of *Phrynosoma* and do not make species level identifications.

Morphotype B: Fossils of morphotype B share with *P. orbiculare* and species in the *P. douglasii* species complex a broad, flat posterodorsal surface in lateral view (Montanucci 1987). Fossils of morphotype B also have a row of several low, rounded tubercles. This is similar to the condition described for *P. taurus*, *P. braconnieri*, and species in the *P. douglasii* species complex. *Phrynosoma taurus* and *P. braconnieri* differ in having more triangular shaped jugals (Montanucci 1987). *Phrynosoma orbiculare* also has low, rounded tubercles anteriorly, but may have a more distinct posterior tubercle (Montanucci 1987). The fossils are similar to species in the *P. douglasii* species complex, but we were unable to examine features on the jugal for all species of *Phrynosoma* making this identification tentative.

### Postorbital

#### Description

TxVP 41229-27368 serves as the basis for our description and is a right postorbital (Fig. 28). The dorsal process widens mediolaterally, and projects anteriorly with a lateral facet for articulation with the frontal and parietal. Laterally, the postorbital is twisted with a long, ventrally pointed process. Posterolaterally, there is a squamosal articulation facet, and the lateral margin has a groove for articulation with the jugal. The dorsal surface has several long tubercles. Ventrally there is a ridge that runs from the posteromedial corner onto the ventrolateral process.

#### Identification

Postorbitals were identified to Pleurodonta based on a sub-triangular morphology with a distinct ventral process (Estes et al. 1988). Among NA pleurodontans, a broadened dorsal process is found in *Phrynosoma*, *Iguana iguana*, and *Corytophanes* (Presch 1969; Lang 1989; Evans 2008). The dorsal process in examined *Iguana iguana* is not expanded to the extent seen in the fossils. The postorbital of *Corytophanes* differs in having an anteriorly projecting spine extending from the dorsal process that contacts the prefrontal (Lang 1989). On this basis fossils were identified to *Phrynosoma*. The fossils share with *P. solare*, *P. braconnieri*, *P. cornutum*, *P. coronatum*, *P. ditmarsi*, *P. mcallii*, *P. modestum*, *P. obiculare*, *P. platyrhinos*, and *P. taurus* a rugose lateral surface with distinct tubercles (Montanucci 1987). *Phrynosoma mcallii* differs in having a posteriorly expanded postorbital that restricts the supratemporal fossa (Norris and Lowe 1951).

### Squamosal

#### Description

Morphotype A: TxVP 41229-27225 is a right squamosal that serves as the basis for our description (Fig. 28). The anterior edge is concave with a posteromedial process that is broken at the distal end. The dorsal surface has two long medial horns and one short lateral horn. There is a short posteroventrally projecting inferior process (*sensu* Powell et al. 2017) with a lateral facet for the supratemporal.

Morphotype B: TxVP 41229-25835 is a left squamosal missing the posteromedial process and a portion of the anterior edge (Fig. 28). The dorsal surface has two short medial horns and one minute lateral horn. Posteriorly, there is a short posteroventraly projecting inferior process with a lateral facet for the supratemporal.

#### Identification

The squamosal of *Phrynosoma* is unique among North American lizards in having horns along the dorsal surface (Presch 1969). Fossils were identified to *Phrynosoma*.

Morphotype A: The number and length of the squamosal horns varies among extant *Phrynosoma*. The number of horns was previously considered to be of limited diagnostic use because of considerable overlap between extant species (Presch 1969; Bell 1993) and because fossils are often broken, so exact horn number is often indeterminate. Fossil squamosals of morphotype A have long horns similar to *P. cornutum*, *P. modestum*, *P. asio*, *P. coronatum*, *P. mcallii*, *P. platyrhinos*, and *P. solare* (Parmley and Bahn 2012). *Phrynosoma asio* differs in only having two horns and *P. solare* has four (Montanucci 1987; Bell 1993). *Phrynosoma modestum* differs in having a long posterior horn and shorter anterior horns (Parmley and Bahn 2012).

Morphotype B: Fossil squamosals of morphotype B have short horns. The short lateral horns are similar to *P. braconnieri*, *P. ditmarsi*, and species in the *P. douglasii* species complex (Parmley and Bahn 2012; Powell et al. 2017). *Phrynosoma braconnieri* and *P. ditmarsi* differ in having a relatively broader squamosal in dorsal view (Powell et al. 2017). Fossils are most similar to species in the *P. douglasii* species complex.

### Quadrate

#### Description

TxVP 41229-25903 serves as the basis for our description and is a left quadrate (Fig. 28). There is a wide, medially slanted central column, and the bone narrows ventrally. There is no pterygoid lappet, and the medial crest is minute. The conch is deep and is obscured from view at its dorsomedial margin by the central column. The cephalic condyle projects posteriorly without extensive dorsal ossification. The dorsolateral margin of the tympanic crest is slightly rounded. The anterior surface is convex. There is a foramen on the anteroventral surface (quadrate foramen of Villa and Delfino 2019) and a foramen medial to the central column.

#### Identification

Fossil quadrates were identified to Pleurodonta based on the absence of a pterygoid lappet (Estes et al. 1988) and widening dorsally relative to the articular surface. The quadrate of *Phrynosoma* (except *P. modestum* and *P. mcallii*) differs from other pleurodontans in having a conch that is deep and obscured from view at its dorsomedial margin by the central column. In examined *P. modestum* and *P. mcallii* the conch is shallow. The fossil shares with *P. cornutum*, *P. coronatum*, *P. mcallii*, *P. modestum*, *P. platyrhinos*, and *P. solare* a minute medial crest (Presch 1969).

### Dentary

#### Description

Morphotype A: TxVP 41229-27134 serves as the basis for our description and is a left dentary with 19 tooth positions (Fig. 28). Teeth are unicuspid and slender. The Meckelian groove is open medially for its entire length, and the dentary is tall posteriorly. The suprameckelian lip is short anteriorly. The dental shelf is narrow and there is an intramandibular lamella. The posterior end of the dentary is bifurcated. The angular process is broken at the distal end but is flat ventrally and curves far medially. There is a small lateral tubercle at the base of the angular process. There are seven nutrient foramina on the anterolateral surface.

Morphotype B: TxVP 41229-28744 serves as the basis for our description and is a right dentary with 20 tooth positions (Fig. 28). Teeth are unicuspid and slender. The suprameckelian and inframeckelian lips meet midway along the tooth row, and the dentary is tall posteriorly. The suprameckelian lip is short anteriorly. The dental shelf is narrow. The angular process is flat ventrally and curves far medially. The lateral surface is smooth and lacks tubercles, and there are five nutrient foramina anteriorly.

#### Identification

Morphotype A: Dentaries of Morphotype A share with Pleurodonta and some teiids pleurodont teeth and an inframeckelian lip that curls dorsolingually, producing a medial exposure of the Meckelian groove along the mid-length of the dentary (Gauthier et al. 2012; Bochaton et al. 2017). Fossil dentaries differ from teiids in lacking a broad subdental shelf (Estes et al. 1988), lacking asymmetric bicuspid teeth, and lacking large amounts of cementum deposits at base of teeth (Estes et al. 1988; Nydam et al. 2007; Scarpetta 2020). Fossils are assigned to Pleurodonta. *Phrynosoma* differ from other NA pleurodontans in having unicuspid teeth (except for some slightly tricuspid posterior teeth in *P. asio*, *P. coronatum*, and *P. mcallii*; Presch 1969), a relatively flat posteroventral surface, and a posteroventral lamina of bone that is curved far medially (Mead et al. 1999). Fossils are assigned to *Phrynosoma* on this basis. Fossils of morphotype A share with *P. asio*, *P. cornutum*, *P. mcallii*, *P. modestum*, and *P. platyrhinos* an open Meckelian groove (Presch 1969). Of those species, *P*. *asio* differs in having a smooth lateral surface (Presch 1969; Mead et al. 1999). *Phrynosoma modestum* was reported to have a smooth lateral surface by Presch (1969); however, other authors note that *P. modestum* has a rugose lateral surface (Montanucci 1987; Mead et al. 1999; Parmley and Bahn 2012), which is supported by our observations.

Morphotype B: Dentaries of Morphotype B share with Pleurodonta, some teiids, and some scincids pleurodont teeth and suprameckelian and inframeckelian lips that meet to close the Meckelian groove (Greer 1974; Gauthier et al. 2012; Bochaton et al. 2017; Scarpetta 2021). Fossil dentaries differ from teiids and scincids in lacking a broad subdental shelf (Estes et al. 1988) Fossils further differ from teiids in lacking asymmetric bicuspid teeth and lacking large amounts of cementum deposits at base of teeth (Estes et al. 1988; Nydam et al. 2007; Scarpetta 2020). Fossils are assigned to Pleurodonta. Fossils of morphotype B share a closed Meckelian groove with *P. braconnieri*, *P. coronatum*, *P. orbiculare*, *P. solare*, and species in the *P. douglasii* species complex (Presch 1969). We also observed one specimen of *P. modestum* (TxVP M-14818) with a closed Meckelian groove. *Phrynosoma solare* and examined *P. modestum* differ in having a rugose lateral surface of the dentary (Presch 1969; Montanucci 1987).

### Coronoid

#### Description

TxVP 41229-27341 is a left coronoid (Fig. 28). The coronoid process is tall and wide, and gradually declines anteriorly, giving the coronoid a triangular appearance. The anteromedial process is elongated, with a medial articulation facet for the dentary, and a small ventrally-projecting lamina of bone. The posteromedial process is thin and directed posteroventrally. There is a distinct medial crest that extends from the coronoid process onto the posteromedial process. There is a small, rounded lateral process. There is a vertically oriented lateral crest that terminates at the posterior margin of the lateral process. The dorsal articulation facet for the surangular is narrow. TxVP 41229-26885 and TxVP 41229-25848 differ in having lateral crests that end anteriorly on the lateral process.

#### Identification

Fossil coronoids share with several pleurodontans and xantusiids the absence of a distinct anterolateral process (Estes et al. 1988). Xantusiids differ in having an anterior groove extending onto the coronoid process (Savage 1963). Fossils differ from examined NA pleurodontans, except for some *Phrynosoma* (see fig. 4 in Meyers et al. 2006), in having a triangular-shaped coronoid process that is sloped at a low angle anteriorly. Furthermore, fossils differ from examined NA pleurodontans, except for some *Phrynosoma* (e.g., *P. cornutum* TxVP M-6405 and *P. douglasii* TxVP M-8526), in having a thin, posteriorly directed posteromedial process. Fossils were assigned to *Phrynosoma*. Fossils differ from *P. braconnieri*, *P. mcallii*, and *P. solare* in having a lateral process to overlap the dentary and surangular (Presch 1969). Examined *P. coronatum*, *P. ditmarsi*, *P. orbiculare*, and *P. hernandesi* differ in having a more steeply sloped anterior margin of the coronoid process (see fig. 4 of Meyers et al. 2006).

### Compound bone

#### Description

Morphotype A: TxVP 41229-10024 is a left compound bone missing the prearticular and the anterior end of the surangular that serves as the basis for our description (Fig. 28). There is a broad articular surface. The retroarticular process (post-condylar process of Presch 1969) is mediolaterally flat, tall, and oriented in a near vertical plane. The posteroventral end of the retroarticular process is rounded and extends posterior to the dorsal portion. On the surangular there are three lateral horns. The posterior-most horn is long and narrow, and the other horns are short and broad.

Morphotype B: TxVP 41229-26299 serves as the basis for our description and is a right compound bone that is missing the prearticular (Fig. 28). There is a broad articular surface and a short medial process. The retroarticular process is mediolaterally flat and slanted ventrolaterally with a rounded lateral end. The lateral surface of the surangular is smooth, the dorsal margin has a dorsally expanded crest where it articulates with the coronoid, and the ventral margin is slightly convex. There is a posterior surangular foramen and an anterior surangular foramen on the lateral surface.

#### Identification

The compound bone of *Phrynosoma* is unlike that of other North American lizards in having a retroarticular process that is twisted in a near vertical plane with dorsal and ventral tubercles on the posterior end (Presch 1969). Fossils were assigned to *Phrynosoma* on that basis.

Morphotype A: Fossils of morphotype a share with *P. cornutum*, *P. ditmarsi*, *P. modestum*, *P. mcallii*, *P. platyrhinos*, *P. goodei*, and *P. solare* horns on the lateral surface (Presch 1969; Montanicci 1987). *Phrynosoma ditmarsi* differs in having the horns oriented more ventrally (Montanicci 1987; see also fig. 4 in Meyers et al. 2006). *Phrynosoma modestum* differs in having shorter horns relative to the fossils, and *P. platyrhinos* and *P. solare* differ in only having two lateral horns (Presch 1969). Fossils have a comparable horn morphology to *P. cornutum* and *P. mcallii*.

Morphotype B: Fossils of morphotype B have a flat lateral surface without horns, similar to that in *P. asio*, *P. cerroense, P. coronatum*, *P. orbiculare*, *P. taurus*, *P. braconnieri*, and species in the *P. douglasii* species complex (Presch 1969).

**Anguimorpha Fürbringer, 1900**

**Anguidae Gray, 1825**

**Referred specimens**: See Table S3.

### Coronoid

#### Description

TxVP 41229-27599 is a left coronoid (Fig. 29). The coronoid process is tall, broad, and rounded. The anteromedial process is elongate, but the distal end is missing. The anteromedial process has a medial splenial facet that, with the anteriorly projecting lateral process, forms a narrow facet for the coronoid process of the dentary. The posteromedial process is missing the distal end, but the remaining portion is posteroventrally directed with an expanded posterior lamina of bone to articulate medially with the surangular. The medial crest extends from the coronoid process onto the posteromedial process. There is a vertically oriented lateral crest that ends at the posterior margin of the lateral process. The anteromedial process has a notch, which may convey a foramen.

**Figure 29.**
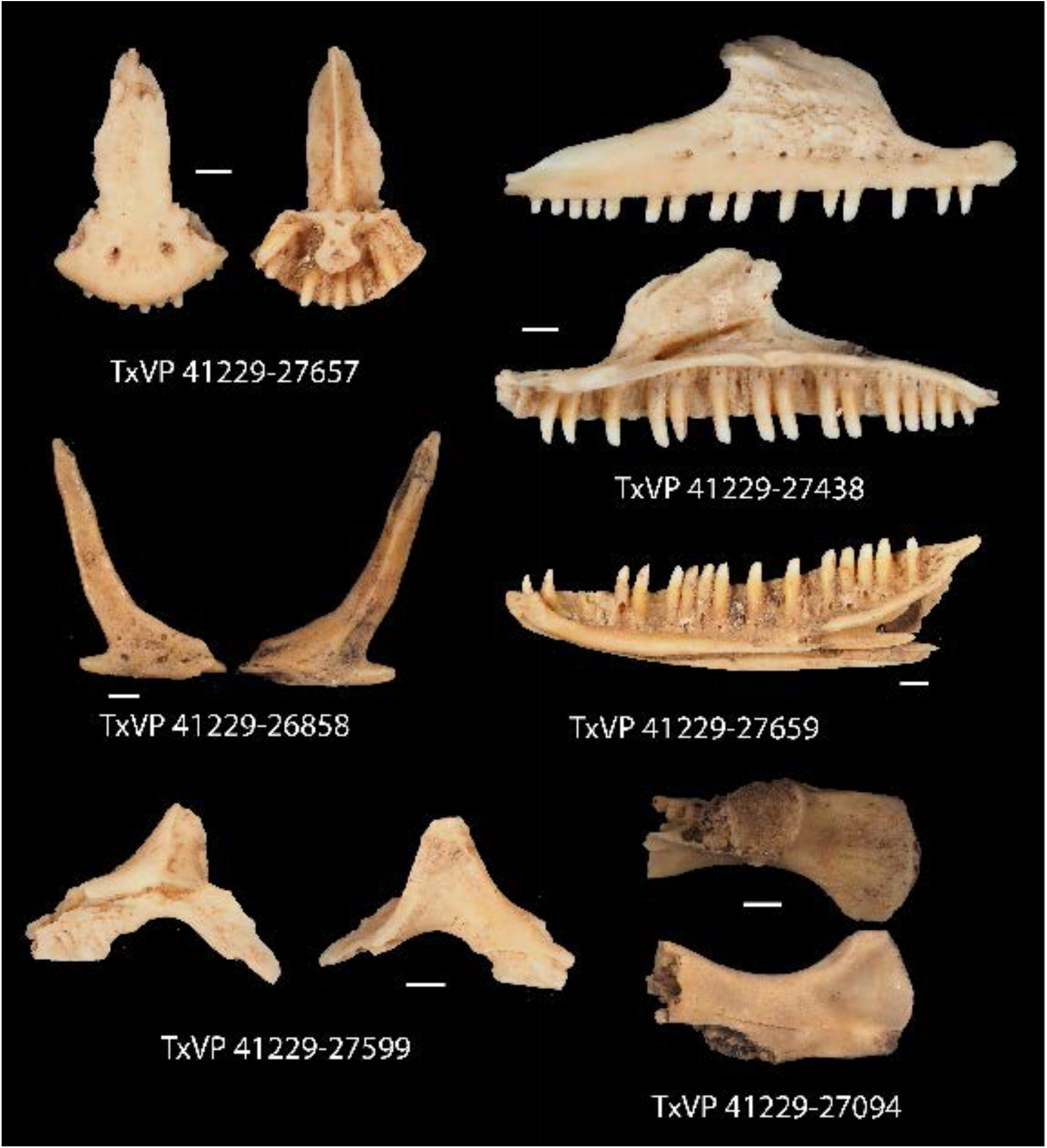
Fossil anguids. TxVP 41229-27657 Dorsal and ventral view of premaxilla; TxVP 41229-27438 Lateral and medial view of right maxilla; TxVP 41229-26858 Lateral and medial view of right jugal; TxVP 41229-27659 Medial view of right dentary; TxVP 41229-27599 Lateral and medial view of left coronoid; TxVP 41229-27094 Dorsal and ventral view of right compound bone. Scale bars = 1 mm.

#### Identification

The fossil coronoid differs from xantusiids and some pleurodontans in having an anterolateral process (Estes et al. 1988; Fig. 30). The fossil differs from examined NA pleurodontans with an anterolateral process (see above) in having a more posteriorly oriented posteromedial process.

**Figure 30.**
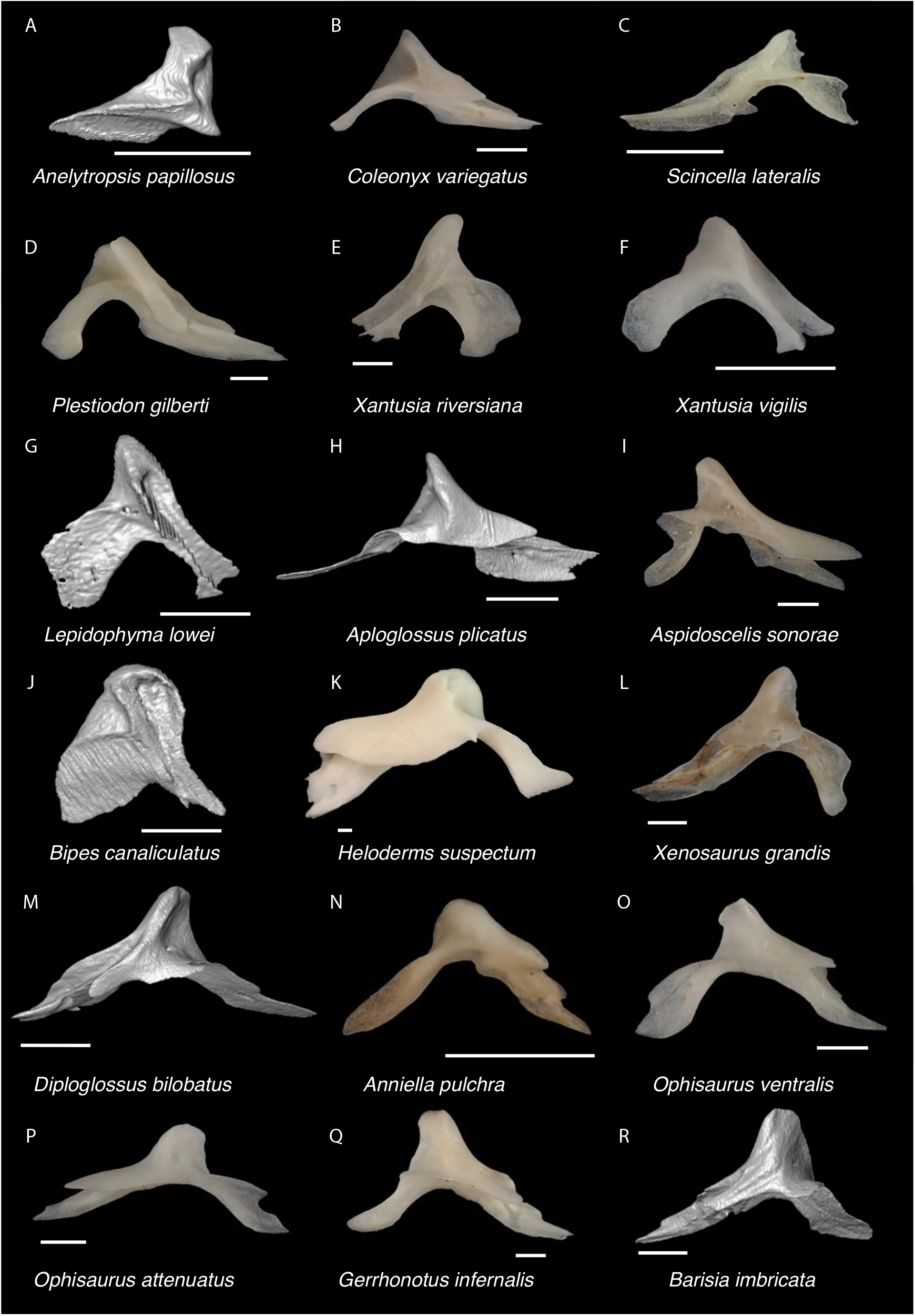
Non-Pleurodontan Coronoid bones. Coronoids in lateral view– **A**. *Anelytropsis papillosus* UF 86708; **B**. *Coleonyx varegatus* TxVP M-13892; **C**. *Scincella lateralis* TxVP M-4489; **D**. *Plestiodon gilberti* TxVP M-8587; **E**. *Xantusia riversiana* TxVP M-8505; **F**. *Xantusia virigatus* TxVP M-12130; **G**. *Lepidophyma lowei* LACM 143367; **H**. *Aploglossus plicatus* TNHC 34481; **I**. *Aspidoscelis sonorae* TxVP M-15670; **J**. *Bipes canaliculatus* CAS 134753; **K**. *Heloderma suspectum* TxVP M-9001; **L**. *Xenosaurus grandis* TxVP M-8960; **M**. *Diploglossus bilobatus* TNHC 31933; **N**. *Anniella pulchra* TxVP M-8678; **O**. *Ophisaurus ventralis* TxVP M-8585; **P**. *Ophisaurus attenuatus* TxVP M-8979; **Q**. *Gerrhonotus infernalis* TxVP M-13441; **R**. *Barisia imbricata* TNHC 76984. Scale bars = 1 mm.

TxVP 41229-27599 differs from examined *Aspidoscelis* and *Ameiva* in lacking a deeply notched posterior edge, forming dorsal and ventral rami (Tedesco et al. 1999). Furthermore, *Aspidoscelis*, *Ameiva*, and *Pholidoscelis* differ in having a distinct lateral crest running from the apex of the coronoid process anteroventrally onto the anterolateral process (Tedesco et al. 1999; Bochaton et al. 2019). Fossils differ from gymnophthalmoids in having a relatively shorter anterolateral process (Bell et al. 2003; Roscito and Rodrigues 2010; Morales et al. 2019). North American geckos such as *Coleonyx variegatus*, *C. brevis*, *Sphaerodactylus roosevelti*, and *Thecadactylus rapicauda* differ from the fossil in having a thinner posteromedial process (Kluge 1962; Daza et al. 2008; Bochaton et al. 2018). The posteromedial process is slightly wider in *C. elegans* and *C. mitratus* compared to other species (Kluge 1962) but examined *Coleonyx*, including *C. elegans*, have a coronoid process sloped at a posteriorly lower angle compared to the fossil. The coronoid of *Coleonyx variegatus* and *C. brevis* also differ from the fossil in having a dorsoventrally expanded anterolateral process (Kluge 1962). The lateral crest on the coronoid process is almost vertically oriented in the fossil, but in examined *Plestiodon* and *Scincella* the crest is more obliquely oriented. Furthermore, the anterolateral process is more anteriorly oriented in the fossil but more ventrally oriented in *Plestiodon* (see also fig. 4 in Nash 1970). Based on differences from other NA lizards, the fossil coronoid is referable to Anguimorpha. *Xenosaurus*, except for *X. rackhami*, differs from the fossil in having a foramen on the anterolateral process (Evans 2008; Bhullar 2011). *Heloderma* differs in having a dorsoventrally expanded anterolateral process and *Anniella* differs in having a shorter coronoid process (Evans 2008). On this basis the fossil was identified to Anguidae.*mbricata*

### Compound bone

#### Description

TxVP 41229-27094 is a right compound bone missing much of the portion anterior to the articular surface (Fig. 29). The retroarticular process is broadened and medially oriented. The dorsal surface of the retroarticular process is slightly concave, and the ventral surface bears a distinct sub-triangular depression. There is a ventral angular articulation facet. The articular surface is broad and saddle shaped. There are two small foramina posterior to the articular surface.

#### Identification

The fossil shares with anguimorphs, geckos, and scincids a medially directed and broadened retroarticular process (Estes et al. 1988). Geckos differ from the fossil in having a distinct notch on the medial margin of the retroarticular process (Estes et al. 1988). Scincids differ in having a tubercle or flange on the medial margin of the retroarticular process (Estes et al. 1988); however, this feature was not obvious in all examined specimens, particularly for *Scincella* (Fig. 31). The fossil differs from examined scincids in having a distinct sub-triangular depression on the ventral surface of the retroarticular process. Based on differences from other NA lizards, the fossil is referable to Anguimorpha. Examined *Anniella* differ in having a more medially slanted retroarticular process and examined *Xenosaurus* and *Heloderma* have a more slender retroarticular process (Evans 2008). The fossil is referable to Anguidae. A distinct sub-triangular depression on the ventral surface of the retroarticular process was observed in gerrhonotines and diploglossines but was absent in examined *Ophisaurus*. We refrain from making a more refined identification pending examination of additional skeletal material of diploglossines.

**Figure 31.**
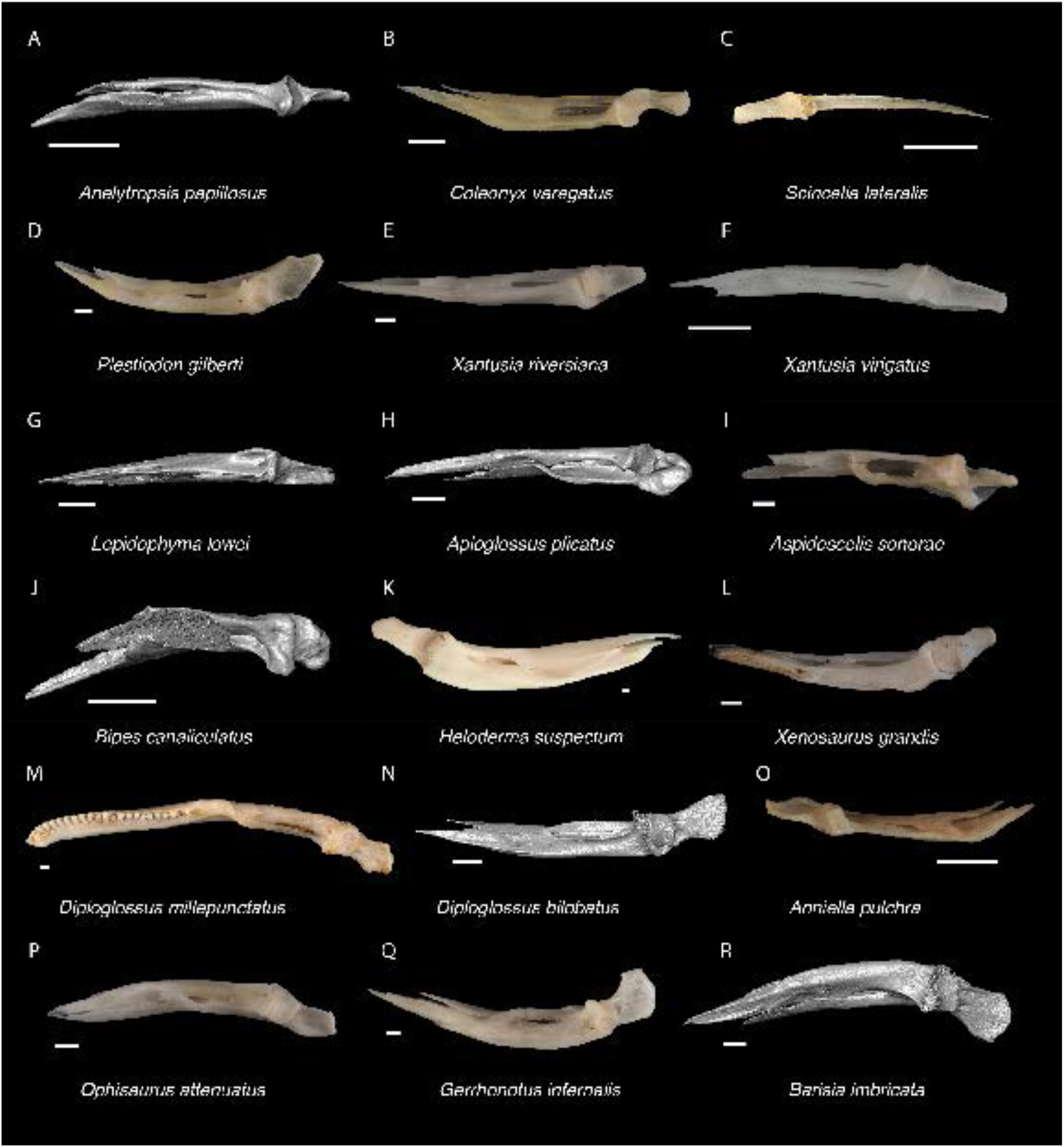
Non-Pleurodontan Compound bones. Compound bones in dorsal view– **A**. *Anelytropsis papillosus* UF 86708; **B**. *Coleonyx varegatus* TxVP M-13892; **C**. *Scincella lateralis* TxVP M-5531; **D**. *Plestiodon gilberti* TxVP M-8587; **E**. *Xantusia riversiana* TxVP M-8505; **F**. *Xantusia virigatus* TxVP M-12130; **G**. *Lepidophyma lowei* LACM 143367; **H**. *Aploglossus plicatus* TNHC 34481; **I**. *Aspidoscelis sonorae* TxVP M-15670; **J**. *Bipes canaliculatus* CAS 134753; **K**. *Heloderma suspectum* TxVP M-9001; **L**. *Xenosaurus grandis* TxVP M-8960; **M**. *Diploglossus millepunctatus* TxVP M-9010; **N**. *Diploglossus bilobatus* TNHC 31933; **O**. *Anniella pulchra* TxVP M-8678; **P**. *Ophisaurus attenuatus* TxVP M-8979; **Q**. *Gerrhonotus infernalis* TxVP M-13441; **R**. *Barisia imbricata* TNHC 76984. Scale bars = 1 mm.

**Anguinae Gray, 1825**

***Ophisaurus* Daudin, 1803**

**Referred specimens**: See Table S3.

### Premaxilla

#### Description

TxVP 41229-28734 is a premaxilla with nine tooth positions (Fig. 32). Teeth are unicuspid. The rostral surface is rounded, and the nasal process is strongly curved posteriorly. The nasal process is thin and slightly waisted at the base and has a short posterior keel. There are lateral maxillary facets but no dorsal ossifications on the alveolar plate. The palatal plate has short posterior projections. The incisive process is large, round, and bilobed. There are small foramina posterolateral to the base of the nasal process and there is a single midline anterior foramen.

**Figure 32.**
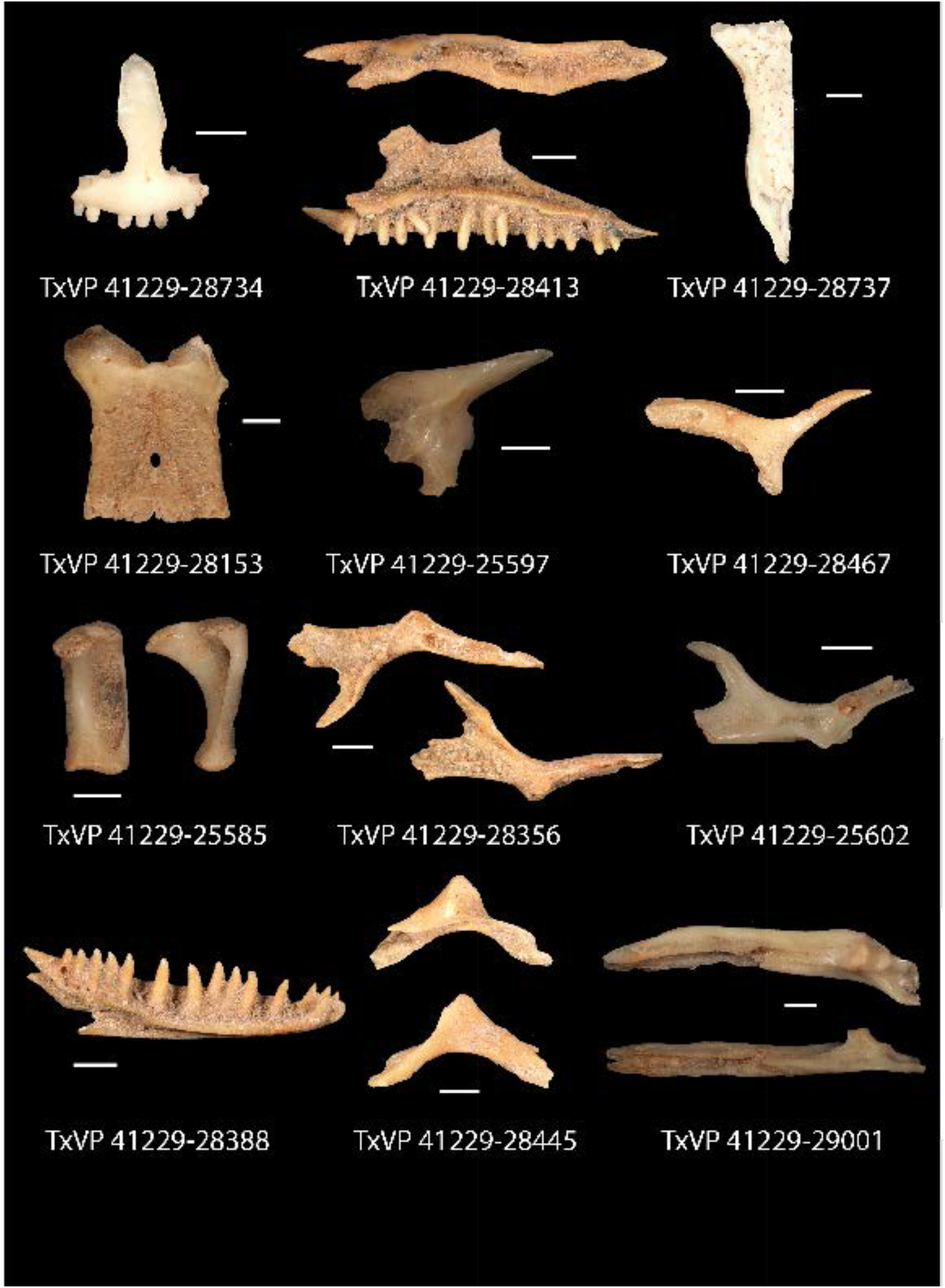
Fossil anguines. TxVP 41229-28734 Anterior view of premaxilla; TxVP 41229-28413 Dorsal and medial view of right maxilla; TxVP 41229-28737 Dorsal view of frontal; TxVP 41229-28153 Dorsal view of parietal; TxVP 41229-25597 Lateral view of prefrontal; TxVP 41229-28467 Dorsal view of postfrontal; TxVP 41229-25585 Posterior and lateral view of quadrate; TxVP 41229-28356 Dorsal and ventral view of pterygoid; TxVP 41229-25602 Dorsal view of pterygoid; TxVP 41229-28388 Medial view of dentary; TxVP 41229-28445 Lateral and medial view of coronoid; TxVP 41229-29001 Dorsal and medial view of compound bone. Scale bars = 1 mm.

#### Identification

The fossil premaxilla is assigned to Anguimorpha based on having a large, round, and bilobed incisive process (Evans 2008; Fig. 33). The incisive process is relatively smaller and less distinctly bilobed in pleurodontans, scincids (when present), and xantusiids (Evans 2008). The fossil shares with anguids and *Xenosaurus* nine tooth positions (Conrad et al. 2011; Bhullar 2011; Scarpetta 2018). The fossil differs from *Xenosaurus* in lacking a rugose rostral surface of the premaxilla (Bhullar 2011). The fossil shares with Anguinae and Diploglossinae a forked palatal process (Evans 2008; Conrad et al. 2011; Scarpetta 2018). The fossil differs from Diploglossinae and is assigned to Anguinae based on the absence of a dorsal ossification on the palatal plate posterior to the medial ethmoidal foramen (Scarpetta 2018).

**Figure 33.**
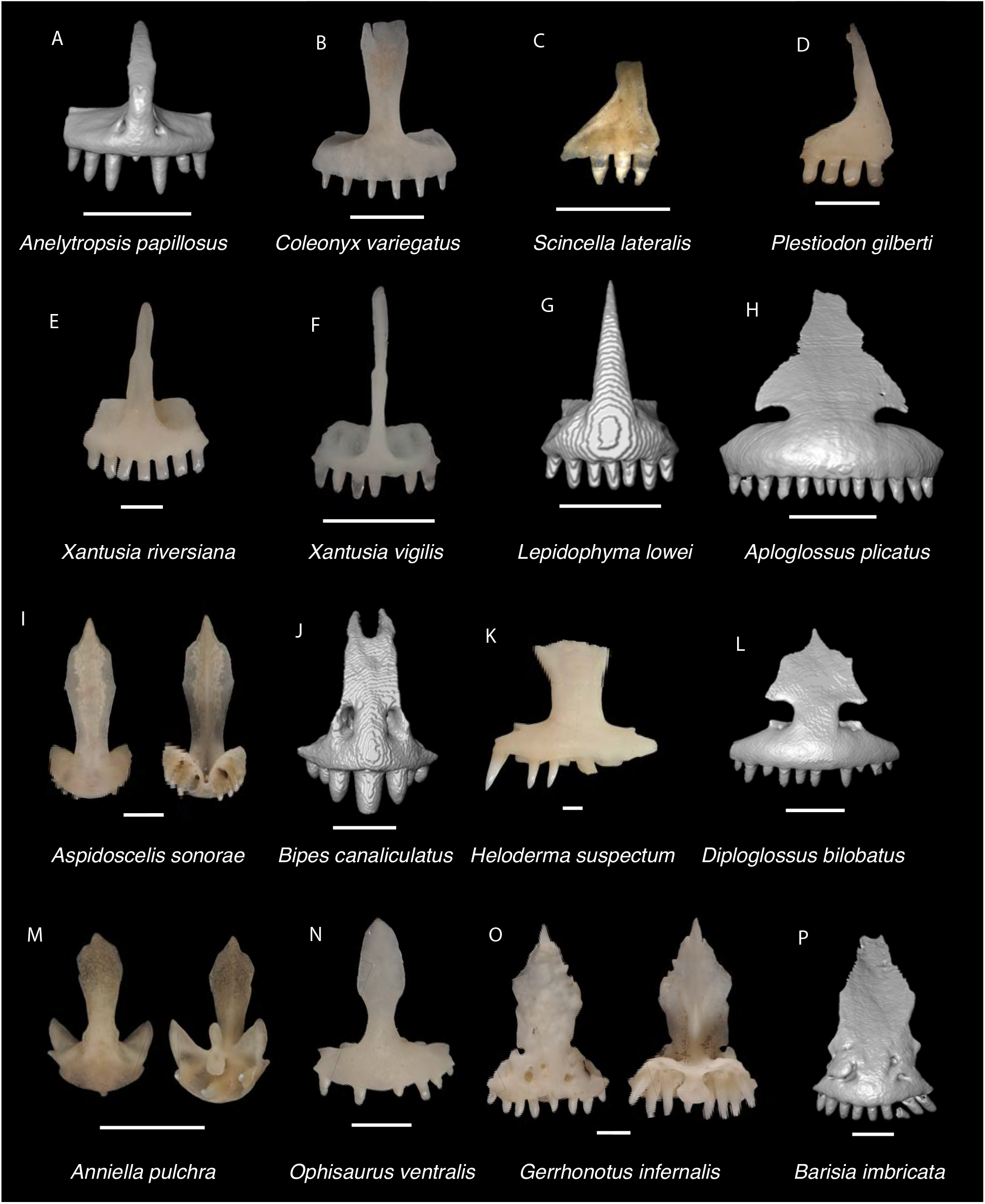
Non-Pleurodontan Premaxillae. Premaxillae in anterior, dorsal, and ventral views– **A**. *Anelytropsis papillosus* UF 86708; **B**. *Coleonyx variegatus* TxVP M-12109; **C**. *Scincella lateralis* TxVP M-4489; **D**. *Plestiodon gilberti* TxVP M-8587; **E**. *Xantusia riversiana* TxVP M-8505; **F**. *Xantusia virigatus* TxVP M-12130; **G**. *Lepidophyma lowei* LACM 143367; **H**. *Aploglossus plicatus* TNHC 34481; **I**. *Aspidoscelis sonorae* TxVP M-15670; **J**. *Bipes canaliculatus* CAS 134753; **K**. *Heloderma suspectum* TxVP M-9001; **L**. *Diploglossus bilobatus* TNHC 31933; **M**. *Anniella pulchra* TxVP M-8678; **N**. *Ophisaurus ventralis* TxVP M-8585; **O**. *Gerrhonotus infernalis* TxVP M-13441; **P**. *Barisia imbricata* TNHC 76984. Scale bars = 1 mm.

### Maxilla

#### Description

TxVP 41229-28413 serves as the basis for our description and is a right maxilla (Fig. 32). There are 17 tooth positions and teeth are unicuspid with medial striations. The dorsal portion of the facial process is broken, but it is broad. The anterior face of the facial process gently curves medially and has an anterodorsal projection. The premaxillary process is strongly bifurcated with a longer, pointed lateral projection and a shorter medial lappet. The crista transversalis trends anteromedially from the facial process. There is a narrow palatal shelf with a rounded palatine process. The palatal shelf becomes especially narrow anterior to the palatine process. There is a deep recess on the medial surface of the facial process. The lateral wall of the posterior orbital process is short with a small notch posteriorly. The dorsal surface of the postorbital process has a shallow jugal groove. There is a large superior alveolar foramen on the palatal shelf medial to the palatine process, and five lateral nutrient foramina.

#### Identification

Fossil maxillae share with some scincids and anguimorphs unicuspid teeth with striated crowns (Estes 1963; Smith 2009b; Fig. 34). Fossils differ from examined scincids in lacking a large notch at the end of the posterior orbital process (see also fig. 5 in Čerňanský and Syromyatnikova 2021). *Scincella* differs in lacking striations on the crowns (Townsend et al. 1999). Examined *Plestiodon* differ in having the crista transversalis abruptly trend medially anterior to the facial process and having a much wider palatal shelf. Based on these differences, the fossils are referable to Anguimorpha.

*Anniella* and *Heloderma* differ from the fossils in having more pointed and recurved teeth as well as fewer tooth positions (up to seven in *Anniella* and up to ten in *Heloderma*; Torien 1950; Evans 2008). *Xenosaurus* differs in having fused osteoderms on the lateral surface of the facial process (Bhullar 2011). Fossils are assigned to Anguidae. Fossils share with anguines and diploglossines a deeply notched premaxillary process (Meszoely 1970). Fossil maxillae share with Anguinae sharp and widely spaced teeth (Meszoely 1970). Some species of *Abronia* (e.g., *Abronia mixteca*; Scarpetta et al. 2021) and perhaps some diploglossines (e.g., *Diploglossus fasciatus*; see fig. 2 in Syromyatnikova and Aranda 2022) also have sharp and widely spaced teeth, but in *Abronia* and diploglossines, the palatal shelf anterior to the palatine process does not narrow to the same degree as in fossils and examined anguines.

**Figure 34.**
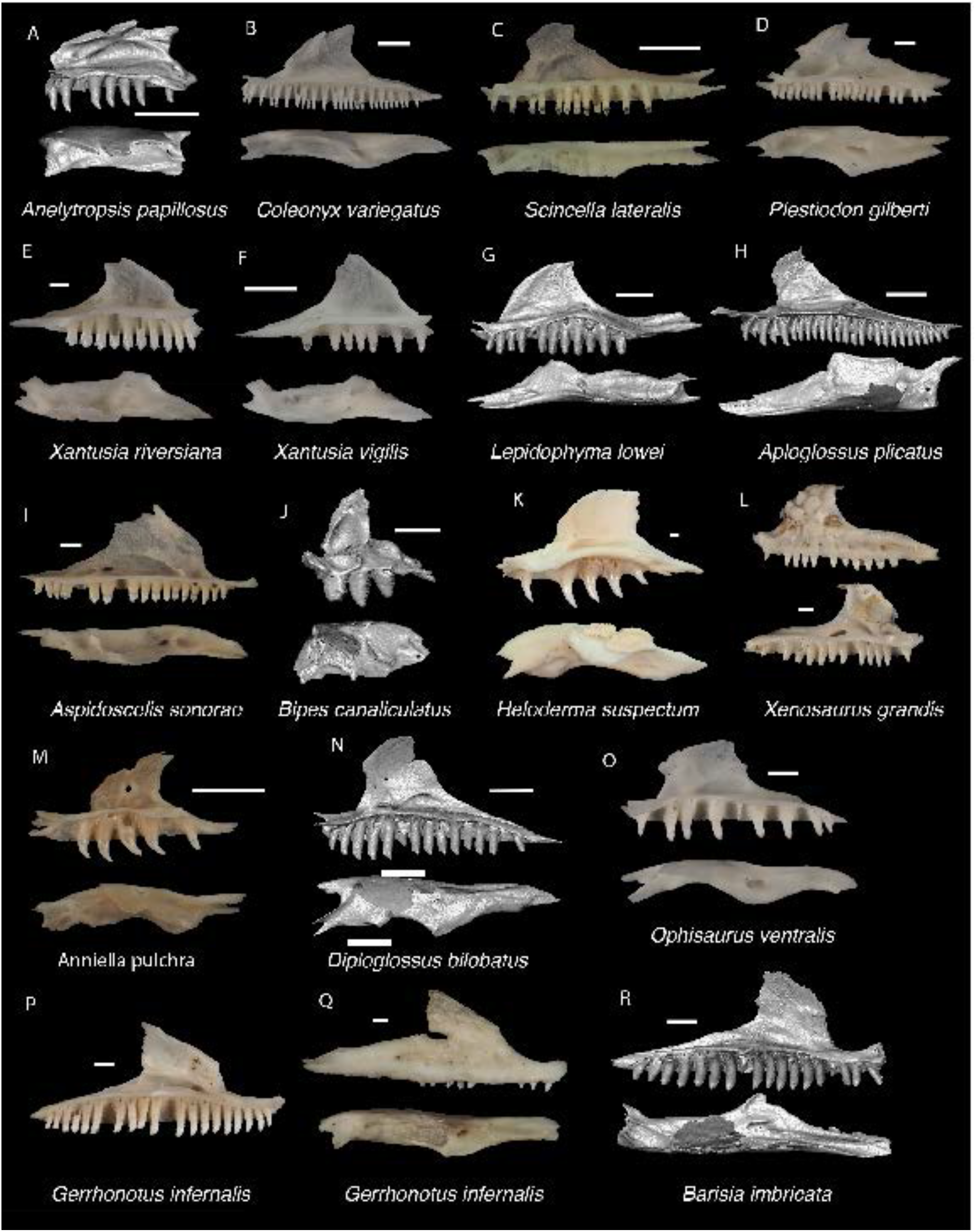
Non-Pleurodontan Maxillae. Maxillae in medial and dorsal views– **A**. *Anelytropsis papillosus* UF 86708; **B**. *Coleonyx variegatus* TxVP M-12109; **C**. *Scincella lateralis* TxVP M-4489; **D**. *Plestiodon gilberti* TxVP M-8587; **E**. *Xantusia riversiana* TxVP M-8505; **F**. *Xantusia virigatus* TxVP M-12130; **G**. *Lepidophyma lowei* LACM 143367; **H**. *Aploglossus plicatus* TNHC 34481; **I**. *Aspidoscelis sonorae* TxVP M-15670; **J**. *Bipes canaliculatus* CAS 134753; **K**. *Heloderma suspectum* TxVP M-9001; **L**. *Xenosaurus grandis* TxVP M-8960; **M**. *Anniella pulchra* TxVP M-8678; **N**. *Diploglossus bilobatus* TNHC 31933; **O**. *Ophisaurus ventralis* TxVP M-8585; **P**. *Gerrhonotus infernalis* TxVP M-13441; **Q**. *Gerrhonotus infernalis* TxVP M-13440; **R**. *Barisia imbricata* TNHC 76984. Scale bars = 1 mm.

### Frontal

#### Description

TxVP 41229-28737 is an unfused right frontal (Fig. 32). There are osteoderms with a pitted texture fused to the frontal much of its dorsal surface. The lateral frontal sulcus separates the larger frontal shield from the smaller posterolateral frontoparietal shield, and the medial frontal sulcus separates a minute interfrontal shield (*sensu* Klembara et al. 2017). Anterolaterally, there is a prefrontal facet, and anterodorsally, there is a nasal facet. The anterior end is elongated and pointed without a distinct anterolateral process. The medial margin and interorbital margins are straight, and the posterolateral processes gently curve laterally. The posterior margin is straight with a posterolateral parietal articulation facet. There is a postfrontal facet along the posterolateral edge. The anteroventral portion of the crista cranii is broken, but the crest is well-developed, tall, and anteroposteriorly long.

#### Identification

The fossil shares with some anguimorphs and most scincids co-ossified osteoderms (Estes et al. 1988), well-developed and ventrally directed cristae cranii (=subolfactory processes of McDowell and Bogert 1954), and an unfused frontal (Estes et al. 1988). The sculpting on the dorsal surface in scincids is more vermiculate compared to the fossil and to extant anguimorphs which have a pitted texture, such as *Ophisaurus* and *Elgaria* (Conrad 2008; Fig. 35). *Plestiodon* differ in having exceptionally slender cristae cranii (Nash 1970), and *Scincella* and *Mabuya* differ in having a fused frontal (Greer 1970). Examined *Plestiodon* also differs from the fossil and anguines, except for *Anguis fragilis* (Klembara et al. 2017), in having a broader and less pointed anterior portion of the unfused frontal. Frontals are assigned to Anguimorpha. Among NA anguimorphs, unfused frontals occur in Anguinae, Diploglossinae, *Heloderma*, and *Anniella* (Evans 2008). *Heloderma* and *Anniella* differ in having the cristae cranii curve to meet at the midline, forming an enclosed olfactory canal (Evans 2008). In the fossil and other anguines except for *Anguis fragilis* (Klembara et al. 2017), there is a pointed anterior portion of the unfused frontal. The anterior portion of the frontal is broader in diploglossines (Evans 2008). Frontals are assigned to Anguinae based on the presence of an unfused frontal and a pointed anterior portion of the frontal.

**Figure 35.**
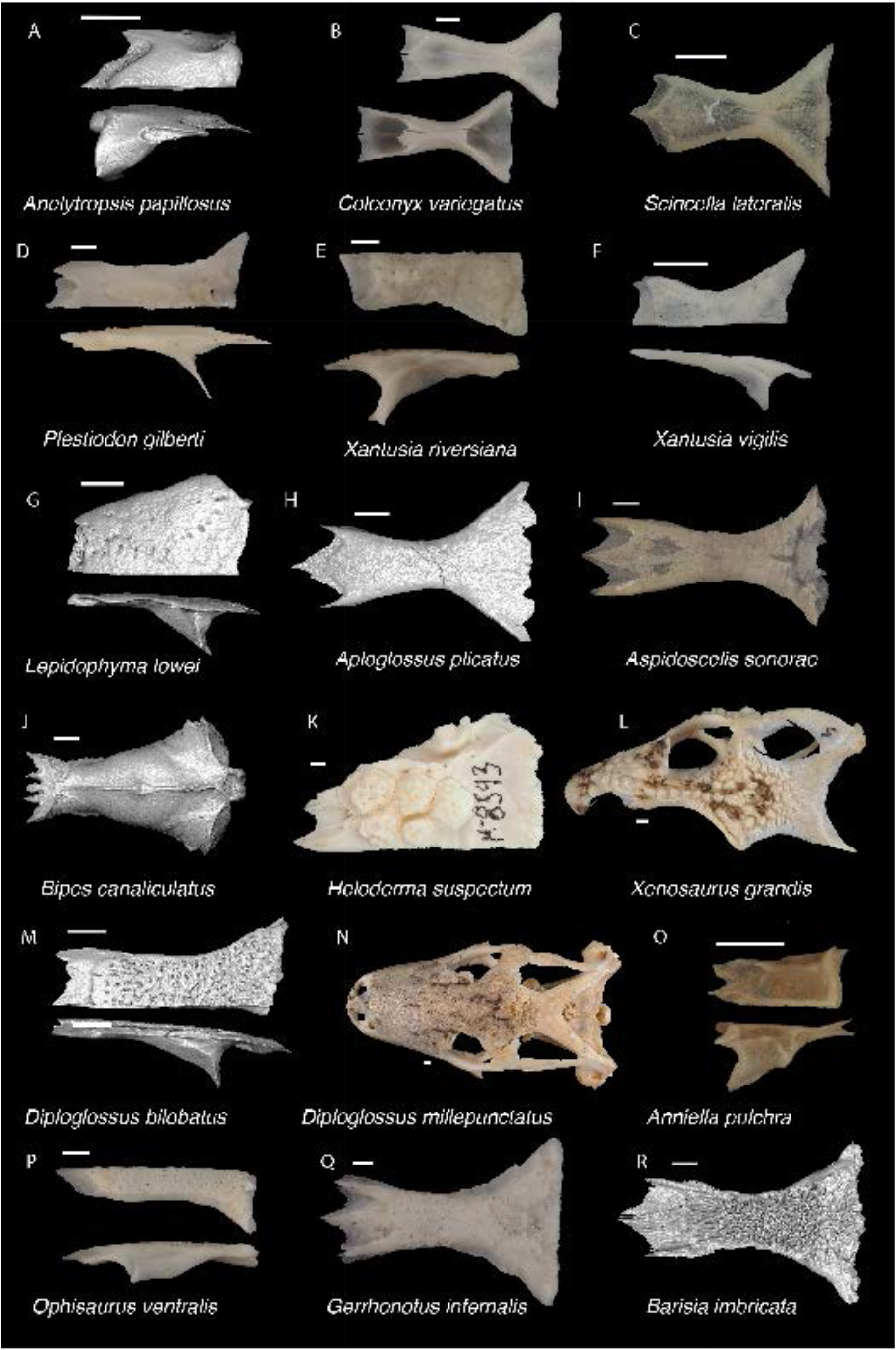
Non-Pleurodontan Frontals. Frontals in dorsal and lateral views– **A**. *Anelytropsis papillosus* UF 86708; **B**. *Coleonyx variegatus* TxVP M-12109; **C**. *Scincella lateralis* TxVP M-4489; **D**. *Plestiodon gilberti* TxVP M-8587; **E**. *Xantusia riversiana* TxVP M-8505; **F**. *Xantusia virigatus* TxVP M-12130; **G**. *Lepidophyma lowei* LACM 143367; **H**. *Aploglossus plicatus* TNHC 34481; **I**. *Aspidoscelis sonorae* TxVP M-15670; **J**. *Bipes canaliculatus* CAS 134753; **K**. *Heloderma suspectum* TxVP M-8593; **L**. *Xenosaurus grandis* TxVP M-8960; **M**. *Diploglossus bilobatus* TNHC 31933; **N**. *Diploglossus millepunctatus* TxVP M-9010; **O**. *Anniella pulchra* TxVP M-8678; **P**. *Ophisaurus ventralis* TxVP M-8585; **Q**. *Gerrhonotus infernalis* TxVP M-13441; **R**. *Barisia imbricata* TNHC 76984. Scale bars = 1 mm.

### Parietal

#### Description

TxVP 41229-28153 serves as the basis for our description and is a parietal missing only the ends of the postparietal processes (Fig. 32). The parietal table is rectangular and is covered dorsally in co-ossified osteoderms with a pitted texture. The interparietal sulcus separates the triangular interparietal shield from the lateral shields (*sensu* Klembara et al 2017). The interparietal shield reaches the posterior smooth region of the parietal table. The anterior edge is straight with small frontal tabs and interlacing articulation facets for the frontal. The posterior edge between the postparietal processes is characterized by two small depressions (nuchal fossae) separated by a small ridge. The postparietal processes are broad and flat at the bases. The ventrolateral crests are low, positioned along the lateral margins, and border the cerebral vault. There is a low ridge anterior to the pit for the processus ascendens. The ventrolateral crests curve medially onto the postparietal processes, and together with a ventrolateral ridge on the postparietal process, define distinct depressions. There is a large parietal foramen within the interparietal shield.

#### Identification

The fossil shares with some anguimorphs and scincids co-ossified osteoderms with dorsal sculpting (Estes et al. 1988), a parietal foramen enclosed by the parietal (Estes et al. 1988), and ventrally projecting parietal crests or processes (Estes et al. 1988; Evans 2008; Ledesma and Scarpetta 2018; Fig. 36). Scincids differ in having long posterior projections (median extensions of Evans 2008) on the posterior edge of the parietal table between the postparietal processes (Evans 2008; Gauthier et al. 2012; see also fig. 8 in Jerez et al. 2015). Scincids further differ from the fossil in having distinct ventrolateral crests that include long, thin, ventral projections (Nash 1970; Evans 2008) and the postparietal processes in examined scincids are more separated relative to the fossil. Parietals are assigned to Anguimorpha. *Heloderma* differs in lacking a parietal foramen (Estes et al. 1988) and *Anniella* differs in having the ventrally projecting parietal crests developed into extensive sheets of bone (Evans 2008). Xenosaurids differ in having heavily sculptured dorsal roofing bones with many bumpy, dome-like co-ossified osteoderms (Gauthier 1982; Bhullar 2010, 2011). Parietals are assigned to Anguidae.

**Figure 36.**
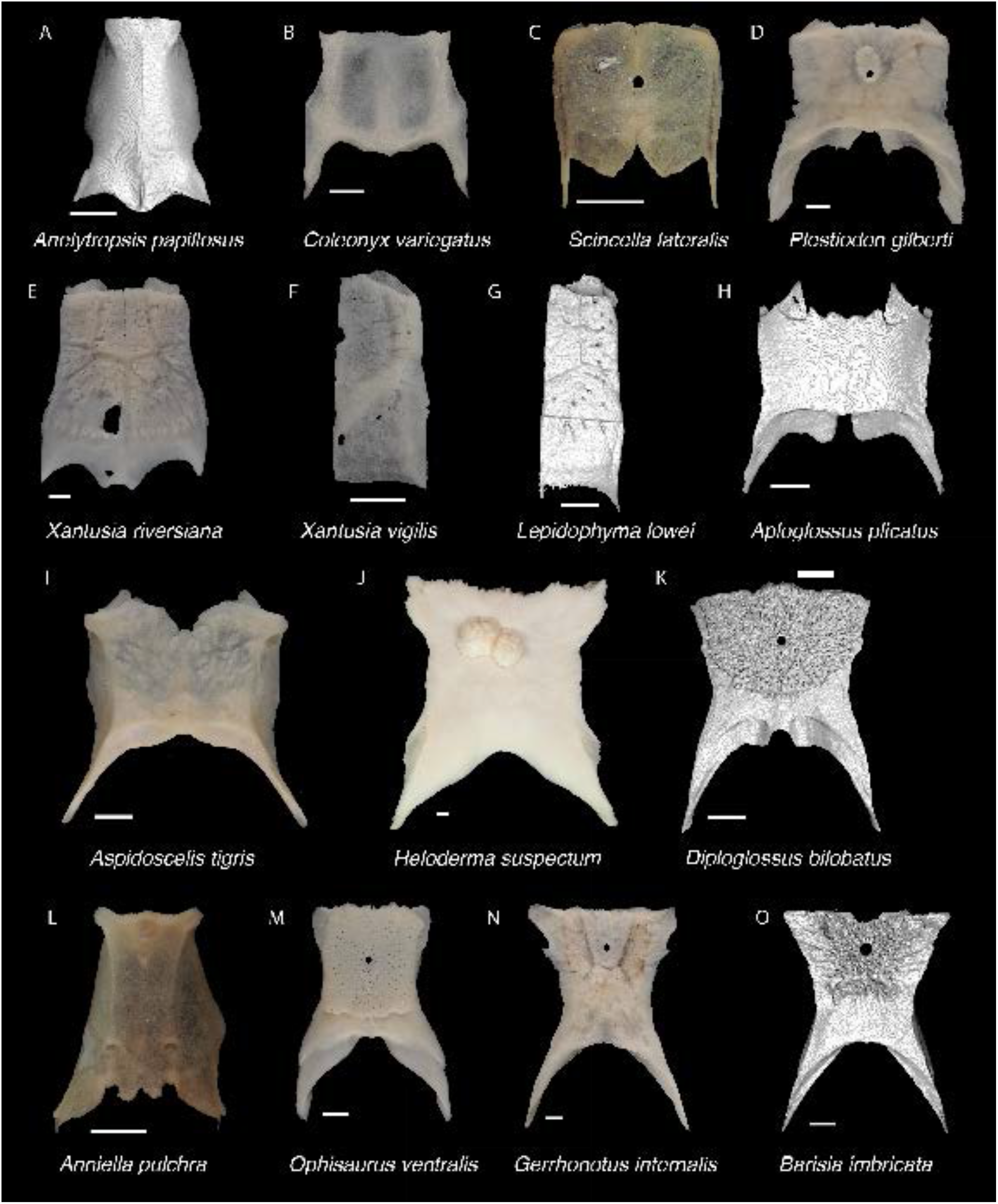
Non-Pleurodontan Parietals. Parietals in dorsal view– **A**. *Anelytropsis papillosus* UF 86708; **B**. *Coleonyx variegatus* TxVP M-12109; **C**. *Scincella lateralis* TxVP M-4489; **D**. *Plestiodon gilberti* TxVP M-8587; **E**. *Xantusia riversiana* TxVP M-8505; **F**. *Xantusia virigatus* TxVP M-12130; **G**. *Lepidophyma lowei* LACM 143367; **H**. *Aploglossus plicatus* TNHC 34481; **I**. *Aspidoscelis tigris* TxVP M-15667; **J**. *Heloderma suspectum* TxVP M-8593; **K**. *Diploglossus bilobatus* TNHC 31933; **L**. *Anniella pulchra* TxVP M-8678; **M**. *Ophisaurus ventralis* TxVP M-8585; **N**. *Gerrhonotus infernalis* TxVP M-13441; **O**. *Barisia imbricata* TNHC 76984. Scale bars = 1 mm.

The lateral margins of the parietal table are generally more concave in gerrhonotines and diploglossines, whereas in the fossil and in anguines, the lateral margins are straighter (Evans 2008). Furthermore, examined gerrhonotines and diploglossines usually have a larger smooth area on the posterior portion of the parietal table and a more rounded posterior terminus of the interparietal shield (see also fig. 9 in Ledesma et al. 2021). The fossil shares with examined *Ophisaurus* a small posterior smooth area on the parietal table and a pointed posterior terminus of the interparietal shield (see also figs. 2-4 in Klembara 2015). Fossils are assigned to Anguinae.

### Prefrontal

#### Description

TxVP 41229-25597 is a left prefrontal (Fig. 32). It is triradiate with a long and pointed orbital process, a short ventral process, and an anterior sheet. The anterior sheet is slightly broken and has a broad articulation facet for the facial process of the maxilla. There is a small ridge on the lateral surface near the base of the orbital process. The ventral process is missing the posteroventral tip but is narrow and squared off. There is a distinct notch for the lacrimal foramen, and the ventral process forms the posterior border of the foramen. Dorsal to the lacrimal foramen notch is a small overhanging lamina. Medially, the boundary of the olfactory chamber is a smooth, rounded, and concave surface. Dorsal to the olfactory chamber is a shallow groove for articulation with the frontal. The orbitonasal flange is broad with a distinct medial projection for articulation with the palatine.

#### Identification

The fossil differs from NA pleurodontans in lacking a lateral prefrontal boss (Estes et al. 1988; Smith 2009b; Smith 2011; Gauthier et al. 2012; Fig. 37), lacking a strong lateral canthal ridge (reported in *Anolis* and *Polychrus*; Smith 2011), lacking a supraorbital spine (present in *Phrynosoma* and *Corytophanes*; Smith 2009a), and lacking a thin, crescent shape with a distinct laterally projecting lamina (present in examined phrynosomatines). NA teiids and gymnophthalmoids differ in having a distinct laterally projecting lamina (lacrimal flange of Bell et al. 2003) with a distinct articulation facet for the facial process of the maxilla (Tedesco et al. 1999). Examined scincids differ in having a relatively shorter orbital process and, excluding *Scincella*, a more elongate, oblong anterior process (Nash 1970). Xantusiids differ in having the lacrimal fused to the prefrontal (Savage 1963) with a suborbital foramen nearly or entirely enclosed within the prefrontal, and in having a distinct vertical articulation ridge or flange (e.g., in *X. riversiana*) that articulates with the maxilla. Examined *Coleonyx* differ in having an orbitonasal flange that extends farther medially. *Sphaerodactylus roosevelti* has a smaller notch for the lacrimal foramen (Daza et al. 2008). Based on differences with other NA lizards, the fossil is identified to Anguimorpha. *Heloderma* differs in having a much broader orbital process and the prefrontal of *Anniella* is much smaller than the fossil. Diploglossines and gerrhonotines differ in having an orbitonasal flange that does not extend as far medially as in the fossil. The fossil is assigned to Anguinae.

**Figure 37.**
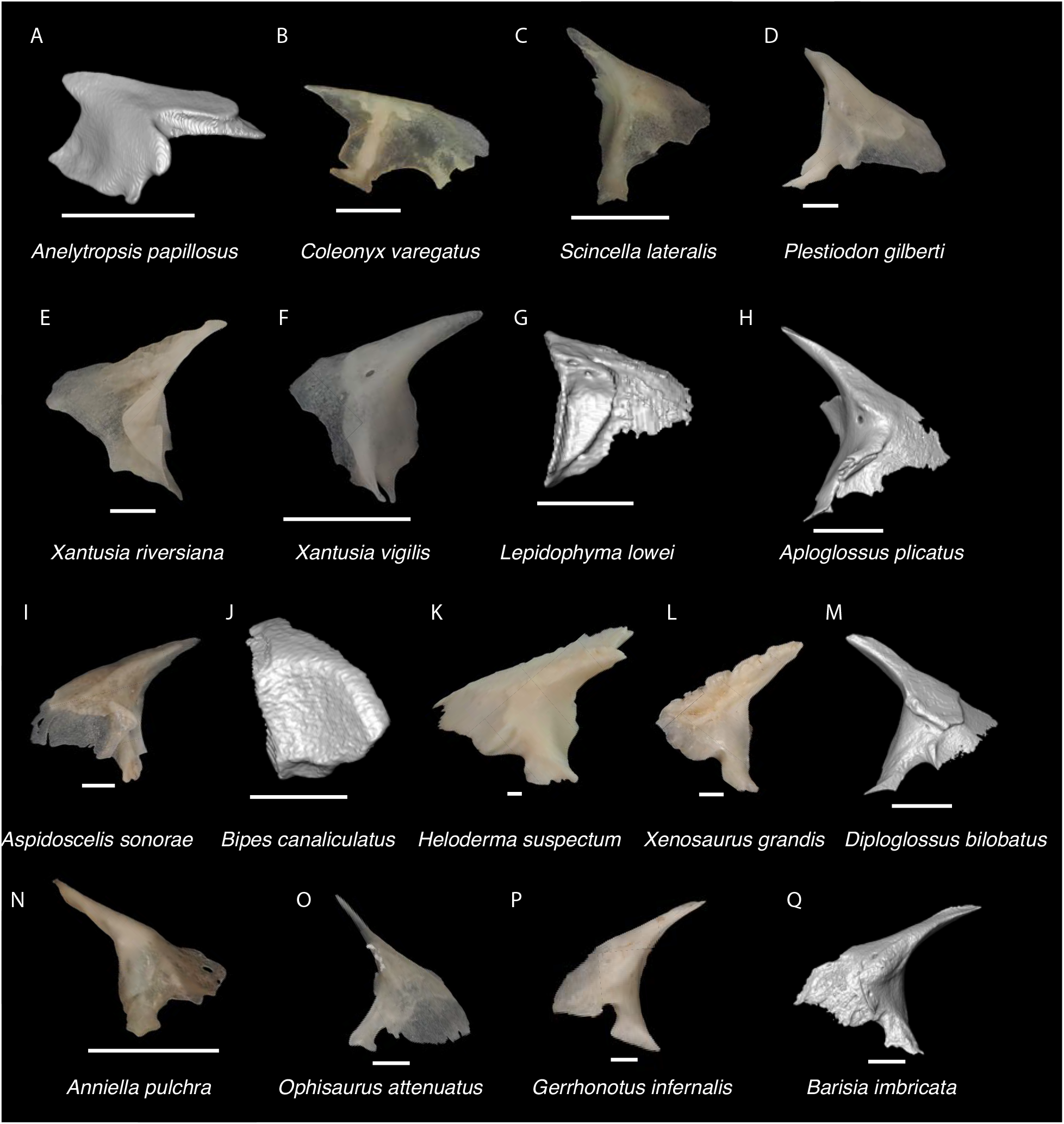
Non-Pleurodontan Prefrontals. Prefrontals in lateral view– **A**. *Anelytropsis papillosus* UF 86708; **B**. *Coleonyx variegatus* TxVP M-13892; **C**. *Scincella lateralis* TxVP M-4489; **D**. *Plestiodon gilberti* TxVP M-8587; **E**. *Xantusia riversiana* TxVP M-8505; **F**. *Xantusia virigatus* TxVP M-12130; **G**. *Lepidophyma lowei* LACM 143367; **H**. *Aploglossus plicatus* TNHC 34481; **I**. *Aspidoscelis sonorae* TxVP M-15670; **J**. *Bipes canaliculatus* CAS 134753; **K**. *Heloderma suspectum* TxVP M-9001; **L**. *Xenosaurus grandis* TxVP M-8960; **M**. *Diploglossus bilobatus* TNHC 31933; **N**. *Anniella pulchra* TxVP M-8678; **O**. *Ophisaurus attenuatus* TxVP M-8979; **P**. *Gerrhonotus infernalis* TxVP M-13441; **Q**. *Barisia imbricata* TNHC 76984. Scale bars = 1 mm. Figure 41. Non-Pleurodontan Pterygoids. Pterygoids in dorsal view– **A**. *Anelytropsis papillosus* UF 86708; **B**. *Coleonyx variegatus* TxVP M-12109; **C**. *Scincella lateralis* TxVP M-4489; **D**. *Plestiodon gilberti* TxVP M-8587; **E**. *Xantusia riversiana* TxVP M-8505; **F**. *Xantusia virigatus* TxVP M-12130; **G**. *Lepidophyma lowei* LACM 143367; **H**. *Aploglossus plicatus* TNHC 34481; **I**. *Aspidoscelis sonorae* TxVP M-15670; **J**. *Bipes canaliculatus* CAS 134753; **K**. *Heloderma suspectum* TxVP M-9001; **L**. *Xenosaurus grandis* TxVP M-8960; **M**. *Diploglossus bilobatus* TNHC 31933; **N**. *Anniella pulchra* TxVP M-8678; **O**. *Ophisaurus attenuatus* TxVP M-8979; **P**. *Gerrhonotus infernalis* TxVP M-13441; **Q**. *Barisia imbricata* TNHC 76984. Scale bars = 1 mm.

### Postfrontal

#### Description

TxVP 41229-28467 is a right postfrontal (Fig. 32). It is a triradiate, delicate bone with a thin, elongate anterior process, a small, rounded lateral process, and an elongate posterior process. There is a jugal articulation facet on the lateral process and a postorbital articulation facet along the lateral margin of the posterior process. The medial margin is widely curved to clasp the frontal and parietal. There is a large dorsal foramen on the posterior process.

#### Identification

Most NA pleurodontans differ from the fossils in either lacking a postfrontal or having relatively small postfrontal that lacks a facet for clasping the fronto-parietal articulation (Evans 2008; Smith 2009a). Some iguanids (e.g., *Sauromalis ater* TNHC 18483) have a comparatively larger postfrontal that does clasp the fronto-parietal articulation; however, the posterior process is shorter compared to the fossil. *Xantusia*, *Xenosaurus*, and NA teiids differ in having a fused postorbitofrontal (Conrad 2008; Fig. 38). *Coleonyx variegatus* and *C. brevis* differ in lacking a lateral process (Kluge 1962). *Coleonyx elegans* and *C. mitratus* reportedly have a lateral projection (Kluge 1962), but similar to *Sphaerodactylus roosevelti* (Daza et al. 2008) and *Phyllodactylus baurii*, the lateral process is much shorter than in the fossil. Examined *Plestiodon* differ in lacking a distinct facet on the lateral projection for articulation with the jugal, and *Scincella* and *Mabuya* differ in having a much longer posterior process compared to the anterior process (see also fig. 8 in Jerez et al. 2015). Some gymnophthalmoids have a separate, triradiate postorbital (Evans 2008); however, to our knowledge, none have the large dorsal foramen on the posterior process that is present in the fossil and several anguimorphs (Klembra et al. 2017; Ledesma et al. 2021). Based on these differences, fossils were assigned to Anguimorpha. *Anniella* differs in lacking a lateral process (Torien 1950; Rieppel 1980) and *Heloderma* differs in having a broad postorbitofrontal (Rieppel 1980; Evans 2008). Fossils were assigned to Anguidae. Some diploglossines differ in having a fused postorbitofrontal (Evans 2008). Examined *Diploglossus* with a separate postfrontal differ from the fossil in having a wider posterior process. Examined gerrhonotines differ in having the angle between the anterior and posterior processes closer to 90 degrees (see also fig. 17 in Ledesma et al. 2021). Fossils are assigned to Anguinae.

**Figure 38.**
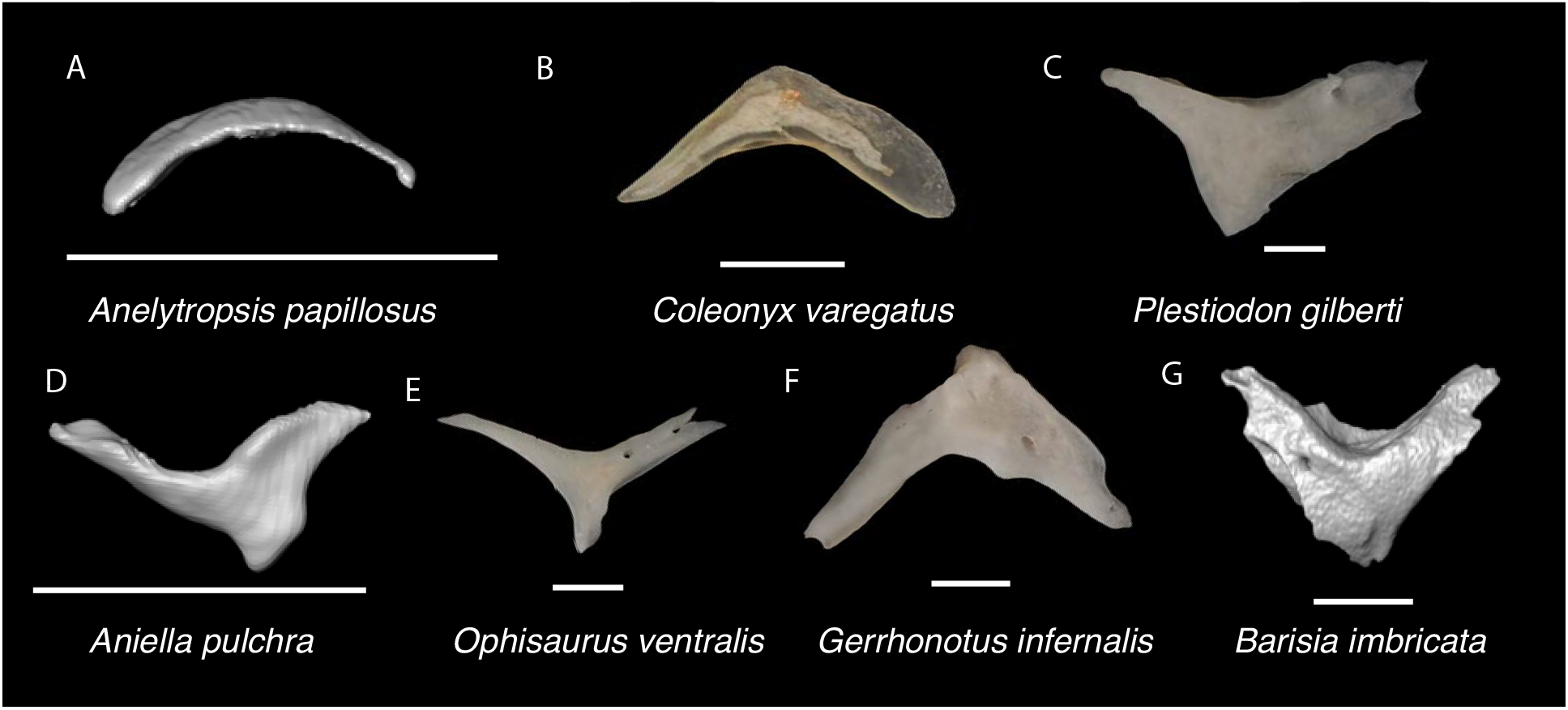
Non-Pleurodontan Postfrontals. Postfrontals in dorsal view– **A**. *Anelytropsis papillosus* UF 86708; **B**. *Coleonyx varegatus* TxVP M-13892; **C**. *Plestiodon gilberti* TxVP M-8587; **D**. *Anniella pulchra* FMNH 130479; **E**. *Ophisaurus ventralis* TxVP M-8585; **F**. *Gerrhonotus infernalis* TxVP M-13441; **G**. *Barisia imbricata* TNHC 76984. Scale bars = 1 mm unless otherwise noted.

### Quadrate

#### Description

TxVP 41229-25585 is a right quadrate (Fig. 32). The bone is thin and parallel-sided. There is no pterygoid lappet. There is a moderately developed medial crest that is directed anteriorly. The conch is deep and narrow. The cephalic condyle projects posteriorly without extensive dorsal ossification. There is an anteriorly expanded dorsal tubercle. There is a quadrate foramen on the anteroventral surface and a foramen medial to the central column. TxVP 41229-25582 differs in having two quadrate foramina on the anterior surface.

#### Identification

The fossil shares with geckos, *Scincella*, xantusiids, some pleurodontans, and anguimorphs except for *Heloderma* the absence of a distinct pterygoid lappet (Estes et al. 1988; Evans 2008; Fig. 39). Examined *Scincella* differ in having a curved tympanic crest. Examined *Xantusia riversiana* and *Lepidophyma lowei* differ in having a slightly more curved tympanic crest and examined *Xantusia vigilis* have a much more slender quadrate (see also figs. 16-17 in Savage 1963). Examined *Coleonyx* differ in having a more narrow quadrate that slightly narrows ventrally and has a notched dorsolateral margin (Kluge 1967). *Sphaerodactylus roosevelti* differs in having a more narrow quadrate with a curved tympanic crest (Daza et al. 2008). Most examined NA pleurodontans differ in having the dorsal portion much wider compared to the articular surface. In *Anolis* and *Uma*, the lateral margins are parallel but examined *Anolis* differ in having a distinct boss at the ventromedial margin of the quadrate and *Uma* differ in having a laterally slanted cephalic condyle and central column. Fossils were assigned to Anguimorpha.

**Figure 39.**
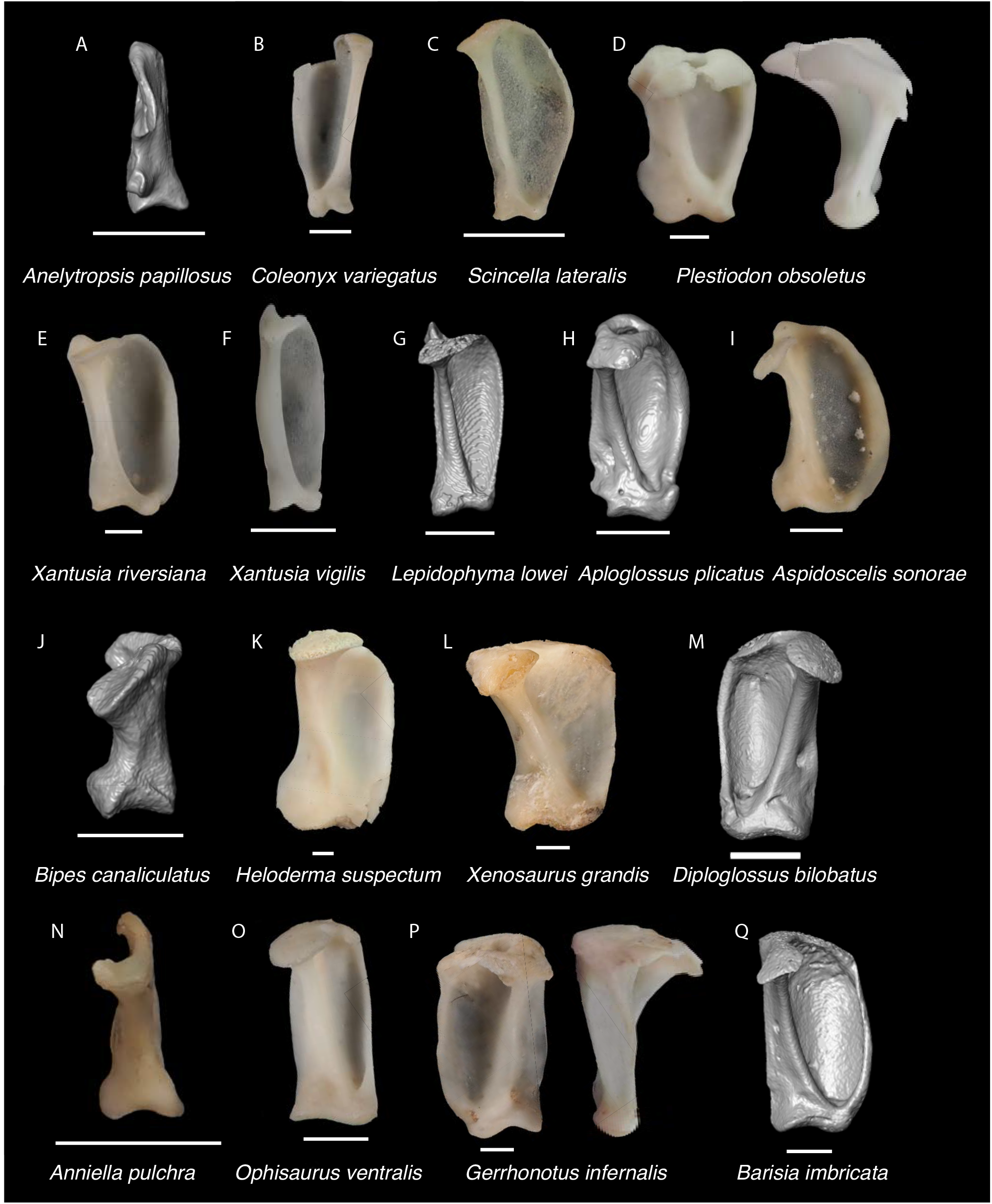
Non-Pleurodontan Quadrates. Quadrates in posterior and lateral views– **A**. *Anelytropsis papillosus* UF 86708; **B**. *Coleonyx variegatus* TxVP M-12109; **C**. *Scincella lateralis* TxVP M-4489; **D**. *Plestiodon obsoletus* TxVP M-8574; **E**. *Xantusia riversiana* TxVP M-8505; **F**. *Xantusia virigatus* TxVP M-12130; **G**. *Lepidophyma lowei* LACM 143367; **H**. *Aploglossus plicatus* TNHC 34481; **I**. *Aspidoscelis sonorae* TxVP M-15670; **J**. *Bipes canaliculatus* CAS 134753; **K**. *Heloderma suspectum* TxVP M-8593; **L**. *Xenosaurus grandis* TxVP M-8960; **M**. *Diploglossus bilobatus* TNHC 31933; **N**. *Anniella pulchra* TxVP M-8678; **O**. *Ophisaurus ventralis* TxVP M-8585; **P**. *Gerrhonotus infernalis* TxVP M-13441; **Q**. *Barisia imbricata* TNHC 76984. Scale bars = 1 mm.

*Xenosaurus* differs from the fossils in having a wider quadrate with a shallower lateral conch (Bhullar 2011). *Anniella* differs in having a thin quadrate in posterior view that is widened in medial and lateral views (Toerien 1950). Fossils were assigned to Anguidae. Examined gerrhonotines (see fig. 19 in Ledesma and Scarpetta 2021) and diploglossines (see also fig. 2 in Bochaton et al. 2016b) differ from the fossil and examined *Ophisaurus* in having a more laterally extensive pterygoid lamina and a more curved tympanic crest. Fossils are assigned to Anguinae.

### Pterygoid

#### Description

TxVP 41229-28356 is a left pterygoid that is missing the distal end of the palatine process (Fig. 32). The palatine process is narrow with an anterior palatine facet. The transverse process is pointed and extends anterolaterally. The transverse process bears a ridge on the dorsal surface for insertion of the superficial pseudotemporal muscle and a developed ectopterygoid facet. There is a large ridge on the ventral surface for insertion of the pterygomandibular muscle, and on the dorsal surface there is a small, deep fossa columella. The epipterygoid crest is anteromedial to the fossa columella. There is a long postepipterygoid groove (*sensu* Good 1987) on the quadrate process. The quadrate process is elongated, and the medial surface has a shallow groove that serves for insertion of the pterygoideus muscle. There is a small notch in the quadrate process that is likely taphonomic. There is a small medial projection at the floor of the basipterygoid fossa and a broad patch of 19 pterygoid tooth positions with 13 pterygoid teeth present. TxVP 41229-25602 is missing the distal ends of the palatine and quadrate processes and does not differ from TxVP 41229-28356 substantively.

#### Identification

Fossils share with some anguimorphs and pleurodontans a medial projection at the floor of the basipterygoid fossa (Evans 2008; Smith 2009b). Examined NA pleurodontans, except *Petrosaurus*, differ in having the transverse process oriented more medially (Fig. 40). Examined *Petrosaurus* differ in lacking pterygoid teeth (Etheridge 1964) and lacking a long postepiterygoid groove on the quadrate process. Fossils are assigned to Anguimorpha.

*Xenosaurus* and *Anniella* lackpterygoid teeth (Evans 2008). *Heloderma* differs in having only a single row of pterygoid teeth located on an elevated ridge (Evans 2008). Thus, fossils are assigned to Anguidae. The presence and number of teeth are variable in gerrhonotines (Ledesma et al. 2021; Scarpetta et al. 2021) and *Ophisaurus* (Taylor 1940), but diploglossines lack pterygoid teeth (Rieppel 1980; Evans 2008). The epipterygoid crest in examined anguines is more distinct and projects farther medially compared to examined gerrhonotines (Klembara et al. 2017; Ledesma et al. 2021). The palatal process is narrower in examined *Ophisaurus* compared to gerrhonotines and the space between the transverse process and the palatal plate (suborbital incisure of Klembara et al. 2017) is wider in examined gerrhonotines and diploglossines compared to *Ophisaurus*, thus fossils are assigned to Anguinae.

### Dentary

#### Description

TxVP 41229-28388 serves as the basis for our description and is a left dentary with 18 tooth positions and 14 teeth present (Fig. 32). Teeth are unicuspid, but the crowns are slightly eroded. There are three posterolateral processes, including a pointed coronoid process, a long surangular process, and a short angular process. Medially, a minute surangular spine is visible (*sensu* Klembara et al. 2014). The Meckelian canal is open ventrally. The intramandibular septum has a free posteroventral margin. The dental shelf is narrow, and there is a posteriorly projecting splenial spine (*sensu* Klembara et al. 2014). There are seven nutrient foramina arranged in a row on the lateral surface. TxVP 41229-27759 differs in lacking an intramandibular septum with a free posteroventral margin and having a long surangular spine.

#### Identification

Dentaries are assigned to Anguimorpha based on the presence of a discrete surangular process (Gauthier 1982; Good 1988) and a posteriorly projecting splenial spine (Conrad 2008; Čerňanský and Augé 2019; Fig. 41). Dentaries are assigned to Anguidae based on the presence of an intramandibular septum with a free posteroventral margin although this morphology is absent in some anguines (Klembara et al. 2014; Scarpetta 2018; Scarpetta et al. 2021) and is indeed absent in TxVP 41229-27759. Fossils differ from other NA anguimorphs except for *Ophisaurus* in having a surangular spine (Klembara et al. 2014). A surangular spine was reported in some diploglossines (Bochaton et al. 2016b; Syromyatnikova and Aranda 2022); however, this feature is likely a posterior extension on the intramandibular septum described in other anguids (Klembara et al. 2014; Ledesma and Scarpetta 2018). Furthermore, diploglossines differ from the fossils in lacking a splenial spine and having the anterior inferior alveolar foramen completely within the splenial (Syromyatnikova and Aranda 2022). Fossils are assigned to Anguinae.

**Figure 40.**
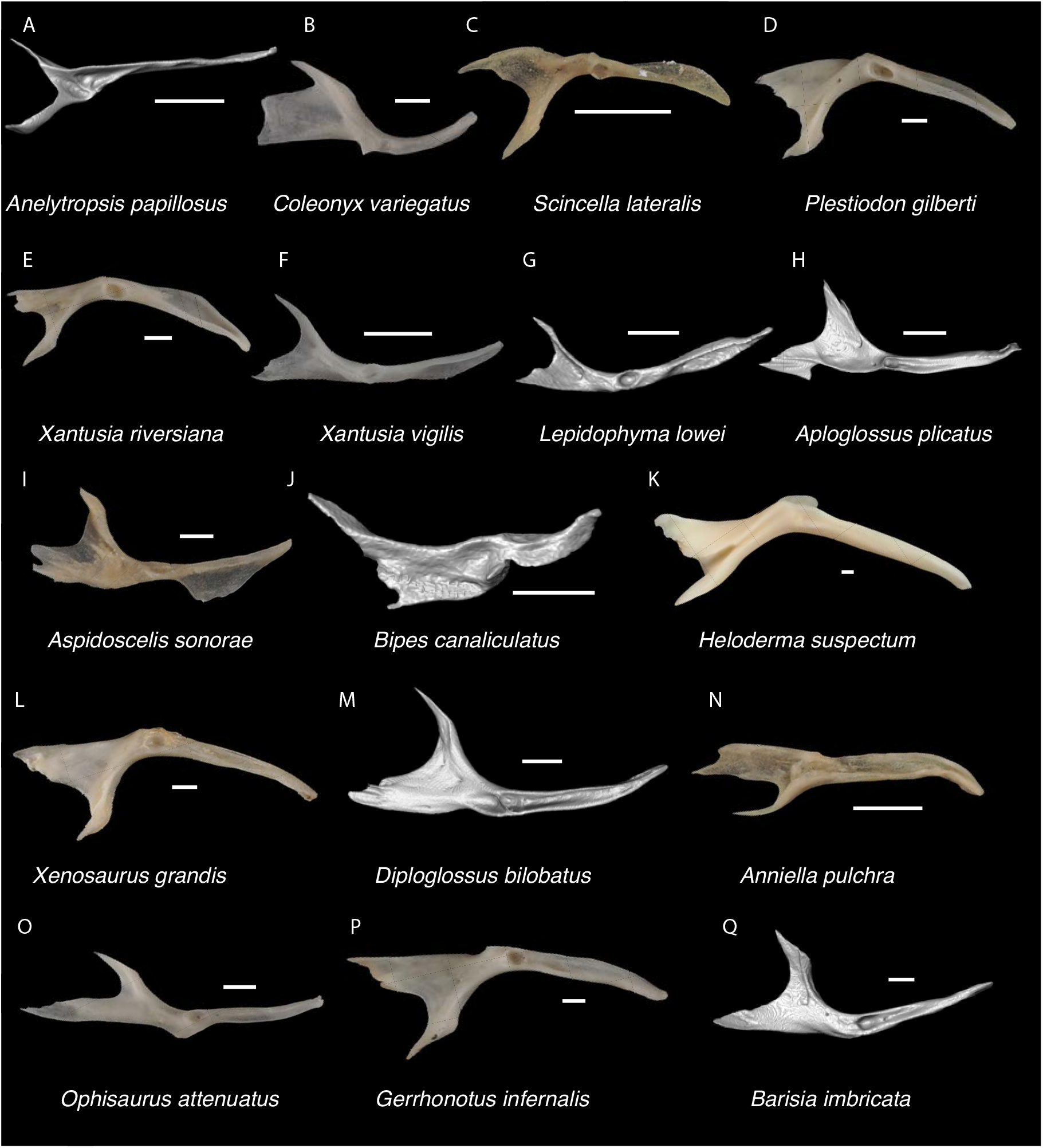
Non-Pleurodontan Pterygoids. Pterygoids in dorsal view– **A**. *Anelytropsis papillosus* UF 86708; **B**. *Coleonyx variegatus* TxVP M-12109; **C**. *Scincella lateralis* TxVP M-4489; **D**. *Plestiodon gilberti* TxVP M-8587; **E**. *Xantusia riversiana* TxVP M-8505; **F**. *Xantusia virigatus* TxVP M-12130; **G**. *Lepidophyma lowei* LACM 143367; **H**. *Aploglossus plicatus* TNHC 34481; **I**. *Aspidoscelis sonorae* TxVP M-15670; **J**. *Bipes canaliculatus* CAS 134753; **K**. *Heloderma suspectum* TxVP M-9001; **L**. *Xenosaurus grandis* TxVP M-8960; **M**. *Diploglossus bilobatus* TNHC 31933; **N**. *Anniella pulchra* TxVP M-8678; **O**. *Ophisaurus attenuatus* TxVP M-8979; **P**. *Gerrhonotus infernalis* TxVP M-13441; **Q**. *Barisia imbricata* TNHC 76984. Scale bars = 1 mm.

**Figure 41.**
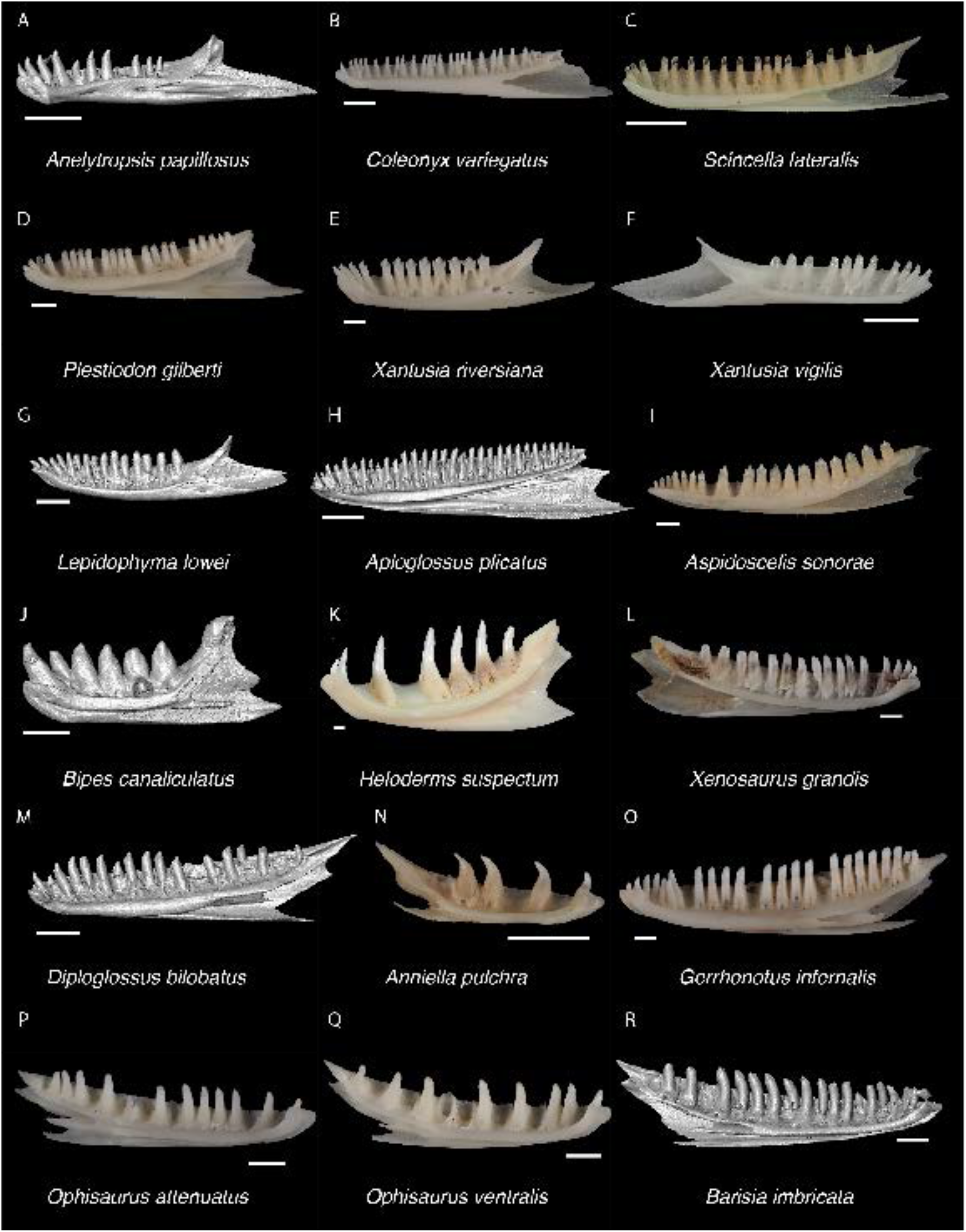
Non-Pleurodontan Dentaries. Dentaries in medial view– **A**. *Anelytropsis papillosus* UF 86708; **B**. *Coleonyx variegatus* TxVP M-12109; **C**. *Scincella lateralis* TxVP M-4489; **D**. *Plestiodon gilberti* TxVP M-8587; **E**. *Xantusia riversiana* TxVP M-8505; **F**. *Xantusia virigatus* TxVP M-12130; **G**. *Lepidophyma lowei* LACM 143367; **H**. *Aploglossus plicatus* TNHC 34481; **I**. *Aspidoscelis sonorae* TxVP M-15670; **J**. *Bipes canaliculatus* CAS 134753; **K**. *Heloderma suspectum* TxVP M-9001; **L**. *Xenosaurus grandis* TxVP M-8960; **M**. *Diploglossus bilobatus* TNHC 31933; **N**. *Anniella pulchra* TxVP M-8678; **O**. *Gerrhonotus infernalis* TxVP M-13441; **P**. *Ophisaurus attenuatus* TxVP M-8979; **Q**. *Ophisaurus ventralis* TxVP M-8585; **R**. *Barisia imbricata* TNHC 76984. Scale bars = 1 mm.

### Coronoid

#### Description

TxVP 41229-28445 is a left coronoid (Fig. 32). The coronoid process is short and rounded and the anteromedial process is elongated but is missing the distal tip. The posteromedial process is directed posteriorly and has a notch on the posterodorsal end. There is a short medial crest that extends from the coronoid process and diminishes on the posteromedial process. There is a large, anteriorly projecting lateral process. The anteromedial process has a medial splenial facet that, together with an anteriorly projecting lateral process, forms a narrow articulation facet for the coronoid process of the dentary. There is no lateral crest and the facet for the dorsal articulation with the surangular is narrow.

#### Identification

The fossil coronoid differs from xantusiids and some pleurodontans in having an anterolateral process (Estes et al. 1988). The fossil differs from examined NA pleurodontans with an anterolateral process (see discussion in Crotaphytidae section above) in having a more posteriorly oriented posteromedial process. The fossil differs from examined *Aspidoscelis* and *Ameiva* in lacking a deeply notched posterior edge forming dorsal and ventral rami (Tedesco et al. 1999; Fig. 30). Furthermore, *Aspidoscelis*, *Ameiva*, and *Pholidoscelis* differ in having a lateral crest running from the apex of the coronoid process anteroventrally onto the anterolateral process (Tedesco et al. 1999; Bochaton et al. 2019). The fossil shares with several gymnophthalmoids a low coronoid process and widely divergent anteromedial and posteromedial processes (Evans 2008; Roscito and Rodrigues 2010). The fossil differs from examined gymnophthalmoids with a low coronoid process in having a relatively broader posteromedial process (see figs. 7-9 in Roscito and Rodrigues 2010; fig. 2 in Holovacs et al. 2020). Examined *Coleonyx variegatus*, *C. brevis*, *Sphaerodactylus roosevelti*, and *Thecadactylus rapicauda* differ from the fossil in having a thinner posteromedial process (Kluge 1962; Daza et al. 2008; Bochaton et al. 2018). The posteromedial process is slightly wider in *C. elegans* and *C. mitratus* compared to other species (Kluge 1962) but examined *Coleonyx* including *C. elegans* differ in having a pronounced lateral crest on the coronoid process (see also fig. 7 in Kluge 1962). The lateral crest on the coronoid process is absent in the fossil, but in examined *Plestiodon* and *Scincella,* the crest is pronounced and obliquely oriented. Furthermore, the anterolateral process is more anteriorly oriented in the fossil but more ventrally oriented in *Plestiodon* (see also fig. 4 in Nash 1970). Based on differences from other NA lizards, the fossil coronoid is referable to Anguimorpha. *Xenosaurus*, except for *X. rackhami*, differs from the fossil in having a foramen on the anterolateral process (Evans 2008; Bhullar 2011). *Heloderma* differs in having a dorsoventrally expanded anteromedial process (Evans 2008). *Anniella* differs in having a facet for the coronoid process of the dentary extending on the anterior face of the coronoid process (Evans 2008). The fossil is assigned to Anguidae. The coronoid process of examined gerrhonotines and diploglossines is generally taller and more distinct than that of the fossil and of some *Ophisaurus*. On that basis, the fossil was identified to Anguinae.

### Compound bone

#### Description

TxVP 41229-29001 is a left compound bone missing the anterior portion of the prearticular and posterior tip of the retroarticular process (Fig. 32). The adductor fossa is narrow. The retroarticular process is broadened and medially oriented. The dorsal surface of the retroarticular process is depressed. The surangular is short and the dorsal margin is rounded. There is a distinct squared-off tubercle anterior to the articular surface. There is a ventral angular articulation facet and an anterolateral coronoid facet. There is a foramen on the surangular just posterior to the adductor fossa. On the lateral surface there are two anterior surangular foramina and one posterior surangular foramen. There is also a dorsal foramen just anterior to the medial process.

#### Identification

The fossil compound bone shares with anguimorphs, geckos, and scincids a medially directed and broadened retroarticular process (Estes et al. 1988). Geckos differ from the fossil in having a distinct notch on the medial margin of the retroarticular process (Estes et al. 1988). Scincids differ in having a tubercle or flange on the medial margin of the retroarticular process (Estes et al. 1988), but this feature was not obvious in all examined specimens, particularly in *Scincella*. Examined scincids differ in having a narrower and taller tubercle anterior to the articular surface and in having an adductor fossa that extends farther posteriorly. Examined *Scincella* differ in having a comparatively elongated and more rectangular retroarticular process. Based on differences from other NA lizards, the fossil is referable to Anguimorpha.

Among anguimorphs, *Xenosaurus* differs in having a deep subcoronoid fossa (Bhullar 2011). Examined *Anniella* differ in having a more retroarticular process that slants medially, and *Heloderma* have a more slender retroarticular process (Evans 2008). The fossil is referable to Anguidae. Among anguids, the fossil differs from many examined gerrhonotines and diploglossines in having a very short adductor fossa with a distinctly separate posteromedial surangular foramen. Some examined gerrhonotines also have a short adductor fossa with a distinctly separate posteromedial surangular foramen but differ in having a more mediolaterally expanded retroarticular process. On this basis, the fossil is referred to Anguinae.

**Gerrhonotinae Tihen, 1949**

**Referred specimens**: See Table S3.

### Dentary

#### Description

TxVP 41229-27659 is a right dentary with 26 tooth positions (Fig. 29). Teeth bear medial striations. Mesial teeth are unicuspid, pointed, and slightly recurved while distal teeth are near-bicuspid and more squared-off. The coronoid process is well-developed and pointed, and the angular process is present but broken posteriorly. There is no surangular process. The Meckelian canal is open ventrally. The intramandibular septum has a free posteroventral margin. The dental shelf is broad and is slanted ventromedially. There is a posteriorly projecting splenial spine. There are six nutrient foramina arranged in a row on the lateral surface.

#### Identification

The fossil is assigned to Anguimorpha based on the presence of a posteriorly projecting splenial spine (Conrad 2008; Čerňanský and Augé 2019). Dentaries are assigned to Anguidae based on the presence of an intramandibular septum with a free posteroventral margin (Gauthier 1982; Klembara et al. 2014; Scarpetta 2018; Scarpetta et al. 2021). Diploglossines differ from the fossils in lacking a splenial spine and in having the anterior inferior alveolar foramen contained within the splenial (Syromyatnikova and Aranda 2022). *Ophisaurus* differs from the fossil in having a long surangular process and having a surangular spine (Klembara et al. 2014; Scarpetta et al. 2021). The fossil is assigned to Gerrhonotinae.

**Clade composed of (*Gerrhonotus*, (*Barisia*, *Abronia*))**

**Referred specimens**: See Table S3.

### Premaxilla

#### Description

TxVP 41229-27657 is a premaxilla with ten tooth positions (Fig. 29). Teeth are unicuspid. The anterior rostral surface is rounded, and the nasal process is curved posteriorly and broken at the posterior end. The nasal process is wide and has lateral notches near the base. The nasal process tapers posteriorly and has a well-developed posterior keel. On the alveolar plate, there are lateral maxillary facets and dorsal ossifications that form an ossified bridge with the nasal process. Posteriorly, the palatal plate is slightly broken and has a wavy posterior edge. The incisive process is large, round, and slightly bilobed. There are large foramina posterolateral to the base of the nasal process and an anterior foramen between the ossified bridge, the nasal process, and the alveolar plate on either side.

#### Identification

The fossil premaxilla is assigned to anguimorpha based on having a large, round, and bilobed incisive process (Evans 2008). The incisive process is comparatively smaller and less distinctly bilobed in pleurodontans, scincids (when present), and xantusiids (Evans 2008). Among NA anguimorphs, the fossil shares with anguids and *Xenosaurus* nine or greater tooth positions (Conrad et al. 2011; Bhullar 2011; Scarpetta 2018). The fossil differs from *Xenosaurus* in lacking a rugose rostral surface of the premaxilla (Bhullar 2011). Anguinae and Diploglossinae differ from the fossil in having a forked palatal process (Evans 2008; Conrad et al. 2011; Scarpetta 2018). The fossil is assigned to Gerrhonotinae.

The fossil is identifiable to the clade (*Desertum*, (*Gerrhonotus*, (*Barisia*, *Abronia*))) based on the presence of an ossified bridge between the nasal process and the body of the premaxilla (Scarpetta et al. 2021). An ossified bridge also occurs in some species of *Elgaria*, but the bridge is much thinner (Ledesma et al. 2021). Furthermore, most *Elgaria* have a midline anterior foramen, which is absent on the fossil (Scarpetta et al. 2021). Examined *Desertum lugoi* differ in having a thinner nasal process that sometimes narrows towards the base (Ledesma et al. 2021). On that basis, the fossil is assigned to the clade (*Gerrhonotus*, (*Barisia*, *Abronia*)).

### Jugal

#### Description

TxVP 41229-26858 is a right jugal that is missing much of the suborbital process (Fig. 29). There is a distinct ventral maxillary facet on the suborbital process. The quadratojugal process (jugal spur) is long and pointed. The postorbital process is long and is directed dorsally and has a postorbital facet on the anteromedial surface. On the medial surface, there is a medial ridge that is located at the midline of the suborbital and postorbital processes. The medial ridge defines the anterior border of a small depression on the medial surface anterior to the quadratojugal process. There are two medial and four lateral foramina near the inflection point.

#### Identification

TxVP 41229-26858 is assigned to Anguimorpha based on the presence of a medial ridge located at the midline of the suborbital and postorbital processes (Čerňanský et al. 2014). The fossil differs from NA pleurodontans in having a quadratojugal process and a dorsally directed postorbital process (Gauthier et al. 1988; Smith 2009a; Fig. 42). Geckos have a highly reduced jugal, and dibamids lack a jugal (Greer 1985; Evans 2008). Examined scincids differ in having a long and thin suborbital process with a distinct notch on the ventral margin and a large depression (coronoid recess of Smith 2011) on the medial surface near the inflection point. Teiioids differ in having a distinct medial ectopterygoid process, which is also present, albeit smaller, in some gymnophthalmoids (Evans 2008). Xantusiids differ in having a short suborbital process, and *Xantusia riversiana* and *Lepidophyma* have an anteroposteriorly widened postorbital process (Savage 1963).

**Figure 42.**
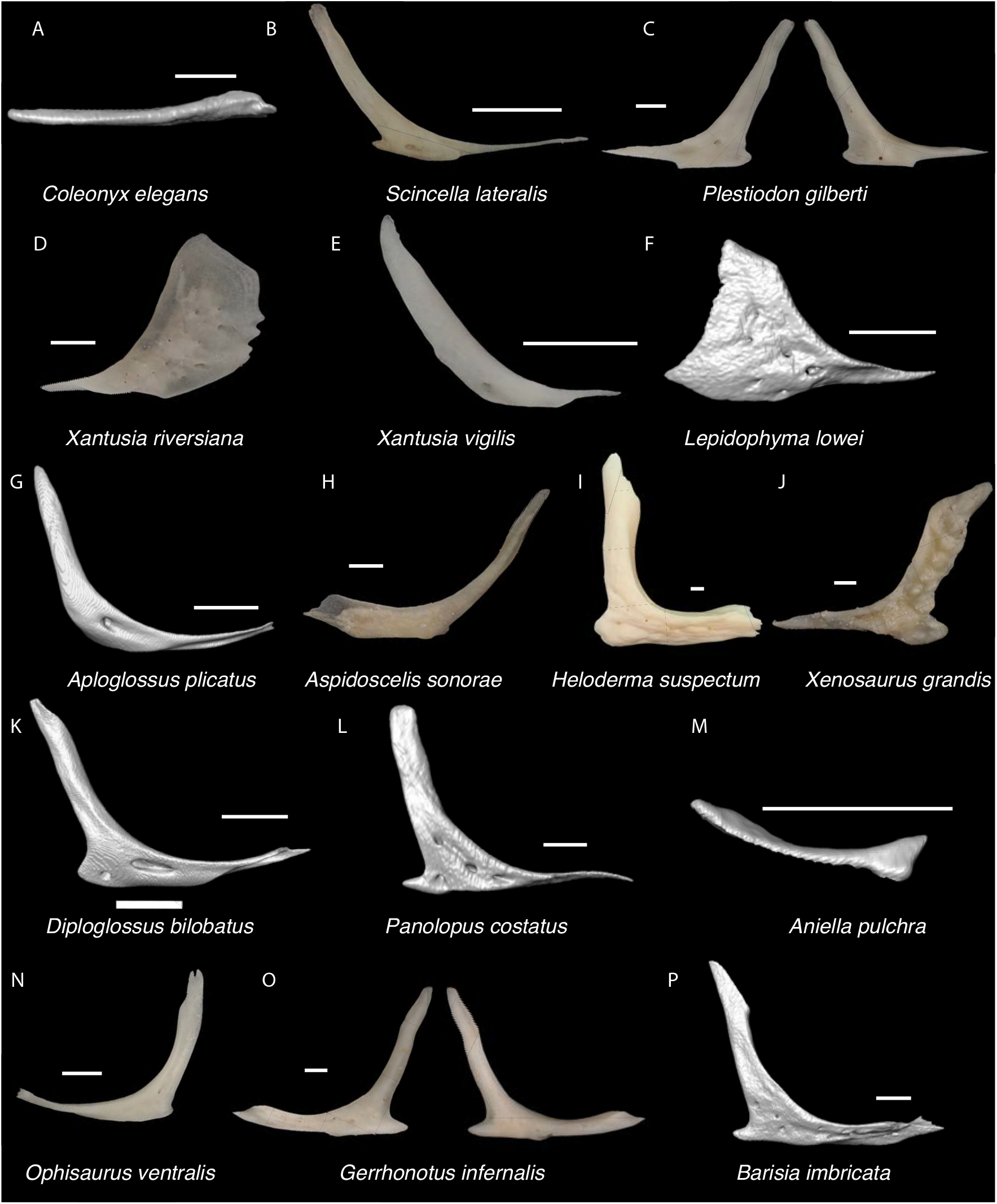
Non-Pleurodontan Jugals. Jugals in lateral view– **A**. *Coleonyx elegans* UF 11258; **B**. *Scincella lateralis* TxVP M-4489; **C**. *Plestiodon gilberti* TxVP M-8587; **D**. *Xantusia riversiana* TxVP M-8505; **E**. *Xantusia virigatus* TxVP M-12130; **F**. *Lepidophyma lowei* LACM 143367; **G**. *Aploglossus plicatus* TNHC 34481; **H**. *Aspidoscelis sonorae* TxVP M-15670; **I**. *Heloderma suspectum* TxVP M-8593; **J**. *Xenosaurus grandis* TxVP M-8960; **K**. *Diploglossus bilobatus* TNHC 31933; **L**. *Panolopus costatus* UF 59382; **M**. *Anniella pulchra* FMNH 130479; **N**. *Ophisaurus ventralis* TxVP M-8585; **O**. *Gerrhonotus infernalis* TxVP M-13441; **P**. *Barisia imbricata* TNHC 76984. Scale bars = 1 mm.

Among NA anguimorphs, *Anniella* differs in having a reduced jugal (Toerien 1950) and *Heloderma* differs in that the anterior and posterior processes form a right angle (Evans 2008). *Xenosaurus* has co-ossified osteoderms or sculpturing on the lateral surface of the jugal (Bhullar 2011). The fossil was assigned to Anguidae. A quadratojugal process is usually present in anguids (Evans 2008; Klembara et al. 2017; Scarpetta 2021). The length of the quadratojugal process in the fossil is longer than that of examined *Ophisaurus* but is similar to that present in some species of *Gerrhonotus*, *Barisia*, and *Abronia* (Klembara et al. 2017; Ledesma et al. 2021; Scarpetta et al. 2021). An examined *Celestus enneagrammus* (FMHN 108860) and *Panolopus costatus* (UF 59382) also have a long quadratojugal process but differ from the fossil in having a relatively shorter orbital process. Examined *Desertum lugoi* have a relatively shorter quadratojugal process. Jugals are assigned to the clade (*Gerrhonotus*, (*Barisia*, *Abronia*)) on this basis. *Barisia* differs in having sculpturing on the lateral surface of the jugal (Scarpetta et al. 2021) so the fossil likely represents either *Gerrhonotus* or *Abronia*.

***Gerrhonotus* Wiegmann, 1828**

**Referred specimens**: See Table S3.

### Maxilla

#### Description

TxVP 41229-27438 is a right maxilla (Fig. 29). There are 22 tooth positions. The teeth are unicuspid with medial striations on the crowns, and most are recurved. The lateral surface of the maxilla is highly sculptured. The facial process is tall and broad, and gently curves dorsomedially. The medial margin of the facial process has a distinct nasal facet, and the posterior margin has a large notch. The premaxillary process is long and bifurcated with a longer lateral projection and a shorter medial lappet. The crista transversalis is low and trends anteromedially from the facial process, defining the medial border of a shallow depression on the premaxillary process. There is a narrow palatal shelf without a distinctly projecting palatine process. There is a deep, elongate medial recess on the medial surface of the facial process. The lateral wall of the posterior orbital process is short. The dorsal surface of the postorbital process has an elongate, deep jugal groove and a small ectopterygoid facet at the posterior end. There is an opening for the superior alveolar canal anterior to the facial process. There are three large superior alveolar foramina (maxillary trigeminal foramina of Evans 2008) on the palatal shelf medial to the palatine process and eight lateral nutrient foramina.

#### Identification

Fossil maxillae share with some scincids and anguimorphs unicuspid teeth with striated crowns (Estes 1963; Smith 2009b). Fossils differ from examined scincids in lacking a large notch at the end of the posterior orbital process (see also fig. 5 in Čerňanský and Syromyatnikova 2021). Furthermore, *Scincella* differs in lacking striations on the crowns (Townsend et al. 1999) and examined *Plestiodon* differ in that the crista transversalis abruptly trends medially anterior to the facial process and the palatal shelf is wider. Based on these differences, the fossils are referable to Anguimorpha.

*Anniella* and *Heloderma* differ from the fossils in having a greater number of pointed and recurved teeth as well as fewer tooth positions (up to seven in *Anniella* and up to ten in *Heloderma*; Torien 1950; Evans 2008). *Xenosaurus* differs in having fused osteoderms laterally on the facial process (Bhullar 2011). Fossils are assigned to Anguidae. The fossil is assigned to Gerrhonotinae and differs from anguines and diploglossines in having an elongated premaxillary process that lacks a deep anterior notch (Meszoely 1970). Some gerrhonotines have a notch where the lacrimal articulates with the posterior edge of the facial process; however, a large notch as seen on the fossil only occurs only in *Gerrhonotus infernalis* among examined gerrhonotines (Ledesma et al. 2021) and is absent in other examined anguids.

**Scincidae Gray, 1825**

**Referred specimens**: See Table S3.

### Compound bone

#### Description

TxVP 41229-26992 is a right compound bone missing only the anterior portion of the prearticular (Fig. 43). The adductor fossa is anteroposteriorly long. The retroarticular process is narrow and elongated posteriorly. There is a medial ridge that extends from the posterior margin of the articular surface to the medial edge of the retroarticular process, which bears a small medial tubercle. The dorsal and ventral surfaces of the retroarticular process are characterized by slight depressions. There is a tall, rounded tubercle anterior to the articular surface. The anterodorsal margin of the surangular has a dorsally expanded crest for articulation with the coronoid. There is a ventral angular articulation facet and a “v” shaped facet on the anterolateral surface of the surangular for articulation with the dentary. There are both anterior and posterior surangular foramina. There is also a dorsal foramen just anterior to the dorsal tubercle and a foramen posterior to the articular surface. TxVP 41229-26998 does not differ substantially.

**Figure 43.**
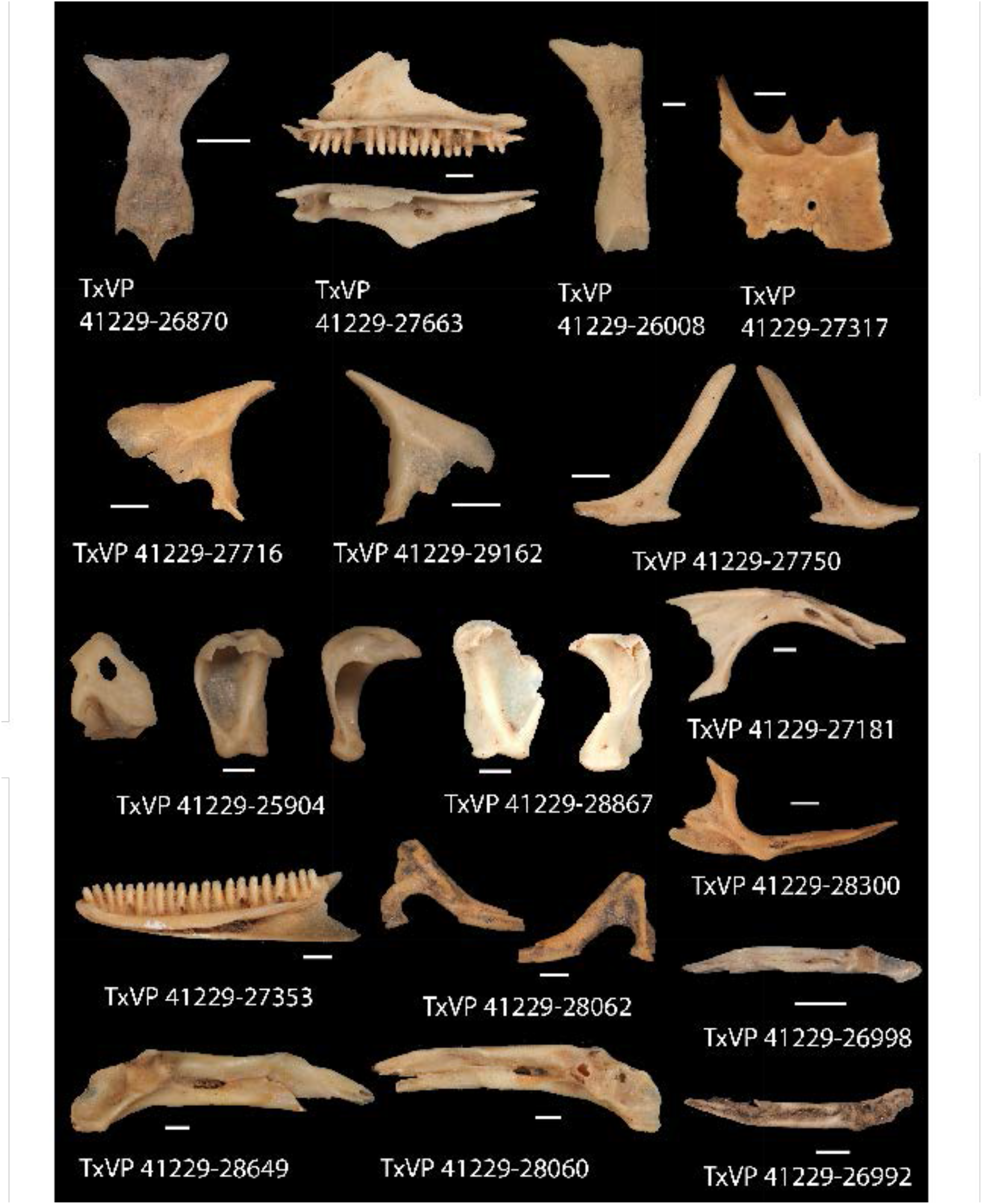
Fossil scincids. TxVP 41229-26870 Dorsal view of frontal; TxVP 41229-27663 Medial and dorsal view of right maxilla; TxVP 41229-26008 Dorsal of frontal; TxVP 41229-27317 Dorsal view of parietal; TxVP 41229-27716 Lateral view of left prefrontal; TxVP 41229-29162 Lateral view of right prefrontal; TxVP 41229-27750 Lateral and medial view of left jugal; TxVP 41229-25904 Dorsal, posterior, and lateral view of left quadrate; TxVP 41229-28867 Posterior view of right quadrate; TxVP 41229-27181 Dorsal view of left pterygoid; TxVP 41229-28300 Dorsal view of right pterygoid; TxVP 41229-27353 Medial view of right dentary; TxVP 41229-28062 Lateral and medial view of right coronoid; TxVP 41229-26998 Dorsal view of right compound bone; TxVP 41229-28649 Dorsal view of left compound bone; TxVP 41229-28060 Dorsal view of right compound bone; TxVP 41229-26992 Dorsal view of left compound bone. Scale bars = 1 mm.

#### Identification

Fossils share with scincids, anguimorphs, and geckos a medially directed retroarticular process (Estes et al. 1988). Fossils differ from geckos in lacking a notch on the medial margin of the retroarticular process (Estes et al. 1988). Fossils differ from anguimorphs and geckos in having a medial tubercle on the retroarticular process, although a small tubercle was found in a few anguimorph taxa (e.g., *Diploglossus millepunctatus* TxVP M-9010). Fossils differ from anguimorphs in having a “v” shaped facet on the anterolateral surface of the surangular (see also fig. 10E of Estes et al. 1988) which is present on all examined skinks except specimens of *Plestiodon gilberti*. Fossils also differ from anguimorphs in having a narrower and more rounded to pointed tubercle anterior to the articular surface. Examined anguimorphs tend to have a shorter and more squared-off tubercle. Fossils were assigned to Scincidae on this basis. Fossils share with examined *Scincella*, mabuyids (Čerňanský and Syromyatnikova 2021), and *Plestiodon tetragrammus* a relatively narrow retroarticular process. We refrain from making a more refined identification pending examination of additional skeletal material.

**Sphenomorphinae Welch, 1982**

***Scincella* Mittleman, 1950**

**Referred specimens**: See Table S3.

### Frontal

#### Description

TxVP 41229-26870 is a frontal (Fig. 43). Sculpturing is restricted to the posterodorsal surface of the bone. There are anterolateral facets for the prefrontal, and the anterior face of the frontal is defined by two short dorsal nasal facets formed by two anterolateral processes and an anteromedial process. The interorbital margins are waisted, and the posterolateral processes extend laterally. The posterior edge is slightly wavy and has interlacing articulation facets for the parietal. There are small postfrontal facets laterally on the posterolateral processes. The cristae cranii are low and diverge posteriorly along the lateral margins of the ventral surface. There is a broad ridge on the ventral surface near the interorbital margin creating separate anterior and posterior depressions.

#### Identification

The fossil frontal shares with Pleurodonta, Teiioidea, Mabuyinae and *Scincella* a fused frontal with reduced cristae cranii (Greer 1970; Estes et al. 1988). Examined teiids differ in having a frontal that is relatively more elongate relative to the width and gymnophthalmoids differ in having frontal tabs (Estes et al. 1988). Examined NA pleurodontans differ from the fossil in lacking a distinct anteromedial process that extends far anterior relative to the anterolateral processes. Furthermore, among examined pleurodontans, the cristae cranii converge to a greater degree compared to the fossil. The fossil can be attributed to Scincidae. Examined *Mabuya* differ from *Scincella* in having more dermal sculpting on the dorsal surface and a relatively wider anterior portion. The fossil is referred to *Scincella*.

**Scincinae Gray, 1825**

**Referred specimens**: See Table S3.

### Maxilla

#### Description

TxVP 41229-27663 serves as the basis for our description (Fig. 43). TxVP 41229-27663 is a right maxilla with 19 tooth positions. Teeth are unicuspid with medial striations. The facial process is tall, broad and gently curves dorsomedially. The medial margin of the facial process has a distinct nasal facet, and the anterior margin has a small projection. The premaxillary process is short and bifurcated with a longer lateral projection and a shorter medial lappet. The crista transversalis is low in height and trends medially from the facial process and defines the posterior border of a shallow depression on the premaxillary process. There is a broad palatal shelf with a triangular palatine process. The medial surface of the facial process bears a distinct ridge that extends anteriorly onto the palatal shelf and defines the posterior border for a deep medial recess on the anteromedial surface of the facial process. The lateral wall of the posterior orbital process is tall, and the posterior end of the process is slightly bifurcated. The dorsal surface of the postorbital process has an elongate, shallow jugal groove. There is an anterior inferior alveolar foramen anterior to the facial process. There are two superior alveolar foramina on the palatal shelf medial to the palatine process, 13 lateral nutrient foramina dorsal to the tooth row, and two foramina anterodorsally on the facial process.

#### Identification

Fossil maxillae share with some scincids and anguimorphs unicuspid teeth with striated crowns (Estes 1963; Smith 2009b). Fossils differ from examined anguimorphs, except some *Diploglossus* (Bochaton et al. 2016b; Syromyatnikova and Aranda 2022), in having a large notch at the end of the posterior orbital process. Examined anguids differ from the fossils in having a comparatively shorter lateral lamina along the posterior orbital process and a crista transversalis that trends anteromedially instead of medially from the facial process (Smith 2009b). Fossils can be further differentiated from the NA anguimorphs *Anniella* (6–7 tooth positions) and *Heloderma* (6–10 tooth positions) in having more tooth positions (Riepell 1980; Evans 2008; this study) lacking sharp recurved teeth (Evans 2008). *Anniella* lack a crista transversalis (Smith 2009b) and *Xenosaurus* have fused osteoderms on the lateral surface of the facial process (Bhullar 2011). Fossils are assigned to Scincidae.

Examined *Scincella* and at least some mabuyine skinks (Čerňanský and Syromyatnikova 2021) differ from the fossils in lacking striations on the crowns (Townsend et al. 1999), having a crista transversalis that trends anteromedially from the facial process, and having a much narrower palatal shelf posterior to the palatal process. Fossils are assigned to Scincinae. We refrain from making a more refined identification pending examination of additional skeletal material of scincine lizards.

### Frontal

#### Description

TxVP 41229-26008 serves as the basis for our description and is an unfused right frontal (Fig. 43). There is a small area of co-ossified osteoderms on the dorsal surface. There is an anterolateral prefrontal facet and an anterodorsal nasal facet. The anterior face of the bone is flat and has a well-developed anterolateral process. The interorbital margin is only slightly narrower than the anterior end. The posterolateral process gently curves laterally and projects posterolaterally past the posteromedial edge. The posterior edge has a shallow parietal articulation facet. There is a postfrontal facet along the posterolateral edge. The ventral portion of the crista cranii is broken, but the crest is well-developed, tall, and anteroposteriorly long. Ventrally, there is a series of small transverse ridges and corresponding grooves medial to the crista cranii.

#### Identification

The fossil shares with some anguimorphs and scincids co-ossified osteoderms with dorsal sculpturing (Estes et al. 1988), well-developed and ventrally directed cristae cranii (=subolfactory processes of McDowell and Bogert 1954), and an unfused frontal (Estes et al. 1988). Osteoderm sculpturing has a more pitted texture in many anguimorphs (Conrad 2008). Examined *Ophisaurus* and other anguines, except for *Anguis fragilis* (Klembara et al. 2017), also differs from the fossil in having a pointed anterior portion of the unfused frontal. Fossils are assigned to Scincidae.

*Scincella* and mabuyine skinks are excluded because they both have a fused frontal (Greer 1970). In one examined extant specimen, *Plestiodon skilitonianus* TxVP M-8498, the frontals are separate but are held together by fused osteoderms. Fossils are assigned to Scincinae.

### Parietal

#### Description

The exemplar TxVP 41229-27317 is missing the anterolateral corner and ends of the postparietal processes (Fig. 43). The parietal table is rectangular and smooth with scattered small foramina. The anterior edge is straight with small frontal tabs. The posterior edge between the postparietal processes has two small nuchal fossae separated by a small ridge, as well as two long posterior projections that create a medial notch. The right postparietal process is broad and flat at its base. There is a flat muscle attachment surface lateral to the ventrolateral crests. The ventrolateral crests are low and are broken at the ends. There is a large ventral swelling at the pit for the processus ascendens. The posterior projections and a ventrolateral ridge on the postparietal process define small ventral depressions. There is a large parietal foramen within the middle of the parietal table.

#### Identification

Parietals are assigned to Scincomorpha based on the presence of long posterior projections (median extensions of Evans 2008) on the posterior edge of the parietal table between the postparietal processes (Evans 2008; Gauthier et al. 2012). Xantusiids, except for *Cricosaura*, some *Lepidophyma* (e.g., *L. gaigeae*), and *Xantusia riversiana*, differ in having a paired parietal (Savage 1963; Maisano 2002b; Evans 2008). *Cricosaura* and *Xantusia riversiana* differ in lacking long postparietal processes (Evans 2008). Posterior projections on the posterior edge of the parietal table are found in some anguid lizards (see fig. 9 in Ledesma et al. 2021), but the projections in anguids are much smaller than those found on the fossils and scincids. Fossils are assigned to Scincidae. Examined *Scincella* and mabuyine skinks have less developed adductor crests compared to that in the fossils (Jerez et al. 2015) and lack long, thin, ventral projections, the base of which is preserved in fossils. Fossils are assigned to Scincinae.

### Prefrontal

#### Description

TxVP 41229-27716 is a left prefrontal (Fig. 43). It is triradiate with a short orbital process, a short ventral process, and an elongate anterior facial sheet. The anterior facial sheet has a broad articulation facet for the facial process of the maxilla, which extends dorsally on the prefrontal. There is a small ridge on the lateral surface near the base of the orbital process. The ventral process is pointed, projects posteriorly, and has a small lateral ridge. There is a distinct notch for the lacrimal foramen, with the ventral process forming the posterior border. Medially, there is a smooth, rounded, and concave wall of the olfactory chamber. Dorsal to the olfactory chamber and medially on the orbital process is a deep groove for articulation with the frontal. The orbitonasal flange is broad and slightly broken the medial margin. There is a small foramen anterodorsal to the notch for the lacrimal foramen. TxVP 41229-29162 differs from TxVP 41229-27716 in having a small overhanging lamina of bone dorsal to the notch for the lacrimal foramen and in having a notch in the medial margin of the orbitonasal flange.

#### Identification

The fossil differs from NA pleurodontans in lacking a lateral prefrontal boss (Estes et al. 1988; Smith 2009b; Smith 2011; Gauthier et al. 2012), lacking a strong lateral canthal ridge (reported in *Anolis* and *Polychrus*; Smith 2011), lacking a supraorbital spine (present in *Phrynosoma* and *Corytophanes*; Smith 2009a), and lacking a thin, crescent shaped prefrontal with a distinct laterally projecting lamina (present in examined phrynosomatines). NA teiids and gymnophthalmoids differ from the fossil in having a laterally projecting lamina (lacrimal flange of Bell et al. 2003) with a distinct articulation facet for the facial process of the maxilla (Tedesco et al. 1999). Xantusiids are excluded because the lacrimal is fused to the prefrontal (Savage 1963), the suborbital foramen is nearly or entirely enclosed within the prefrontal, and there is a vertical articulation ridge (most *Xantusia*) or flange (e.g., *X. riversiana*) that articulates with the maxilla. Examined *Coleonyx* differ in having an orbitonasal flange that extends farther medially compared to the fossil. *Sphaerodactylus roosevelti* has a smaller notch for the lacrimal foramen (Daza et al. 2008). Examined anguimorphs differ in lacking an oblong, elongate anterior process. Based on differences with other NA lizards, the fossil is identified to Scincidae. The prefrontal of examined *Scincella* lacks a posterior procession on the ventral process, has a less elongate anterior process, and is much smaller than the fossil. The fossil is assigned to Scincinae. The fossil, *Plestiodon gilberti*, and *P. multivarigatus* share a relatively short posterodorsal process, but the process is long in examined *P. obsoletus*.

### Jugal

#### Description

TxVP 41229-27750 is a right jugal (Fig. 43). There is a distinct maxillary facet on the lateral and ventral surface of the suborbital process. The suborbital process thins anteriorly. There is a large, pointed quadratojugal process. The postorbital process is long, posterodorsally oriented, and has a shallow postorbital facet on the anteromedial surface. There is a medial ridge that is located on the midline of the suborbital process and anteriorly on the postorbital process. The medial ridge defines the anterior border for a small depression on the medial surface at the base of the quadratojugal process and the base of the postorbital process. There are two medial and two lateral foramina at the inflection point, as well as one medial foramen on the postorbital process. TxVP 41229-29020 differs from TxVP 41229-27750 in having a round medial boss near the inflection point and in only having one medial foramen near the inflection point.

#### Identification

Jugals share with other NA lizards except for dibamids, geckos, and some pleurodontans an angulated jugal (Conrad 2008). Fossils differ from NA pleurodontans in having a quadratojugal process with a dorsally oriented postorbital process (Gauthier et al. 1988; Smith 2009a). Teiioids differ in having a distinct medial ectopterygoid process, which is also sometimes present, albeit comparatively smaller, in gymnophthalmoids (Evans 2008). Examined xantusiids differ in lacking a quadratojugal process and in having a short suborbital process (Evans 2008). *Xantusia riversiana* and *Lepidophyma* have an anteroposteriorly widened postorbital process (Savage 1963). Among NA anguimorphs, *Anniella* differs in having a reduced jugal (Toerien 1950) and *Heloderma* differs in having the postorbital and suborbital processes form a right angle (Evans 2008). *Xenosaurus* has co-ossified osteoderms or sculpturing on the lateral surface of the jugal (Bhullar 2011). The fossil differs from anguids in having a medial ridge located midline on the suborbital process and anteriorly on the postorbital process (Čerňanský et al. 2014). Furthermore, fossils differ from examined anguids in having a distinct notch on the ventral margin of the suborbital process. Examined *Scincella* differ in having a jugal that is more smoothly curved with a relatively thinner suborbital process. Fossils are assigned to Scincinae.

### Quadrate

#### Description

TxVP 41229-25904 is a left quadrate (Fig. 43). The central column narrows and curves laterally at its base. There is a small pterygoid lappet and a well-developed, anteromedially directed medial crest. The conch is deep and gradually slants laterally from the central column. The cephalic condyle projects far posteriorly and there is extensive ossification dorsally enclosing the squamosal foramen, along with an anteriorly expanded dorsal tuber (*sensu* Klembara et al. 2017). There is a foramen medial to the central column and three foramina on the anterior surface. TxVP 41229-28867 differs in having a large pterygoid lappet and a foramen on the posterior surface dorsal to the mandibular condyle.

#### Identification

The fossil shares a distinct pterygoid lappet with scincids, teiids, gymnophthalmoids, some pleurodontans, *Heloderma*, *Xenosaurus*, and some gerrhonotines (Estes et al. 1988; Evans 2008; Ledesma et al. 2021). Most examined NA pleurodontans differ from the fossil in having a quadrate that is much wider dorsally compared to the articular surface. In *Anolis* and *Uma*, the lateral margins are parallel; however, examined *Anolis* differ in having a distinct boss at the ventromedial margin of the quadrate, and *Uma* differ in having a laterally slanted cephalic condyle and central column. *Xenosaurus* differ in having a more shallow lateral conch (Bhullar 2011) and examined *Heloderma* differ in having a dorsal head with more distinct anterior projection. The central column in the fossils is curved posteriorly to a greater degree than that of examined gerrhonotines. Examined teiids and gymnophthalmoids differ in having a more curved tympanic crest. Fossils were assigned to Scincidae on this basis. Examined *Scincella* differ in having a relatively narrow quadrate in anterior and posterior view. Fossils are assigned to Scincinae.

### Pterygoid

#### Description

TxVP 41229-28300 is a right pterygoid (Fig. 43). The palatine process is broad with an anterior palatine facet. The transverse process has a distinct posterolateral corner and a triangular and anterolaterally pointed distal end. The transverse process has a shallow ectopterygoid facet. Ventrally, the transverse process has a large ridge that slightly curls anteriorly for insertion of the pterygomandibular muscle, but there is no dorsal ridge on the transverse process. On the palatal plate there is a dorsal ridge that trends anteriorly and forms the lateral border for a medial depression. There is a deep fossa columella and an epipterygoid crest anteromedial to the fossa columella. There is no postepipterygoid groove (*sensu* Good 1987) on the quadrate process. The quadrate process is elongated, and the medial surface has a shallow groove that serves for insertion of the pterygoideus muscle. The ventrolateral surface of the quadrate process bears a distinct groove. The medial margin of the pterygoid, including the quadrate process and the palatal plate, is curved laterally. There is no medial projection at the floor of the basipterygoid fossa and there is a small patch of seven pterygoid tooth positions containing four teeth. There is a dorsal foramen on the palatal plate. TxVP 41229-27181 is missing the dorsal ridge and the epipterygoid crest, and TxVP 41229-28682 has an epipterygoid crest that does not project as far medially as in TxVP 41229-28300.

#### Identification

Fossils share the presence of pterygoid teeth with scincids, teiids, some NA gymnophthalmoids (e.g., *Gymnophthalmus* and *Ptychoglossus*), some pleurodontans, and some anguimorphs (Estes et al. 1988; Evans 2008). Examined NA pleurodontans, except *Petrosaurus*, differ in having the transverse process oriented more medially. *Petrosaurus* also lack pterygoid teeth (Etheridge 1964). Examined NA teiids differ in having an extensive, medially directed lamina of bone on the quadrate process (Du Bois 1953). *Ptychoglossus* differs in having a posteromedial process along the medial edge of the palatal plate (Morales et al. 2019) and *Gymnophthalmus* differs in having a more slender palatal plate (Evans 2008). Examined anguimorphs differ in having a postepipterygoid groove on the quadrate process (Evans 2008). *Xenosaurus* and *Anniella* can be further differentiated in lacking pterygoid teeth (Evans 2008) and *Heloderma* differs in having a single row of pterygoid teeth located on an elevated ridge (Evans 2008). The medial edge of the pterygoid is more strongly curved in fossils compared to examined anguids. Examined *Scincella* and mabuyines differ in lacking pterygoid teeth (Mausfeld et al. 2002). Fossils were assigned to Scincinae on this basis.

### Dentary

#### Description

TxVP 41229-27353 is a right dentary with 22 tooth positions and serves as the basis for our description (Fig. 43). Teeth are unicuspid with medial striations. There is a pointed coronoid process, no surangular process, and an angular process that is broken posteriorly. The Meckelian groove is open ventrally, and the suprameckelian lip is tall. The intramandibular septum does not extend to the distal teeth. There is a deep dental gutter. There are four nutrient foramina arranged in a row on the lateral surface of the bone.

#### Identification

Fossils share with Scincomorpha and Anguimorpha unicuspid teeth with striated crowns (Estes 1963; Smith 2009b). Anguimorphs differ from the fossils in having a posteriorly extended intramandibular septum near the posterior end of tooth row (Pregill 1981; Estes et al. 1988). Xantusiids differ in having a fused spleniodentary (Savage 1963; Mead and Bell 2001; Evans 2008). Fossils were assigned to Scincidae on this basis. Mabuyines have a closed or fused Meckelian groove (Greer 1974; Čerňanský and Syromyatnikova 2021). *Scincella* differs in lacking striations on the crowns and in having a closed but unfused Meckelian groove (Townsend et al. 1999). Fossils were assigned to Scincinae on this basis.

### Coronoid

#### Description

TxVP 41229-28062 is a right coronoid and serves as the basis for our description (Fig. 43). The coronoid process is tall, rounded and slopes anteroventrally. The anteromedial process is elongated with a medial splenial facet. The posteromedial process is oriented ventrally and has a small posterior notch. There is a medial crest that extends from the coronoid process and extends onto the posteromedial process. There is a large, anteroventrally projecting lateral process with a dorsal dentary articulation facet. There is a anteroventrally oriented lateral crest that extends along the anterolateral process. There is a small articulation surface for the surangular, located ventral to the anterolateral process.

#### Identification

Fossil coronoids differ from xantusiids and some pleurodontans in having an anterolateral process (Estes et al. 1988). The fossil differs from examined NA pleurodontans with an anterolateral process (see Crotaphytidae section above) in having a lower and more rounded coronoid process. The fossil differs from examined *Aspidoscelis* and *Ameiva* in lacking a deeply notched posterior edge forming dorsal and ventral rami (Tedesco et al. 1999). The fossil differs from examined gymnophthalmoids in having the posteromedial process oriented ventrally (Evans 2008; Roscito and Rodrigues 2010). Examined geckos differ from the fossils in having an anterolateral process that is not directed as far ventrally (Kluge 1962; Daza et al. 2008; Bochaton et al. 2018). Examined NA anguimorphs also differ in having an anterolateral process that does not project as far ventrally as in the fossils. Fossils were assigned to Scincidae. Examined *Scincella* have a more posteriorly oriented posteromedial process and examined mabuyines have a taller and more pointed coronoid process (Čerňanský and Syromyatnikova 2021). Fossils were assigned to Scincinae.

### Compound bone

#### Description

TxVP 41229-28649 is a left compound bone that is missing the anterior portion of the prearticular (Fig. 43). The adductor fossa is short. The retroarticular process is broad and medially oriented. There is a medial ridge that extends from the posterior margin of the articular surface to a medial tubercle on the retroarticular process. The dorsal surface of the retroarticular process is a semicircular depression, and the ventral surface is smooth. There is a tall, rounded tubercle anterior to the articular surface. The anterodorsal margin of the surangular has a dorsally expanded crest where it articulates with the coronoid. There is a ventral angular articulation facet and an anterolateral coronoid facet. There are anterior and posterior surangular foramina. There is also a dorsal foramen just anterior to the medial process, and a foramen posterior to the articular surface. TxVP 41229-28060 differs in having a low ridge lateral to the medial ridge on the retroarticular process. TxVP 41229-28053 differs in lacking a medial boss and lacking an anterodorsal crest on the surangular.

#### Identification

Fossils share a medially directed and broadened retroarticular process with scincids, anguimorphs, and geckos (Estes et al. 1988). Fossils differ from geckos in lacking a notch on the medial margin of the retroarticular process (Estes et al. 1988). Fossils, except for TxVP 41229-28053, differ from anguimorphs and geckos in having a medial tubercle on the retroarticular process, although a small tubercle was found in a few anguimorph taxa as well (e.g., *Diploglossus millepunctatus* TxVP M-9010). Fossils differ from anguimorphs in having a narrower tubercle anterior to the articular surface. Examined anguimorphs tend to have a shorter and more squared-off tubercle. In examined *Scincella* and mabuyines, (Čerňanský and Syromyatnikova 2021) the retroarticular process is more narrow compared with the fossils. The retroarticular process is also narrow in examined *Plestiodon tetragrammus*. Fossils were assigned to Scincinae. Based on the shape of the retroarticular process (and given the good possibility that the fossils represent *Plestiodon*), the compound bones likely represent a species other than *Plestiodon tetragrammus*.

**Teiidae Gray, 1827**

**Referred specimens**: See Table S3.

### Premaxilla

#### Description

TxVP 41229-27594 is a premaxilla with eight tooth positions (Fig. 44). Teeth are unicuspid. The rostral surface of the premaxilla is rounded. The nasal process is slightly broken posterolaterally, but it is evident that it is strongly curved posteriorly and tapers to a point. The process is slightly widened near the midpoint, and ventrally, there is a low keel. On the alveolar plate, there are lateral maxillary facets with small lateral notches. Posteriorly, the palatal plate is strongly incised and “v” shaped. There is no incisive process. There are two bilateral ethmoidal foramina posterior to the base of the nasal process, but no anterior foramen.

**Figure 44.**
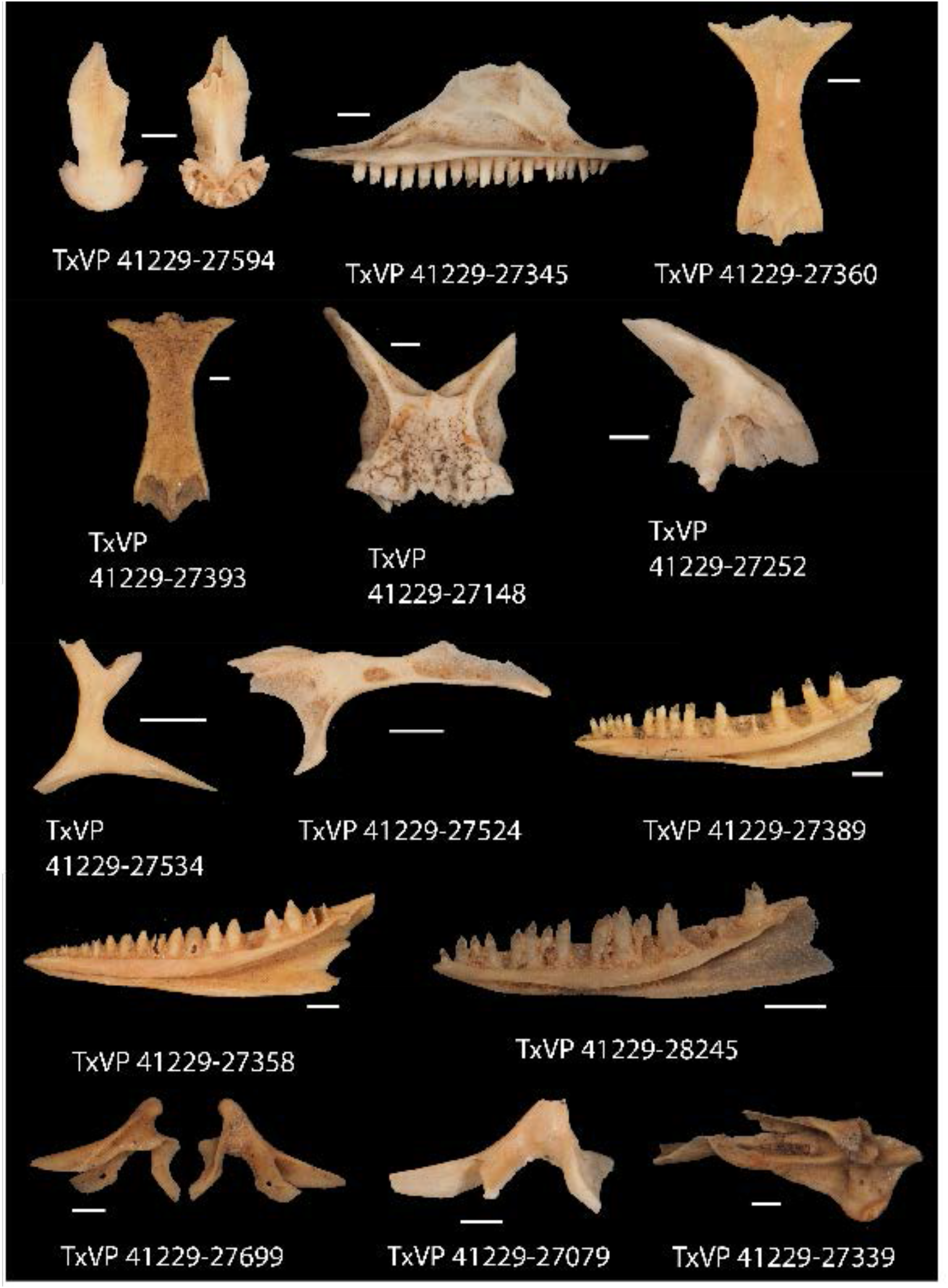
Fossil teiids. TxVP 41229-27594 Dorsal and ventral view of premaxilla; TxVP 41229-27345 Medial view of left maxilla; TxVP 41229-27360 Dorsal view of frontal; TxVP 41229-27393 Dorsal view of frontal; TxVP 41229-27148 Dorsal view of parietal; TxVP 41229-27252 Lateral view of right prefrontal; TxVP 41229-27534 Dorsal view of postorbitofrontal; TxVP 41229-27524 Dorsal view of left pterygoid; TxVP 41229-27389 Medial view of right dentary; TxVP 41229-27358 Medial view of right dentary; TxVP 41229-28245 Medial view of right dentary; TxVP 41229-27699 Lateral and medial view of left coronoid; TxVP 41229-27079 Medial view right coronoid; TxVP 41229-27339 Dorsal view of right compound bone. Scale bars = 1 mm.

#### Identification

The fossil is assigned to Teiioidea based on the absence of an incisive process (Gao and Norell 1998; Conrad 2008; Morales et al. 2019) without a large median tooth (Smith 2009b). Alopoglossids differ in having between 10 to 14 tooth positions (Morales et al. 2019). Many gymnophthalmids differ in having a laterally expanded nasal process (Evans 2008; Morales et al. 2019). *Gymnophthalmus speciosus* has parallel margins of the nasal process, but that species differs from the fossil in having a nasal process with a more rectangular posterior end (Hoyos 1998; Evans 2008). With over 300 species of gymnophthalmoid lizards, there is still much to learn about patterns of osteological variation in this group and so we tentatively assign the fossil to Teiidae.

### Maxilla

#### Description

TxVP 41229-27345 is a left maxilla that serves as the basis for our description (Fig. 44). There are 20 tooth positions. The distal-most tooth is weakly tricuspid with a minute distal cusp, and the mesial teeth, except for the most-mesial tooth, are bicuspid with a smaller mesial cusp. There are substantial deposits of cementum at the tooth bases. The facial process is tall and broad, and faces vertically, except for the anterodorsal apex, which curves dorsomedially. The dorsomedial margin of the facial process has a distinct nasal facet. The premaxillary process is long and bifurcated with a short lateral projection and a long anteriorly directed medial lappet. The crista transversalis is low and diminishes on the premaxillary process. There is a narrow palatal shelf without a distinct palatine process. The medial surface of the facial process bears a distinct ridge with an overhanging lamina that extends anteriorly onto the palatal shelf and defines the posterior border for a medial recess on the anteromedial surface of the facial process. The lateral wall of the posterior orbital process is tall, and the dorsal surface of the postorbital process has an elongate, shallow jugal groove and a small ectopterygoid facet. There is an anterior inferior alveolar foramen positioned anterior to the facial process and medial to the crista transversalis. There is one superior alveolar foramen on the palatal shelf medial to the palatine process and posterior to the facial process. The facial process is slightly concave laterally. There are seven lateral nutrient foramina dorsal to the tooth row, and several foramina scattered on the lateral surface of the facial process.

#### Identification

Maxillae are assigned to Teiioidea based on the presence of asymmetrically bicuspid distal teeth with a smaller anterior cusp that is anteriorly directed (Bell 1993; Evans 2008; Scarpetta 2020). Gymnophthalmoids differ in generally having a relatively short facial process anteroposteriorly compared to the fossils (Evans 2008). Furthermore, gymnophthalmoids lack large cementum deposits at the base of the teeth (Estes et al. 1988; Nydam et al. 2007). Maxillae are assigned to Teiidae.

### Frontal

#### Description

TxVP 41229-27360 and TxVP 41229-27397 serve as the basis for our description (Fig. 44). Anteriorly, there are lateral prefrontal facets that extend to the ventral surface, and two dorsal nasal facets defined by shorter anterolateral processes separated by a longer anteromedial process. The interorbital margins are waisted and the posterolateral processes curve laterally such that the posterior end is slightly wider than the anterior end. The posterior edge is wavy with narrow parietal facets. There are small postorbitofrontal facets laterally and ventrally on the posterolateral processes. There is a small depression on the posterodorsal portion of the frontal. The cristae cranii are low, approach one another in the interorbital region, and bound a groove for attachment of the solium supraseptale. The cristae cranii diverge posteriorly and extend along the lateral margins of the ventral surface. TxVP 41229-27397 differs in having sculpturing on the posterior half of the dorsal surface.

#### Identification

The fossils share a fused frontal with reduced cristae cranii with Pleurodonta, Teiioidea, Mabuyinae and *Scincella* (Greer 1970; Estes et al. 1988). Examined Mabuyinae (see fig. 8 in Jerez et al. 2015) and *Scincella* differ in having a frontal that is comparatively shorter relative to the width. Examined pleurodontans differ in having more strongly constricted interorbital margins (Estes et al. 1988) with a relatively wider posterior margin (Evans 2008). Juvenile teiids have more constricted interorbital margins compared to adults, but the posterior margin is not widened as in pleurodontans (Bochaton et al. 2019). Gymnophthalmoids differ in having frontal lappets on the posterior margin (Estes et al. 1988; Evans 2008). Fossils are assigned to Teiidae.

### Parietal

#### Description

TxVP 41229-27148 is a parietal with extensive sculpturing on the dorsal surface and serves as the basis for our description (Fig. 44). The anterior edge has frontal tabs. The adductor crests do not meet posteriorly, giving the parietal table a trapezoidal appearance. The ventrolateral crests are tall and are visible in dorsal view. The anterolateral processes curve laterally and have lateral facets for articulation with the postorbitofrontal. The posterior edge between the postparietal processes is characterized by two distinct depressions (nuchal fossae). The postparietal process has a dorsal crest that slants laterally. The ventral surface has a deep depression (cerebral vault). There is a deep pit for the processus ascendens along the posteroventral edge. Flanges are present at the bases of the postparietal processes.

#### Identification

Parietals share with teiioids and pleurodontans parietal lappets and a parietal foramen that is not fully enclosed by the parietal (Estes et al. 1988). Pleurodontans differ in lacking long descending parietal crests (parietal downgrowths of Estes et al. 1988). Many gymnophthalmoids differ in having the anterior margin of the parietal more distinctly trifurcate with shorter postparietal processes (Evans 2008). Furthermore, examined gymnophthalmoids have a broad parietal without the descending crests visible in dorsal view. Fossils are assigned to Teiidae.

### Prefrontal

#### Description

TxVP 41229-27252 is a right prefrontal that serves as the basis for our description (Fig. 44). It is triradiate with a long and pointed orbital process, a short ventral process, and an anterior sheet. The anterior sheet has a broad, distinct articulation facet for the facial process of the maxilla and a lateral facet for the lacrimal, formed by a laterally extending flange (lacrimal flange of Bell et al. 2003). There is a small ridge on the dorsolateral surface near the base of the orbital process. The ventral process is thin with a slightly widened distal end. There is a distinct notch for the lacrimal foramen, and the ventral process forms the posterior and ventral border of the foramen. Medially, the boundary of the olfactory chamber is a smooth, rounded, and concave surface. Dorsal to the olfactory chamber there is a broad groove for articulation with the frontal. The orbitonasal flange extends far medially with a notch dorsally.

#### Identification

The fossil shares with teiids and gymnophthalmoids a distinct lacrimal flange (Bell et al. 2003) with an articulation facet for the facial process of the maxilla (Tedesco et al. 1999). Examined gymnophthalmoids differ in having a less extensive anterior process (facial sheet of Morales et al. 2019). The fossil is assigned to Teiidae.

### Postorbitofrontal

#### Description

TxVP 41229-27534 is a left postorbitofrontal missing the distal end of the anteromedial process (Fig. 44). The bone is slender and quadraradiate. The anteromedial process is slightly longer than the posteromedial processes. Between the medial processes, there is a facet for articulation with the frontal and parietal. The lateral margin is curved, and the posterolateral process is longer than the anterolateral process. The anterolateral process has a lateral facet for the jugal, and the posterolateral process has a lateral facet for articulation with the squamosal. There is a small ventral foramen on the central shaft of the bone.

#### Identification

The postfrontal and postorbital are fused into a single postorbitofrontal in some teiioids, xantusiids, some *Xenosaurus*, and some anguids (Estes et al. 1988; Evans 2008; Fig. 45). The fossil is quadraradiate as in some teiids and some *Xenosaurus*. The postorbitofrontal of *Xenosaurus* differs in having dorsal sculpturing and being much broader. The postfrontal and postorbital are separate in many gymnophthalmoids (Evans 2008). The postorbitofrontal is triradiate in examined gymnophthalmoids in which the postfrontal and postorbital are fused (Morales et al. 2019). The fossil is assigned to Teiidae.

**Figure 45.**
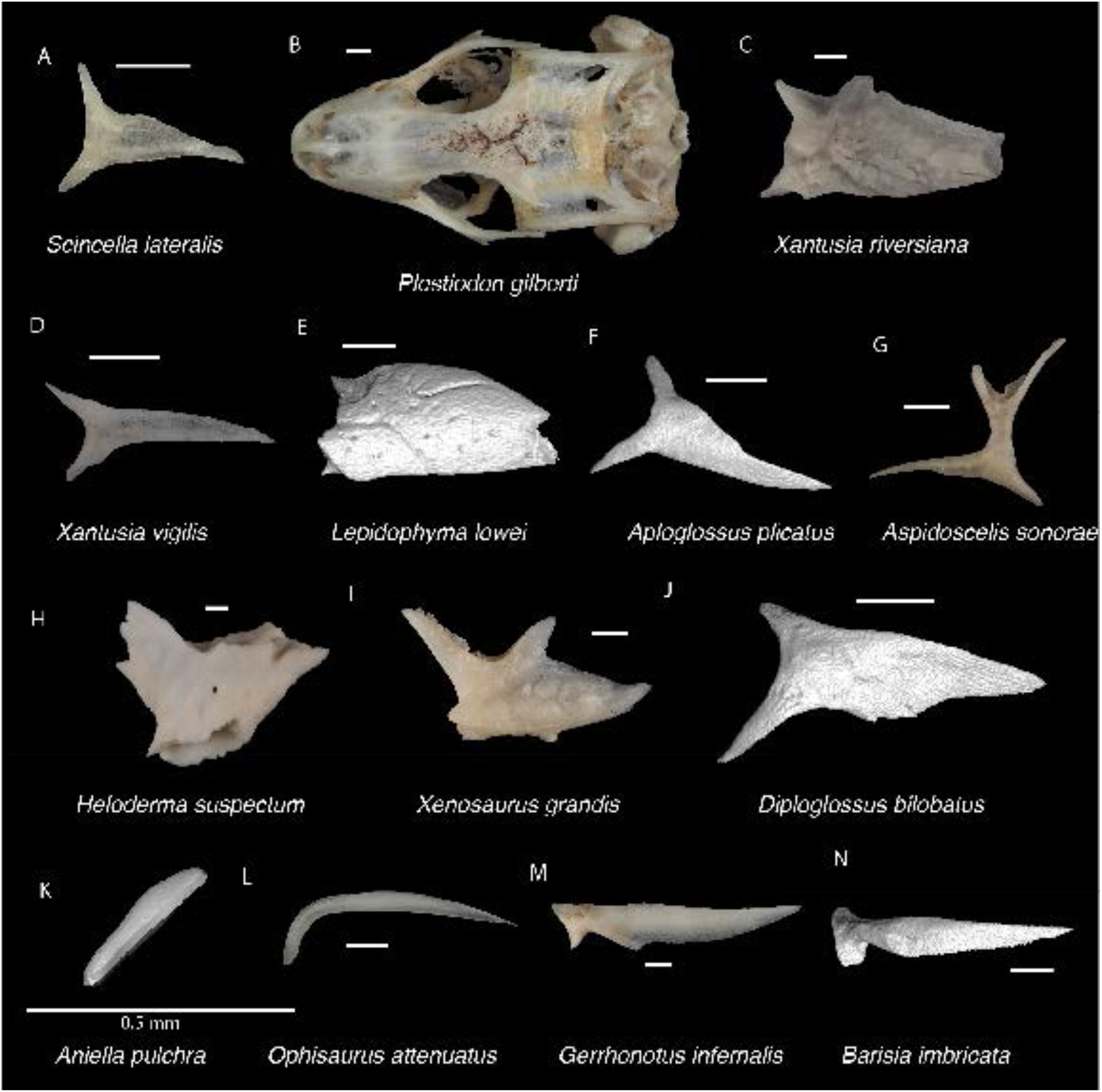
Non-Pleurodontan Postorbitals. Postorbitals and postorbitofrontals in dorsal and lateral views– **A**. *Scincella lateralis* TxVP M-4489; **B**. *Plestiodon gilberti* TxVP M-15662; **C**. *Xantusia riversiana* TxVP M-8505; **D**. *Xantusia virigatus* TxVP M-12130; **E**. *Lepidophyma lowei* LACM 143367; **F**. *Aploglossus plicatus* TNHC 34481; **G**. *Aspidoscelis sonorae* TxVP M-15670; **H**. *Heloderma suspectum* TxVP M-9001; **I**. *Xenosaurus grandis* TxVP M-8960; **J**. *Diploglossus bilobatus* TNHC 31933; **K**. *Anniella pulchra* FMNH 130479; **L**. *Ophisaurus attenuatus* TNHC 98712; **M**. *Gerrhonotus infernalis* TxVP M-13441; **N**. *Barisia imbricata* TNHC 76984. Scale bars = 1 mm.

### Pterygoid

#### Description

TxVP 41229-27524 is a left pterygoid (Fig. 44). The palatine process is thin with an anterior palatine facet. The transverse process projects laterally from the palatal plate and has a well-developed posterolateral corner and a triangular and anterolaterally pointed distal end. The transverse process has a deep ectopterygoid facet. The transverse process has no ridge on the ventral surface for insertion of the pterygomandibular muscle, but there is a dorsal ridge for insertion of the superficial pseudotemporal muscle. The quadrate process is elongated and has a tall, thin lateral ridge and medially directed lamina of bone. On the quadrate process, there is a deep fossa columella, but no postepipterygoid groove. There is no medial projection at the floor of the basipterygoid fossa. There is a ridge on the ventral surface of the palatal plate with four empty tooth positions.

#### Identification

Fossils share with teiioids the presence of pterygoid teeth (Estes et al. 1988) and an extensive, medially directed lamina of bone on the quadrate process (Du Bois 1943; Tedesco et al. 1999). Some gymnophthalmoids have a medially directed lamina of bone on the quadrate process (e.g., *Gymnophthalmus* and *Alopoglossus*), but the lamina does not extend as far medially as in the fossils (Evans 2008; Morales et al. 2019). *Calyptommatus* differs from the fossils in having the medial lamina on the quadrate process much more expanded (Roscito and Rodrigues 2010; Holovacs et al. 2020). *Alopoglossus* is further differentiated in having a posteromedial process along the medial edge of the pterygoid (Morales et al. 2019). The fossils are assigned to Teiidae.

### Dentary

#### Description

TxVP 41229-27389, TxVP 41229-27358, and TxVP 41229-28245 serve as the basis for our description (Fig. 44). TxVP 41229-27389 is a right dentary with 21 tooth positions. Teeth, except the three mesial-most teeth, are bicuspid with a smaller anterior cusp. There are substantial deposits of cementum at the tooth bases. The coronoid process is dorsally pointed. The angular process is broken and there is no surangular process. Anteriorly, the Meckelian groove opens ventrally, and near the middle of the tooth row the groove is briefly closed ventrally by the infra- and suprameckelian lips. Posteriorly, the Meckelian groove is tall. The suprameckelian lip is tall anteriorly. The intramandibular septum does not extend to the distal teeth. There is a narrow dental shelf. There is a distinct coronoid facet on the posterolateral surface, and there are five nutrient foramina arranged in a row on the lateral surface. TxVP 41229-27358 has eroded tooth crowns and differs from TxVP 41229-27389 in having 20 tooth positions, a small surangular process, and an entirely open Meckelian groove. TxVP 41229-28245 differs from TxVP 41229-27389 in having tricuspid distal teeth and an open Meckelian groove.

#### Identification

Fossils are assigned to Teiidae based on the presence of asymmetrically bicuspid distal teeth, a large amount of cementum deposits at base of teeth, a tall Meckelian groove for hypertrophied splenial, and a distinct intramandibular lamella (Estes et al. 1988; Nydam et al. 2007; Scarpetta 2020).

### Coronoid

#### Description

TxVP 41229-27699 is a left coronoid and serves as the basis for our description (Fig. 44). The coronoid process is tall and rounded, and slopes anteroventrally. There is a thin shelf of bone posterior to the coronoid process to articulate dorsally with the surangular. The anteromedial process is elongated with a medial splenial facet. The posteromedial process is oriented posteroventrally and has a wide notch (emargination of the adductor fossa *sensu* Evans 2008) along the posterior margin resulting in a distinct surangular process (*sensu* Evans 2008). There is a medial crest that extends from the coronoid process onto the posteromedial process. There is a large, anteriorly projecting lateral process with a dorsal dentary articulation facet. There is an anteroventrally oriented lateral crest that extends along the dorsal margin of the anterolateral process. There is a deep and narrow ventral groove for articulation with the surangular. There is a foramen on the anteromedial process.

#### Identification

Fossil coronoids differ from xantusiids and some pleurodontans in having an anterolateral process (Estes et al. 1988). The fossil differs from examined NA pleurodontans with an anterolateral process (see above) in having a more posteriorly deflected posteromedial process. Examined scincids and anguimorphs, except for *Heloderma* and *Ophisaurus attenuatus*, have a lateral process that does not extend as far anteriorly. *Heloderma* differs in having a shorter coronoid process with a vertically oriental lateral crest (Evans 2008) and examined *Ophisaurus* have a much smaller emargination of the adductor fossa, if present, along the posteromedial process. Examined geckos also lack a wide notch along the posterior margin of the posteromedial process (Kluge 1962; Daza et al. 2008; Bochaton et al. 2018). Examined gymnophthalmoids, except *Vanzosaura rubricauda* (Guerra and Montero 2009), differ in lacking a distinct surangular process (Bell et al. 2003; Evans 2008; Roscito and Rodrigues 2010; Morales et al. 2019). Examined gymnophthalmoids can be further distinguished in lacking a distinct anteroventrally oriented lateral crest. Fossils are assigned to Teiidae.

### Compound bone

#### Description

TxVP 41229-27339 is a right compound bone (Fig. 44). The anterior portion of the prearticular and much of the surangular are missing. The adductor fossa is widely open dorsally. The retroarticular process is narrow. There is a medially oriented and broad angular process with a low ridge. A sheet of bone connects the retroarticular, and angular processes and another sheet connects the angular process to the anterior prearticular portion of the compound bone. There is a medial crest (tympanic crest of McGuire 1996) that extends along the retroarticular process. There is a distinct coronoid articulation facet on the medial surface that extends ventral to the adductor fossa. There is an anteroventrally oriented crest on the lateral surface. There is a foramen on the angular process and a posterior surangular foramen on the lateral surface.

#### Identification

The fossil shares with teiioids the presence of a distinct angular process and a widely opened adductor fossa (Estes et al. 1988). Examined gymnophthalmoids, except *Calyptommatus* (Guerra and Montero 2009) and *Bachia* (Tarazona et al. 2008), differ in having a wider and more rounded retroarticular process (Bell et al. 2003; Evans 2008; Roscito and Rodrigues 2010; Morales et al. 2019). *Calyptommatus* and *Bachia* differ in having a smaller angular process (Guerra and Montero 2009; Tarazona et al. 2008). Fossils are assigned to Teiidae.

## Results

Our fossil identifications from Hall’s Cave resulted in a minimum of 11 lizard taxa, including five lizard taxa previously unknown from Hall’s Cave (Table 1). In most cases, we could not replicate the taxonomic level of previous fossil identifications. Our apomorphic framework permitted less taxonomically specific identifications most of the time. We recovered 17 separate lizard skull bones represented as fossils from Hall’s Cave. For most skeletal elements, identification could only be made to the family level; however, some elements (e.g., the premaxilla, maxilla, dentary, and jugal) permitted identification to the subfamily or genus level. Many of the skeletal elements that we recovered as fossils were relatively more robust skull bones (e.g., the frontal, parietal, dentary, maxilla, and compound bone), though we also recovered some smaller and more fragile elements (e.g., premaxilla, postfrontal, postorbital, jugal, and squamosal). Phrynosomatidae is the family with the most identified distinct taxa. This diversity in the fossil assemblage is recapitulated in the modern-day diversity with phrynosomatids being the most taxonomically diverse lizard family in the region. Most of the identified fossil lizard taxa inhabit the area around Hall’s Cave today, except for lizards within the *Phrynosoma douglasii* species complex (Table 2). Anguines (specifically *Ophisaurus attenuatus*) are reported to occur in Kerr County today; however, an examination of records of *Ophisaurus attenuatus* revealed only a single specimen (USNM 32826) in the county collected from three miles north of Kerrville in 1897. The next closest records of *Ophisaurus* are over 100 km east in Blanco and Hayes counties. There were a few extant lizard taxa that had records within 100 km of Hall’s Cave that we did not detect as fossils, including *Coleonyx brevis*, *Anolis carolinensis*, and *Phrynosoma modestum*. Possible *Phrynosoma modestum* fossils were previously reported from Hall’s Cave (Toomey 1993), yet, when we reexamined these fossils we found evidence arguing against assignment to *Phrynosoma modestum* (see Systematic Paleontology above).

**Table 2.** Identified fossil lizard taxa represented in Hall’s Cave from this study compared to native extant species within 100 km of Hall’s Cave.

| Family | Paleofauna at Hall's Cave identified in this study | Native extant species within 100 km of Hall's Cave (GBIF.org 2023) |
| --- | --- | --- |
| Eublepharidae |  | *Coleonyx brevis* |
| Dactyloidae |  | *Anolis carolinensis* |
| Crotaphytidae | Crotaphytidae | *Crotaphytus collaris* |
| Phrynosomatidae | Sceloporinae | *Sceloporus olivaceus*, *S. poinsettii*, *S. variabilis*, *S. consobrinus*, *Uta stansburiana* |
|  | *Urosaurus* | *Urosaurus ornatus* |
|  | Callisaurini | *Cophosaurus texanus*, *Holbrookia lacerata*, *H. propinqua* |
|  | *Phrynosoma* cf. *P. cornutum* | *Phrynosoma cornutum* |
|  | *Phrynosoma douglasii* species complex | None |
|  |  | *Phrynosoma modestum* |
| Anguidae | *Gerrhonotus* | *Gerrhonotus infernalis* |
|  | Anguinae | *Ophisaurus attenuatus* |
| Scincidae | *Scincella* | *Scincella lateralis* |
|  | Scincinae | *Plestiodon obsoletus*, *P. tetragrammus* |
| Teiidae | Teiidae | *Aspidoscelis gularis*, *A. marmoratus*, *A. inornatus*, *A. exsanguis*, *A. sexlineatus* |

## Discussion

An apomorphy-based fossil identification framework provides a replicable basis for employing the fossil record to understand the past. Such a solid foundation for fossil identifications is paramount for conducting larger evolutionary and ecological analyses using those data. Our study provides apomorphy-based identifications of fossil lizards from Hall’s Cave, and we add five new lizard taxa to the known diversity of the cave fauna. This work increases understanding of the past herpetofaunal diversity on the Edwards Plateau and sets the stage for further analyses examining past responses to environmental change. Additionally, since much of the previous work on Hall’s Cave focused on mammals (Toomey 1993) and plants (Cordova and Johnson 2019), our work on the fossil lizards adds a new dimension towards examining dynamics of the larger ecosystem in the region through time.

Many of our apomorphic identifications are at higher taxonomic levels (i.e., at the family or genus level) compared to previous efforts to identify fossil lizards from this locality, most of which yielded species-level identifications (Parmley 1988; Toomey 1993). Fossil identifications at the genus or family level were also achieved from other Pleistocene sites when using an apomorphic identification framework (Bell et al. 2004). Higher-level fossil identifications make taxon-based paleoecological reconstructions less precise (Bell et al. 2004). Nevertheless, it is important to recognize the limitations of our data and acknowledge that morphology is not always able to provide robust species-level fossil identifications without making assumptions about changes in geographic distributions. Identifying and accepting the limitations of our data will lead to more robust and well-supported evolutionary and paleoecological reconstructions. Identification of fossils using apomorphies generally results in identifications at higher taxonomic levels relative to identifications made from a phenetic framework or based on comparative samples restricted by modern distributions. However, alternative or supplemental methods can be used to refine fossil identifications further. For example, quantitative morphological methods like geometric morphometrics have become more commonly applied towards identification of fossil remains (Gray et al. 2017). In some cases, fossil identifications may be hindered due to substantial intraspecific morphological variation that overwhelms any interspecific signal. Furthermore, fossils are often disarticulated and broken skeletal elements, which hampers their identification. Paleogenomic and paleoproteomic methods are useful for identifying fossils, broken and intact, from more recent geological times, and often can provide species-level identification of fossil remains (Kehlmaier et al. 2017; Harvey et al. 2019). However, preservation of organic molecules differs between geographic regions and time periods, and paleogenomic data are not always recoverable from fossil specimens (Reed et al. 2003). Despite its limitations, morphology-based identification methods are applicable across the vastness of geologic time and continue to be an important approach for interpreting fossil remains.

Although higher-level fossil identifications using apomorphies can make paleoecological and paleoenvironmental interpretation less precise, it is still possible to glean insights from these identifications (e.g., Mead et al. 2008; Ramm et al. 2021; Scarpetta 2021). For example, here we used apomorphies to provide additional support for the presence of an extirpated clade of short-horned lizards (*Phrynosoma douglasii* species complex) on the Edwards Plateau during the late Pleistocene to the early Holocene. The closest living species in the *Phrynosoma douglasii* species complex to Hall’s Cave, *P. hernandesi*, is found 435 km west in Jefferson County, Texas. However, there is a closer preserved specimen of *P. hernandesi* (MCZ R-8216) 285 km northwest in Mitchell County, Texas, collected in 1997. Toomey (1993) postulated that the extirpation of lizards in the *P. douglasii* species complex near Hall’s Cave may have been driven by complex ecological interactions, including climatic changes and potential changes in resource availability. Additional study, perhaps using stable isotope data, will shed more light on the cause of this extirpation.

There are a few extant lizard taxa (e.g., *Coleonyx brevis*, *Anolis carolinensis*) that live within 100 km of Hall’s Cave today that we did not detect as fossils. The absence of these taxa from Hall’s Cave may represent a true absence of these taxa from the area around Hall’s Cave. Alternatively, preservation or accumulation biases, particularly against the small and delicate bones of smaller taxa, like *Coleonyx brevis*, may also explain these absences. Although we found evidence contradicting the previous identifications of some fossils to *Phrynosoma modestum*, examination of additional material may confirm this taxon in the Hall’s Cave sequence. The absence of *Anolis* from the Hall’s Cave sequence is interesting because some skull elements of *Anolis* (e.g., dentary, parietal, frontal, maxilla) are relatively robust, easily recognizable, and commonly preserved as fossils in other Quaternary deposits (Auffenberg 1956). Although *Anolis carolinensis* is widespread in Texas today, including the areas around Hall’s Cave, there is a striking lack of Pleistocene fossil anoles in Texas (Holman 1969, 1995). Fossil *Anolis* are known from at least the late Oligocene of Florida (Chovanec 2014) and *Anolis carolinensis* was estimated to have experienced range expansion out of Florida during the Pleistocene (Campbell-Staton et al. 2012). An eastward expansion of the species range from Florida was estimated to have started around 300,000 years ago (Tollis and Boissinot 2014). A lack of fossil *Anolis* from the Pleistocene of Texas may indicate that they did not reach Texas until more recently; however, increased sampling of fossil sites, especially in east Texas, is needed to understand the dispersal pattern. Although our study represents only a single locality, it preserves evidence that for at least some lizard taxa, stability was not the rule during parts of the Quaternary. Additional study in other fossil localities across NA is necessary to form a more complete synthesis of herpetofaunal dynamics during the Quaternary.

A main goal of this work is to facilitate the identification of North American lizard fossils in a phylogenetically explicit context. Our hope is that researchers familiar with the lizard skeletal system will find our synthesis of previously reported and new apomorphies useful for identifying fossil lizard remains and will spur new investigation into patterns of morphological variation within the lizard skeletal system. Our figures are intended to showcase cranial osteological variation in North American lizards with diverse specimen and taxon sampling. Although we obtained a broad comparative sample, we were unable to obtain comparative specimens for all North American lizard families, genera, and species, especially those for which CT or dry-skeletal material is scarce or non-existent. Broadly, there is a paucity of information in the published literature on morphological variation in many vertebrate clades, including lizards (Evans 2008). An incomplete understanding of patterns of variation hinders our ability to make well-substantiated claims using the fossil record. Modern comparative morphological data provide a basis by which to interpret the fossil record (Bell et al. 2010), particularly for fossils from the more recent past (i.e., the Quaternary). Continued investigations into patterns of morphological variation in modern taxa are necessary for understanding patterns encountered in the fossil record (Bell et al. 2010), including interspecific variation and intraspecific variation (e.g., sexual dimorphism, ontogenetic variation, etc.).

Many of the fossils examined here were found in vials of fossils labeled “scrap bone.” These “scrap” fossils were often elements other than the commonly described upper and lower tooth bearing elements that are commonly described in the Quaternary paleoherpetological literature (Bell et al. 2010). It is important that these additional elements are recognized because, as we have shown here, there are apomorphies on various skeletal elements that are as useful or more useful for fossil identification (Smith 2009a, 2011). Increased recognition of lizard remains can lead to novel insights and discoveries related to NA lizards as well as a more holistic view of ancient faunal assemblages. We therefore specifically worked to include images of mainly disarticulated skeletal elements, as would likely be encountered in the fossil record, from a diverse set of NA lizards. This will be especially useful for researchers who do not specialize on lizards, and it is our hope that the images we provide here lead to an increased recognition and identification of lizard remains in fossil deposits.

## Notes

### Competing Interest Statement

The authors have declared no competing interest.

