## Supplemental Table 2 for "Identification of Late Pleistocene and Holocene fossil lizards from Hall’s Cave and a primer on morphological variation in North American lizard skulls"

*Denotes clades exhibiting known exceptions for an apomorphic feature

| Premaxilla |  |  |
| --- | --- | --- |
| **Apomorphy** | **Character evolution**  **hypothesis** | **References** |
| Fused premaxilla | Apomorphy of Squamata, lost in Scincidae and Gekkota | Gauthier et al. 1988 |
| Less than seven tooth positions on the premaxilla | Apomorphy of Iguania* | Smith 2009a |
| Incisive process absent | Apomorphy of Laterata* | Gao and Norell 1998; Conrad 2008; Gans and Montero 2008; Morales et al. 2019; Villa and Delfino 2020 |
| Deeply incised palatal plate | Apomorphy of Teiioidea* |  |
| Enlarged median tooth on the premaxilla | Apomorphy of Amphisbaenia | Smith 2009b |
| Anterior premaxillary foramina (across Squamata) | Apomorphy of Iguania* independently evolved in Anguimorpha*, Cordyloidea, and Amphisbaenia | Smith 2009b; Scarpetta 2019 |
| Multicuspid premaxillary teeth | Apomorphy of Iguanidae* within Iguania | de Queiroz 1987; Scarpetta 2019 |
| Nasal facets anteriorly on the nasal process | Apomorphy of Iguanidae* within Pleurodonta, independently evolved in Crotaphytidae and the clade (Liolaemidae, (Polychrotidae, (Hoplocercidae, (Opluridae, Leiosaurudae))))* | Adapted from Etheridge and de Queiroz 1988; Daza et al. 2012 |
| Anterior premaxillary foramina (within Phrynosomatidae) | Apomorphy of Phrynosomatinae within Phrynosomatidae and also occurs in *Uta** | Scarpetta 2019 |
| Anterior rostral face flat and flush with nasal process | Apomorphy of Phrynosomatinae within Phrynosomtidae | Scarpetta 2019 |
| Rectangular alveolar plate | Apomorphy of sand-lizard clade within Phrynosomatidae | Scarpetta 2019 |
| Nasal process directed dorsally | Apomorphy of *Phrynosoma* within Phrynosomtidae | Scarpetta 2019 |
| Base of nasal process nearly the same width and alveolar plate | Apomorphy of *Phrynosoma* within Phrynosomtidae | Scarpetta 2019 |
| Forked palatal process | Apomorphy of Anguinae and Diploglossinae within Anguimorpha | Meszoely 1970; Evans 2008; Conrad et al. 2011; Scarpetta 2018 |
| Dorsal ossification on the alveolar plate posterior to medial ethmoidal foramen | Apomorphy of Diploglossinae independently evolved in Gerrhonotinae within Anguidae | Scarpetta 2018 |
| Ossified bridge extending laterally from nasal process that encloses the medial ethmoidal foramen | Apomorphy of Gerrhonotinae* within Anguidae | Scarpetta et al. 2021 |

| Maxilla |  |  |
| --- | --- | --- |
| **Apomorphy** | **Character evolution hypothesis** | **References** |
| Elongate depression on the palatal shelf (gutter) encompassing the superior alveolar nerve and maxillary artery | Apomorphy of Iguania* | Smith 2006, 2009a |
| Foramina for both the subnarial artery and the anterior inferior alveolar nerve on the premaxillary process | Apomorphy of Pleurodonta* | Smith 2009a |
| Deep jugal groove on the postorbital process | Apomorphy of the clade (Crotaphytidae, (Corytophanidae, Leiocephalidae)) independently evolved in Iguanidae and Opluridae within Pleurodonta | Smith 2009a; Scarpetta 2021 |
| Large sub-triangular palatine process | Apomorphy of Phrynosomatidae also present in Tropiduridae, Crotaphytidae, Leiocephalidae*, and Opluridae | Smith 2009a; Scarpetta 2021 |
| Medially folded facial process with distinct canthal crest and anterodorsal facing surface | Apomorphy of Phrynosomatidae* also present in Tropiduridae and *Anolis* within Pleurondonta | Smith 2009a |
| Narrow, triangular facial process | Apomorphy of *Phrynosoma** within Pleurodonta | Bell et al. 2004; Scarpetta 2021 |
| Strongly asymmetric palatine process | Apomorphy of *Phrynosoma** within Pleurodonta | Smith 2009a |
| Tall, prominent crista transversalis | Apomorphy of *Phrynosoma** within Phrynosomatidae | Scarpetta 2021 |
| Elongate premaxillary process with a low, anteromedially trending crista transversalis | Apomorphy of Gerrhonotinae within Anguidae | Smith 2009b |
| Deeply notch premaxillary process | Apomorphy of Diploglosinae also present in Anguinae within Anguidae | Meszoely 1970 |

| Nasal |  |  |
| --- | --- | --- |
| **Apomorphy** | **Character evolution hypothesis** | **References** |
| Distinct supranarial process | Apomorphy of Iguania* | Gauthier et al. 2012 |

| Frontal |  |  |
| --- | --- | --- |
| **Apomorphy** | **Character evolution hypothesis** | **References** |
| Fused frontal | Apomorphy of Gekkota independently evolved in Teiioidea, Iguania, and some members of Anguimorpha* and scincidae* | Estes et al. 1988; Greer 1970 |
| Strongly waisted interorbital margins of the frontal | Apomorphy of Pleurodonta* independently evolved in some xenosaurs, gerrhonotines, and members of Teiioidea* | Estes et al. 1988; Evans 2008; Morales et al. 2019 |
| Reduced descending cristae cranii of the frontal | Apomorphy of Iguania independently evolved in Teiioidea* | Preach 1980; Estes et al. 1988; Evans 2008 |
| Parietal foramen partially or fully within the frontal | Apomorphy of Iguania* | Etheridge 1959; Etheridge 1969; de Quieroz 1987; Smith 2011 |
| Co-ossified osteoderms on dorsal surface | Apomorphy of Anguimorpha*  independently evolved in Scincomorpha* and Lacertidae* | Estes et al. 1988; Evans 2008 |
| Frontal lappets | Apomorphy of Gymnphthalmoidea independently evolved in Chamaeleonidae | Estes et al. 1988 |

| Parietal |  |  |
| --- | --- | --- |
| **Apomorphy** | **Character evolution hypothesis** | **References** |
| Fused parietal | Apomorphy of Squamata* | Estes et al. 1988; Gauthier et al. 1988 |
| Long ventrally projecting parietal crests or processes | Apomorphy of Scincomorpha* independently evolved in Teiioidea* and Anguimorpha | Estes et al. 1988; Evans 2008; Ledesma and Scarpetta 2018 |
| Long posterior projections (median extensions of Evans 2008) on the posterior edge of parietal table between the postparietal processes | Apomorphy of Scincomorpha* | Evans 2008; Gauthier et al. 2012 |
| Co-ossified osteoderms on dorsal surface | Apomorphy of Anguimorpha*  independently evolved in Scincidae* | Estes et al. 1988; Evans 2008 |
| Parietal foramen not fully enclosed by the parietal | Apomorphy of Gekkota+Dibamidae independently evolved in Amphisbaenia*, Teiioidea*, Helodermatidae, and Pleurodonta* | Estes et al. 1988 |
| Parietal lappets | Apomorphy of Lacertoidea* independently evolved in Xantusiidae and some iguanians, cordylids, and scincids | Estes et al. 1988 |

| Prefrontal |  |  |
| --- | --- | --- |
| **Apomorphy** | **Character evolution hypothesis** | **References** |
| Prefrontal boss | Apomorphy of Iguania* also present in Teiinae | Estes et al. 1988; Gauthier et al. 2012; Smith 2009b |
| Well developed lateral projection buttressing the lacrimal | Independently evolved apomorphy of Teiioidea, Corytophanidae, and Scincidae | Bell et al. 2003; Smith 2009a |
| Lacrimal fused to prefrontal | Independently evolved apomorphy of Xantusiidae, Gymnopthamoidea*, and Amphisbaenia | Savage; 1963; Presch 1980; Estes et al. 1988; Evans 2008 |

| Jugal |  |  |
| --- | --- | --- |
| **Apomorphy** | **Character evolution hypothesis** | **References** |
| Quadratojugal process absent | Apomorphy of Iguania* | Gauthier et al. 1988; Smith 2009a |
| Angulated jugal | Apomorphy of Unidentata* (sensu Burbrink et al. 2020), lost in Iguania* | Conrad 2008 |
| Medial ectopterygoid process | Apomorphy of Laterata* | Evans 2008 |
| Medial ridge located at the midline of the suborbital and postorbital processes | Apomorphy of Anguimorpha* | Cernansky et al. 2014 |
| Posteriorly deflected distal end of the postorbital process | Apomorphy of Phrynosomatidae independently evolved in Tropiduridae, Crotaphytidae, and Opluridae within Pleurodonta | Smith 2009a |
| Wide postorbital process in lateral view | Apomorphy of Xenosauridae independently evolved in some iguanians, and Xantusiidae | Savage 1963; Smith 2009a; Gauthier et al. 2012 |
| Long quadratojugal process | Apomorphy of Gerrhonotinae* independently evolved in some diploglossines within Anguidae | Rieppel 1980; Ledesma and Scarpetta 2021 |

| Postfrontal |  |  |
| --- | --- | --- |
| **Apomorphy** | **Character evolution hypothesis** | **References** |
| Not elongated in mediolateral or anteroposterior plane | Apomorphy of Iguania* | Conrad 2008; Smith 2009a |

| Postorbital |  |  |
| --- | --- | --- |
| **Apomorphy** | **Character evolution hypothesis** | **References** |
| Postorbital forms more than half of orbital border and has a distinct ventral process | Apomorphy of Iguania* | Estes et al. 1988 |
| Dorsal facet for frontoparietal articulation on dorsal process | Apomorphy of Crotaphytidae also occurs in Corytophanidae, Polychrotidae, Dactyloidae*, and Leiosauridae within Iguania | Adapted from Smith 2009a |
| Lateral tubercle mid-height on the postorbital | Apomorphy of Crotaphytidae independently evolved in Tropiduridae and Opliuridae* within Pleurodonta | Smith 2009a |
| Convex dorsal margin of posterior process | Apomorphy of Corytophanidae independently evolved in some oplurids, *Leiocephalus*, and agamids within Iguania | Smith 2009a |
| Anteriorly projecting supraorbital spine | Apomorphy of Corytophanidae independently evolved in *Phrynosoma* | Smith 2009a |
| Fused postorbitofrontal | Apomorphy of Xantusiidae independently evolved in Xenosauridae*, Lacertoidea*, some anguimorphs*, and some scincids* | Estes et al. 1988; Conrad 2008 |

| Squamosal |  |  |
| --- | --- | --- |
| **Apomorphy** | **Character evolution hypothesis** | **References** |
| Posterodorsal process | Apomorphy of Iguania* independently evolved in some clades within Teiidae and Xenosauridae | Estes et al. 1988; Tedesco et al. 1999; Evans 2008; Smith 2009a |

| Quadrate |  |  |
| --- | --- | --- |
| **Apomorphy** | **Character evolution hypothesis** | **References** |
| Notch or foramen on dorsal surface for articulation with the squamosal | Apomorphy of Squamata | Gauthier et al. 1988 |
| Distinct pterygoid lappet present | Apomorphy of Lacertoidea* and independently evolved in Helodermatidae, Xenosauridae, Scincidae*, and Iguania* | Estes et al. 1988; Evans 2008 |
| Low ridge on the posterior surface of the conch | Apomorphy of Corytophanidae independently evolved in some members of Hoplocercidae, Dactyloidae, Polychrotidae, Phrynosomatidae, and Agamidae within Iguania | Smith 2009a |

| Pterygoid |  |  |
| --- | --- | --- |
| **Apomorphy** | **Character evolution hypothesis** | **References** |
| Absence of pterygoid tooth loci | Apomorphy of Gekkota+Dibamidae, independently evolved in Xantusiidae, Amphisbaenia, and some clades within Anguimorpha* and Iguania* | Estes et al. 1988 |
| Well-developed ventromedial projection at the floor of the basipterygoid fossa | Apomorphy of Iguania* independently evolved in some clades within Anguimorpha*  –alternatively an apomorphy of Toxicodera lost in some clades | Evans 2008; Smith 2009b |

| Ectopterygoid |  |  |
| --- | --- | --- |
| **Apomorphy** | **Character evolution hypothesis** | **References** |
| Elongate posterolateral process | Apomorphy of Iguania independently evolved in Xantusiidae and Xenosauridae  –alternatively a plesiomorphy of Squamata retained in these clades | Smith 2009b |
| Broad ectopterygoid | Apomorhy of Teiidae* independently evolved in Xantusiidae*, Amphisbaenia, and Dibamidae | Gauthier et al. 2012 |

| Marginal Dentition |  |  |
| --- | --- | --- |
| **Apomorphy** | **Character evolution hypothesis** | **References** |
| Pleurodont teeth (i.e. tooth position lingual relative to labial wall of tooth bearing bone) | Apomorphy of Lepidosauramorpha* | Simões et al. 2018 |
| Tricuspid teeth | Apomorphy of Pleurodonta independently evolved within Xantusiidae and Teiioidea* | Savage 1963; Tedesco et al. 1999; Bell et al. 2003; Scarpetta 2021 |
| Asymmetrically bicuspid distal teeth | Apomorphy of Lacertoidea* | Scarpetta 2020 |
| Large amount of cementum deposits at base of teeth | Apomorphy of Teiidae | Estes et al. 1988; Nydam et al. 2007; Scarpetta 2020 |
| Unicuspid teeth with striated crowns | Apomorphy of Scincomorpha independently evolved within Anguimorpha | Estes 1963; Smith 2009b |
| Tooth bases much wider than the crowns | Apomorphy of Dactyloidae independently evolved in Crotaphytidae within Pleurodonta | Etheridge and de Queiroz 1988; Smith 2009b |

| Dentary |  |  |
| --- | --- | --- |
| **Apomorphy** | **Character evolution hypothesis** | **References** |
| Meckelian groove open ventrally, anterior to the inferior alveolar foramen | Apomorphy of Scincidae* and independently evolved within Teiioidea*, Anguimorpha, and some chamaeleonids  –alternatively an apomorphy of Unidentata lost in Iguania* and Cordyloidea* | Nash 1970; Estes et al. 1988; Lang 1991; Evans 2008; Morales et al. 2019; Scarpetta 2020 |
| Inframeckelian lip curls dorsolingually, producing a medial exposure of the Meckelian groove along the mid-length of the dentary | Apomorphy of Iguania* and independently evolved within some teiids and Cordyloidea* | Lang 1991; Evans 2008; Gauthier et al. 2012; Bochaton et al. 2017 |
| Suprameckelian and inframeckelian lips constrict or close Meckelian groove | Apomorphy of Iguania* and independently evolved within some scincids and teiids | Greer 1974; Gauthier et al. 2012; Bochaton et al. 2017; Scarpetta 2021 |
| Suprameckelian and inframeckelian lips indistinguishably fused, enclosing the Meckelian groove | Apomorphy of Gekkota+Dibamidae independently evolved in Amphisbaenians, Xantusiidae, Gymnophthalmoidea*, Pleurodonta, and some scincids  Within Pleurodonta apomorphy of Iguanidae, Dactyloidae, Tropiduridae, Leiocephalidae, Polychrotidae, Leiosaurudae*, and some members of Opluridae, Corytophanidae, and Phrynosomatidae | Greer 1974; Mead et al. 1985; Estes et al. 1988; Evans 2008; Smith 2011; Gauthier et al. 2012; Scarpetta 2021 |
| Broad subdental shelf | Apomorphy of Gekkota+Dibamidae, and independently evolved in Scincomorpha, and Lacertoidea*  –alternatively small subdental shelf is an apomorphy of Toxicofera | Estes et al. 1988 |
| Surangular notch or presence of surangular process | Apomorphy of Anguimorpha* | Gauthier 1982; Estes et al. 1988; Good 1988 |
| Posteriorly extended intramandibular septum near the posterior end of tooth row | Apomorphy of Anguimorpha independently evolved in some clades within Iguania* | Pregill 1981; Estes et al. 1988 |
| Intramandibular septum with free posteroventral margin | Apomorphy of Anguidae* within Anguimorpha | Gauthier 1982; Conrad et al. 2011 |
| Intramandibular septum fused to posterior inner wall of dentary | Apomorphy of Anguinae, also occurs in Diploglossinae within Anguidae | Pregil 1981; Klembara et al. 2014; Syromyatnikova and Aranda 2022 |
| Surangular spine | Apomorphy of *Ophisaurus* within Anguidae | Klembara et al. 2014 |
| Splenial spine | Apomorphy of Anguimorpha* | Conrad 2008; Čerňanský and Augé 2019 |
| Distinct intramandibular lamella | Apomorphy of Teiidae* independently evolved in some clades within Pleurodonta* | Denton and O’Neill 1995; Smith 2009a; Scarpetta 2020 |
| Tall Meckelian groove for hypertrophied splenial | Apomorphy of Teiidae | Scarpetta 2020 |

| Coronoid |  |  |
| --- | --- | --- |
| **Apomorphy** | **Character evolution** | **References** |
| Distinct coronoid process formed only by coronoid | Apomorphy of Squamata | Gauthier et al. 1988 |
| Distinct anteriorly projecting lateral process | Apomorphy of Squamata, lost in Xantusiidae and some clades within Iguania* | Estes et al. 1988 |

| Splenial |  |  |
| --- | --- | --- |
| **Apomorphy** | **Character evolution** | **References** |
| Anterior inferior foramen enclosed by splenial | Apomorphy of Teiioidea* independently evolved in some clades within Iguania*, in Scincomorpha*, and some clades within Anguimorpha* | Evans 2008; Gauthier et al. 2012; Morales et al. 2019 |
| Anterior inferior alveolar foramen posterodorsal to anterior mylohyoid foramen | Apomorphy of Teiioidea* | Gauthier et al. 2012 |
| Splenial fused to dentary | Apomorphy of Xantusiidae | Gautheir et al. 2012 |

| Compound bone |  |  |
| --- | --- | --- |
| **Apomorphy** | **Character evolution hypothesis** | **References** |
| Condyle formed only by articular | Apomorphy of Lepidosauria | Gauthier et al. 1988 |
| Medial inflection of retroarticular process | Apomorphy of Scincidae independently evolved in Anguimorpha and Gekkota | Estes et al. 1988 |
| Lack of oblique torsion of retroarticular process | Apomorphy of Lactertoidea independently evolved in Pygopodidae and Iguania* | Estes et al. 1988 |
| Tubercle or flange on posteromedial margin of retroarticular process | Apomorphy of Scincidae | Estes et al. 1988 |
| Notch on the medial margin of the retroarticular process | Apomorphy of Gekkota | Estes et al. 1988 |
| Distinct angular process | Apomorphy of Iguania* and independently evolved in Teiioidea | Estes et al. 1988; Smith 2009a |
| Anterior surangular foramen ventrally located on surangular | Apomorphy of Acrodonta independently evolved in Hoplocercidae and some members of Iguanidae | de Queiroz 1987; Frost and de Queiroz 1988; Smith 2009a |
| Broadened retroarticular process | Apomorphy of Anguimorpha independently evolved in Gekkota and Scincomorpha* | Estes et al. 1988 |
| Widely open adductor fossa | Apomorphy of Teiioidea independently evolved in some clades within Scincidae | Estes et al. 1988 |
| Prearticular crest present | Apomorphy of Teiioidea independently evolved in Xantusiidae | Gauthier et al. 2012 |
